# Harnessing P450 enzyme plasticity for generalizable peptide biaryl macrocyclization

**DOI:** 10.1101/2024.10.06.616194

**Authors:** Runze Liu, Cuiping Pang, Gengfan Wu, Bei-Bei He, Bo Huang, Hongyu Ren, Zhuanglin Shen, Jing Liu, Zhi-Man Song, Jiahai Zhou, Yong-Xin Li

## Abstract

Macrocyclic peptides containing biaryl motifs, predominantly derived from natural products, are valuable scaffolds due to their structural rigidity and potent bioactivity. However, current synthetic methods remain constrained by the absence of broadly applicable strategies across chemocatalytic and biocatalytic platforms. Here, we discover a versatile P450 enzyme, GpeC, capable of facilitating oxidative C-C/O/N cross-coupling for peptide biaryl macrocyclization (PBC) with ‘leader-independent’ activity. Crystal structure analysis of GpeC rationalizes leader-independent property and reveals ‘adaptive recognition’ that enables access to diverse biaryl-linked macrocycles. GpeC exhibits exceptional substrate promiscuity, accommodating 19 of 20 canonical amino acids and diverse noncanonical analogs within a minimal tetrapeptide scaffold. The efficient semi-synthesis of the natural product Rubrin further demonstrated the versatility of GpeC. We further introduced ‘lytic to tetraregion’ (LTT), a one-step, single enzyme synthesis for modular synthesis of biaryl-cyclized tetrapeptide. Overall, GpeC’s robustness and programmability position it as a broadly applicable biocatalyst for the synthesis of biaryl macrocyclic peptides.

## Main

Biaryls constitute a structurally constrained motif of profound significance in the design of bioactive molecules and functional materials^1,2^. Among these, biaryl-crosslinked macrocyclic peptides stand out due to their precisely rigid three-dimensional architectures, which confer exceptional stability and pharmacological potential. The macrocyclization of peptides via biaryl linkages enhances proteolytic stability, cellular permeability, and bioactive conformation stabilization, thereby imbuing these molecules with drug-like properties^3-6^. Notable examples include the clinically pivotal antibiotic vancomycin and arylomycin analog G0775, both of which exemplify the therapeutic relevance of this structural class (Fig. 1A)^7,8^. The potent bioactivity of biaryl-cyclized macrocyclic peptides has prompted the development of the modular synthesis of such scaffolds^6,9^. Retrosynthesis of such compounds remains case-specific, as they are largely governed by the intrinsic steric and electronic properties of the substrate side chains^9-14^. Achieving broadly generalizable peptide biaryl cyclization remains a persistent challenge.

**Figure 1.**
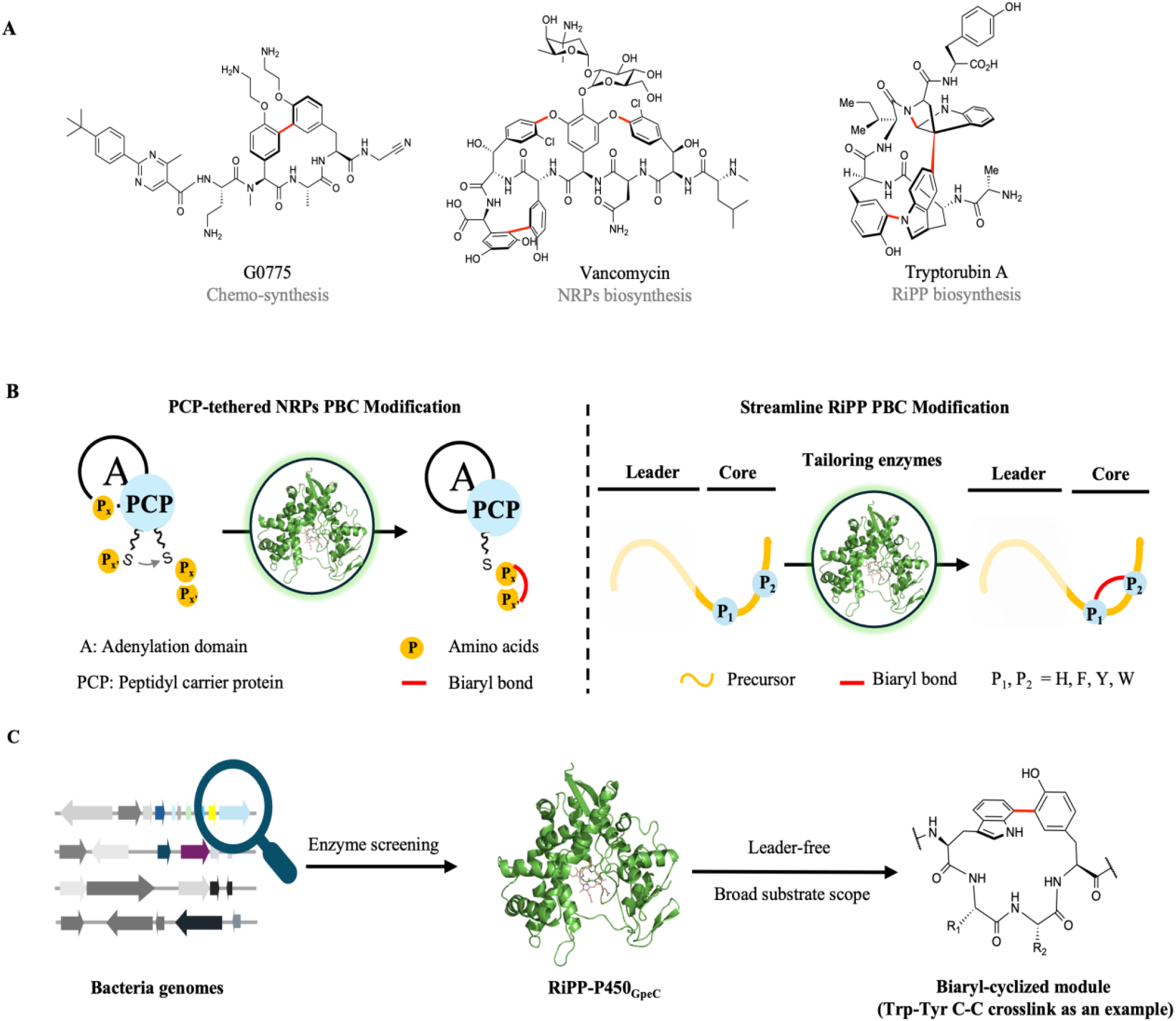
Representative biaryl crosslinked macrocyclic peptide and their synthetic strategies. A) Representative biaryl crosslinked macrocyclic peptides as potent antibiotics. B) Contrasting enzymatic strategies for peptide biaryl cyclization (PBC). Left: Non-ribosomal peptide synthetase (NRPS)-dependent PBC requiring peptidyl carrier protein (PCP)-tethered substrates. Right: Ribosomally synthesized and post-translationally modified peptide (RiPP)-associated PBC enabled by substrate-adaptive recognition. C) This study: P450 GpeC catalyzes leader-free biaryl cyclization under mild conditions with broad substrate tolerance.

This unmet challenge finds intriguing parallels in nature’s enzymatic machinery, where cytochrome P450s have evolved as precision biocatalysts for complex molecular synthesis^15-19^. These heme-containing oxidases leverage radical-mediated oxidative machinery to catalyze diverse reactions, particularly notable for their demonstrated capacity for peptide biaryl cyclization (PBC). While the catalytic ability of PBC-P450s is evident from their biosynthesis of complex biaryl natural products, their adaptation as a programmable biocatalytic toolbox remains constrained. Current characterization efforts have predominantly focused on PBC-P450s associated with non-ribosomal peptide synthetase (NRPS) pathways, where enzymatic activity strictly depends on peptidyl carrier protein (PCP)-tethered substrates (Fig. 1B)^7,20,21^. This fundamental limitation starkly contrasts with the emerging biosynthetic enzymes of ribosomal synthesis post-translational modification peptide (RiPP)^22-24^, whose concise biosynthetic logic and substrate plasticity address the ‘generalizability bottleneck’ (Fig. 1A, B)^6,25^. The expanding repertoire of RiPP PBC-P450s demonstrates remarkable catalytic potential, mediating inter-residue couplings across aromatic residues (Trp, Tyr, Phe, His) in microbial hosts^26-32^; however, it remains underdeveloped for enzymatic applications. P450_blt_, the sole member biochemically characterized *in vitro*, exhibits tricyclic specificity, especially on the stereo-hindered residues^27,33^. Motivated by RiPP P450s’ unparalleled catalytic ability for peptide C-C, C-O, and C-N biaryl cross-coupling reactions, we’re interested in discovering new RiPP PBC-P450s to establish biocatalytic strategies, aiming to achieve a broadly applicable method for PBC.

In this study, via genome mining, we identified a robust RiPP-P450, GpeC, that catalyzes tetrapeptide biaryl macrocyclization. Strikingly, GpeC overcomes the substrate specificity constraints inherent to both transitional metal-catalyzed peptide biarylation and NRPS-dependent P450s, facilitating efficient cyclization of tetrapeptides containing diverse aromatic pairings Trp-Tyr [C-C], Tyr-Tyr [C-O-C], and Tyr-Trp [C-C/C-N] (Fig. 1C). Structural and computational analyses reveal GpeC’s substrate-binding plasticity by the ‘adaptive recognition’ mechanism that enables it to accommodate both canonical and noncanonical residues to facilitate generalizable PBC. We demonstrate the versatility of GpeC by semi-synthesis of the natural product Rubrin. Taking advantage of this adaptability, we formulated a one-step, single-enzyme synthesis called ‘lytic to tetraregion’ (LTT) for the modular synthesis of biaryl-cyclized tetrapeptides. GpeC’s leader peptide independence, wide substrate tolerance, and versatility position it as an enzymatic toolbox for broadly generalizable peptide biaryl cyclization. These advances bridge a critical gap in the efficient and sustainable synthesis of biaryl cyclic peptides, offering a straightforward platform for rapidly generating drug-like biaryl macrocycles for pharmaceutical discovery.

## Results

### Genomics-guided discovery of P450 peptide biaryl macrocyclase GpeC

We have previously established a genome mining workflow called ‘Short Peptide Enzyme Co-Occurrence’ (SPECO) analysis, which effectively identifies RiPP biosynthetic gene clusters (BGCs) without relying on sequence similarity to the known RiPPs^34^. Fascinated by the catalytic ability of biaryl cyclization by RiPP PBC-P450s^28-30,35^, we assume their role in transformative biocatalysts was largely unexplored. To address this, we applied SPECO to systematically analyze 10,228 small peptide-P450 pairs across bacterial genomes (Fig. 2A, Supplementary Figs. S1, 2), and four RiPP-associated P450s were selected for *in vitro* functional characterization. The expression condition for the selected P450s in *E. coli* was optimized to overcome issues related to solubility and heme incorporation to procure functional enzymes (Supplementary Fig. S3). This led us to identify GpeC, which demonstrated *in vitro* enzymatic activity (Fig. 2C). The biosynthesis of the *gpe* gene cluster consists of 2 hypothetical precursor peptides, GpeA_1_ (35-mer) and GpeA_2_ (27-mer) with a typical ‘WxxY’ core sequence (GpeA_1_ as ‘WVMY’, GpeA_2_ as ‘WMLY’), and we proposed them as the native substrates for GpeC (Fig. 2B). Catalytic competence of GpeC was confirmed via the CO difference spectrum assay (Supplementary Fig. S4). The *in vitro* assay of GpeC with GpeA_2_ (**2**), involving NADPH and spinach-oriented ferredoxin-ferredoxin reductase pairs, produced a catalytic product after 5 hours at 28°C (Fig. 2C). UPLC-HRMS analysis revealed GpeC catalyzed the macrocyclization of the native precursor *in vitro*, with the modification localized to the ‘WMLY’ motif as supported by tandem MS analysis (Fig. 2D, Supplementary Fig. S5).

**Figure 2.**
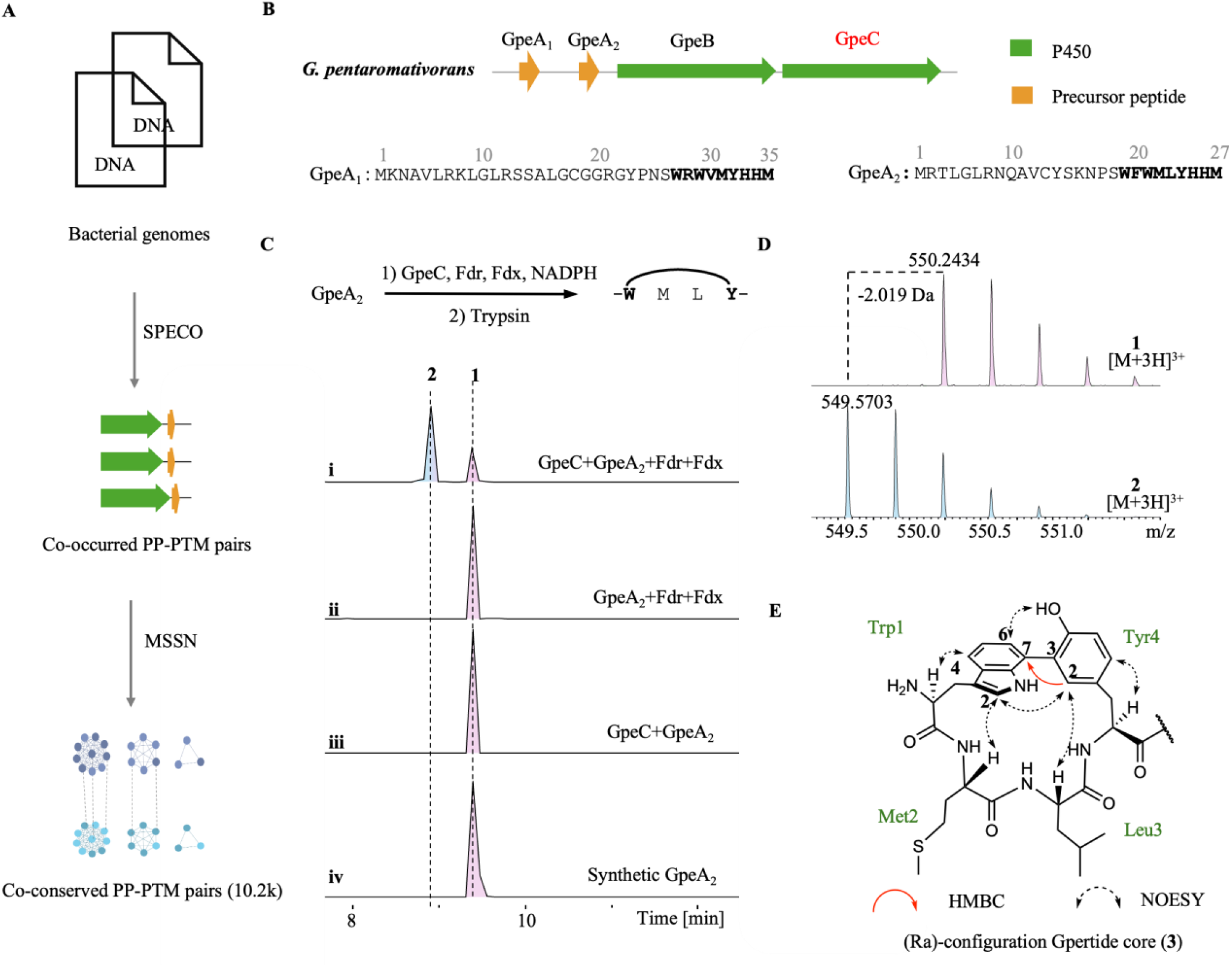
*In vitro* assay of GpeC and structure elucidation of Gpertide. A) General genome mining workflow of RiPP-P450 enzymes. PP, short for precursor peptide; PTM, short for post-translational modification enzyme. B). BGC architecture of *Gpe* family biosynthesis. C) *In vitro* assay of GpeC-catalyzed reaction on precursor GpeA_2_. D) Comparative UPLC-HRMS analysis reveals a 2H loss between substrate **1** and product **2**. E) Structural determination of biaryl-crosslinked Gpertide via NMR (**3**).

To identify the modified WMLY motif, we co-expressed GpeA_2_-GpeC *in vivo* (Supplementary Fig. S6). Chymotrypsin digestion of the full-length cyclized precursor yielded Gpertide (**3**), a 7-mer cyclic peptide containing a Trp1-Tyr4 biaryl crosslink. Comparative UPLC-HRMS analysis demonstrated structural identity between the *in vivo* and *in vitro* products (Supplementary Fig. S7), confirming the catalytic fidelity of GpeC. Comprehensive NMR analysis established the biaryl crosslink between Trp1-C7 and Tyr4-C3, evidenced by the singlet signal of Tyr4-H2 and an HMBC correlation from Tyr4-H2 to Trp1-C7. NOESY spectra further supported the biaryl system, revealing interactions between Tyr4’s phenolic OH hydrogen and Trp1-H6, as well as between Trp1-H2 and Tyr4-H2 (Fig. 2E, Supplementary Fig. S8). Marfey’s analysis confirmed L-configured methionine, leucine, and histidine residues in Gpertide (Supplementary Fig. S8). HPLC and NMR analyses revealed a single product, suggesting stereochemical fidelity in the biaryl C-C bond formation. NOE correlations (Fig. 2E) indicated restricted rotation of the biaryl system, supporting a *R*a configuration. This conclusion was corroborated by matching the experimental electronic circular dichroism (ECD) spectrum to the calculated spectra of predicted isomers (Supplementary Fig. S9).

### GpeC accepts truncated precursor peptides with a dynamic ‘adaptive recognition’

To elucidate the structural basis of GpeC’s catalytic versatility, we resolved its X-ray crystal structure at 2.62 Å resolution (PDB: 9ULW, Fig. 3A and Supplementary Table 7). The asymmetric unit contained two molecules, each adopting the canonical P450 fold with 15 α-helices, 13 β-strands, and a central heme group coordinated by a thiolate ligand (Supplementary Fig. S10A, B). Structural alignments using the Dali Sever^36^ confirmed that GpeC exhibits significant similarity to P450_Blt_ (PDB: 8U3N^33^, Z-scores of 25.6), with structural conservatism in the B/I/K- and L-helix regions (Supplementary Fig. S10C). Divergences emerged in the F-helix, G-helix, and BC-loop (RMSD = 5.435, Supplementary Fig. S10C), where GpeC’s helices adopt outward conformations, expanding its substrate-binding pocket. This structural adaptation correlates with a 57% larger active cavity volume than that of P450_Blt_ (2574 Å^3^, Fig. 3B and Supplementary Fig. S11), rationalizing its capacity to process bulkier substrates (Fig. 3A).

**Figure 3.**
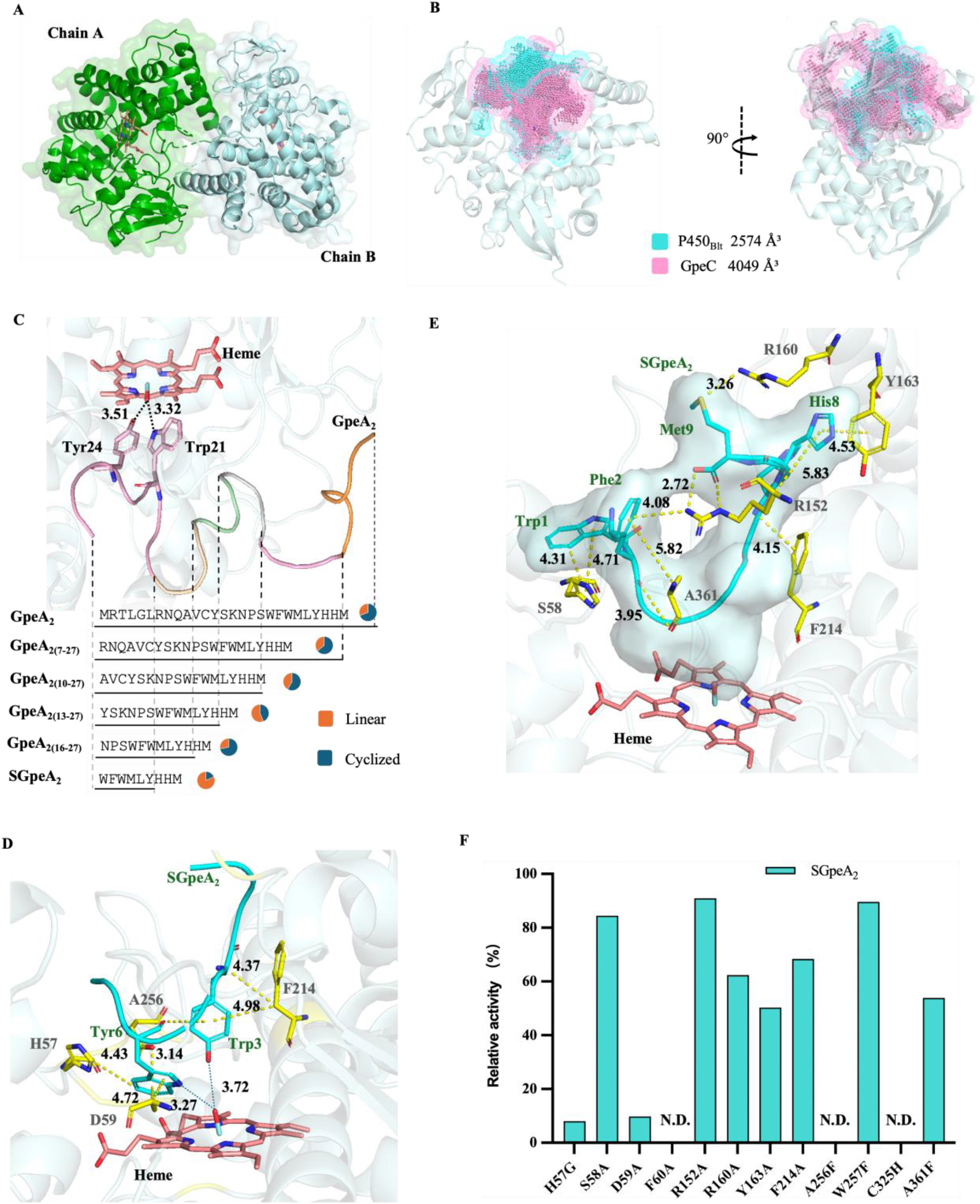
Structural characteristics and mutational analysis of GpeC. A) Crystal structure of GpeC. Green represents A-chain. Blue represents B-chain. B) Substrate cavity comparison of GpeC and P450_Blt_, with a size of 4049 Å^3^ (pink represents GpeC) and 2574 Å^3^ (blue represents P450_blt_). C) Conformation and truncation mode of GpeA_2_ in the substrate pocket of GpeC. D) Relative positions of the heme plane and the SGpeA_2_ (cyan sticks and cartoon) in the active pocket. E) Binding modes of SGpeA_2_’s N-terminus and C-terminus in the substrate pocket. F) Site-directed mutagenesis of amino acids related to the active pocket.

To understand how GpeC accommodates the native substrate GpeA_2_, we attempted to determine a co-crystal structure, but were hindered by the substrate’s flexibility and low binding affinity (K_*d*_=31.35 µM) (Supplementary Fig. S12A). Thus, molecular docking and dynamics simulations were employed to probe the substrate binding mode and identify key interacting residues. Molecular dynamics (MD) simulations revealed that the expansive substrate-binding cavity of GpeC is highly flexible. This cavity specifically accommodates the 27-mer GpeA_2_ peptide, precisely orienting the cyclization residues Trp21 and Tyr24 near the Fe-O intermediate at distances of 3.32 Å and 3.51 Å, respectively (Fig. 3C). Notably, the N-terminal 17 amino acids of the substrate extend outside the cavity (Supplementary Fig. S12B). GpeC’s unique binding mode with GpeA_2_ prompted us to test the leader region’s necessity in the catalytic process. Therefore, we systematically constructed N-terminal truncation mutants of GpeA_2_-derived substrates with lengths of 21, 18, 15, 12, and 9 amino acids and assessed the cyclization activity of GpeC (Fig. 3C). The results revealed that GpeC maintains catalytic efficiency with a minimal 9-mer substrate (SGpeA_2_, WFWMLYHHM). To verify structural consistency, we analyzed the enzymatic reaction products of SGpeA_2_ using UPLC-HRMS. Analysis revealed a -2 Da mass shift in the triply charged ion (observed [M+3H]^3+^ = 450.1964 vs. substrate [M+3H]^3+^ = 450.8685), consistent with biaryl bond formation. Tandem MS confirmed that modification occurred specifically within the ‘WMLY’ motif. These results align with observations from the native GpeA_2_ precursor (Supplementary Fig. S13). The retention of substrate recognition and catalytic activity upon truncation to a 9-residue core demonstrates that the leader peptide is dispensable.

To further elucidate the structural insights into the binding between SGpeA_2_ and GpeC, MD revealed a substrate-binding cavity stabilized by hydrogen bonds, hydrophobic interactions, and ionic bridges between the peptide and the enzyme (Fig. 3D and Supplementary Fig. S14). Root-mean-square fluctuation (RMSF) analysis revealed restricted SGpeA_2_ mobility, with flexibility primarily observed at Trp1, Phe2, Met4, Leu5, His7/8 and Met9 (Supplementary Fig. S15). The core region maintains rigidity to orient catalytic residues through an intricate interaction network (Fig. 3D), while terminal plasticity enables adaptive binding (Fig. 3E). In the catalytic conformation, the critical cyclization residues Trp3-Nδ1 and Tyr6-Oη of the SGpeA_2_ are positioned 3.27 Å and 3.72 Å away from the oxygen of the Fe-oxygen intermediate, respectively, suggesting that this distance ensures effective electron transfer during the reaction process (Fig. 3D). Concurrently, Met4 and Leu5 within the ‘WMLY’ motif exhibit conformational plasticity that accommodates local fluctuations while preserving precise spatial alignment between catalytic residues and reaction sites (Trp3/Tyr6) (Supplementary Fig. S16). This modular binding mechanism, characterized by weak terminal interactions positioning cyclization residues, core rigidity stabilizing catalytic geometry, and motif-level flexibility, enables maintaining optimal reaction conformations.

The substrate-binding site of cytochrome P450 enzymes is characteristically positioned adjacent to the heme group, encompassing the B-C loop, F-G loop, and helices F/G/I that collectively form the substrate recognition portal^37^. Guided by structural insights, site-directed mutagenesis of GpeC’s substrate pocket systematically elucidated critical residues governing enzymatic catalysis (Fig. 3D-F, Supplementary Fig.S17). The C325H mutation at the heme axial ligand position caused the characteristic red color to vanish, indicating the loss of cysteine thiolate-to-iron coordination, which resulted in the loss of catalytic activity. Mutational screening of B-C loop residues (H57-F60) revealed divergent functional consequences: H57G abolished the π-π T-shaped interaction with Trp1 and, along with D59G, disrupted the π-anion interaction with Trp3. F60A interfered with the π-amide stacked interaction with Tyr-6 and the π-π stacked and π-alkyl interactions with heme, resulting in a loss of catalytic activity. Conversely, S58A exhibited negligible activity alterations (<16% variation) due to insignificant spatial hindrance of residue A in the substrate cavity, barely affecting Trp1 binding. For R160A and Y163A mutations in the G-helix, along with R152A in the F-G loop and F214A mutations in the I-helix, despite disrupting critical interactions with His8-Met9 and Leu5-Tyr6 (hydrogen bond, π-π stacked, salt bridge, charge-charge interaction, Fig. 3E and Supplementary Fig. S16), retained partial activity (>50%), implicating auxiliary substrate-binding interactions and underscoring the multifactorial complexity of the catalytic interactome. W257F retained full activity (optimal van der Waals complementarity facilitated substrate positioning), while complete inactivation via A256F mutation arose from multiple disruptions of the hydrophobic microenvironment and Trp3 and heme π-alkyl interaction, confirming their synergistic role in active site integrity. These findings collectively highlight that this systematic mutagenesis deciphered a residue interaction network governing substrate recognition and catalytic efficiency, and the substrate fixation mode further guided us in the design of synthetic utility.

### GpeC is a versatile peptide biaryl macrocyclase through C-C, C-O, and C-N bonds

The total turnover number (TTN) of the SGpeA_2_ was not suitable as a substrate template for the establishment of a tetrapeptide biaryl cyclization platform (TTN ∼10.2) (Supplementary Fig. S18), as compared with the estimated TTN of P450_Blt_ (∼21, ∼85% conversion overnight)^38^. This limitation is likely due to SGpeA_2_’s poor solubility, stemming from its high content of hydrophobic aromatic amino acids (W, F, L, Y). We turned our sight on the other native precursor, GpeA_1_, and rationally truncated it into the 9-mer substrate WRWVMYHHM (SGpeA_1_). We proposed that the incorporation of the R residue highly increases the solubility of the substrate and facilitates the enzymatic capacity. Assaying SGpeA_1_ with GpeC, we found a higher TTN of ∼98.2 than P450_blt_ (Supplementary Fig. S19). Combined with molecular dynamics simulations and RMSF analysis, we found that SGpeA_1_ exhibits significantly higher flexibility than SGpeA_2_. The conformational flexibility of SGpeA_1_ enables active site residues to dynamically reposition its reactive atoms, specifically, the oxygen atom of Trp3 and the nitrogen atom of Tyr6, towards their optimal orientation for catalysis through dynamic interactions such as hydrogen bond reorganization, thereby bringing these atoms into proximity (∼3.2–3.6 Å) to the Fe-O intermediate (Supplementary Fig. S20). However, further truncation to 8-mer abolished activity, implicating critical roles in maintaining the 9-mer length for the catalytic process (Supplementary Figs. S21-22).

Given prior reports of P450 enzymes utilizing the peroxide shunt pathway^39^, we assessed GpeC’s activity under hydrogen peroxide (H_2_O_2_) at 28°C for 5 hours. UPLC-HRMS analysis of the reaction with SGpeA_1_ revealed a +30 Da product (**6**′) corresponding to a Trp-Tyr cyclized species (Fig. 4A, Supplementary Fig. S23). The substrate underwent sequential +16 Da oxidations (likely methionine sulfoxidation) and a concurrent cyclization within the ‘WVMY’ motif. To assess product consistency across redox systems, parallel reactions were conducted using the canonical Fdr/Fdx/NADPH system. After quenching with methanol and supplementing H_2_O_2_, the resulting products matched those from the H_2_O_2_-only condition (Supplementary Fig. S24), demonstrating GpeC’s capacity to utilize cost-effective, widely accessible peroxide as an alternative electron source. Kinetic and TTN assays revealed that GpeC was more catalytically efficient with substrate SGpeA_1_ under the NADPH-dependent system than the peroxide-driven system, with a *k*_cat_/*K*_m_ of 349.75 M^-1^S^-1^ (NADPH) and 287.46 M^-1^S^-1^ (Peroxide) and TTN of ∼74.6 (Peroxide) (Supplementary Fig.S24).

**Figure 4.**
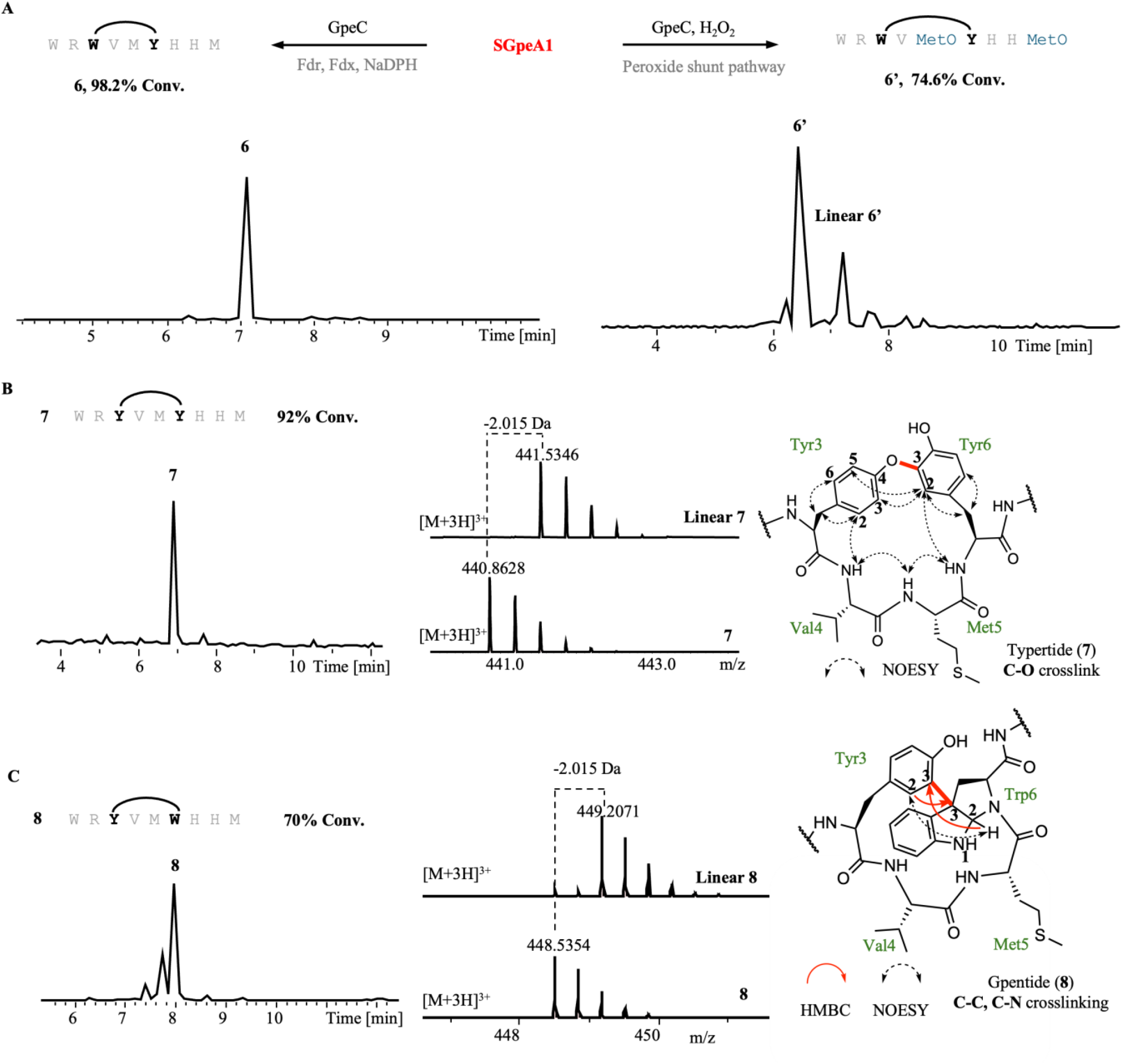
Catalytic versatility of GpeC. A) GpeC cyclization of 9-mer substrate SGpeA_1_ through redox partner pathway or peroxide shunt pathway. B) YVMY motif cyclization to Typertide (**7**) through GpeC-catalyzed C–O bond formation. Left: Comparative HRMS profile of modified product (**7**) and linear substrate. Right: NMR analysis of the corresponding biaryl cyclized module of Typertide (**7**). C) YVMW motif cyclization to Gpentide (**8**) via dual C–C/C–N cyclization by GpeC. Left: Comparative HRMS profile of modified product (**8**) and linear substrate. Right: NMR analysis of the corresponding biaryl cyclized module of Gpentide (**8**).

Beyond its native Trp3-Tyr6 crosslinking activity, GpeC catalyzed non-native Tyr3-Tyr6 (**7**) and Tyr3-Trp6 (**8**) cyclization products, expanding its utility for diverse biaryl architectures. To further characterize GpeC’s catalytic product, we synthesized Typertide (**7**) and Gpentide (**8**) via the enzymatic reaction of the 9-mer substrate WR**Y**VM**Y**HHM and WR**Y**VM**W**HHM, allowing for detailed structural analysis via NMR (Fig. 4B, C). NMR analysis of (**7**) confirmed the biaryl crosslink between Tyr3-O and Tyr6-C3, as indicated by the singlet signal of Tyr6-H2 and the CNMR chemical shift of Tyr6-C3 (δC =143.5). The NOESY spectra further substantiated the biaryl cyclized system by revealing interactions between Tyr3-H3 and Tyr6-H2 (Supplementary Fig. S25). Single product formation was observed via HPLC and NMR, leading us to hypothesize that the C-O bond is formed with stereochemical fidelity within the biaryl system, as dictated by the specificity of GpeC’s catalytic process. However, the NOE correlations unravel the symmetry signal of the hydrogen on Tyr3-H2 and Tyr3-H6 to Tyr3-Hβ, together with correlations between Tyr6-H2 with Tyr3-H3, and Tyr3-H5, suggesting that our bi-tyrosine cyclized ring does not exhibit atropo-chirality (Supplementary Fig. S26). NMR analysis of (**8**) confirmed the C-C crosslink between Tyr3-C3 to Trp6-C3, and a C-N bond was formed between Trp6-C2 to backbone N, as indicated by the HNMR of a singlet peak of Trp6-H2 (δH =5.94, s) and the CNMR of Trp6 C2 (δC =81.5), Trp6-C3 (δC =59.3). The HMBC spectra further verified the crosslink pattern of Tyr3-C3 to Trp6-C3 by the signal of Tyr3-H2 to Trp6-C3 and Trp6-H2 to Tyr3-C3. The NOESY spectra revealed the close spatial distance between Tyr3 and Trp6, as indicated by the signal of Tyr3-H2 to Trp6-H2. Marfey’s analysis verified L-configuration for all unmodified residues of compounds **7** and **8** (tryptophan, arginine, valine, methionine, histidine) (Supplementary Figs. S27-28).

### Broad substrate scope of GpeC’s governing the generalizable peptide biaryl cyclization

To probe GpeC’s substrate plasticity, we first analyzed phylogenetically related RiPP precursors sharing a similar biosynthetic logic to *Gpe*, based on our hypothesis that GpeC might retain catalytic compatibility with *Gpe*-like precursors. We found that these *Gpe*-like precursor peptides are conserved in the ‘WxxY’ catalytic motif but have divergent leader sequences. Sequence logo analysis of *Gpe*-like peptides guided our selection of native core regions for functional validation (Supplementary Fig.S29). Thus, we chemically synthesized 9-mer core sequences of prioritized candidates to assess GpeC’s cross-reactivity. Interestingly, we found that in the N-terminal region I W1R2, most of the mined precursors contained aromatic amino acids in the 1^st^ position, and most of them had R in the 2^nd^ position. Substitution of W1Y was acceptable, while further verification of W1F did not yield any cyclized product. Substitutions in the C-terminal region II H7H8M9 (H7S, H8N, H8D, M9I, M9L) retained compatibility with GpeC-mediated cyclization (Figure 5A, Supplementary Fig. S30-34), underscoring the enzyme’s tolerance for structural variation.

**Figure 5.**
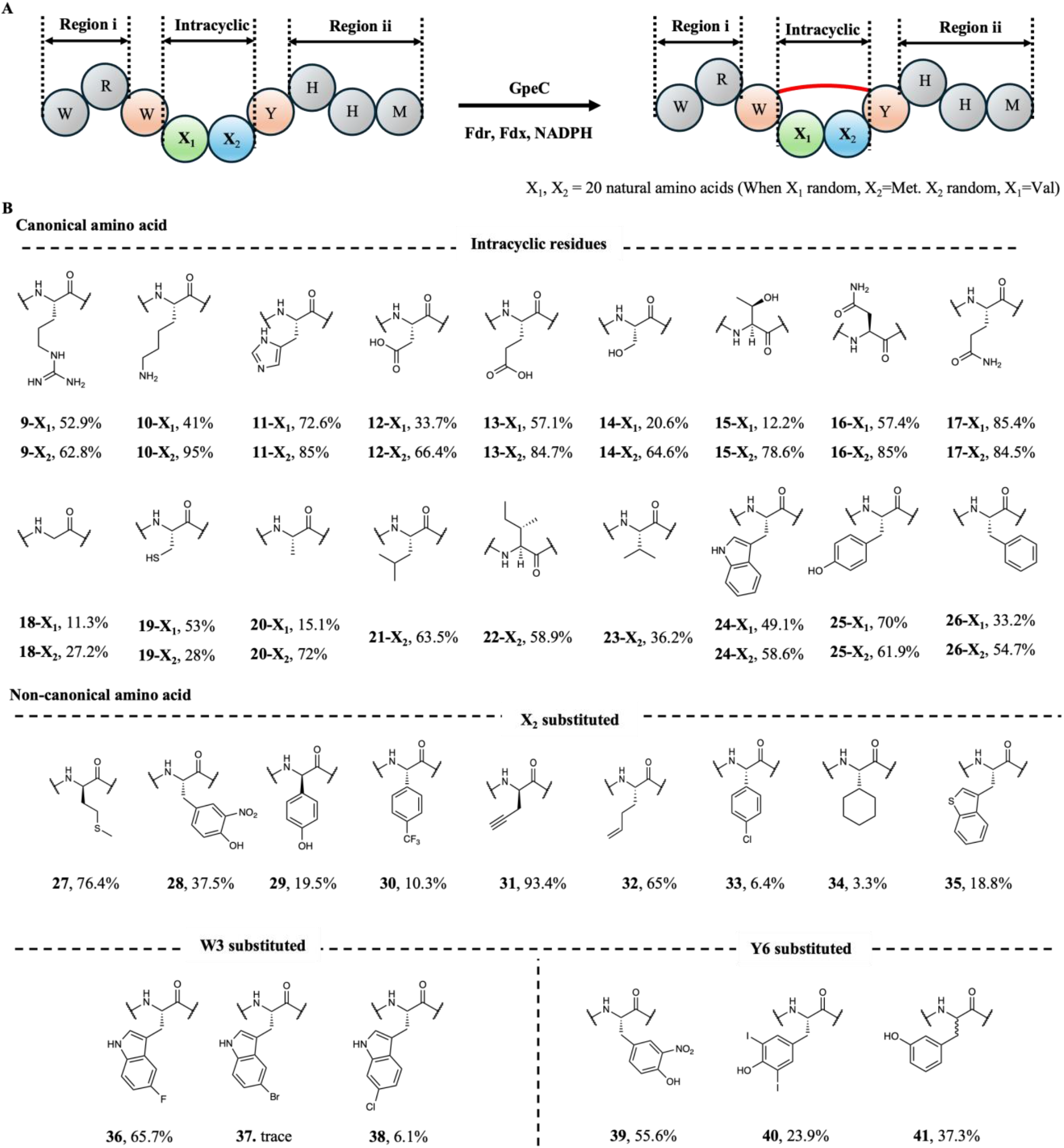
Substrate scope of GpeC. (A) Catalytic promiscuity assessment using diversified 9-mer peptides based on the SGpeA_1_ scaffold. (B) Comprehensive tolerance for 19 canonical amino acids and diverse noncanonical residues at intracyclic positions (X_1_, X_2_) and biaryl crosslinking sites (W3, Y6). *Top*: Canonical substitutions (X_1_/X_2_ positions). *Bottom*: Noncanonical amino acids (300 µM substrate, 3 µM GpeC, Fdx/FdR/NADPH system).

To systematically characterize GpeC’s tolerance for intracyclic residues in biaryl-cyclized scaffolds, we designed a panel of 9-mer precursor peptides derived from the SGpeA_1_ template, including WRWX_1_MYHHM and WRWVX_2_YHHM, in which positions X1 and X2 were individually substituted with each of the 20 canonical amino acids (Supplementary Fig. S35). Reactions were conducted at 28°C for 5 h in the presence of Fdr/Fdx/NADPH, quenched with methanol, and analyzed by UPLC-HRMS. GpeC exhibited broad substrate promiscuity, effectively accepting 19 of 20 canonical amino acids at both X1 and X2 positions, with average conversion rates of 47.2% (X1) and 68.6% (X2). Charged residues (R, H, K, D, E) were well tolerated, particularly H and E, which showed the highest conversions at X1 (72.6% and 57.1%) and a strong average at X2 (78.8%). Polar uncharged residues (N, Q) performed well at X1 (57.4% and 85.4%), whereas S and T significantly reduced activity (20.6% and 12.2%), suggesting incompatibility of terminal hydroxyl groups. Interestingly, at X2, all polar residues were well accommodated (78.2% average). Hydrophobic and aromatic residues showed size-dependent trends: small residues like G and A yielded low conversions (11.3% at X1), while bulkier side chains (L, I, V, W, Y, F) were efficiently processed (56.3% at X1; 60.2% at X2). As expected, proline disrupted cyclization at both positions due to its conformational rigidity (Supplementary Fig. S36). While the substrate concentration was not quantified in the mixed peptide groups, we selected four 9-mer peptides containing residues amenable to access late-stage modification and synthesized to quantitatively evaluate the enzymatic efficiency of GpeC (0.3 mM substrate, 3 µM enzyme). As expected, GpeC exhibited functional versatility, efficiently processing sterically hindered residues (e.g., Y, 70%) and functionally tractable residues (e.g., E, 85%, K, 95%) (Supplementary Figs. S37-40).

GpeC exhibited exceptional catalytic plasticity toward noncanonical amino acids. Quantitative assays (0.3 mM substrate, 3 μM enzyme) revealed D-type amino acid tolerance: while X_1_-D-valine substitution abolished activity, X_2_-D-methionine was accepted (51.7% conversion). We rationalized this differential tolerance by indicating a spatially accommodating X_2_ pocket. Capitalizing on this flexibility, we incorporated diverse non-native motifs, including nitrated/halogenated aromatics, cycloalkanes, sulfur heterocycles, and alkenyl/alkynyl handles, into the X_2_ position. Strikingly, GpeC processed all variants with retained efficiency (Fig. 5B; Supplementary Figs. S41-49). We further probed GpeC’s tolerance for noncanonical residues at crosslinking positions W3 and Y6. Substrates featuring halogenated (F/Cl/Br), nitrated, and meta-hydroxylated tyrosine analogs were synthesized. GpeC efficiently processed 5-F-Trp and 6-Cl-Trp at W3, evidenced by characteristic -2H mass shifts, while 5-Br-Trp proved recalcitrant (Supplementary Figs. S50-52). Remarkably, at Y6, GpeC accepted 3,5-diiodo-Tyr through deiodination and C-C bond formation. Nitrated-Tyr and m-Tyr were well-tolerated as well (Figure 5B, Supplementary Figs. S53-55). In summary, GpeC’s unparalleled substrate scope, spanning 19 canonical amino acids and diverse noncanonical residues under mild aqueous conditions, establishes it as a transformative biocatalyst for generalizable peptide biaryl macrocyclization.

### Versatility of GpeC in biocatalysis

The synthesis of intramolecular Trp-Tyr biaryl macrocyclic peptides, first demonstrated by Baran^14^ and later by Nakamura^11,13^ using palladium catalysis, exemplifies remarkable precision in controlling both site- and atroposelectivity. However, their seminal work also underscores the inherent challenges of synthetic routes, including laborious substrate-specific optimization of reaction conditions and cumbersome protecting-group strategies required to access biaryl-containing macrocyclic peptides (Final yield lower than 10%). Capitalizing on GpeC’s substrate versatility, we leveraged its biocatalytic potential for the de novo synthesis of complex macrocyclic natural products, Rubrin^30^. Our two-step biocatalytic approach involved: (1) solid-phase synthesis of the linear precursor WRWHHYHHM, and (2) GpeC-mediated biaryl cyclization (25 mL scale, 8 mg substrate, 5 μM GpeC) followed by trypsin cleavage. Purification via HPLC yielded 3 mg of cyclized product (45% crude yield, 37.5% isolated yield), which was unambiguously confirmed as rubrin’s core by NMR analysis, including signature Trp-Tyr C–C bond correlations matching natural rubrin (**42**) (Fig. 6A, Supplementary Table 6). The efficient biocatalytic synthesis of rubrin’s biaryl-cyclized core, demonstrating GpeC’s utility as a mild and direct platform for synthesizing structurally intricate peptide macrocycles with precise site selectivity on Trp C-7 to Tyr C-3, while proving GpeC’s activity with double mutations in its conserved “WxxY” catalytic motif at the same time, highlighting its exceptional versatility in catalyzing diverse substrates.

**Figure 6.**
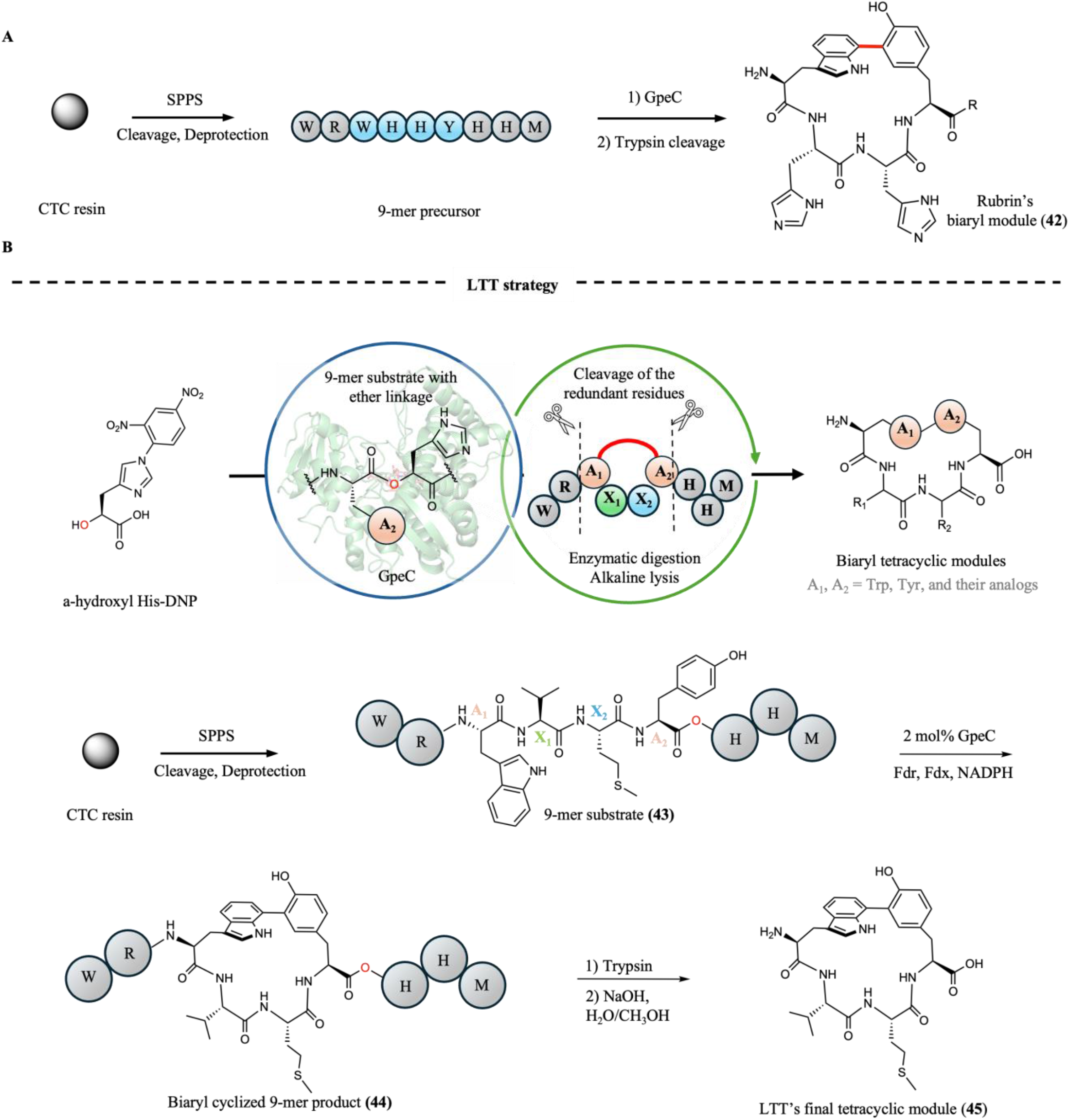
Versatility of GpeC in biocatalyst. A) Enzymatic semi-synthesis of the biaryl-cyclized core of a natural product, Rubrin. B) General chemo-enzymatic synthesis workflow of ‘LTT’ to generate the biaryl-cyclized tetrapeptide module.

While flanking residues in the native scaffold are essential for enzyme recognition, their structural constraints hinder post-cyclization diversification. To overcome this constraint, we engineered a “lytic-to-Tetraregion” (LTT) strategy, enabling direct construction of biaryl-cyclized tetrapeptides from GpeC’s 9-mer catalytic product. Key to this approach was replacing the Tyr6-His7 amide bond in substrate (**43**) with an ether linkage, which permitted GpeC-mediated cyclization to yield (**44**) (Fig. 6B). Subsequent trypsin cleavage of the N-terminal WR motif and alkaline hydrolysis (NaOH in a 1:1 H_2_O/CH_3_OH solution) of the C-terminal HHM generated a streamlined biaryl-cyclized tetrapeptide (**45**) (Fig. 6B; Supplementary Fig. S56). By excising nonessential structural elements, the LTT strategy further advances the GpeC’s peptide biaryl cyclization into a modular synthesis and unlocks rapid access to the structure-unique biaryl macrocyclic peptide motif.

## Discussion

Integrating biaryl motifs into macrocyclic peptides unveils a vast chemical space with untapped therapeutic potential. However, current methodologies, whether transitional metal-catalyzed peptide biarylation or NRPS-dependent enzymatic approaches, suffer from a generalizability bottleneck, limiting their broad applicability. While advances in retrosynthetic strategies for biaryl macrocyclization have improved catalytic selectivity, these catalysts tend to be intricate, expensive, and time-consuming to synthesize. Here, we report GpeC, the first generalizable biocatalytic platform for constructing biaryl-cyclized tetrapeptides. GpeC’s catalytic versatility is exemplified by its ability to forge three bond types: C–C (Trp–Tyr), C–O (Tyr–Tyr), and C–C/C–N (Tyr–Trp) cross-couplings, with acceptability on 9-mer leader-free substrates (SGpeA_1/2_). Structural analysis of the GpeC crystal structure rationalizes the leader-independent activity and reveals a spacious catalytic pocket, while molecular dynamics simulations on substrate RMSF calculations showed restricted ligand mobility specifically within the ‘WxxY’ motif. Although substrate co-crystallization proved elusive, computational docking and molecular dynamics simulations identified critical residues that govern substrate binding. Mutagenesis validated their role in a dynamic recognition mechanism we term ‘adaptive recognition’, wherein GpeC’s plastic binding site undergoes conformational adjustments to accommodate diverse substrates. This adaptive plasticity rationalizes its accommodation of 19 canonical amino acids and diverse noncanonical analogs while maintaining catalytic fidelity.

Capitalizing on GpeC’s versatility, we established a two-step biocatalytic synthesis of the biaryl-cyclized core of Rubrin. Leveraging GpeC’s inherent selectivity and catalytic efficiency, the reaction enables precise bond Trp C-7 to Tyr C-3 formation under mild, aqueous conditions. The synthesis of the biaryl-cyclized ‘WHHY’ module of Rubrin highlights GpeC’s expansive substrate scope, as the enzyme efficiently processes substrates containing double mutations within its conserved ‘WxxY’ motif while maintaining catalytic fidelity, indicating an over 19*19 substrate acceptability. Taking advantage of this adaptability, we formulated a one-step, single-enzyme synthesis ‘LTT’ to modularly construct biaryl-cyclized tetrapeptides. The establishment of the biocatalytic biaryl cyclized tetrapeptide synthesis method ‘LTT’ serves as a rapid access to these pharmacologically relevant motifs and facilitates late-stage diversification on the cyclized tetrapeptide motif.

The discovery of GpeC represents a significant advancement in the biocatalytic toolkit for generizable tetrapeptide biaryl cyclization. Nonetheless, further engineering of GpeC is required to enhance its catalytic efficiency, as evidenced by its currently low TTN. Additionally, the enzyme’s catalytic activity remains restricted to tetrapeptide substrates, with limited efficacy toward tri-or pentapeptide architectures. This limitation is likely attributable to the cyclization’s dependence on a stabilized radical intermediate, which necessitates precise spatial alignment of reactive residues within the tetrapeptide framework. Therefore, a thorough study of the catalytic mechanism is necessary to guide GpeC engineering rationally, encompassing both catalytic efficiency (e.g., TTN) and substrate scope. Despite these limitations, GpeC’s robust *in vitro* production and exceptional solubility position it as an ideal candidate as the peerless starting point for PBC. Future research might leverage GpeC’s characteristics to engineer variants capable of catalyzing structurally diverse biaryl-linked macrocycles, overcoming current limitations confined to tetrapeptide substrates. We firmly believe that the synergy between GpeC’s bioorthogonal and catalytic promiscuity, combined with peptide display technologies, positions this system as a transformative platform for discovering bioactive macrocycles, offering unprecedented potential to accelerate the identification of lead compounds against undruggable targets soon. In all, this study unveils nature’s enzymatic potential for constructing architecturally complex biaryl macrocyclic peptides, establishing GpeC as a benchmark biocatalytic platform that circumvents traditional synthetic limitations.

## Method

### General chemical reagents

All chemicals and culture medium, including kanamycin, Chloramphenicol, Spectinomycin, and isopropyl β-D-1-thiogalactopyranoside (IPTG), are purchased from Sangon Biotech (Shanghai), MACKLIN, or Sigma (USA). Marfey’s reagent Nα-(5-Fluoro-2,4-dinitrophenyl)-L-leucinamide (L-FDLA), Nα-(5-Fluoro-2,4-dinitrophenyl)-D-leucinamide (D-FDLA), and Tioglycolic acid (TGA) were purchased from TCI and MACKLIN. Oligonucleotide primers were synthesized from Ruibotech (Guangzhou). DNA polymerase Phanta Max Master Mix was purchased from Vazyme. TEV protease, endonucleases BamHI, Hind III, XhoI, NcoI, NdeI, XbaI, and Gibson assembly kit NEBuilder® HiFi DNA Assembly were purchased from New England Biolabs. Chymotrypsin was purchased from MACKLIN. Trypsin and PD-10 MidiTrap desalting columns were purchased from Sigma. HISPUR NI−NTA resin was purchased from Thermo Fisher. NMR solvent Dimethyl sulfoxide-d^6^ was purchased from Cambridge Isotope Laboratories.

### Bioinformatic analysis

SPECO pipeline was used to calculate co-occurred sORF-P450 pairs. Sequence clustering analysis of sORF and P450 enzyme sequences was conducted using MMseqs2^40^ with the following parameters: mmseqs --easy-cluster --min-seq-id [0.6 for P450, 0.35 for small peptide] --seq-id-mode 1 --cluster-mode 1 --similarity-type 2 --single-step-clustering --dbtype 1. Multilayer sequence similarity network (MSSN) analysis of sORF-P450 pairs was performed by employing the Python package NetworkX and visualized by using Matplotlib.

### Cloning and plasmid construction

Redox partner gene synthesis and construction. For spinach ferredoxin and ferredoxin reductase, we synthesized the same nucleotide sequences (sequence id: *Spinach ferredoxin* and *Spinach ferredoxin reductase*) as reported by Prof. Xudong Qu et al.^41^. The *E. coli* thioredoxin gene was fused to the N-terminal of ferredoxin to increase the solubility, as reported by Prof. Xudong Qu et al^41^. The maltose binding protein (MBP) was fused to the N-terminal of ferredoxin reductase to increase the solubility. These two fusion proteins were respectively cloned into multiple cloning sites 1 of pCDF-Duet-1 and pRSF-Duet-1 vectors, generating pRSF-fdr and pCDF-fdx.

Precursor and P450 enzyme cloning and plasmid construction. For precursor cloning, GpeA_1_, GpeA_2_ were individually cloned by using each genome as a template and inserted into the multiple cloning site 1 of pRSF-Duet-1 with an N-terminal His^6^-SUMO-TEV tag, yielding pRSF-SUMO-GpeA_1_ and pRSF-SUMO-GpeA_2_ plasmids. For precursor and P450 coexpression, the corresponding P450 enzyme GpeC was cloned and inserted into the multiple cloning site 2 of pRSF-SUMO-GpeA_1_ and pRSF-SUMO-GpeA_2_, generating pRSF-SUMO-GpeA_1_C and pRSF-SUMO-GpeA_2_C. P450 enzyme cloning, GpeC was individually cloned by using each genome as a template and inserted into the multiple cloning site 1 of pRSF-Duet-1 with an N-terminal His^6^-tag, yielding pRSF-His-GpeC.

### Transformation, protein overexpression, and purification

For precursor expression, each redox partner, including pRSF-fdr and pCDF-fdx, was individually transformed with *E. coli* Rosetta to generate strain E. coli-redox. Plasmids pRSF-SUMO-GpeA_2_ and pRSF-SUMO-GpeA_1_ were respectively transformed with *E. coli* Rosetta (DE3). For co-expression of precursor and P450, pRSF-SUMO-GpeA_2_-GpeC were respectively transformed with *E. coli*-redox or *E. coli* Rosetta (DE3).

After transformation, a single colony of each recombinant strain was inoculated into 5 mL fresh LB broth with 34 μg/mL chloramphenicol (native resistance for *E. coli* Rosetta (DE3)) and 50 μg/mL kanamycin (for pRSF-Duet-1 backbone) and shaken at 37 °C overnight. Then, 1 mL of the culture was incubated in 100 mL fresh TB medium (24 g/L yeast extract, 12 g/L casein hydrolysate, 9.4 g/L K_2_HPO_4_ and 2.2 g/L KH_2_PO_4,_ and 8 ml/L glycerol) with antibiotics. After an additional 2-4 hours of growth, the culture broth was cooled to 24 °C when OD_600_ reached 0.8. A final concentration of 0.4 mM IPTG was added to induce protein expression, followed by a 20-hour induction at 16 °C, 150 rotations per minute (rpm). The cells were harvested at 4 °C and centrifuged at 10,000 rpm for five minutes.

Each His^6^-SUMO-tagged peptide was purified using a denaturing method. The following buffers were used for purification: denature buffer A (50 mM Tris, 500 mM NaCl, 6 M guanidine hydrochloride, adjusted pH to 8.0 using HCl), denature buffer B (30 mM imidazole in denature buffer A) and buffer C (50 mM Tris, 500 mM NaCl, 500 mM imidazole, adjusted pH to 8.0 using HCl). The cells were first resuspended using denaturing buffer A and then lysed using Sonics VCX-750 Vibra-Cell Ultrasonic Liquid Processor with a 13 mm probe, 3 seconds on and 3 seconds off for 10 minutes. The cell lysis was then centrifuged at 4 °C, 10,000 rpm for 1 hour, and the supernatant was loaded into pre-equilibrated Ni−NTA resin (using denaturing buffer A) three times to maximize target peptide yield. 5 mL denaturing buffer B was used to wash away non-specifically bound proteins. The target peptide was then eluted by using 1 mL buffer C 4 times, i.e., 4 mL in total. The purified peptide was desalted using a PD-10 desalting column. TEV protease and trypsin digestion were conducted at 30 °C for 1 hour and 37 °C for 12 hours, respectively. After digestion, the reaction mixture was centrifuged at 10000 rpm for 10 minutes, and the supernatant was subjected to UPLC-HRMS analysis.

For purifying the P450 enzymes, the pGroEL/ES chaperone plasmid was transformed into *E. coli* BL21 first, generating a chaperone-based *E. coli* BL21-c. Then, pRSF-Histag-GpeC was transformed into E. coli BL21-c. After transformation, a single colony of each recombinant strain was inoculated into 3 mL fresh LB broth with 34 μg/mL chloramphenicol (native resistance for chaperone plasmid) and 50 μg/mL kanamycin (for pRSF-Duet-1 backbone) and shaken at 37 °C overnight. Then, 1 mL of the culture was incubated in 500 mL fresh LB medium (10 g/L tryptone, 5 g/L yeast extract, 10 g/L NaCl) with antibiotics. After an additional 3 hours of growth, when OD_600_ reached 0.5, the culture broth was cooled to 30 °C, and a final concentration of 6 g/L arabinose, 1.5 mM 5-aminolevulinic acid was added to the culture medium. Keep shaking until OD_600_ reaches 0.8. A final concentration of 0.4 mM IPTG was added to induce protein expression, followed by a 20-hour induction at 24 °C, 150 rotations per minute (rpm). The cells were harvested at 4 °C and centrifuged at 4900 rpm for an hour. The following buffers were used for purification: buffer A (50 mM Tris, 500 mM NaCl, 5% glycerine, adjusted pH to 8.0 using HCl), buffer B (30 mM imidazole in buffer A) and buffer C (50 mM Tris, 500 mM NaCl, 500 mM imidazole, adjusted pH to 8.0 using HCl). The cells were first resuspended using buffer A and then lysed using Sonics VCX-750 Vibra-Cell Ultrasonic Liquid Processor with a 13 mm probe, 3 seconds on and 4 seconds off for 30 minutes. The cell lysis was then centrifuged at 4 °C, 10,000 rpm for 2 hours, and the supernatant was loaded into pre-equilibrated Ni−NTA resin (using buffer A) three times to maximize target protein yield. 5 mL buffer B was used to wash away non-specifically bound proteins. The target peptide was then eluted by using 1 mL buffer C for 4 times, i.e., 4 mL in total. The purified protein was desalted using a PD-10 desalting column.

### GpeC general biaryl-cyclization assay

The P450 enzyme GpeC and substrate mixtures (50 µL) containing 0.3 mM substrate, 3 µM P450, 3 µM Ferredoxin reductase, 3 µM Ferredoxin, and 1 mM NADPH in 50 mM Tris-HCl buffer with 5% glycerol (pH 7.5) were incubated at 28 °C for 5 h. The reactions were quenched by methanol, and the precipitated protein was removed by centrifugation. The supernatant was subjected to LC–MS analysis. Conversions were calculated by the following,

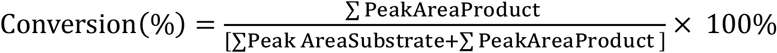

### GpeC biaryl-cyclization Michelle-Menten kinetics assay

The P450 enzyme GpeC and substrate mixtures (50 µL) containing 5, 10, 20, 40, 80, 120, 160,240, 320, 480 µM substrate, 3 µM P450, 3 µM Ferredoxin reductase, 3 µM Ferredoxin, and 1 mM NADPH in 50 mM Tris-HCl buffer with 5% glycerol (pH 7.5) were incubated at 28 °C for 30 min. The reactions were quenched by the addition of methanol, and the precipitated protein was removed by centrifugation. The supernatant was subjected to LC–MS analysis.

### Protein crystallization and structure determination

The GpeC protein was crystallized at a concentration of 10-14 mg/mL in SEC buffer with reservoir solution (10% w/v PEG 4000, 20% v/v glycerol, 0.03 M magnesium chloride, 0.03 M calcium chloride, 0.1 M MES/imidazole pH 7.25) by sitting drop vapor diffusion technique. After 7 days of growth at 16°C, crystals were observed in 0.8 μL sitting drops composed of a 1:1 mixture of protein solution and reservoir solution. The crystals were then cryoprotected by briefly soaking in the corresponding reservoir solution containing an additional 5% (v/v) ethylene glycol before being transferred to liquid nitrogen. Data collection was carried out on the BL18U1 beamline at Shanghai Synchrotron Radiation Facility (SSRF, Shanghai, China). The dataset was indexed and integrated using XDS^42^ and scaled by AIMLESS from the CCP4i suite^43^. The GpeC crystals achieved a diffraction resolution of 2.62 Å and belonged to the space group P4_3_2_1_2, with unit cell parameters a=b=78.40 Å, c=326.90 Å, α=β=γ=90.0°. There were two molecules in the asymmetric unit. The structure was phased by PHASER2^44^ using predicted GpeC coordinates by Alphafold3^45^ as the molecular replacement model. The models were completed through manual model building using COOT4^46^, interspersed with crystallographic refinement using REFMAC5^44^ with TLS restraints. Phenix.table_one^44^ was utilized to create the crystallographic statistics table (excluding the Rwork/Rfree values), which indicated that 96.11% of the residues were in the most favored regions of the Ramachandran plot. Data collection, phasing, and refinement statistics are detailed in Table 7. Structure figures were generated using PyMOL2.3.4.

### System Setup

The structure of protein GpeC was used to build the model system by the crystallized structure. The native protein was called WT protein. And we used PyMol^47^ Mutagenesis module to mutate one residue of the original protein, which is called WT or F62G protein. The peptide ligands were docked into the active site of the P450 protein via AutoDock Vina^48^. The associated atomic interactions between residues were described by the classical force field Amber ff14sb. The molecular structures of peptides were optimized at the MP2/6-31G* level of theory by Orca^49^. The HEM parameters are from Cpd I^50^.

### The classical molecular dynamics simulation

The solvents were explicitly represented by the TIP3P model^51^ under periodic boundary conditions (PBC). Na^+^ and Cl^-^ were added to make neutralize the system. The initial atomic coordinates of WT/F62G systems were with implicit solvent. In an isothermal–isobaric (NPT) ensemble, the system temperature and pressure were maintained at 300 K and 1 bar using the respectively Langevin thermostat^52^ and Monte Carlo barostat^53^. The electrostatic interactions were calculated via the Particle mesh Ewald (PME) method^54^. The cutoff of electrostatic and vdW interactions are both 1.2 nm. The simulation box is cubic, and its size is adjusted so that its boundary is more than 1.0 nm away from the protein. After energy minimization and NPT equilibrium, 200 ns of MD simulations were performed. The simulation of each system is performed three times in parallel. The time step was set as 2 fs. All classical MD simulations were performed with OpenMM^55^. And the binding free energy is calculated by MMPBSA.py^56^ in AmberTools^57^.

## Acknowledgments

This work is partially funded by the Research Grants Council of Hong Kong (17115322, 17102123, and 17101324) to Y-X.L. and Jiangsu Basic Research Center for Synthetic Biology Grant (BK20233003 to J.Z.). The authors would like to acknowledge and thank Yi-Man Eva Fung, Jo Yip, Xiaolin Zou, and Bonnie Yan for their help in MS and NMR analysis. We thank Yuqi Shi for her help with the primary bioassay. We also thank the staff of beam line BL18U1 of Shanghai Synchrotron Radiation Facility for access and help with the X-ray data collection.

## Author contributions

R-Z.L. and B-B.H., and Y-X.L. designed the research. G-F.W. and R-Z.L. conducted the bioinformatic analysis. R-Z. L. characterized the BGC, isolated the compounds, and characterized the structures.C-P.P. and H-Y.R. characterized the crystal structure. Z-L.S performed in silico calculations, R-Z. L. and J.L. contributed to *in vitro* enzyme characterization. R-Z. L and Z-M.S. conducted the Advanced Marfey’s experiment. R-Z.L. and B.H. synthesized all peptides used in this study. All authors analyzed data and discussed results. All authors participated in preparing the manuscript. J-H. Z. and Y.X. L. supervised the project.

## Additional information

Supplementary information

## Data availability

The paper, supplementary information contain data supporting this study’s results. The corresponding authors can provide any other data or datasets from this study upon reasonable request.

## Code availability

Data from genome mining, mass calculation and mapping were analyzed using scripts written in the programming language Python. The code of the scripts is available from https://github.com/yxllab-hku.

