## Supplementary Information for "Harnessing P450 enzyme plasticity for generalizable peptide biaryl macrocyclization"

### Table of Contents


**1. Supplementary Methods**

**2. Supplementary Tables**

Supplementary Table 1: Primers used in this study.

Supplementary Table 2: DNA sequences used in this study.

Supplementary Table 3: NMR spectroscopic data for Gpertide (**3**).

Supplementary Table 4: NMR spectroscopic data for Typertide (**7**).

Supplementary Table 5: NMR spectroscopic data for Gpentide (**8**).

Supplementary Table 6: NMR spectroscopic data for biocatalyzed ’WHHY‘ module.

Supplementary Table 7: Crystal data collection and refinement statistics.

**3. Supplementary Figures**

Supplementary Fig.1: General genome mining of RiPP-P450s.

Supplementary Fig.2: Bioinformatic analysis of GpeC analogs.

Supplementary Fig.3: Expression systems used in this work.

Supplementary Fig.4: Carbon monoxide binding assay performed with $P450_{GpeC}$ and $P450_{GpeC\text{-}F62G}$.

Supplementary Fig.5: GpeC enzymatic assay with $GpeA_2$, then digested by trypsin.

Supplementary Fig.6: In vivo characterization of the *Gpe* family.

Supplementary Fig.7: Comparative analysis of the consistency of the in vivo and in vitro catalyzed product.

Supplementary Fig.8: Structure elucidation of Gpertide (**3**).

Supplementary Fig.9: Electronic Circular Dichroism (ECD) analysis of Gpertide (**3**).

Supplementary Fig.10: Analysis of the GpeC structural characteristics.

Supplementary Fig.11: Analysis of substrate cavity of GpeC and $P450_{Blt}$.

Supplementary Fig.12: Analysis of GpeC binding affinity and optimal conformation with $GpeA_2$.

Supplementary Fig.13: Truncated N-terminal leader region of the native precursor peptide $GpeA_2$.

Supplementary Fig.14: Analysis of the interaction between GpeC and substrate $SGpeA_2$.

Supplementary Fig.15: Variation diagram of ligand RMSF during MD simulation process.

Supplementary Fig.16: Interaction network between GpeC and $SGpeA_2$.

Supplementary Fig.17: Site-directed mutagenesis of GpeC.

Supplementary Fig.18: In vitro assay of $SGpeA_2$ with GpeC.

Supplementary Fig.19: In vitro assay of $SGpeA_1$ with GpeC.

Supplementary Fig.20: Binding conformation of $SGpeA_1$ and GpeC.

Supplementary Fig.21: In vitro assay of WRWVMYHH with GpeC.

Supplementary Fig.22: In vitro assay of 8-mer substrate RWVMYHHM with GpeC.

Supplementary Fig.23**:** In vitro assay of $SGpeA_1$ with GpeC by peroxide shunt pathway.

Supplementary Fig.24: Catalytic consistency of GpeC under redox and the peroxide shunt pathway.

Supplementary Fig.25: GpeC enzymatic assay with WRYVMYHHM.

Supplementary Fig.26: Structure elucidation of Typertide (**7**).

Supplementary Fig.27: GpeC enzymatic assay with WRYVMWHHM.

Supplementary Fig.28: Structure elucidation of Gpentide (**8**).

Supplementary Fig.29: GpeC enzymatic assay of a series of bioinformatic-guided 9-mer substrates.
Supplementary Fig.30: GpeC enzymatic assay with WRWVHYHNM.
Supplementary Fig.31: GpeC enzymatic assay with YRWYLYHHI.
Supplementary Fig.32: GpeC enzymatic assay with WRWYLYHSM.
Supplementary Fig.33: GpeC enzymatic assay with WRWVHYHDL.
Supplementary Fig.34: GpeC enzymatic assay with YIWIMYNNI.
Supplementary Fig.35: General synthesis method for the substrate scope screening.
Supplementary Fig.36: Conformation of GpeC with substrates WRWPMYHHM and WRWVPYHHM.
Supplementary Fig.37: GpeC enzymatic assay with WRWYMYHHM.
Supplementary Fig.38: GpeC enzymatic assay with WRWKMYHHM.
Supplementary Fig.39: GpeC enzymatic assay with WRWVKYHHM.
Supplementary Fig.40: GpeC enzymatic assay with WRWVEYHHM.
Supplementary Fig.41-56: GpeC enzymatic assays with substrate incorporating non-native amino acids.

**4. UPLC-HRMS data**

Truncated leader peptide assay.
Mutations on the ring-forming residues.
Random amino acids incorporated into the intracyclic amino acids within the substrate.

**5. NMR spectra**

NMR spectra of **3.**
NMR spectra of **7.**
NMR spectra of **8.**
NMR spectra of **42.**
NMR spectra of **His(OH)-DNP**.

#### Supplementary Method

##### Strain preparation

All wild-type strains used in this study were purchased from the German Collection of Microorganisms and Cell Cultures GmbH (https://www.dsmz.de/). Cultivation was performed by following the guidance from DSMZ. *Escherichia coli* DH5α was used as cloning host. *Escherichia coli* Rosetta (DE3), and *Escherichia coli* BL21 (DE3) were used as expression hosts.

##### Mutation assay of enzymes

*GpeC* mutants were constructed by using PCR and confirmed by sequencing. Each mutant was individually transformed with *E. coli* BL21 containing chaperon plasmid for expression. Mutant purification was conducted by the abovementioned methods.

##### LC-HRMS analysis of purified peptides

For LC-MS analysis, Waters ACQUITY UPLC BEH C18, 130Å, 1.7 μm column was used. LC-HRMS experiments were performed on Thermo Scientific UltiMate 3000 UHPLC system coupled with Bruker impact Mass Spectrometer. The following LC method was used:

| | |
|---|---|
| column temperature | 40˚C |
| flow rate | 0.2 mL/min |
| solvent A | 0.1% formic acid in $H_2O$ |
| solvent B | 0.1% formic acid in acetonitrile |
| 0-2 min | 5% B |
| 2-15 min | 5% to 95% B |
| 15-19 min | 95% B |
| 19-21 min | 5% B |

The MS system was tuned using a sodium formate standard. All the samples were analyzed in positive polarity with m/z range from 150 to 1500 Th, using a data-dependent acquisition mode.

##### Semi-preparation of modified peptides

Larger-scale fermentation of *E. coli* Rosetta (DE3) harboring the recombinant plasmid pRSFDuet-GpeA$_2$-GpeC was performed in a 15 L LB medium with IPTG induction. Protein purification yielded about 100 mg of modified GpeA2 with an N-terminal hexa-histidine-SUMO tag. The crude sample was digested with trypsin at 30 °C for 20 h. After digestion, the crude sample was extracted with n-butanol and then concentrated and dissolved in 1 mL of methanol solution. The sample was further subjected to semipreparative HPLC (Waters 1525, USA) using a Phenomenex column (Kinetx 5 μm XB-C18 100 Å, 250 X10 mm) under the following conditions:

Phase A: 99.95% $H_2O$ with 0.05% Trifluoroacetic acid,

Phase B: 99.95% acetonitrile with 0.05% Trifluoroacetic acid,

0-5 min: 20% B,

5-40 min: 20%-40% B,

40-50 min: 100% B; 50-60 min: 20% B.

The eluted samples were freeze-dried to afford compounds **3**. Compound **3** was dissolved in 200 μL Dimethylsulfoxide-$d_6$ separately for NMR analysis.

Substrate WRYVMYHHM was synthesized by SPPS (solid-phase peptide synthesis). Large-scale reaction of 8mg substrate WRYVMYHHM was performed with 1 uM NADPH, 3 uM P450, 3 uM Fdr, and 3 uM Fdx under 20 mL of 50 mM Tris-HCl buffer with 5% glycerol (pH 7.5). The scale-up reaction was performed at 28 degrees for 5 hours and quenched with an equivalent volume of methanol. The sediment was removed through centrifugation, and the supernatant was subjected to semipreparative HPLC (Waters 1525, USA) using a preparative column (SilGreen C18, 5 μm, 12nm, 250*20mm) under the following conditions:

Phase A: 99.95% $H_2O$ with 0.05% Trifluoroacetic acid,

Phase B: 99.95% acetonitrile with 0.05% Trifluoroacetic acid,

0-5 min: 5% B,

5-40 min: 5%-50% B,

40-50 min: 100% B; 50-60 min: 5% B.

The fragment was freeze-dried to afford compounds **7** (white powder, $t_R$ 22.4 min, 3 mg). Compound **7** was dissolved in 200 μL Dimethylsulfoxide-$d_6$ separately for NMR analysis.

Substrate WRYVMWHHM was synthesized by SPPS (solid-phase peptide synthesis). Large-scale reaction of 10 mg substrate WRYVMWHHM was performed with 1 uM NADPH, 5 uM P450, 5 uM Fdr, and 5 uM Fdx under 25 mL of 50 mM Tris-HCl buffer with 5% glycerol (pH 7.5). The scale-up reaction was performed at 28 degrees for 5 hours and quenched with an equivalent volume of methanol. The sediment was removed through centrifugation, and the supernatant was subjected to semipreparative HPLC (Waters 1525, USA) using an analytical column (Cosmosil PBr, 5 μm, 250*4.6mm) under the following conditions:

Phase A: 99.95% $H_2O$ with 0.05% Trifluoroacetic acid,

Phase B: 99.95% acetonitrile with 0.05% Trifluoroacetic acid,

0-5 min: 25% B,

5-40 min: 25%-60% B,

40-50 min: 100% B; 50-60 min: 25% B.

The fragment was freeze-dried to afford compounds **8** (white powder, $t_R$ 17.5 min, 2.8 mg). Compound **8** was dissolved in 200 μL Dimethylsulfoxide-$d_6$ separately for NMR analysis.

Substrate WRWHHYHHM was synthesized by SPPS (solid-phase peptide synthesis). Large-scale reaction of 8 mg substrate WRWHHYHHM was performed with 1 uM NADPH, 5 uM P450, 5 uM Fdr, and 5 uM Fdx under 20 mL of 50 mM Tris-HCl buffer with 5% glycerol (pH 7.5). The scale-up reaction was performed at 28 degrees for 5 hours and quenched with an equivalent volume of methanol. The sediment was removed through centrifugation, and 5 µl of trypsin enzyme was added to the supernatant overnight, and the mixture was subjected to semipreparative HPLC (Waters 1525, USA) using an analytical column (Cosmosil PBr, 5 µm, 250*4.6mm) under the following conditions:

Phase A: 99.95% $H_2O$ with 0.05% Trifluoroacetic acid,
Phase B: 99.95% acetonitrile with 0.05% Trifluoroacetic acid,
0-5 min: 8% B,
5-40 min: 8%-50% B,
40-50 min: 100% B; 50-60 min: 8% B.

The fragment was freeze-dried to afford compounds **42** (white powder, $t_R$ 18.5 min, 3 mg). Compound **42** was dissolved in 200 µL Dimethylsulfoxide-$d_6$ separately for NMR analysis.

**NMR spectroscopy**

$^1$H NMR, $^{13}$C NMR, HSQC, HSQC-DEPT, $^1$H-$^1$H COSY, and NOESY spectra were acquired on a Bruker Avance 600 MHz spectrometer with Cryoprobe, using DMSO-*d6* as solvent. The Dimethyl sulfoxide-*d6* chemical shifts were used as the internal reference.

**Acid hydrolysis and Advanced Marfey's analysis**[1]

For acid hydrolysis, 0.1 mg dried compounds were prepared and dissolved in 1.5 mL 6M HCl in a glass vial and sealed tightly. Glass vials were then incubated in an oil bath at 110 °C for 16 hours. After hydrolysis, the aqueous phase was dried under a $N_2$ stream. The remaining solid was dissolved in 100 µL $H_2O$ and equally transferred into two glass vials, followed by adding 100 µL 1% L-FDLA and D-FDLA (dissolved in acetone), respectively. For each vial, 20 µL of 1 M $NaHCO_3$ stock solution was added and incubated at 40 °C for 1 hour. After the mixture was cooled to room temperature, 10 µL 2 M HCl was used to quench the reaction and diluted by adding 200 µL methanol. After centrifugation at 10000 rpm for 10 minutes, the reaction mixtures were analyzed by liquid chromatography low-resolution mass spectrometer.

**Electronic Circular Dichroism experiment**

The purified peptides were dissolved in $H_2O$ at the final concentration of 0.1 mg/mL. The CD spectrum of compounds was collected on a Jasco J-815 CD spectrometer with the following parameters:

Band Width, 5 nm

Measure Range, 400-250 nm

Data Pitch, 1.0 nm

Scanning Speed, 100 nm/min

D.I.T, 2 sec

For ECD calculation, the theoretical calculations were carried out using Gaussian 09[2]. At first, all conformers were optimized at PM6. Room-temperature equilibrium populations were calculated according to Boltzmann distribution law, based on which dominant conformers of the population over 1% were kept. The chosen conformers were further optimized at B3LYP/6-31G(d) in the gas phase. Vibrational frequency analysis confirmed the stable structures. ECD calculations were conducted at B3LYP/6-311G(d,p) level in $H_2O$ with IEFPCM model using Time-dependent Density functional theory (TD-DFT). Rotatory strengths for 30 excited states were calculated. The ECD spectrum was simulated using the ECD/UV analysis tool in Yinfo Cloud Computing Platform (https://cloud.yinfotek.com/) by overlapping Gaussian functions for each transition.

**Code availability**

**Python scripts availability**. Source codes of SPECO pipeline and logo sequence generation are available via https://github.com/yxllab-hku/speco and https://github.com/Bio-bbhe.

**Peptide substrate synthesis by solid-phase peptide synthesis[3]**

**Attachment of the first amino acids.** 2-chlorotrityl chloride (CTC) resin (1.08mmol/g, CSBio) in a Libra tube was swollen in dichloromethane (DCM) for 20 minutes before washing the resin with DCM 3 times. The first Fmoc-protected amino acids (2eq) dissolving in DCM and N, N-Diisopropylethylamine (DIPEA) (6eq) were added to the swollen resin for 2 hours. The reaction of attaching the first amino acid was conducted twice in order to maximize the load of the resin. In the end, 3 mL methanol was added to block the unreacted CDC resin, then the Fmoc group was removed by 20% piperidine in DMF solution (15min, 2 times).

**Solid-phase peptide synthesis.** After a thorough wash with DCM and DMF, the amino acids (2eq) were coupled to the resin under the conditions of DIC (4eq) and HOBt (4eq) in DMF for 4h. Then, the resin was washed with DMF and DCM, and treated with 20% piperidine in DMF solution again, followed by the coupling with the next amino acid. This cycle was repeated until the desired 9-mer peptides were synthesized. After removing the last Fmoc group, the peptides were cleaved from the resin by the solution of TFA containing phenol, thioanisole, 1,2-EDT, DMSO, and ammonium iodide and purified by the Waters HPLC system.

**Carbon Monoxide (CO) Binding Assay Performed with P450 GpeC and its Mutant.** Spectra were measured at 30°C within the wavelength range of 300 nm to 600 nm using the Thermo Scientific™

GENESYS™ 150 spectrophotometer. The P450 enzyme was diluted to 2.5 μM using Tris-HCl buffer (50 mM, pH 8.0). CO gas was bubbled through the CYP enzyme solution for about 30 seconds at two bubbles per second, after which sodium dithionite was added to reduce the enzyme to its ferrous form. After the reaction was complete, the system was subjected to UV detection to obtain the CO difference spectrum.

**UV−Vis Absorbance Spectroscopy and Analysis of Substrate Binding.** Two cuvettes were prepared, each containing 1 mL of 3 μM P450 solution (50 mM Tris-HCl buffer, pH 8.0). After equilibrating at 30°C, the initial spectrum was measured between 300 and 600 nm. Substrate was then added stepwise to the sample cuvette, while an equal amount of buffer was added to the reference cuvette to maintain equal protein concentrations in both. After a 2-minute equilibration period following each addition, the spectrum was measured, and the peak-to-trough absorbance difference (ΔA) was recorded. The equilibrium dissociation constant (Kd) was determined by fitting the ΔA values against the substrate concentration using the following equation:

$$\frac{\Delta A}{\Delta A_{max}} = \frac{([E] + [S]K_d - \sqrt{\{([E] + [S]K_d)^2 - 4[E][S]\}}}{2[E]}$$

where ΔA is the peak-to-trough absorbance difference, ΔAmax is the maximum absorbance difference, [S] is the substrate concentration, and [E] is the enzyme concentration.

**Preparation of protein samples for protein crystals**

**Bacterial Cloning and Protein Expression.** *Escherichia coli* DH5α was used for gene cloning, while the native enzyme and its mutants were produced by *Escherichia coli* BL21 (DE3). Overlap PCR was employed to remove the 8 C-terminal amino acids of GpeC, resulting in pRSFDuet-GpeC-C8. The GpeC-C8 gene was then cloned and ligated with the linearized pET-28a (+)-sumo vector to obtain ppSumo-Gpec-C8. For convenience, GpeC will be used in place of GpeC-C8 in subsequent descriptions. Any other mutants derived from this were generated by overlap PCR. The DNA endonuclease DpnI for plasmid digestion and the homologous recombination kit for DNA fragment ligation were purchased from Thermo Fisher Scientific. The plasmid extraction and DNA fragment recovery kits were obtained from TIANGEN Biotech (Beijing, China).

For protein expression, the overnight seed culture was diluted 1:50 into 1 L of Terrific Broth (TB) (containing 12 g·$L^{-1}$ tryptone, 24 g·$L^{-1}$ yeast extract, 6 g·$L^{-1}$ glycerol, 2.31 g·$L^{-1}$ $KH_2PO_4$, 16.84 g·$L^{-1}$ $K_2HPO_4 \cdot 3H_2O$), supplemented with 50 μg·$mL^{-1}$ kanamycin. This was then cultured at 37 °C until the $OD_{600}$ reached 0.6-0.8. At this point, 0.2 mM isopropyl β-D-thiogalactoside (IPTG), 0.3 mM δ-aminolevulinic acid (ALA), and 0.2 mM $FeCl_3$ were added to induce protein expression. The culture was then incubated at 18 °C with shaking at 220 rpm for 22 hours.

**Protein purification.** The cells were harvested by centrifugation and suspended in lysis buffer (20 mM Tris-HCl, pH 8.0, 300 mM NaCl) and disrupted using a high-pressure homogenizer (Union Biotech Inc, Shanghai, China). After that, the cellular debris was removed by centrifugation (16,000 × g, 30 min, 4 °C). The soluble cell lysates was loaded onto a 10 ml nickel affinity column (Smart-Lifesciences, Changzhou, China) and thoroughly washed with a gradient of washing buffers (containing 20 mM Tris-HCl, pH 8.0, 300 mM NaCl, and imidazole at concentrations of 5 mM, 10 mM, 20 mM, and 30 mM), subsequently, the target proteins were eluted from the column with the elute buffer (containing 20 mM Tris-HCl, pH 8.0, 300 mM NaCl, and 300 mM imidazole). The eluted peak fractions were further purified by gel filtration chromatography using a Superdex 200 10/300 GL column (Union-Biotech, Shanghai, China) with a buffer of 20 mM Tris-HCl, pH 8.0, 100 mM NaCl, and 5% glycerol. Protein concentration was measured using a NanoDrop™ One/OneC Micro UV-Vis Spectrophotometer (Thermo Fisher, Waltham, USA), and protein purity was assessed by 10-12% sodium dodecyl sulfate-polyacrylamide gel electrophoresis (SDS-PAGE).

**Protein crystallization.** To elucidate such catalytic reactions, we aimed to characterize the structure of GpeC using protein crystallography. However, after several rounds of screening, it has proven challenging to grow high-resolution crystals of the native GpeC protein. In order to obtain high-resolution diffraction data, we have successively conducted some optimization strategies for protein sequences, such as sumo fusion expression and C-terminal truncation. The GpeC protein was crystallized at a concentration of 10-14 mg/mL in SEC buffer with reservoir solution (10% w/v PEG 4000, 20% v/v glycerol, 0.03 M magnesium chloride, 0.03 M calcium chloride, 0.1 M MES/imidazole pH 7.25) by sitting drop vapor diffusion technique. After 7 days of growth at 16 °C, crystals were observed in 0.8 μL sitting drops composed of a 1:1 mixture of protein solution and reservoir solution. The crystals were then cryoprotected by briefly soaking in the corresponding reservoir solution containing an additional 5% (v/v) ethylene glycol before being transferred to liquid nitrogen. Data collection was carried out on the BL18U1 beamline at Shanghai Synchrotron Radiation Facility (SSRF, Shanghai, China).

**Route for synthesizing the biaryl-cyclized tetrapeptide motif**

**Synthesis of (S)-3-(1H-imidazol-4-yl)-2-hydroxy-propanoic acid.** α-hydroxy histidine was synthesized from sodium nitrite in an acidic aqueous medium following the method[4]. Histidine (3.1 g, 20 mmol) was dissolved in 200 mL ($H_2O$: AcOH=8:2), and the solution was cooled to 0 °C. An aqueous solution of $NaNO_2$ (2.77 g, 40 mmol in 40 mL $H_2O$) was added to the solution slowly, the mixture was stirred to room temperature overnight. After the reaction was completed, the solvents were evaporated in vacuo, and the crude product was dissolved in MeOH(100 mL), and the precipitates were filtered, and the filtrate was evaporated in vacuo.

**Synthesis of (S)-3-[1-(2,4-Dinitro-phenyl)-1H-imidazol-4-yl]- 2-hydroxy-propionic Acid.** α-hydroxy histidine(DNP) was synthesized from the previously obtained α-hydroxy histidine following the method[5]. α-hydroxy histidine (1.8g, 11.5 mmol), 2,6-dinitrofluorobenzene (2.14 g, 11.5 mmol), and triethylamine (3.2 mL, 23 mmol) were dissolved in acetonitrile (100 mL). The mixture was allowed to stir at room temperature in the dark for 16 h. Upon completion, acetonitrile was evaporated. The residue was redissolved in water. Excess 2,6-dinitrofluorobenzene was removed by washing the aqueous layer with hexane. The aqueous layer was lyophilized to afford His-OH(DNP) as triethylamine salt. $^{1}$H NMR (400 MHz, DMSO-$d^{6}$) δ 1.19 ($CH_3$- of $Et_3N$ t, J = 5.5, 9H), δ 2.85 (m, 1H), δ 3.03 (-$CH_2$- of $Et_3N$ J = 5.5 Hz), δ 3.25 (m, 1H), δ 4.36 (dd, J = 3.5, 7.9 Hz, 1H), δ 7.89 (s, 1H), δ 8.22 (d, 1H), δ 8,82 (dd, J = 8.8, 1H), δ 9.01 (d, J = 8.8, 1H), δ 9.59 (d, 1H) 13C NMR (101 MHz, DMSO-$d^{6}$) δ 29.9, 68.5, 120.8, 121.6, 129.5, 131.3, 131.8, 132.9, 136.5, 144.1, 148.3, 174.0.

**Solid phase synthesis of ether-bonded (43).** 0.2 mmol CTC resin (1.08mmol/g, CSBio) was swelled in DCM for 20 minutes before washing the resin with DCM 3 times. Then, 147 mg compound **43** (0.46 mmol) was dissolved in 5 mL DCM, and 200 μL DIPEA (1.15 mmol) was added to the swollen resin overnight. In the end, 3 mL of methanol was added to block the unreacted CTC resin. After a thorough wash with DCM and DMF, the Fmoc-Tyrosine(Tbu) (2eq) was coupled to the resin under the condition of DIPEA (4eq) and Pybop (4eq) in DMF for 12 hours, followed by cleavage with 3 mL 20% hexafluoroisopropanol in DCM and purified by the Waters HPLC system. Light yellow (**37**) was obtained after lyophilized. And (**43**) was synthesized by the abovementioned SPPS method.

**Biaryl bond building and flanking residues cleavage (44, 45).** The biaryl bond of Tryptophan and Tyrosine was synthesized with the P450 enzyme GpeC with the abovementioned in vitro reaction method to give compound (**44**). An equivalent volume of methanol was added after the reaction, and the precipitate was removed through centrifugation. Methanol was then removed in vacuo, and trypsin was added to digest WR on the N-terminal for 2 hours at 37 °C. NaOH and methanol were added to the solution containing a 1:1 $H_2O$: $CH_3OH$ for 1 hour to hydrolyse the ether bond, yielding the final product (**45**).

#### Supplementary Tables

**Supplementary Table 1.** Primers used in this study.

| Primers | Sequences (5’ → 3’) |
|---|---|
| $GpeA_2$_F | aaaacctgtattttcagggcatgcgcaccctcggtctg |
| $GpeA_2$_R | cattatgcggccgcaagcttccatgctattacatgtgatgataaagca |
| GpeC_F | tataagaaggagatatacatgtgaaatttgaaccttcagatgccc |
| GpeC_R | gtttctttaccagactcgagtcatgccagctgagcatg |
| HisGpeC_F | atcaccacagccaggatccgatgaaatttgaaccttcagatg |
| HisGpeC_R | cattatgcggccgcaagctttcatgccagctgagcatgg |
| pRSF_His_R | cggatcctggctgtggtgatgatggt |
| pRSF_Mid_F | aagcttgcggccgcataatgctt |
| pRSF_Sumo_TEV_R | gccctgaaaatacaggttttcg |
| pRSF_Mid_R | atgtatatctccttcttatacttaactaatatactaagatgg |
| pRSF_MCS2_F | ctcgagtctggtaaagaaaccgctgc |
| GpeC_C325_F | catccaatttgactctggagttggtttccgagcttc |
| GpeC_C324G_R | ccagagtcaaattggatgccacgcccttatgttc |
| GpeC_C324H_R | ccagagtcaaattggatgccacgtgcttatgttccc |
| GpeC_N34A_F | tacctgaccgacctagacgatgctaaggcc |
| GpeC_N34A_R | atcgtctaggtcggtcaggtaataagcttgttcaggaaattcaataatg |
| GpeC_Y36A_F | tagacgatgctaaggccgtgctggcc |
| GpeC_Y36A_R | cttagcatcgtctaggtcggtcagagcataattttgttcaggaaattcaat |
| GpeC_H57G_F | tttaccagtcgaggcttgttacagccgcg |
| GpeC_H57G_R | caagcctcgactggtaaaaaagaaatcagagccagggtgataattttcatctttgttggcc |
| GpeC_D59A_F | tttaccagtcgaggcttgttacagccgcg |
| GpeC_D59A_R | caagcctcgactggtaaaaaagaaagcagagtgagggtgataattttcatctttgttgg |
| GpeC_F60A_F | caccctcactctgatgcttttttaccagtcg |
| GpeC_F60A_R | atcagagtgagggtgataattttcatctttgttggc |
| GpeC_F62G_F | cttgttacagccgcgcagccgt |
| GpeC_F62G_R | ctgcgcggctgtaacaagcctcgactggtgccaaagaaatcagagtgagg |
| GpeC_R152A_F | gtgctatccggcaaagctcttgagcacg |
| GpeC_R152A_R | tttgccggatagcactgagcgatgaacg |
| GpeC_R160A_F | tatcgccagcgcaactatcggctgttaaaaaaggag |
| GpeC_R160A_R | atagttgcgctggcgatatagccaagcacgccagcctccgtgc |
| GpeC_Y163A_F | gctggcgtcgctggctagctcgccagcgcaac |
| GpeC_Y163A_R | tagccagcgacgccagcctccg |
| GpeC_D207A_F | gataccgaacttgcagctatttttctgtc |
| GpeC_D207A_R | ctgcaagttcggtatcgctg |
| GpeC_F214A_F | gatatttttctgtcttttcttgctgccactaccggttc |
| GpeC_F214A_R | aagaaaagacagaaaaatatctgcaagttcg |

| | |
|---|---|
| GpeC_S219A_F | tgggtttttgcctgagctggtgtctgc |
| GpeC_S219A_R | gctcaggcaaaaacccagagcaccggtagtggcaaaa |
| GpeC_A256F_F | gactttggcctgtgttttggaatctggc |
| GpeC_A256F_R | cacaggccaaagtcgcagcg |
| GpeC_W257F_F | atctggccaggatgcctgcaaaggac |
| GpeC_W257F_R | gcaggcatcctggccagattgaaggccacagg |
| GpeC_W257G_F | atctggccaggatgcctgcaaaggac |
| GpeC_W257G_R | ggcatcctggccagattgccggccacaggc |
| GpeC_N258G_F | gccaggatgcctgcaaaggaccatcaattgc |
| GpeC_N258G_R | ctttgcaggcatcctggccaggccccaggccacaggccaaag |
| GpeC_N258L_F | gccaggatgcctgcaaaggaccatcaattgc |
| GpeC_N258L_R | ctttgcaggcatcctggccagcagccaggccacaggccaaag |
| GpeC_A361F_F | cgccctcttgcccagtttgcattggctc |
| GpeC_A361F_R | ctgggcaagagggcgctcgc |

**Supplementary Table 2.** DNA sequences used in this study.

| |
|---|
| **>TRX-Spinach ferredoxin** |
| atgagcgataaaattattcacctgactgacgacagttttgacacggatgtactcaaagcggacggggcgatcctcgtcgatttctgggcagagtg<br>gtgcggtccgtgcaaaatgatcgccccgattctggatgaaatcgctgacgaatatcagggcaaactgaccgttgcaaaactgaacatcgatcaa<br>aaccctggcactgcgccgaaatatggcatccgtggtatcccgactctgctgctgttcaaaaacggtgaagtggcggcaaccaaagtgggtgca<br>ctgtctaaaggtcagttgaaagagttcctcgacgctaacctggccggttctggttctggccatatggaattcgccgcctacaaggtgaccctggtg<br>accccgaccggcaacgtggagttccagtgcccggacgacgtgtacatcctggacgccgcagaagaagaggggatcgacctgccgtacagct<br>gtcgcgcaggcagttgcagcagctgcgcaggtaagctgaaaaccggcagcctgaaccaggacgaccagagcttcctggacgacgaccaga<br>tcgacgagggctgggttctgacatgcgccgcctacccggtgagcgatgtgaccatcgagacccacaaagaagaggagctgaccgcctaa |
| **>Spinach ferredoxin reductase** |
| gaattccagatcgcctctgatgtggaggcacctccacctgctcctgctaaggtagagaaacattcaaagaaaatggaggaaggcattacagtta<br>acaagtttaagcctaagaccccttacgttggaagatgtcttcttaacaccaaaattactggggatgatgcacccggagagacctggcacatggttt<br>tttcccatgaaggagagatcccttacagagaagggcaatccgttggggttattccagatggggaagacaagaatggaaagccccataagttga<br>gattgtactcgatcgccagcagtgctcttggtgattttggtgatgctaaatctgtttcgttgtgtgtaaaacgactcatctacaccaatgacgctggag<br>agacgatcaagggagtctgctccaacttcttgtgtgacttgaaacccggtgctgaagtgaagttaacaggaccagttggaaaggagatgctcat<br>gcccaaagaccctaacgcgacaattatcatgcttggaactggaactgggattgctcctttccgttcattcttgtggaagatgttcttcgaaaagcat<br>gatgattacaagtttaacggcttggcttggcttttcttgggtgtacccacaagcagttctcttctctacaaagaggaatttgagaagatgaaggaaaa<br>ggctccagacaacttcaggctggattttgcagtgagcagagagcaaactaacgagaaaggggagaagatgtacattcaaacccgaatggcac<br>aatacgcagttgagctatgggaaatgttgaagaaagataatacttatttctacatgtgtggtctcaagggaatggaaaagggaattgacgacattat<br>ggtttcattggctgctgcagaaggcattgattggattgaatacaagaggcagttgaagaaggcagaacaatggaacgttgaagtctactaa |
| **>$GpeA_2$** |
| atgcgcaccctcggtctgcgtaatcaagccgtgtgctatagcaaaaaccctagctggttttggatgctttatcatcacatgtaa |
| **>GpeC** |
| gtgaaatttgaaccttcagatgccctttttaaagacccttatcattttctggacctgggactggatgccggaagtgacattattgaatttcctgaacaa<br>aattattacctgaccgacctagacgatgctaaggccgtgctggccaacaaagatgaaaattatcaccctcactctgatttctttttaccagtcgagg<br>cttgttacagccgcgcagccgtcaggtggacatggccagaggagctgtaaagtttcttcaacagcaccttaatgcaacaaggcggctgctacca<br>gaaagggtcaaggagcagttacagggacggcacagttggccggatatctccaacttgtttatgttctcacttttcgagccggcattgttggatgct<br>cggggtgataaaaccctaaaaaattgtcttgatgaaatcgttcatcgctcagtgctatccggcaaacggcttgagcacggaggctggcgtcgctg<br>gctatatcgccagcgcaactatcggctgttaaaaaaggagatccgcgcgcgacgccgaggactggtggagtgtcaggatgtaatgtccgcttta<br>gttggagctgcggaccacgggatcagcgataccgaacttgcagatatttttctgtcttttctttttgccactaccggttctctgggtttttgcctgagct<br>ggtgtctgcatctggtcagcgaagcccccgaaaaagctaaacttccttcccagtggctagtgcttgaggcgctgcgactttggcctgtggcctgg<br>aatctggccaggatgcctgcaaaggaccatcaattgccttcgggccataaggtcagcaacaacagcgtggtgatcgcatgcccctactctgttc<br>aacgaaaccccaagcactggcctgacgccgaccgcttcttaccggagcgctggcaggataatagcaatcgccagcgtttacttgcctttgggtg<br>gggggaacataagtgcgtggcatccaatttgactctggagttggtttccgagcttctggatatcattttgggaaattaccatctctcctgcgaatact<br>caggcgagcgccctcttgcccaggcggcattggctcccccatcttttacgttgactctctcagccaagacggaacacaaccatgctcagctggc<br>atga |

**Supplementary Table 3.** NMR spectroscopic data for Gpertide (**3**)

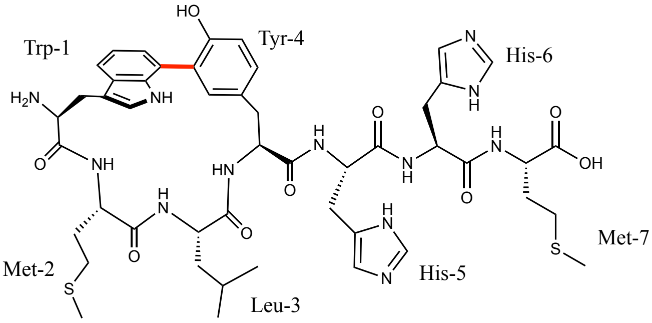


| Residue | Position | $\delta_C$, mult. | $\delta_H$ ($J$ in Hz) | Residue | Position | $\delta_C$, mult. | $\delta_H$ ($J$ in Hz) |
|---|---|---|---|---|---|---|---|
| Trp-1 | $NH_2$ | | 8.33, s | | 2 | 134.1 | 7.52, s |
| | $\alpha$ | 53.0 | 3.74, m | | 3 | 124.8 | - |
| | $\beta$ | 27.4 | 2.96,3.18,m | | 4 | 152.8 | - |
| | 1 | - | 10.73, s | | 4-OH | - | 9.08, s |
| | 2 | 123.5 | 6.59, d (2.3) | | 5 | 115.7 | 6.75, s |
| | 3 | 106.3 | - | | 6 | 129.3 | 6.75, s |
| | 3a | 126.8 | - | His-5 | NH | - | 8.40, d (7.85) |
| | 4 | 115.7 | 7.42, dd (6.50, 2.34 ) | | $\alpha$ | 51.9 | 4.65, m |
| | 5 | 117.8 | 7.04, m | | $\beta$ | 27.9 | 3.04 m |
| | 6 | 122.1 | 7.05, m | | 1 | | |
| | 7 | 123.1 | - | | 2 | 116.5 | 7.25, m |
| | 7a | 135.1 | - | | 3 | - | - |
| | C=O | 167.4 | - | | 4 | 128.6 | 7.23, m |
| Met-2 | NH | - | 7.66, d (8.96) | | 5 | - | - |
| | $\alpha$ | 50.6 | 4.41, m | | C=O | 170.1 | - |
| | $\beta$ | 31.5 | 1.58,1.50, m | His-6 | NH | - | 8.41, d (7.85) |
| | γ | 29.3 | 2.32, m | | $\alpha$ | 51.9 | 4.70, m |
| | δ | 14.6 | 2.01, s | | $\beta$ | 27.5 | 3.04, m |
| | C=O | 169.1 | - | | 1 | | |
| Leu-3 | NH | - | 7.85, s | | 2 | 116.1 | 7.23, m |
| | $\alpha$ | 52.2 | 4.17, m | | 3 | - | - |
| | $\beta$ | 40.4 | 1.28,1.49,m | | 4 | 128.1 | 7.24, m |
| | γ | 23.7 | 1.80, m | | 5 | - | - |
| | δ-Me1 | 21.9 | 0.93, d (6.47) | | C=O | 170.0 | - |
| | δ-Me2 | 22.8 | 0.94, d (6.63) | Met-7 | NH | - | 8.62, brs |
| | C=O | 171.4 | - | | $\alpha$ | 50.3 | 4.36, m |
| Tyr-4 | NH | - | 8.23, d (8.79) | | $\beta$ | 30.4 | 2.00,1.86,m |
| | $\alpha$ | 51.6 | 4.70, m | | γ | 29.0 | 2.43,2.32,m |
| | $\beta$ | 37.3 | 2.96,2.61,m | | δ | 14.3 | 1.99, s |
| | 1 | 126.8 | - | | C=O | 173.3 | - |

**Supplementary Table 4.** NMR spectroscopic data for Typertide (**7**)

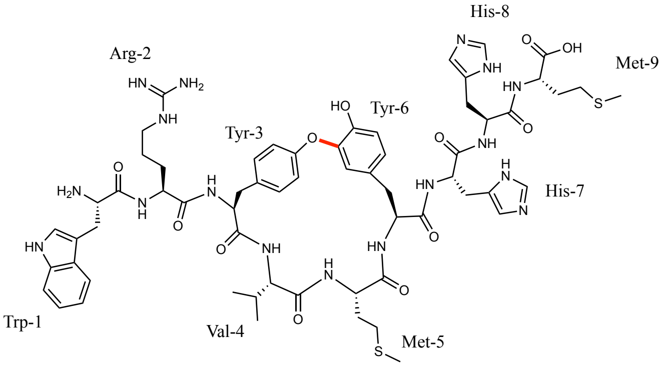


| Residue | Position | $\delta_C$, mult. | $\delta_H$ (*J* in Hz) | Residue | Position | $\delta_C$, mult. | $\delta_H$ (*J* in Hz) |
|---|---|---|---|---|---|---|---|
| Trp-1 | $NH_2$ | - | 8.01, s | | γ | 28.9 | 2.38, m |
| | *α* | 52.5 | 4.10, m | | δ | 14.5 | 1.98, s |
| | *β* | 27.5 | 3.04, 3.25, m | | C=O | 170.1 | |
| | 1 | - | 11.03, d (2.00) | Tyr-6 | NH | - | 7.75, m |
| | 2 | 125.1 | 7.22, d (2.42) | | *α* | 52.2 | 4.67, m |
| | 3 | 106.7 | - | | *β* | 36.5 | 2.77, 2.70, m |
| | 3a | 127.1 | - | | 1 | 127.7 | - |
| | 4 | 118.2 | 7.70, d (8.04) | | 2 | 120.8 | 7.08, s |
| | 5 | 118.4 | 7.02, t (7.57) | | 3 | 143.5 | - |
| | 6 | 120.8 | 7.10, t (7.35) | | 4 | 147.0 | - |
| | 7 | 111.5 | 7.37, d (8.16) | | 4-OH | - | 9.18, s |
| | 7a | 136.4 | - | | 5 | 116.2 | 6.73, d (8.26) |
| | C=O | 168.4 | - | | 6 | 125.5 | 6.66, d (8.12) |
| Arg-2 | NH | - | 8.85, d (7.81) | | C=O | 170.9 | - |
| | *α* | 52.4 | 4.44, m | His-7 | NH | - | 8.26, d (7.66) |
| | *β* | 29.5 | 1.66, m | | *α* | 51.6 | 4.60, m |
| | γ | 25.0 | 1.54, 1.48, m | | *β* | 27.1 | 3.09, 3.00, m |
| | δ | 40.4 | 3.09, m | | 1 | - | - |
| | C=O | 170.1 | - | | 2 | 129.1 | - |
| Tyr-3 | NH | - | 8.38, d (6.99) | | 3 | - | - |
| | *α* | 53.9 | 4.54, m | | 4 | 133.9 | - |
| | *β* | 37.0 | 2.90, 2.79, m | | 5 | 116.8 | 7.27, s |
| | 1 | 130.7 | - | | C=O | 170.1 | |
| | 2 | 129.7 | 7.04, d (6.78) | His-8 | NH | - | 8.43, d (6.91) |
| | 3 | 117.8 | 6.84, d (8.52) | | *α* | 51.7 | 4.60, m |
| | 4 | 155.6 | - | | *β* | 27.1 | 3.02, 2.93, m |
| | 4-OH | - | - | | 1 | - | - |
| | 5 | 117.8 | 6.84, d (8.52) | | 2 | 129.1 | - |
| | 6 | 129.7 | 7.04, d (6.78) | | 3 | - | - |
| | C=O | 169.8 | - | | 4 | 133.8 | - |
| Val-4 | NH | - | 7.97, d (9.5) | | 5 | 116.7 | 7.33, s |
| | *α* | 58.0 | 4.05, m | | C=O | 170.3 | - |
| | *β* | 30.5 | 1.86, m | Met-9 | NH | - | 8.50, d (7.72) |
| | γ-Me1 | 18.2 | 0.75, d (6.62) | | *α* | 51.1 | 4.33, m |
| | γ-Me2 | 19.1 | 0.79, d (6.65) | | *β* | 30.5 | 1.84, m |
| | C=O | 169.9 | - | | γ | 29.5 | 2.46, m |
| Met-5 | NH | - | 7.72, m | | δ | 14.5 | 2.01, s |
| | *α* | 53.0 | 4.10, m | | C=O | 173.1 | - |
| | *β* | 31.9 | 1.76, m | | | | |

**Supplementary Table 5.** NMR spectroscopic data for Gpentide (**8**)

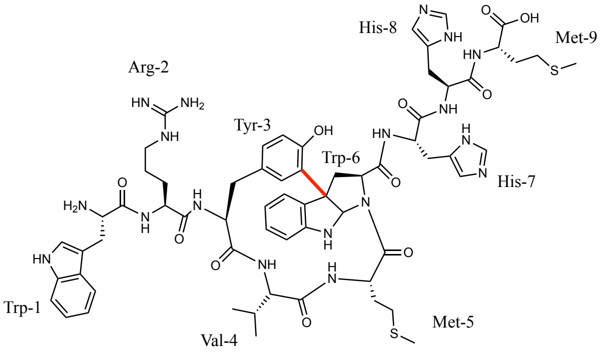


| Residue | Position | $\delta_C$, mult. | $\delta_H$ ($J$ in Hz) | Residue | Position | $\delta_C$, mult. | $\delta_H$ ($J$ in Hz) |
|---|---|---|---|---|---|---|---|
| Trp-1 | $NH_2$ | - | 7.99, d (5.31) | | δ | 14.5 | 2.00, s |
| | *α* | 52.3 | 4.07, m | | C=O | - | - |
| | *β* | 27.6 | 3.26, 3.03, m | Trp-6 | NH | - | - |
| | 1 | - | 11.04, d (2.43) | | *α* | 60.2 | 4.10, m |
| | 2 | 125.2 | 7.25, d (2.49) | | *β* | 39.0 | 2.86, 2.70 m |
| | 3 | 106.7 | - | | 1 | - | - |
| | 3a | 127.0 | - | | 2 | 81.5 | 5.94, s |
| | 4 | 118.6 | 7.74, d, (7.84) | | 3 | 59.3 | - |
| | 5 | 118.5 | 7.02, t (7.49) | | 3a | 132.1 | - |
| | 6 | 121.2 | 7.10, t (7.78) | | 4 | 125.2 | 7.58, d (7.63) |
| | 7 | 111.5 | 7.37, d (7.98) | | 5 | 117.8 | 6.46, t (7.62) |
| | 7a | 136.4 | - | | 6 | 128.1 | 6.93, t (7.74) |
| | C=O | 168.5 | - | | 7 | 108.9 | 6.57, d (7.69) |
| Arg-2 | NH | - | 8.84, d (7.85) | | 7a | 148.3 | - |
| | *α* | 51.6 | 4.46, m | | $C^6$=O | - | - |
| | *β* | 29.5 | 1.50, m | His-7 | NH | - | 8.05, s |
| | γ | 25.1 | 1.47, 1.42, m | | *α* | 51.5 | 4.39, m |
| | δ | 40.4 | 3.06, m | | *β* | 27.5 | 2.93, m |
| | C=O | 170.4 | | | 1 | - | 8.93, s |
| Tyr-3 | NH | - | 7.75, d, ol | | 2 | 129.2 | - |
| | *α* | 53.0 | 4.64, m | | 3 | - | - |
| | *β* | 36.6 | 2.99, m | | 4 | 133.7 | - |
| | 1 | | - | | 5 | 116.5 | 7.23, s |
| | 2 | 127.0 | 6.50, s | | C=O | 169.9 | - |
| | 3 | 126.6 | - | His-8 | NH | - | 8.25, d (7.39) |
| | 4 | 153.7 | - | | *α* | 51.7 | 4.50, m |
| | 4-OH | - | 9.89, s | | *β* | 27.5 | 3.04, m |
| | 5 | 116.5 | 6.82, t (8.22) | | 1 | - | 8.93, s |
| | 6 | 130.8 | 6.81, t (8.44) | | 2 | 128.8 | - |
| | C=O | 170.0 | - | | 3 | - | - |
| Val-4 | NH | - | 8.16, d (8.63) | | 4 | 133,7 | - |
| | *α* | 60.7 | 4.03, m | | 5 | 116.8 | 7.33, s |
| | *β* | 29.0 | 1.95, m | | C=O | 169.9 | - |
| | γ-Me1 | 19.2 | 0.89, d (6.65) | Met-9 | NH | - | 8.34, d (7.69) |
| | γ-Me2 | 19.1 | 0.88, d (6.65) | | *α* | 51.0 | 4.29, m |
| | C=O | 171.4 | - | | *β* | 30.6 | 1.95, m |
| Met-5 | NH | - | 8.70, d (7.94) | | γ | 29.5 | 1.64, m |
| | *α* | 51.0 | 4.78, m | | δ | 14.7 | 2.03, s |
| | *β* | 31.7 | 2.07, m | | C=O | 172.9 | - |
| | γ | 29.3 | 1.51, m | | | | |

**Supplementary Table 6.** NMR spectroscopic data for the biocatalyzed 'WHHY' (**42**) module.

| GpeC catalyzed biaryl module of Rubrin in DMSO-$d^6$ | | | | Rubrin in $D_2O$ | | | |
|---|---|---|---|---|---|---|---|
| **Residue** | **Position** | **$\delta_C$, mult.** | **$\delta_H$ (*J* in Hz)** | **Residue** | **Position** | **$\delta_C$, mult.** | **$\Delta\delta c$** |
| Trp-1 | $NH_2$ | - | 8.46, brs | Trp-1 | $NH_2$ | - | - |
| | $\alpha$ | 53.5 | 3.76, m | | $\alpha$ | 53.1 | 0.4 |
| | $\beta$ | 27.6 | 3.24, 2.95, m | | $\beta$ | 27.6 | 0 |
| | 1 | - | 10.59, s | | 1 | - | - |
| | 2 | 123.9 | 6.64, d (2.55) | | 2 | 124.5 | 0.6 |
| | 3 | 106.4 | - | | 3 | 106.8 | 0.4 |
| | 3a | 126.9 | - | | 3a | 126.1 | 0.8 |
| | 4 | 116.2 | 7.46, m | | 4 | 117.4 | 1.2 |
| | 5 | 118.2 | 7.05, m | | 5 | 119.4 | 1.2 |
| | 6 | 122.6 | 7.06, m | | 6 | 122.9 | 0.3 |
| | 7 | 123.0 | - | | 7 | 122.4 | 0.6 |
| | 7a | 134.8 | - | | 7a | 135.7 | 0.9 |
| | C=O | 167.6 | - | | C=O | 168.7 | 1.1 |
| His-2 | NH | - | 7.89, d (8.97) | His-2 | NH | - | - |
| | $\alpha$ | 50.7 | 4.57, m | | $\alpha$ | 51.1 | 0.4 |
| | $\beta$ | 26.7 | 2.78, m | | $\beta$ | 25.8 | 0.9 |
| | 1 | - | - | | 1 | - | - |
| | 2 | 133.7 | - | | 2 | 133.4 | 0.1 |
| | 3 | - | - | | 3 | - | - |
| | 4 | 128.7 | - | | 4 | 127.5 | 1.2 |
| | 5 | 117.1 | - | | 5 | 117.2 | 0.1 |
| | C=O | 168.5 | - | | C=O | 169.4 | 0.9 |
| His-3 | NH | - | 8.36, brs | His-3 | NH | - | - |
| | $\alpha$ | 53.2 | 4.48, m | | $\alpha$ | 54.9 | 1.7 |
| | $\beta$ | 26.9 | 2.94, m | | $\beta$ | 25.8 | 1.1 |
| | 1 | - | 8.87, s | | 1 | - | - |
| | 2 | 133.7 | - | | 2 | 133.8 | 0.1 |
| | 3 | - | - | | 3 | - | - |
| | 4 | 129.2 | - | | 4 | 127.2 | 2.0 |
| | 5 | 117.1 | | | 5 | 117.6 | 0.5 |
| | C=O | 170.1 | | | C=O | 170.6 | 0.5 |
| Tyr-4 | NH | - | 8.30, brs | Tyr-4 | NH | - | - |
| | $\alpha$ | 51.5 | 4.70, m | | $\alpha$ | 52.4 | 0.9 |
| | $\beta$ | 37.3 | 2.93, 2.70, m | | $\beta$ | 36.4 | 0.9 |
| | 1 | 126.5 | - | | 1 | 127.0 | 0.5 |
| | 2 | 133.7 | 7.45, s | | 2 | 133.5 | 0.2 |
| | 3 | 124.7 | - | | 3 | 125.0 | 0.3 |
| | 4 | 153.0 | - | | 4 | 151.9 | 1.1 |
| | 4-OH | - | 9.15, s | | 4-OH | - | - |
| | 5 | 116.0 | 6.73, m | | 5 | 116.2 | 0.2 |
| | 6 | 129.4 | 6.72, m | | 6 | 130.9 | 1.5 |
| | C=O | 170.1 | - | | C=O | 170.9 | 0.8 |

**Supplementary Table 7.** Crystal data collection and refinement statistics

| **Crystal Parameters** | **Gpec$_{F62G}$ holo** | **Crystal Parameters** | |
|---|---|---|---|
| PDB ID | 9ULW | **Refinement** | |
| Space group | P $4_32_12$ | Resolution range(Å) | 30.17 - 2.62 (2.714 - 2.62) |
| **Cell dimensions** | | Reflections used in refinement | 31728 (3088) |
| a,b,c (Å) | 78.40, 78.40, 326.90 | $R_{work}$[c] | 0.2287 (0.3333) |
| α,β,γ (˚) | 90, 90, 90 | $R_{free}$[d] | 0.2651 (0.3825) |
| **Data Collection** | | RMS bonds (Å) | 0.007 |
| Wavelength (Å) | 0.96183 | RMS angles (°) | 1.02 |
| Resolution (Å) | 30.90-2.62 (2.74-2.62) | Protein residues | 734 |
| No. of measured reflections | 812157 (96096) | **No. of non-hydrogen atom/B-factor(Å$^2$)** | |
| No. of unique reflections | 31770 (3752) | All/average | 5955 / 70.56 |
| Completeness (%) | 99.8 (99.6) | Macromolecules | 5836 / 70.77 |
| Multiplicity | 25.6 (25.6) | Ligands | 86 / 61.05 |
| Mean I/σ(I) | 20.8 (2.6) | Solvent | 33 / 57.49 |
| $CC_{1/2}$ | 0.999 (0.868) | **Ramachandran plot** | |
| $R_{merge}$(%)[b] | 15.8 (201.3) | Favored (%) | 96.11 |
| $R_{meas}$(%)[b] | 16.4 (209.1) | Allowed (%) | 3.89 |
| $R_{pim}$(%)[b] | 4.4 (56.4) | Outliers (%) | 0.00 |

#### Supporting Figures

##### Figure S1. General genome mining of RiPP-P450s

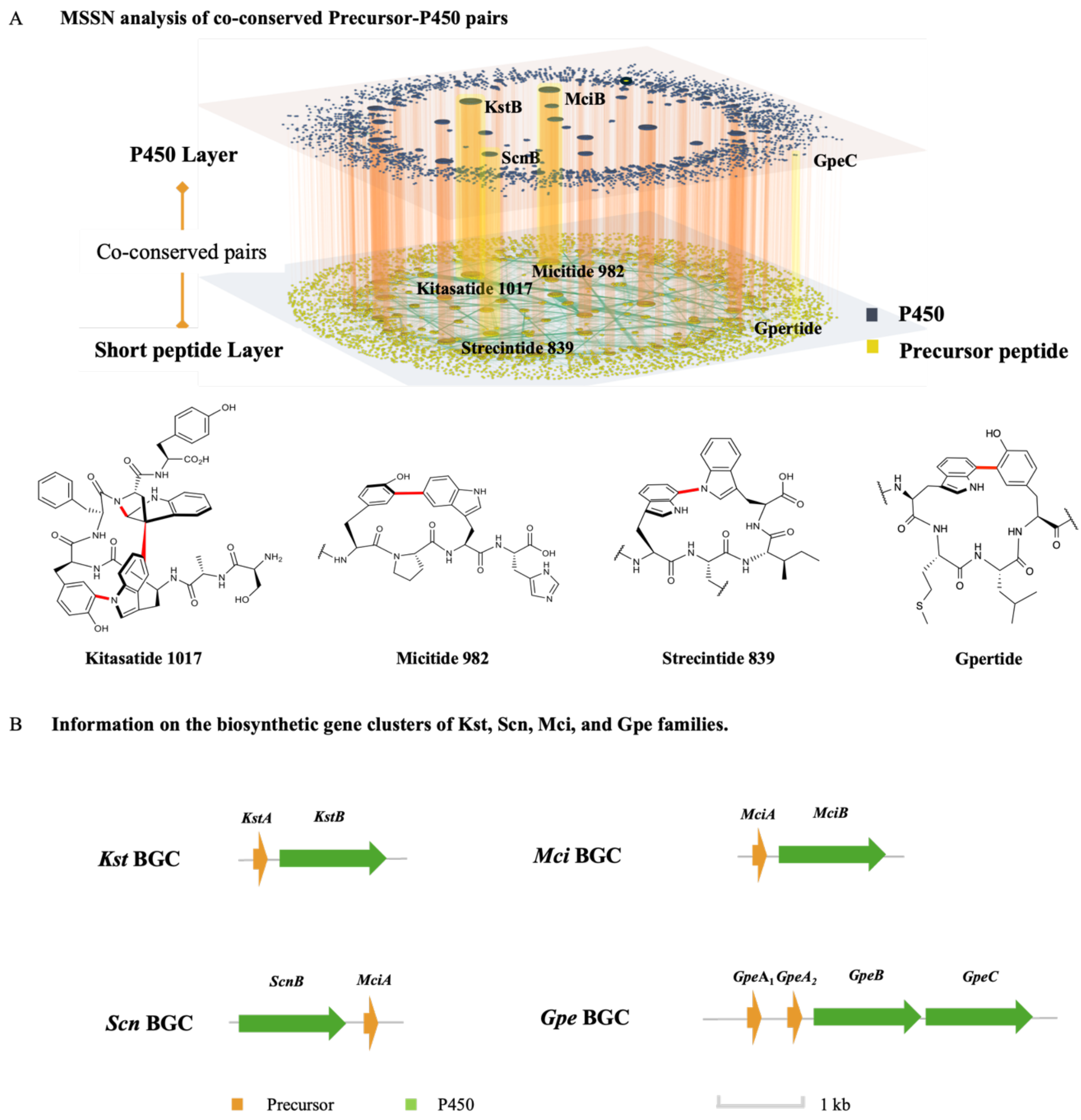


A) Speco-base genome mining workflow and visualization by multi-sequence-similarity network. The upper layer represented the mined P450 enzymes, as the lower layer represented the co-conserved short peptides within the biosynthesis gene cluster. B) Enzyme families that were selected for this work. As catalyzed structures had been elucidated in vivo, we aimed to characterize the P450s' functions in vitro.

**Figure S2.** Bioinformatic analysis of GpeC analogs.

**A. Phylogenetic tree of GpeC related RiPP-P450s**

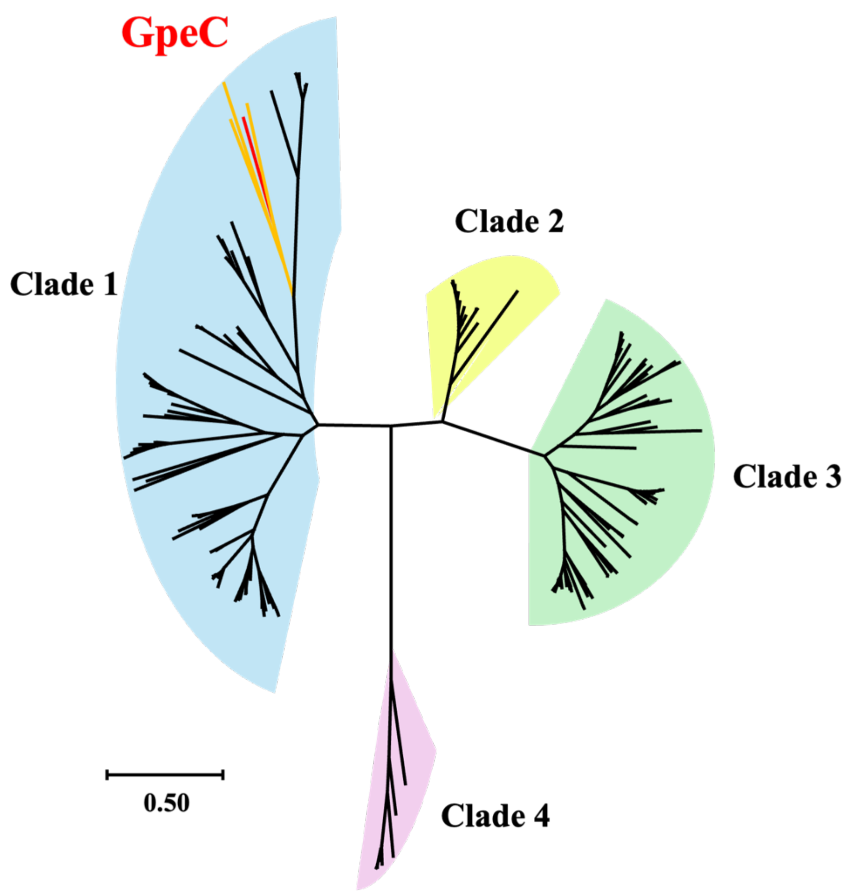


**B. Precursor logo sequence of corresponding P450 clades**

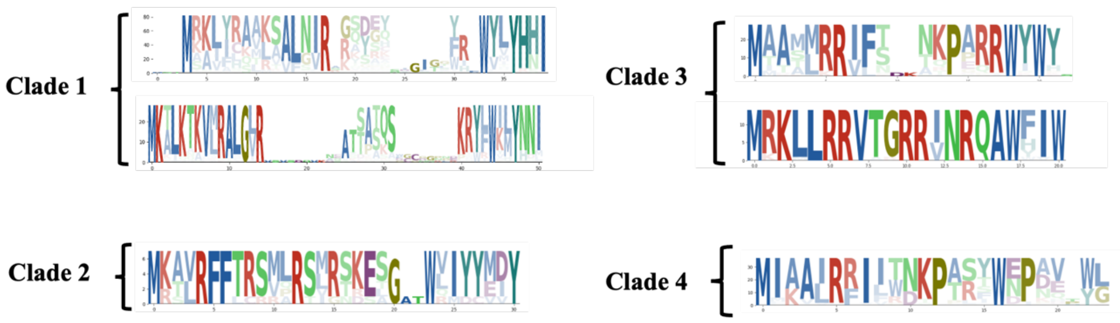


A) Phylogenetic tree of mined P450 enzymes with sequence similarity higher than 0.4 with GpeC. B) Four clades formed with corresponding predicted precursor peptides were clustered, and sequence logos were made. In clade 1, which contains GpeC, precursor sequences are diverse, as a pattern was mainly maintained with W/Y-X-WXXY-H/N-H/N-I.

**Figure S3.** Expression systems used in this work.

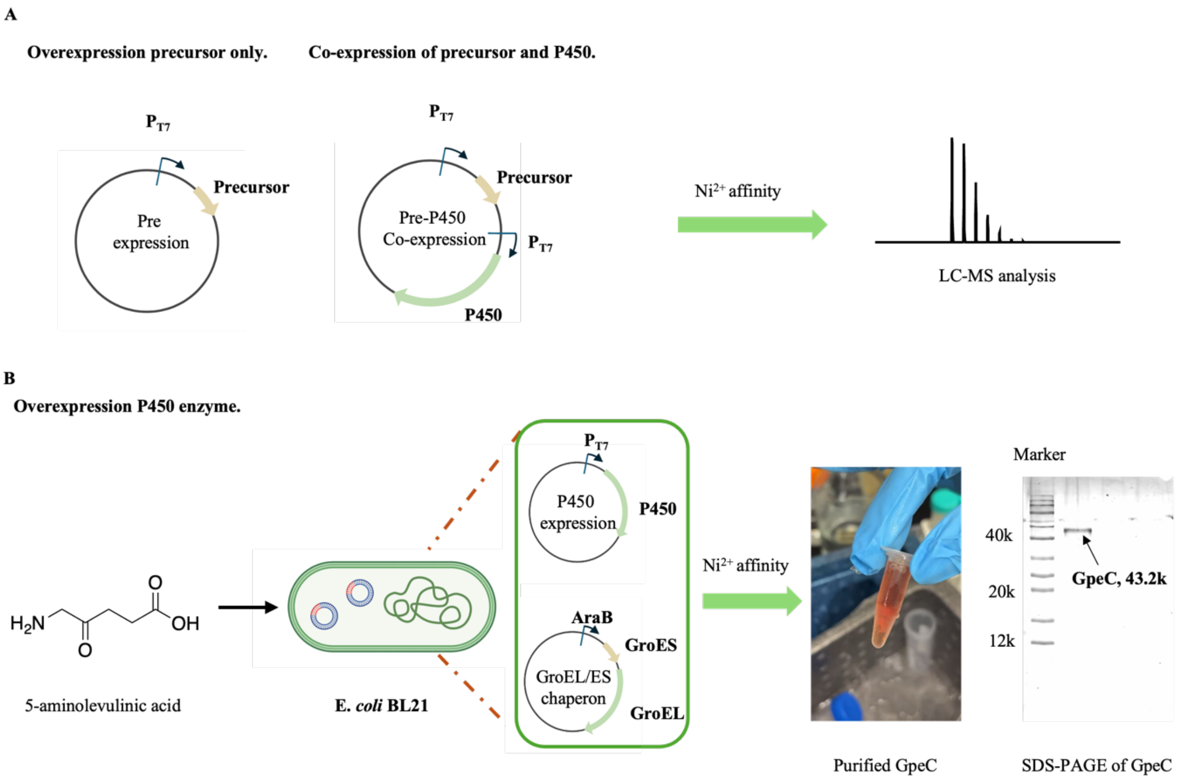


A) Precursor expression was performed on the pRSF-Duet1 plasmid with the T7 promoter. For the co-expression of precursor and P450 enzyme in vivo, precursor and P450 enzyme sequences were separately cloned into the MCS1 and MCS2 sites on the plasmid. Each MCS site was promoted by a T7 promoter. We used the $Ni^{2+}$ affinity column to purify the 6*His fused precursor peptides. B) For expression of P450 enzymes, an *Escherichia coli* BL21(DE3) containing GroEL/ES chaperon plasmid was made, and pRSF-Duet1 containing the P450 sequence was introduced to the system. Expression was performed with the introduction of 5-ALA for the heme biosynthesis.

**Figure S4** Carbon monoxide binding assay performed with $P450_{GpeC}$ and $P450_{GpeC\text{-}F62G}$

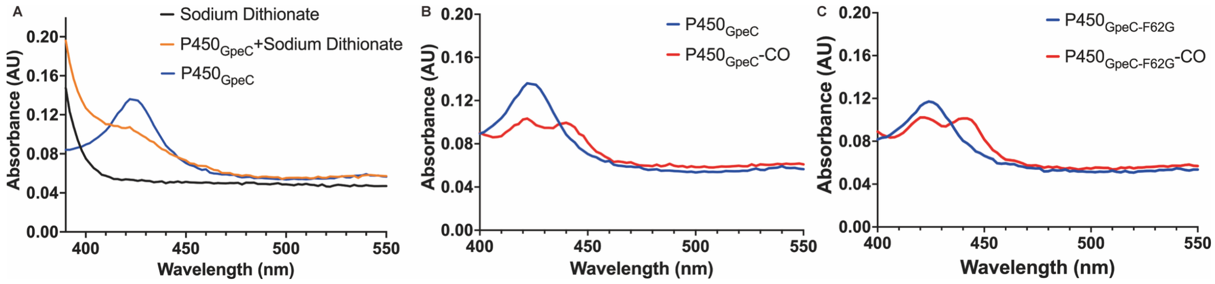


A) The UV-Vis spectra of sodium dithionite and $P450_{GpeC}$ without CO treatment. B) and C) The UV-Vis spectra of $P450_{GpeC}$/$P450_{GpeC\text{-}F62G}$ with/without CO treatment. The UV-Vis spectra of sodium dithionite (black), $P450_{GpeC}$/$P450_{GpeC\text{-}F62G}$ (blue),($Fe^{3+}$) $P450_{GpeC}$ treated with sodium dithionite (orange), and the $Fe^{2+}$-CO complex (red), which was generated by gently bubbling CO through the P450 solution for ~30 seconds.

**Figure S5.** *In vitro* assay of $GpeA_2$ with purified His-GpeC. A 5-hour reaction was performed before quenching by adding an equivalent volume of methanol. Methanol was removed with air, and trypsin was added to digest the catalytic product to NPSWFWMLYHHM. Conversion of the 27-mer peptide was analyzed by the square of the linear peptide (**1**) and the modified one(**2**). Tandem mass analysis of (**2**) revealed the peptide ions of $y_3$, $b_5$, and $y_7$-2H, proving the modification happened within the WMLY motif.

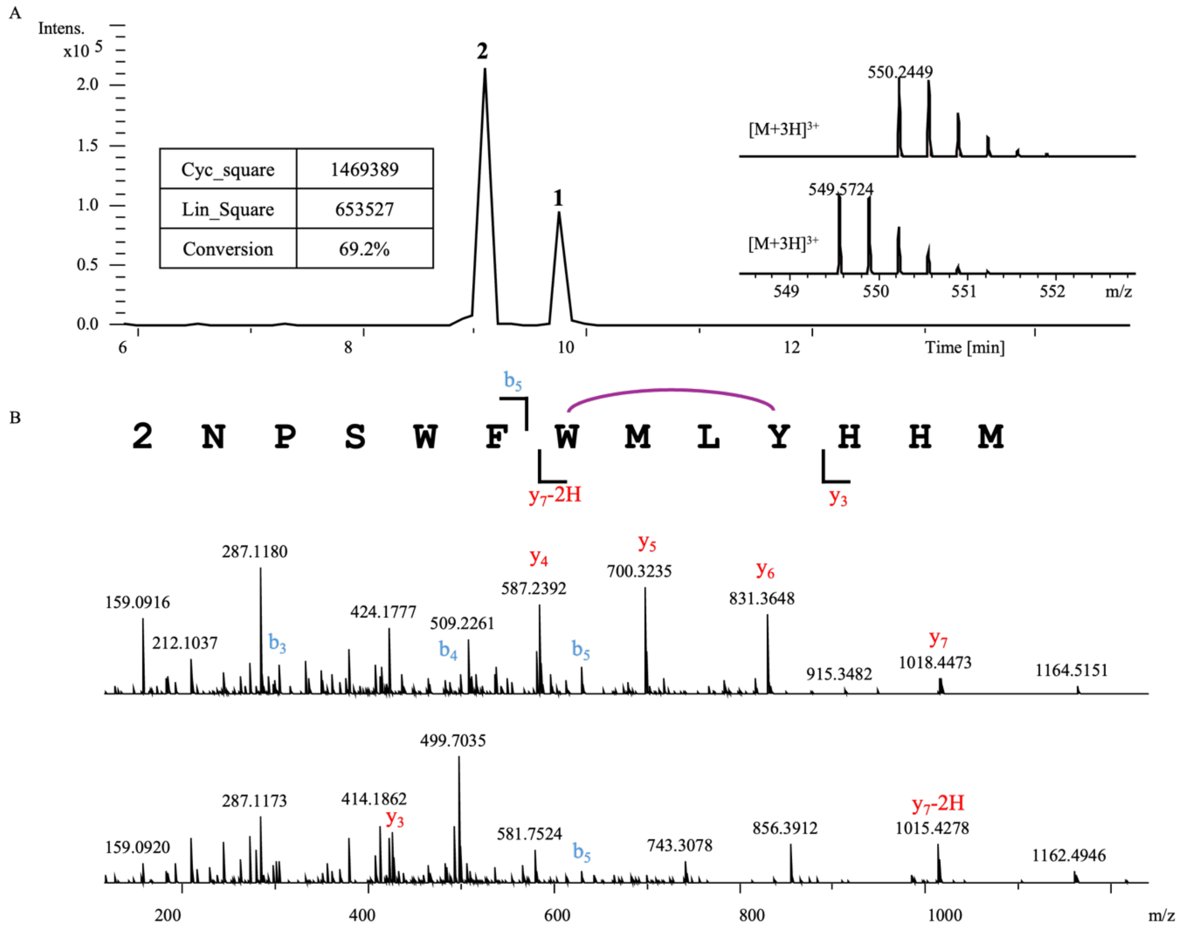


| | | **1** | | | |
|---|---|---|---|---|---|
| **Ions** | **Fragment** | **Observed mass** | **Calculated mass** | **z** | **Δppm** |
| $y_4$ | YHHM | 587.2392 | 587.2395 | 1 | 0.51 |
| $y_5$ | LYHHM | 700.3235 | 700.3235 | 1 | <0.1 |
| $y_6$ | MLYHHM | 831.3648 | 831.364 | 1 | 0.96 |
| $y_7$ | WMLYHHM | 1017.4405 | 1017.443 | 1 | 2.46 |
| $b_2$ | NP | 212.1037 | 212.103 | 1 | 3.3 |
| $b_3$ | NPS | 299.1355 | 299.135 | 1 | 1.67 |
| $b_4$ | NPSW | 485.2169 | 485.2143 | 1 | 5.36 |
| $b_5$ | NPSWF | 632.2831 | 632.2827 | 1 | 0.63 |

| | | **2** | | | |
|---|---|---|---|---|---|
| Ions | Fragment | Observed mass | Calculated mass | z | Δppm |
| $y_3$ | HHM | 424.1757 | 424.1761 | 1 | 0.94 |
| $y_7$-2H | WMLYHHM | 1015.4278 | 1015.428 | 1 | 0.2 |
| $b_5$ | NPSWF | 632.2825 | 632.2827 | 1 | 0.32 |

**Figure S6.** *In vivo* characterization of the *Gpe* family**.** His-SUMO-TEV GpeA₂ and GpeC were heterologously expressed in *Escherichia coli* Rossetta on a pRSF-DUET vector. After 20 hours of fermentation, the bacteria were lysed and purified with a Ni-affinity column. After digesting the purified peptide with trypsin, the sample was collected for UPLC-HRMS analysis. HRMS revealed a -2Da peak, and tandem mass confirmed the modification within the WMLY motif.

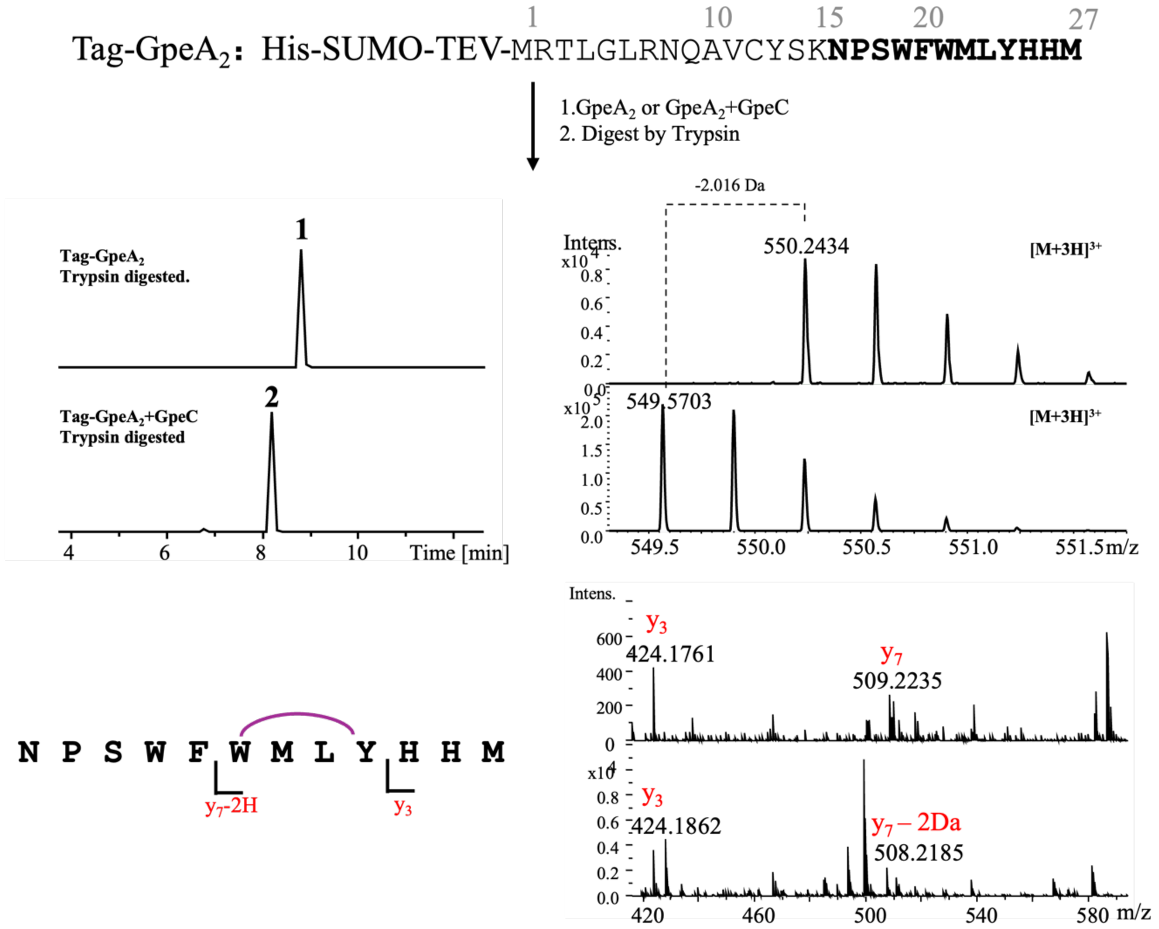

**Figure S7.** Comparative analysis of the consistency of the *in vivo* and *in vitro* catalyzed product. As the figure shows same retention time on the UPLC-HRMS system and the tandem mass profile of the two catalyzed products were totally the same. We proved the consistency of the *in vivo* and *in vitro* catalyzed product.

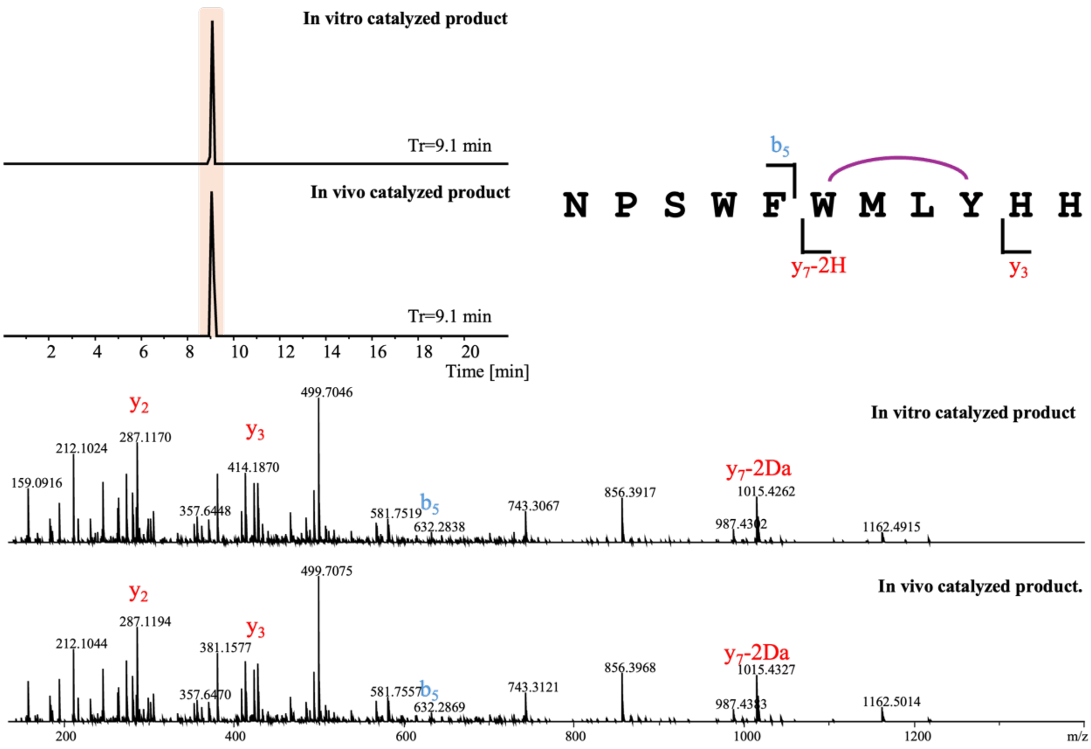

**Figure S8.** Structure elucidation of Gpertide (**3**)

#### Marfey analysis

| Residue | Derivative with L-FDLA | Derivative with D-FDLA | Configuration |
|---|---|---|---|
| Met | 16.6 | 18.2 | L |
| Leu | 16.5 | 18.7 | L |
| His | 12.4 | 11.9 | L |

#### HMBC and H-H Cosy correlations:

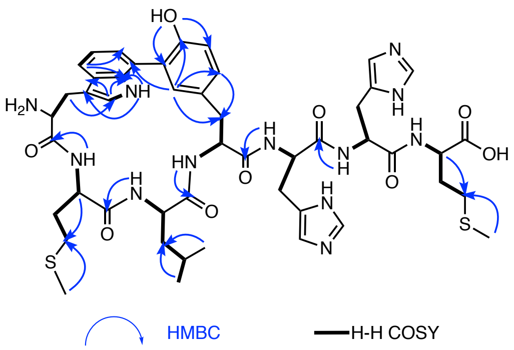


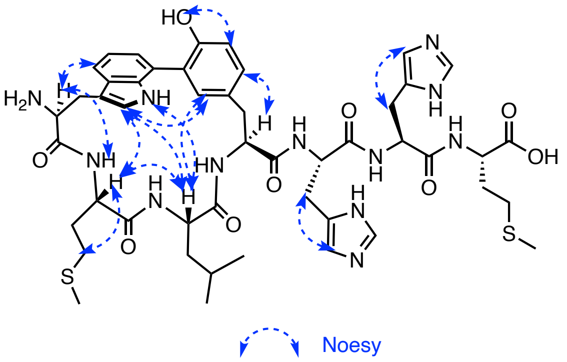


#### Key correlations

HMBC

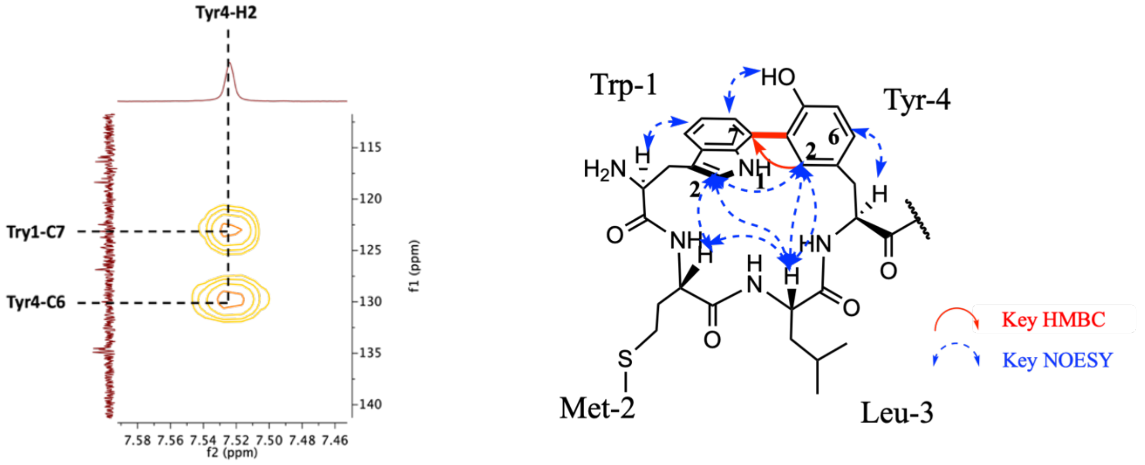


NOESY

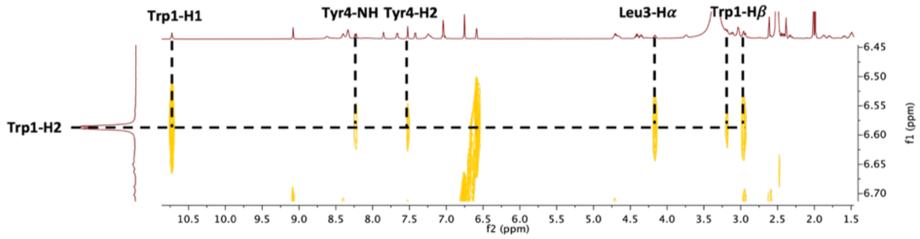

**Structure elucidation of Gpertide**

Temdem mass analysis revealed a 2Da loss on the WMLY module. Marfey's analysis proved that the Met, His, and Leu are all L-configured. We further subjected it to NMR analysis to elucidate the structure of Gpertide. The NH protons ranging from 7.42 to 8.62 in $^1$H NMR are consistent with their peptide nature (WMLYHHM). The signal ($\delta_H$=7.52, s) in $^1$HNMR presented an unusual singlet signal on the aromatic ring, further analysis proved it to be the Tyr4-H2. 2D NMR revealed its correlation with Trp1-C7 in HMBC spectra, which proved that a C-C bond was formed between Trp1-C7 to Tyr4-C3. The WMLY biaryl motif was further proved by the NOESY spectra of Trp1-H6 to Tyr4-OH and Trp1-H2 to Tyr4-H2, showing the close spatial orientation.

**Figure S9.** Electronic Circular Dichroism (ECD) analysis of Gpertide (**3**). (A) Experimental ECD spectra of **3**, calculated ECD of 3, and enantiomer. (B) Molecular model of the representative Gpertide with *Ra* configuration. (C) Detailed calculation information.

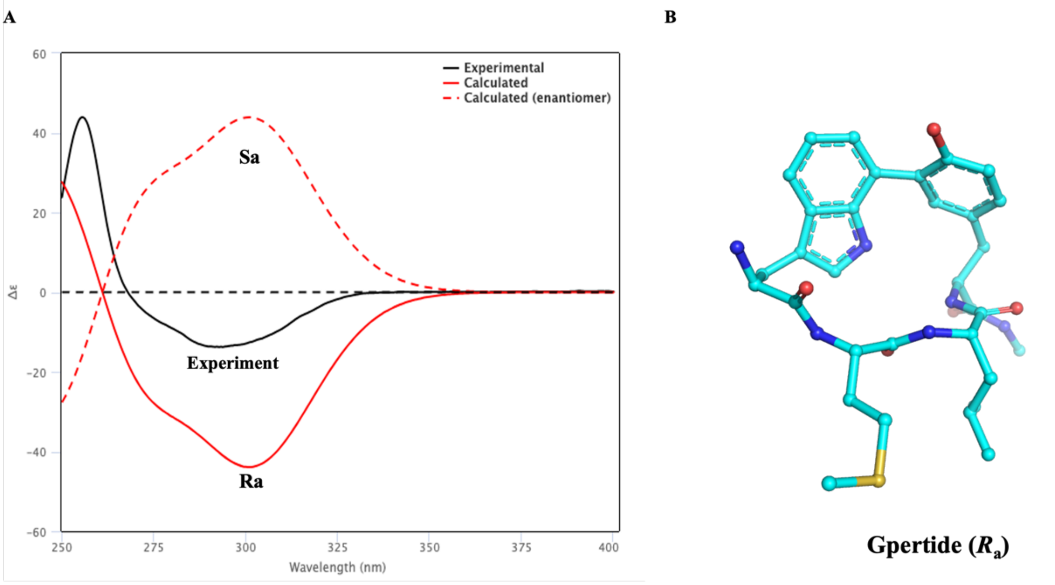


C

| Wave Range | Sigma | Shift |
|---|---|---|
| 250 – 400 nm | 0.26 ev | -18 nm |

| | Energy Hartree | Energy (kcal/mol) | Population (%) |
|---|---|---|---|
| 1 $R_a$ | -2348.15836960 | -1473491.61 | 100 |
| 1’ $S_a$ | - | - | 0 |

**Figure S10.** Analysis of the GpeC structural characteristics.

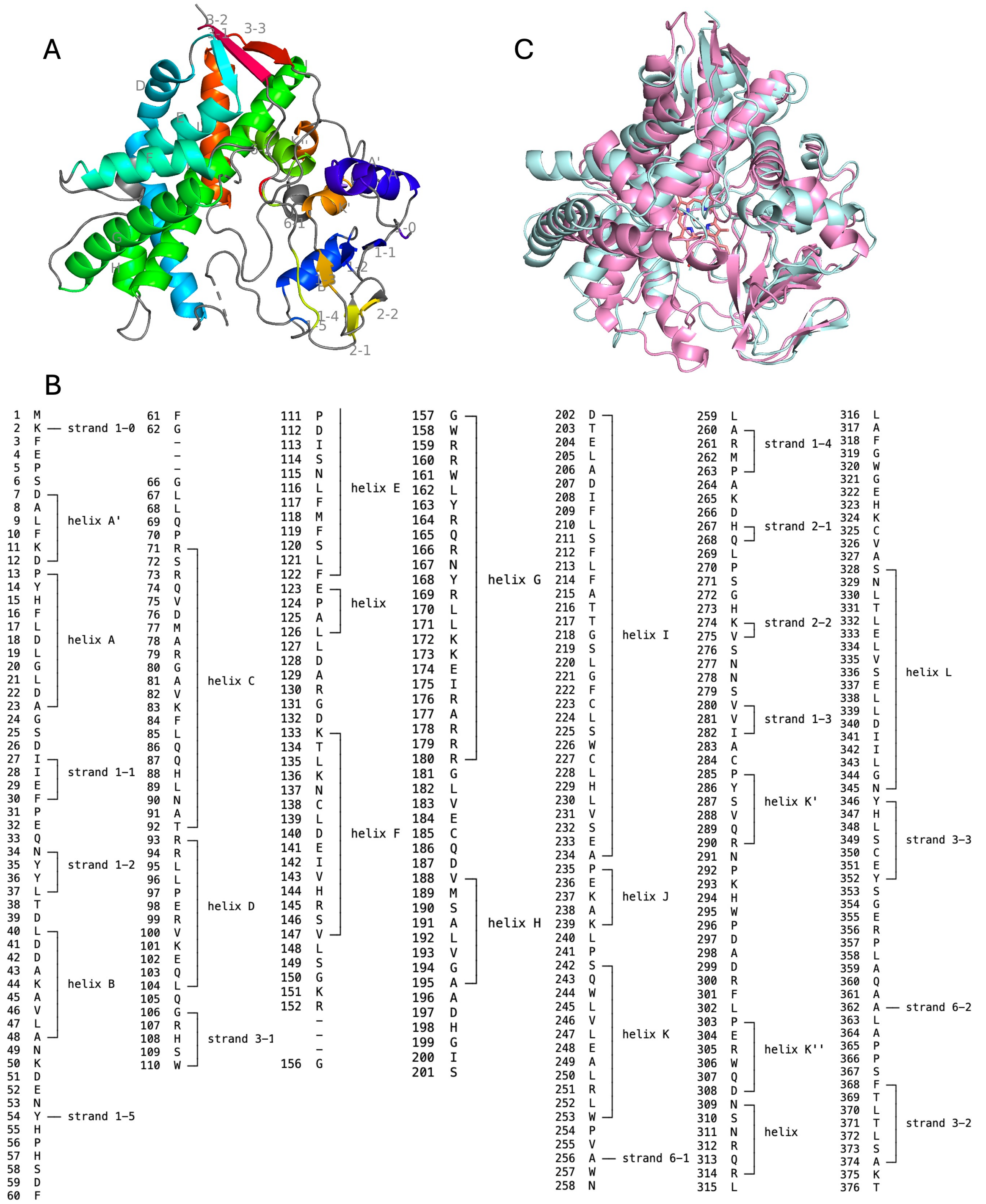


A) GpeC with typical structure of a cytochrome P450, showing the preponderance of a-helices in the structure (labelled A to L) together with one β-sheet (1-1 to 3-2). B) Amino acid sequences corresponding to different domains. The secondary structure of GpeC was predicted through the SecStrAnnotator website, using 2nnjA as a template. C) Structural alignment between GpeC and $P450_{Blt}$. Blue represents GpeC. Magenta represents $P450_{Blt}$. RMSD = 5.435. Z-scores = 25.6. In crystal structures, Z-score is commonly used to evaluate the quality of the structure[7].

**Figure S11.** Analysis of the substrate cavity of GpeC and $P450_{Blt}$.

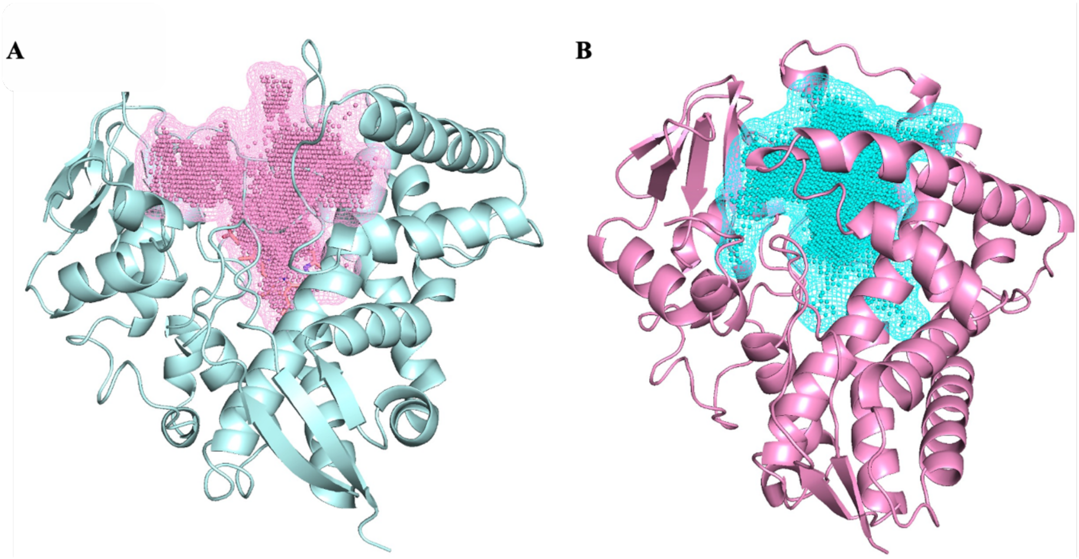


A) Analysis of substrate cavity of GpeC, with a size of 4049 Å³. B) Analysis of substrate cavity of P450Blt, with a size of 2574 Å³. Cavities were calculated using CavitOmiX. For the analysis of the hydrophobicity of the cavities, the corresponding hydrophobicity module of the program VASCo was used. Cavities were calculated using a modified LIGSITE algorithm.

**Figure S12.** Analysis of GpeC binding affinity and optimal conformation with $GpeA_2$.

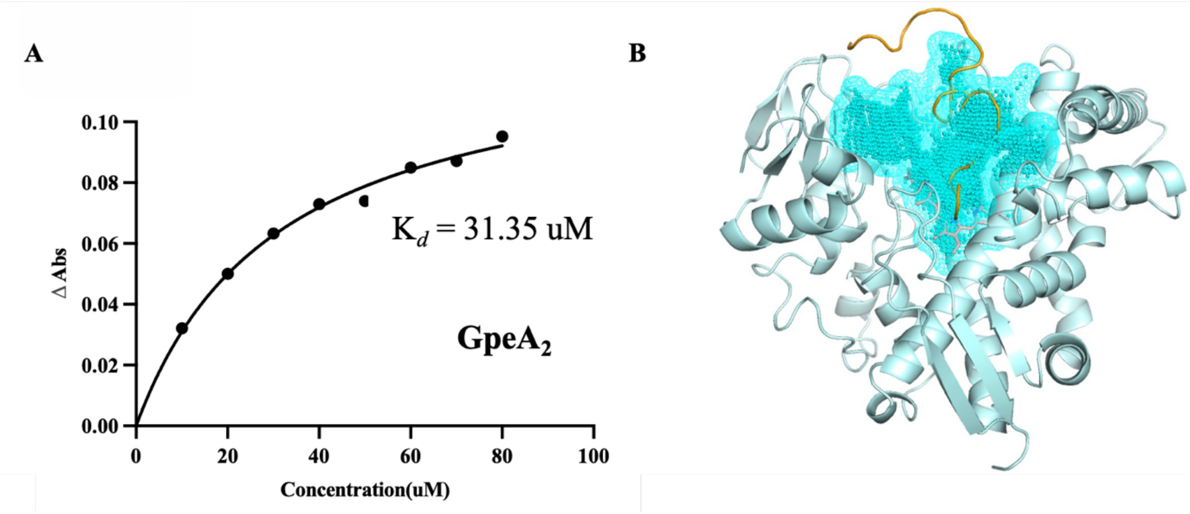


A) $K_d$ value of the binding of GpeC to the $GpeA_2$. B) The 27-mer nature substrate ($GpeA_2$) is positioned in the substrate cavity, with its N-terminal region extending outside the cavity.

**Figure S13.** Systematically truncated the N-terminal leader region of the native precursor peptide GpeA₂. Using stepwise three-residue deletions, we generated a series of constructs with up to 18 residues removed and assayed their activity with purified GpeC *in vitro*.

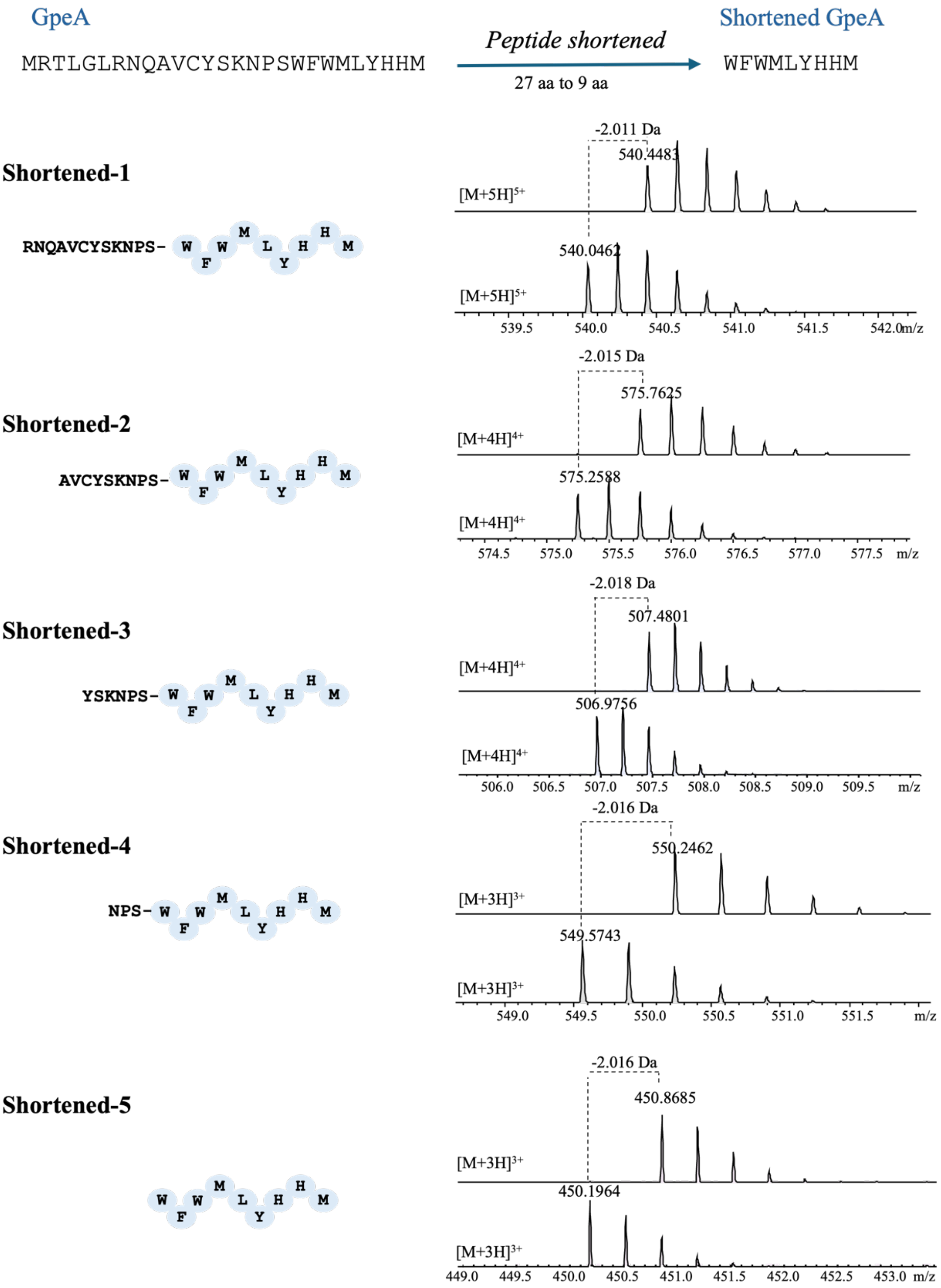

**Figure S14.** Analysis of the interaction between GpeC and substrate SGpeA$_2$.

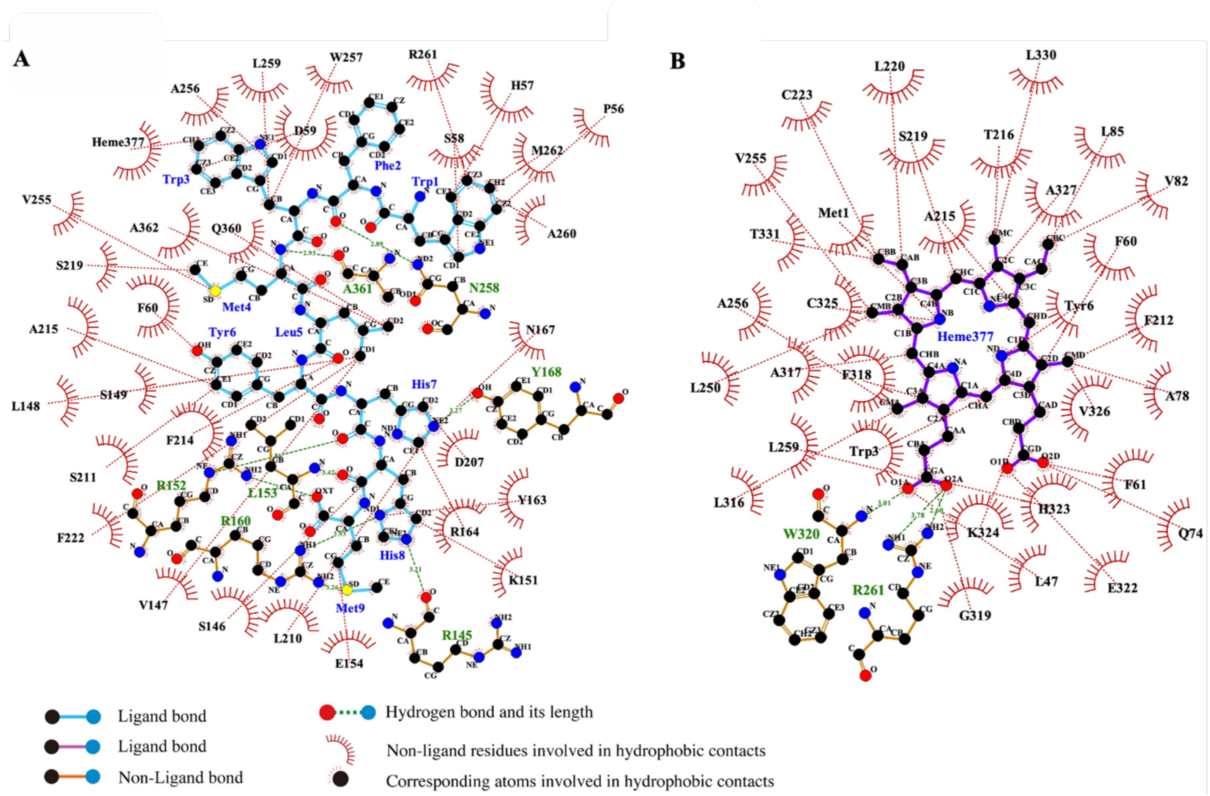


A) Analysis of the interaction among SGpeA$_2$ and surrounding amino acids, heme in GpeC. B) Analysis of the interaction among heme and surrounding amino acids, substrates in GpeC.

**Figure S15**. Variation diagram of ligand RMSF during MD simulation process.

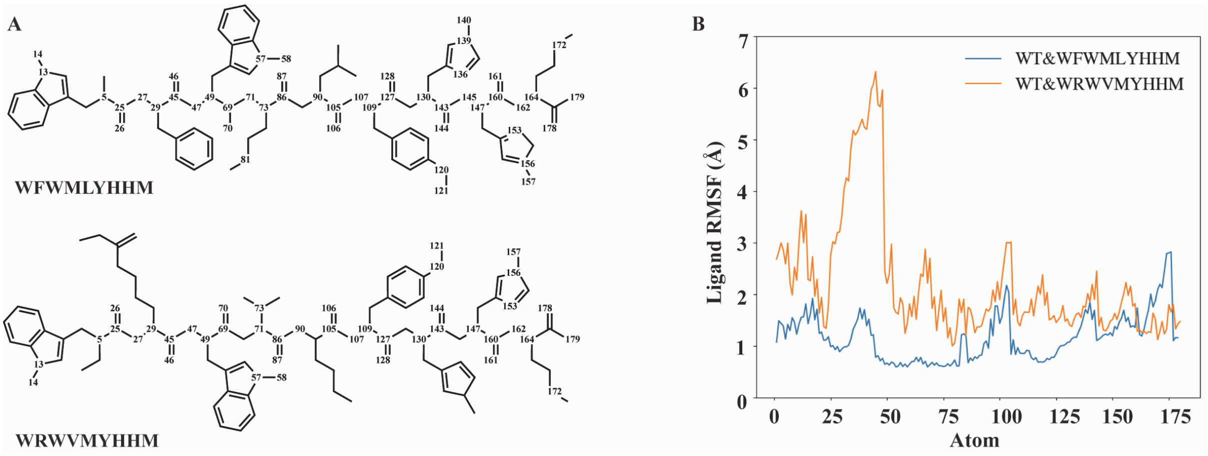


A) Atomic numbers of 9-mer substrates(SGpeA$_1$ and SGpeA$_2$). B) The ligand RMSF relative to GpeC displays peaks corresponding to the atoms of SGpeA$_1$/SGpeA$_2$ ligand residues, suggesting they are somewhat flexible, which is likely due to them being solvent-exposed.

**Figure S16.** Interaction network between GpeC and SGpeA$_2$.

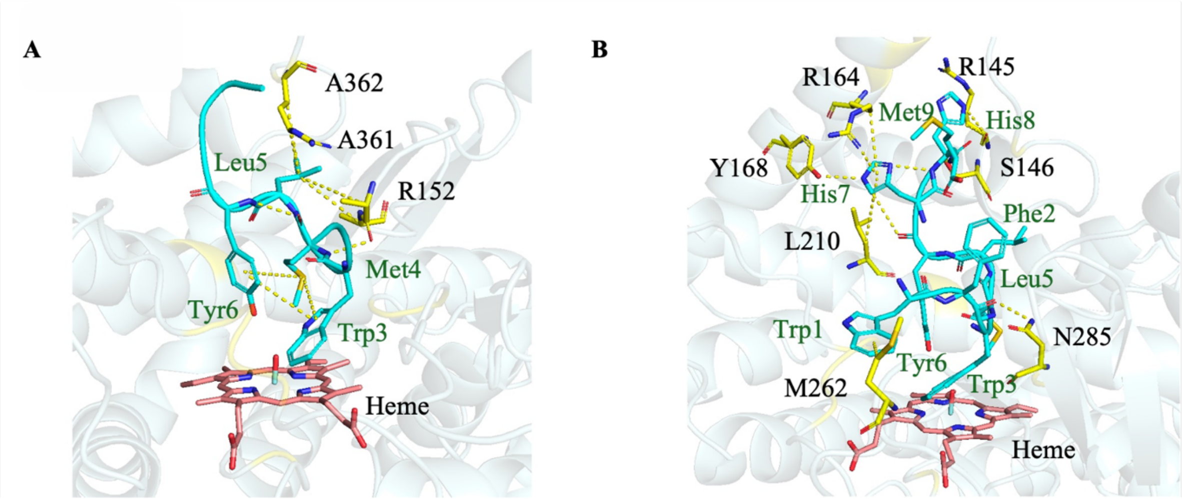


A) Interaction network of the WMLY motif of SGpeA$_2$ in the GpeC cavity. B) Alternative interaction network of WF and HHM motif of SGpeA$_2$ in the GpeC cavity.

**Figure S17.** Site-directed mutagenesis of GpeC.

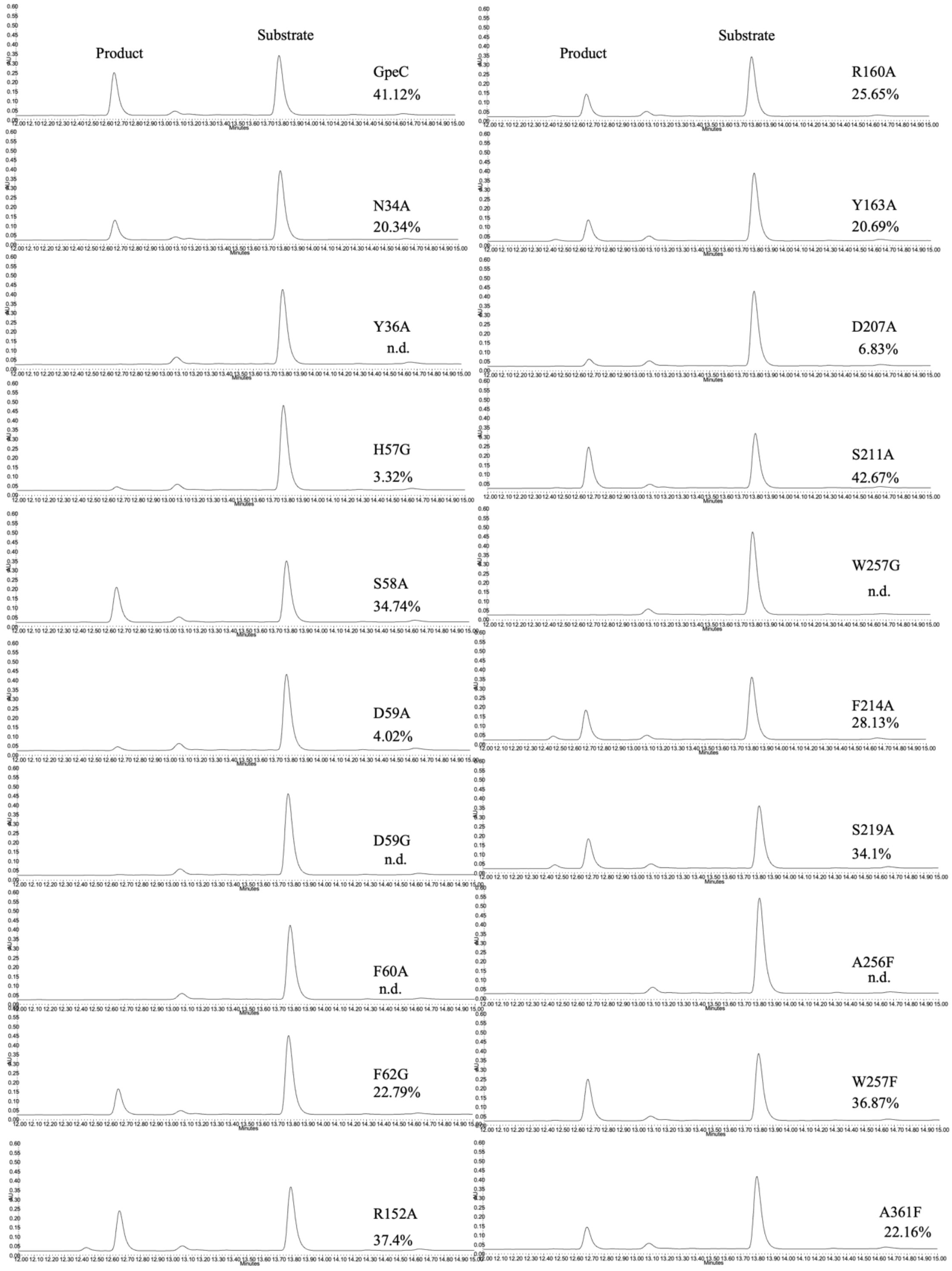


A) Site-directed mutagenesis of GpeC. Mutants were assayed with SGpeA$_2$ under 100uM substrate, 3 uM GpeC. Conversion was calculated by the abovementioned method.

**Figure S18.** *In vitro* assay of $SGpeA_2$ (300 µM) with GpeC (3 µM). A 5-hour reaction was performed before quenching by adding an equivalent volume of methanol. Conversion of the $SGpeA_2$ was calculated by the abovementioned method. Tandem mass analysis revealed the peptide ions of $y_2$, $y_3$, $y_7$-2H, $b_2$, and $b_7$-2H, proving the modification happened within the WMLY motif. TTN was calculated by conversion.

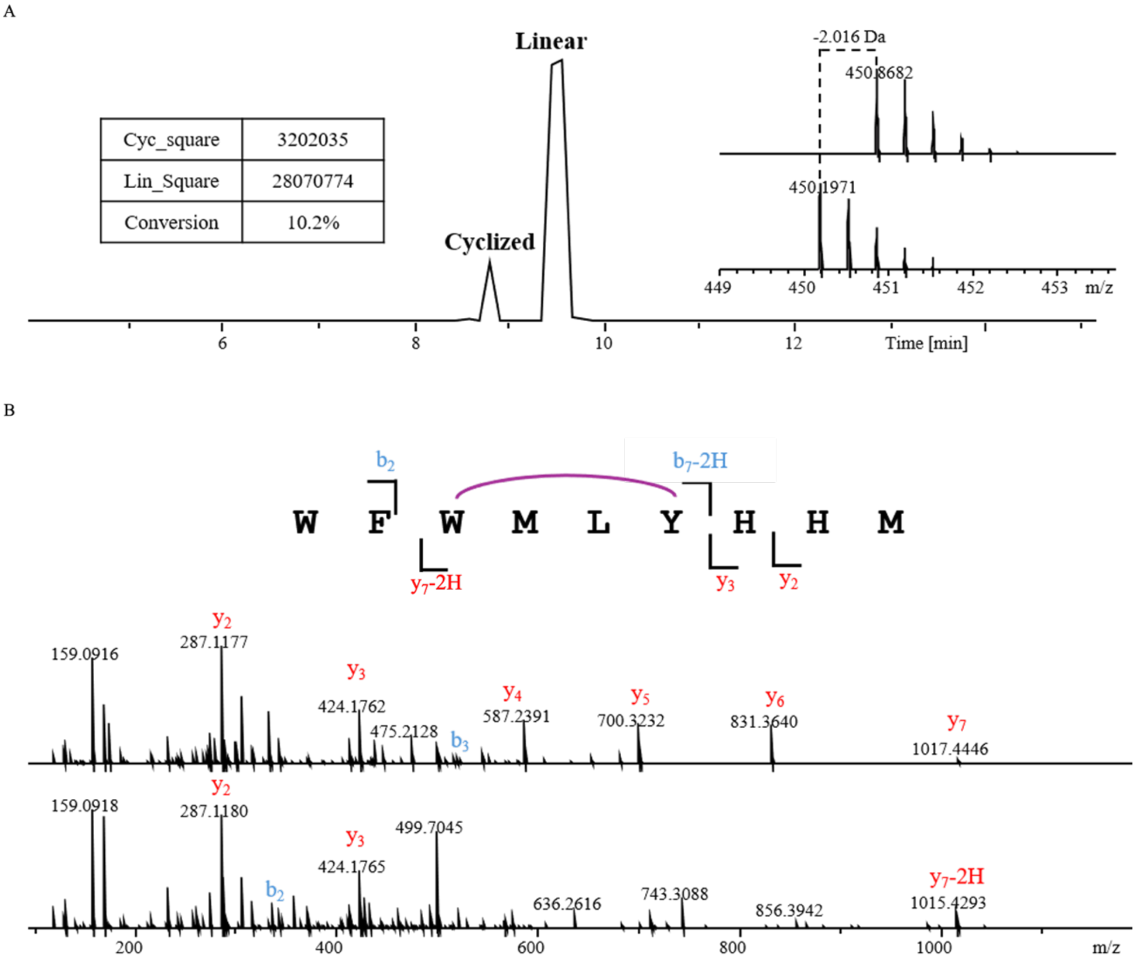


**Linear WFWMLYHHM**

| Ions | Fragment | Observed mass | Calculated mass | z | Δppm |
|---|---|---|---|---|---|
| $y_4$ | YHHM | 587.2391 | 587.2395 | 1 | 0.68 |
| $y_5$ | LYHHM | 700.3232 | 700.3235 | 1 | 0.43 |
| $y_6$ | MLYHHM | 831.3640 | 831.364 | 1 | <0.1 |
| $y_7$ | WMLYHHM | 1017.4446 | 1017.443 | 1 | 1.57 |
| $b_2$ | WF | 334.1552 | 334.155 | 1 | 0.6 |
| $b_3$ | WFW | 520.2339 | 520.2343 | 1 | 0.77 |
| $b_4$ | WFWM | 651.2758 | 651.2768 | 1 | 1.54 |

**Cyclized WFWMLYHHM**

| Ions | Fragment | Observed mass | Calculated mass | z | Δppm |
|---|---|---|---|---|---|
| $y_3$ | HHM | 424.1765 | 424.1761 | 1 | 0.94 |
| $y_7$-2H | WMLYHHM | 1015.4293 | 1015.428 | 1 | 1.28 |
| $b_2$ | WF | 334.1551 | 334.155 | 1 | 0.3 |

**Figure S19.** *In vitro* assay of SGpeA$_1$ (300 μM) with GpeC (3 μM). A 5-hour reaction was performed before quenching by adding an equivalent volume of methanol. Conversion of the SGpeA$_1$ was calculated by the square of the linear peptide and the cyclized one. Tandem mass analysis revealed the peptide ions of $y_2$, $y_3$, $b_2$, and $b_7$-2H, proving the modification happened within the WVMY motif. TTN was calculated by conversion.

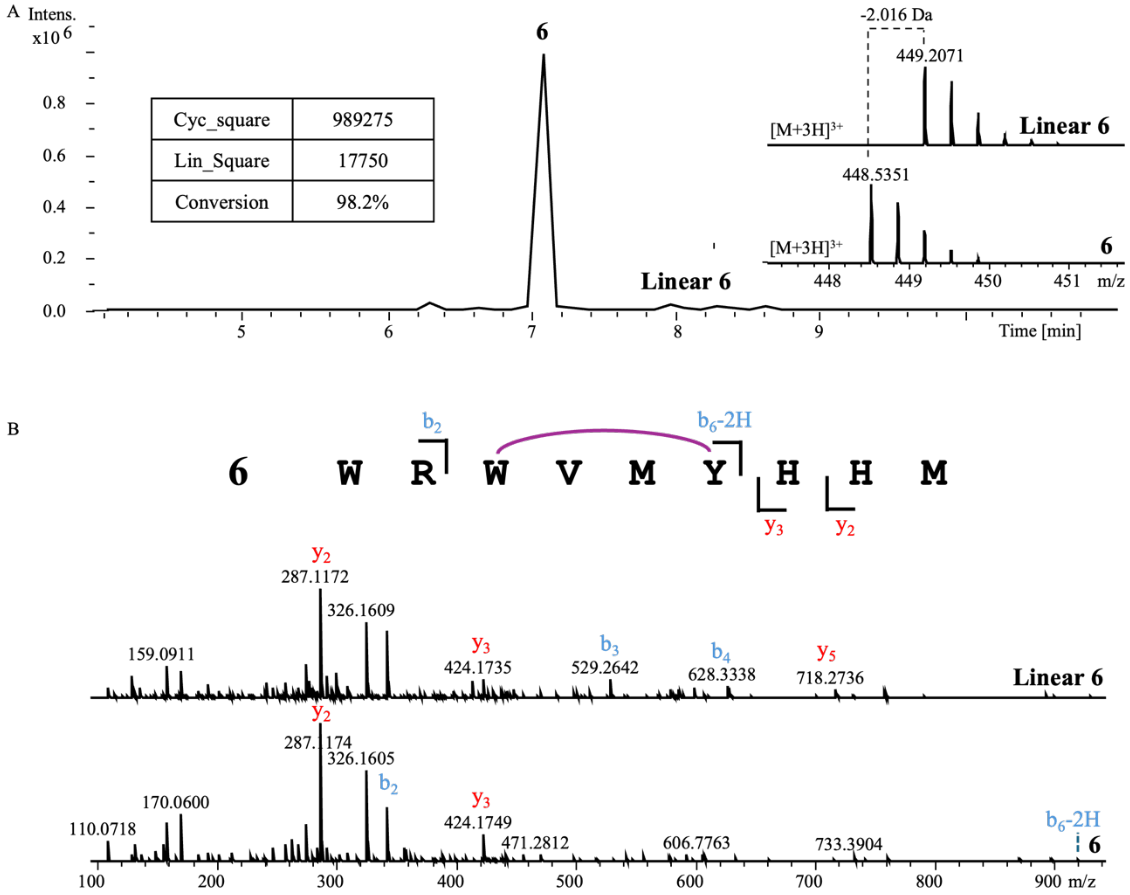


**Linear 6**

| Ions | Fragment | Observed mass | Calculated mass | z | Δppm |
|---|---|---|---|---|---|
| $y_2$ | HM | 287.1172 | 287.1172 | 1 | <0.1 |
| $y_3$ | HHM | 424.1735 | 424.1761 | 1 | 6.13 |
| $y_4$ | YHHM | 587.2347 | 587.2395 | 1 | 8.17 |
| $y_5$ | MYHHM | 718.2736 | 718.28 | 1 | 8.91 |
| $b_2$ | WR | 343.1865 | 343.1877 | 1 | 3.5 |
| $b_3$ | WRW | 529.2642 | 529.267 | 1 | 5.29 |
| $b_4$ | WRWV | 628.3338 | 628.3354 | 1 | 2.55 |

**6**

| Ions | Fragment | Observed mass | Calculated mass | z | Δppm |
|---|---|---|---|---|---|
| $y_2$ | HM | 287.1174 | 287.1172 | 1 | 0.7 |
| $y_3$ | HHM | 424.1749 | 424.1761 | 1 | 2.83 |
| $b_6$-2H | WRWVMY | 920.4228 | 920.4236 | 1 | 0.87 |

**Figure S20.** Binding conformation of SGpeA$_1$ and GpeC.

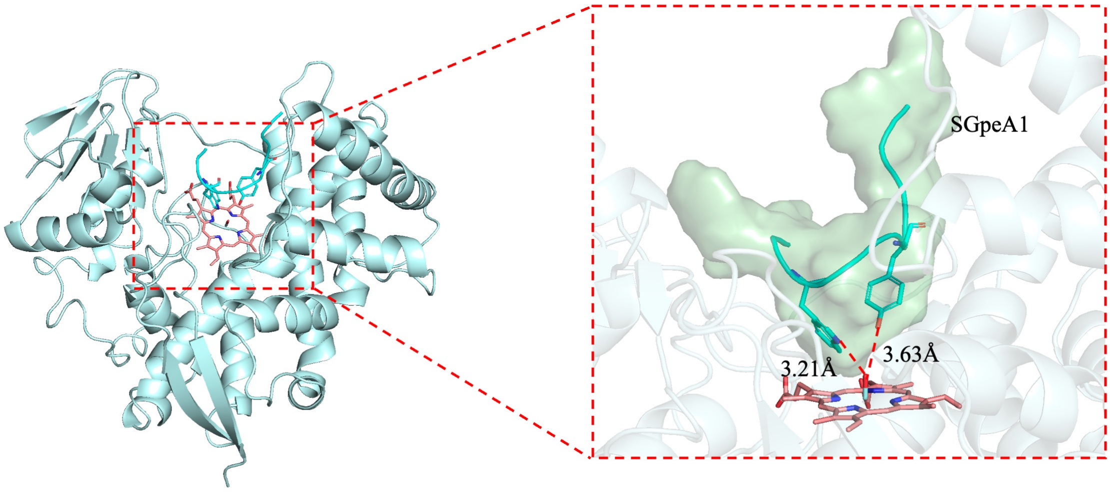


Relative positions of the heme plane and the SGpeA$_1$ (cyan sticks and cartoon) in the active pocket. Reaction residues Trp3 and Tyr6 are within the catalytic region of the heme center.

**Figure S21.** *In vitro* assay of WRWVMYHH with GpeC. A 5-hour reaction was performed before quenching by adding an equivalent volume of methanol. Only a trace amount of modified product was observed (less than 5%). Tandem mass analysis revealed the linear peptide ions of $y_2$, $y_3$, $y_4$, $y_5$, $y_7$, $b_2$, and $b_3$.

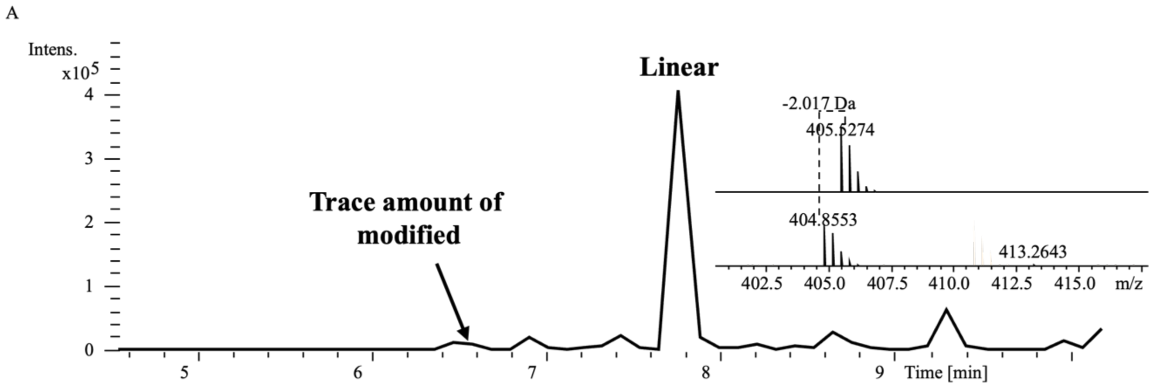


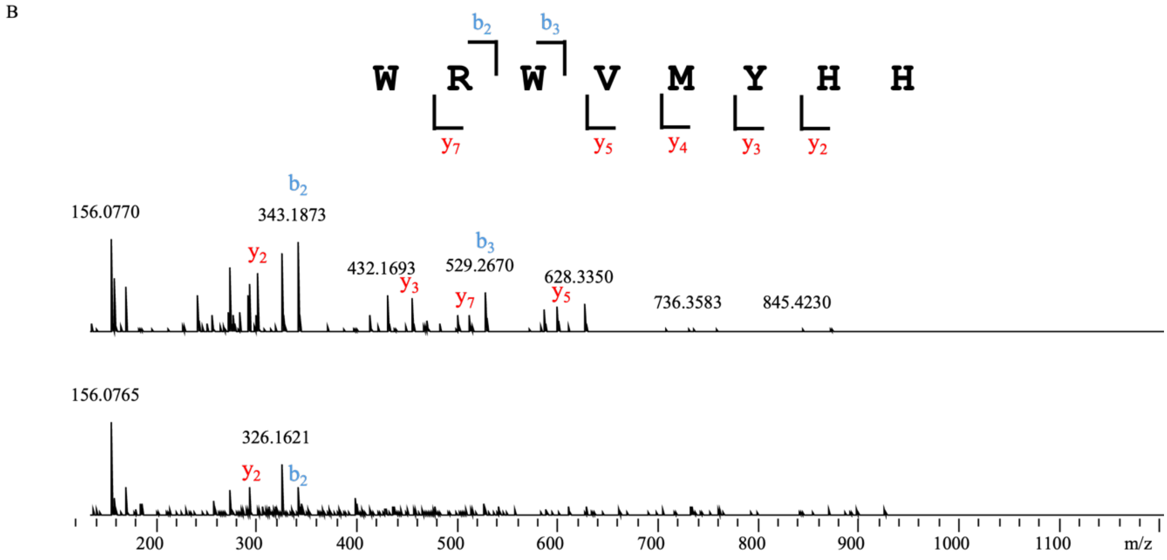


| WRWVMYHH | | | | | |
|---|---|---|---|---|---|
| Ions | Fragment | Observed mass | Calculated mass | z | Δppm |
| $y_2$ | HH | 293.1358 | 293.1357 | 1 | 0.34 |
| $y_3$ | YHH | 456.1991 | 456.199 | 1 | 0.22 |
| $y_4$ | MYHH | 587.2385 | 587.2395 | 1 | 1.7 |
| $y_5$ | VMYHH | 686.3071 | 686.3079 | 1 | 1.17 |
| $y_7$ | RWVMYHH | 514.7463 | 514.7478 | 2 | 2.91 |
| $b_2$ | WR | 343.1873 | 343.1877 | 1 | 1.17 |
| $b_3$ | WRW | 529.267 | 529.267 | 1 | <0.1 |

| WRWVMYHH_modified | | | | | |
|---|---|---|---|---|---|
| Ions | Fragment | Observed mass | Calculated mass | z | Δppm |
| $y_2$ | HH | 293.1367 | 293.1357 | 1 | 3.41 |
| $b_2$ | WR | 343.1846 | 343.1877 | 1 | 9.03 |

**Figure S22.** *In vitro* assay of 8-mer substrate RWVMYHHM with GpeC. A 5-hour reaction was performed before quenching by adding an equivalent volume of methanol. Only linear substrate observed. Tandem mass analysis revealed the linear peptide ions of $y_2$, $y_3$, $y_4$, $y_5$, $b_2$, and $b_3$.

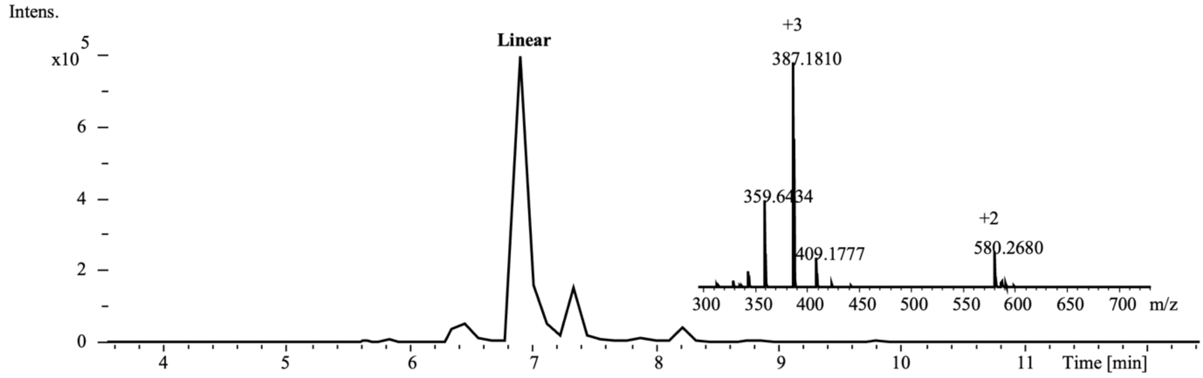


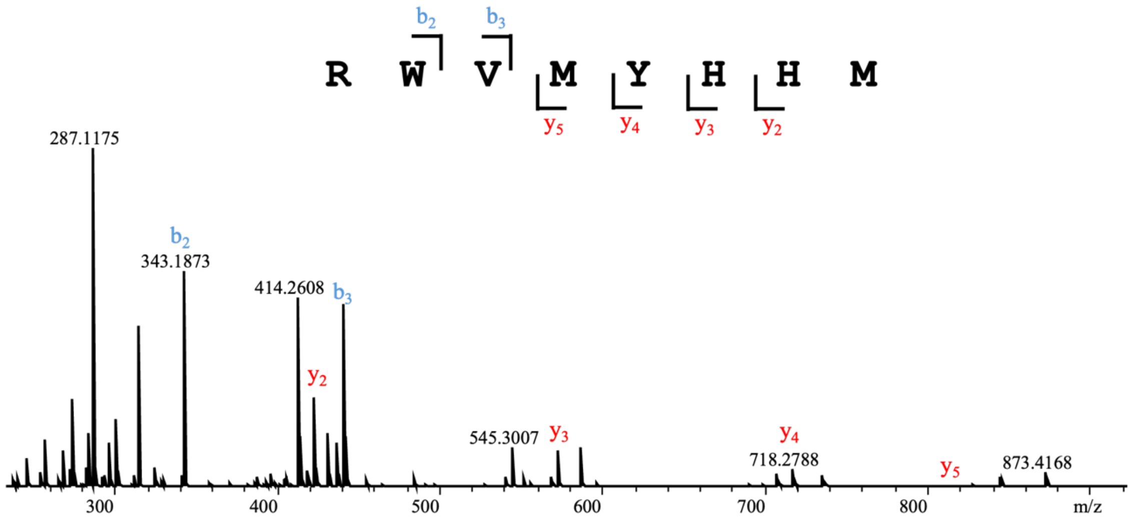


| RWVMYHHM | | | | | |
|---|---|---|---|---|---|
| Ions | Fragment | Observed mass | Calculated mass | z | Δppm |
| $y_2$ | HHM | 424.1759 | 424.1761 | 1 | 0.47 |
| $y_3$ | YHHM | 587.2390 | 587.2395 | 1 | 0.85 |
| $y_4$ | MYHHM | 718.2788 | 718.28 | 1 | 1.67 |
| $y_5$ | VMYHHM | 817.3481 | 817.3484 | 1 | 0.37 |
| $b_2$ | RW | 343.1873 | 343.1877 | 1 | 1.17 |
| $b_3$ | RWV | 442.2561 | 442.2561 | 1 | <0.1 |

**Figure S23.** *In vitro* assay of SGpeA$_1$ with GpeC by peroxide shunt pathway. A 5-hour reaction was performed with 0.5 µL 30% $H_2O_2$, 3 µM GpeC, and 0.3 mM substrate within 50 µL reaction buffer before quenching by adding an equivalent volume of methanol. Cyclized product (**6'**) was found, with a 30 Da mass gained, indicating the two oxidized Met and a C-C bond forming within the WVMY motif. TTN was calculated by conversion.

A

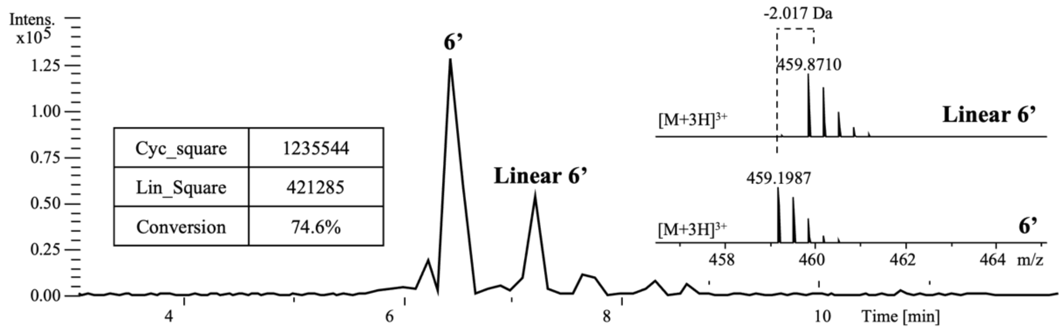


B

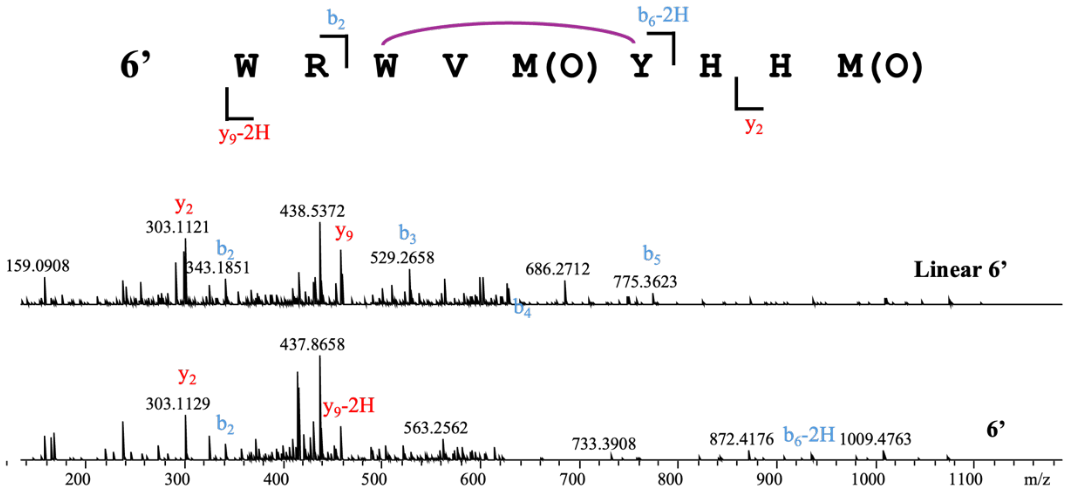


| Linear 6' | | | | | |
|---|---|---|---|---|---|
| **Ions** | **Fragment** | **Observed mass** | **Calculated mass** | **z** | **Δppm** |
| $y_2$ | HM(Met-Oxidized) | 303.1121 | 303.1162 | 1 | 13.5 |
| $y_9$ | WRWVMYHHM (Met-Oxidized) | 459.8715 | 459.8736 | 3 | 4.57 |
| $b_2$ | WR | 343.1851 | 343.1877 | 1 | 7.58 |
| $b_3$ | WRW | 529.2658 | 529.267 | 1 | 2.27 |
| $b_4$ | WRWV | 628.3357 | 628.3354 | 1 | 0.49 |
| $b_5$ | WRWVM(Met-Oxidized) | 775.3623 | 775.3749 | 1 | 16.3 |

| 6' | | | | | |
|---|---|---|---|---|---|
| **Ions** | **Fragment** | **Observed mass** | **Calculated mass** | **z** | **Δppm** |
| $y_2$ | HM(Met-Oxidized) | 303.1129 | 303.1162 | 1 | 10.9 |
| $y_9$-2H | WRWVMYHHM (Met-Oxidized) | 459.1983 | 459.2016 | 3 | 7.19 |
| $b_2$ | WR | 343.1860 | 343.1877 | 1 | 4.95 |
| $b_6$-2H | WRWVMY (Oxidized Met) | 936.4175 | 936.4225 | 1 | 5.34 |

**Figure S24.** Catalytic consistency of GpeC under the ferredoxin/ferredoxin reductase/NADPH pathway and the peroxide shunt pathway. 50 µL reaction was first performed through the Fdr/Fdx/NaDPH pathway for 5 hours, as we precipitated the enzyme with methanol, 0.5 µL 30% peroxide was added for one more hour, and the catalytic product was sent to UPLC-HRMS analysis. As the catalytic product between the two systems showed a similar retention time on the LC system, and the tandem mass profile showed similar fragmentation, we proposed that the bond formation was consistent under the Fdr/Fdx/NADPH pathway and the peroxide shunt pathway. The enzymatic kinetics were measured, demonstrating the catalytic efficiency of the two pathways.

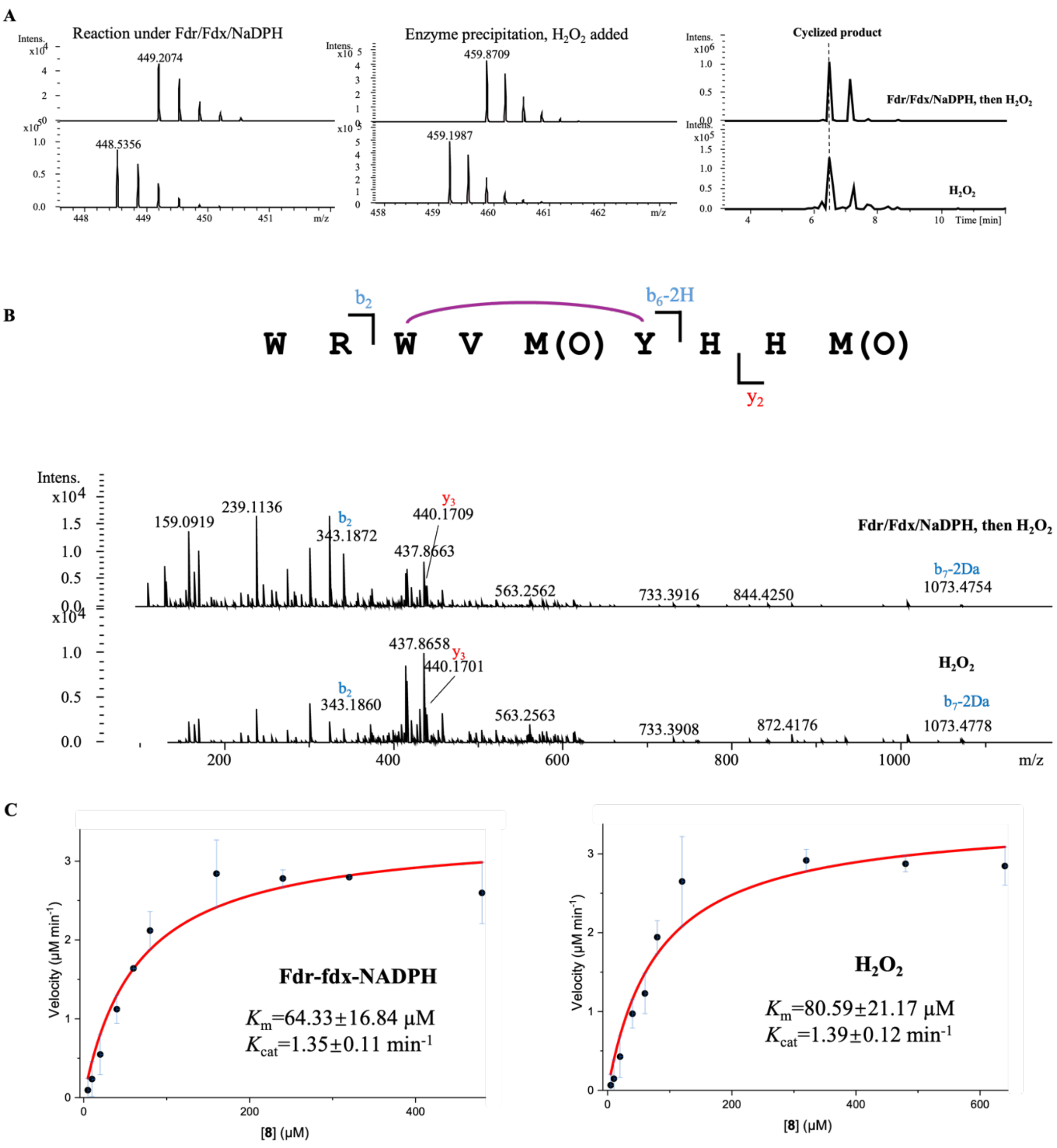

**Figure S25.** GpeC enzymatic assay with WRYVMYHHM. A 5-hour reaction was performed before quenching by adding an equivalent volume of methanol. Almost all of the linear substrate was catalyzed to the biaryl cyclized product (**7**). Tandem mass analysis revealed the compound (**7**) ions of $y_2$, $y_3$, $b_2$, and $b_7$-2H. To isolate the cyclized product for structure elucidation, we performed a large-scale reaction of 25 mL and sent it for HPLC isolation. The compound (**7**) was marked as the target peak in green.

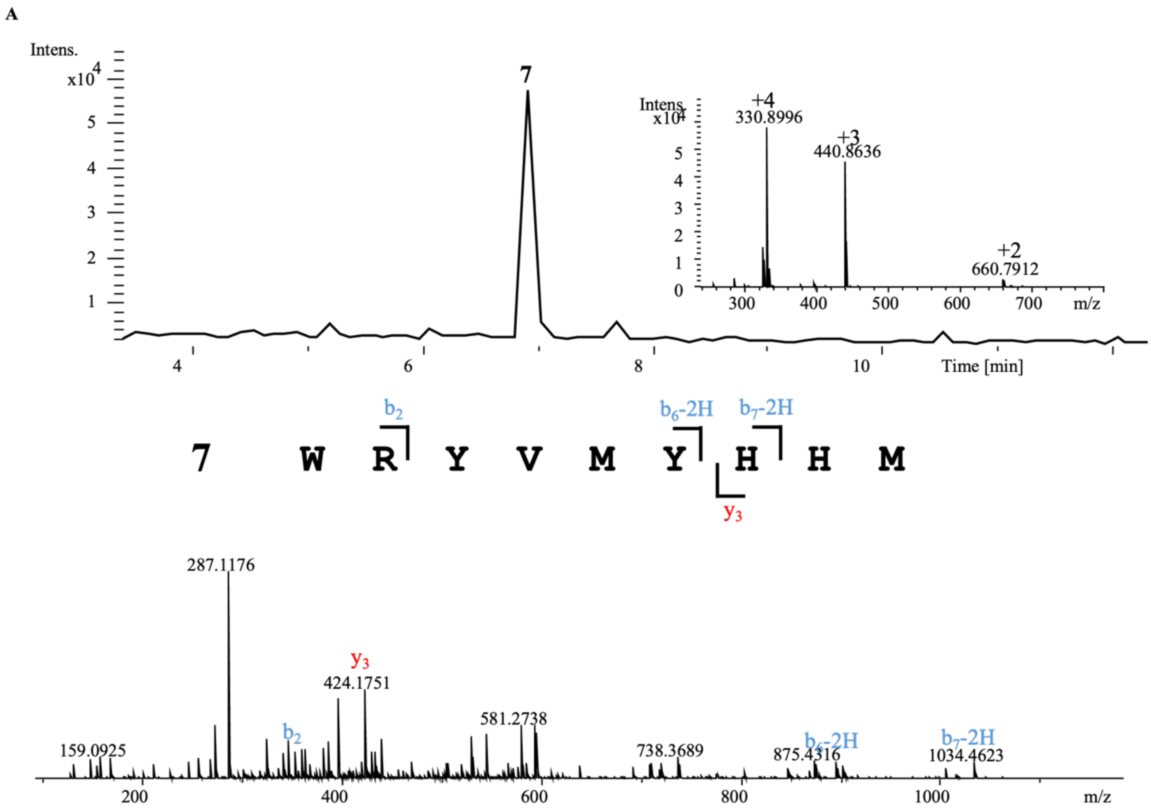


| | | **7** | | | |
|---|---|---|---|---|---|
| **Ions** | **Fragment** | **Observed mass** | **Calculated mass** | **z** | **Δppm** |
| $y_3$ | HHM | 424.1751 | 424.1761 | 1 | 2.36 |
| $b_2$ | WR | 343.1891 | 343.1877 | 1 | 4.08 |
| $b_6$-2H | WRYVMY | 897.4043 | 897.4076 | 1 | 3.68 |
| $b_7$-2H | WRYVMYH | 1034.4623 | 1034.4665 | 1 | 4.06 |

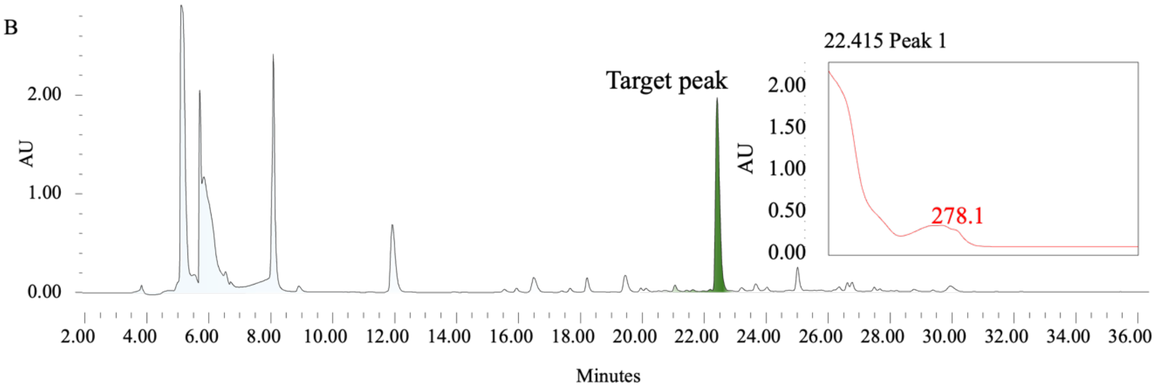

**Figure S26.** Structure elucidation of Typertide (**7**)

| Residue | Derivative with L-FDLA | Derivative with D-FDLA | Configuration |
|---|---|---|---|
| Met | 15.1 | 16.6 | L |
| Val | 15.2 | 17.0 | L |
| His | 11.9 | 11.4 | L |
| Arg | 12.4 | 12.1 | L |
| Trp | 15.9 | 16.8 | L |

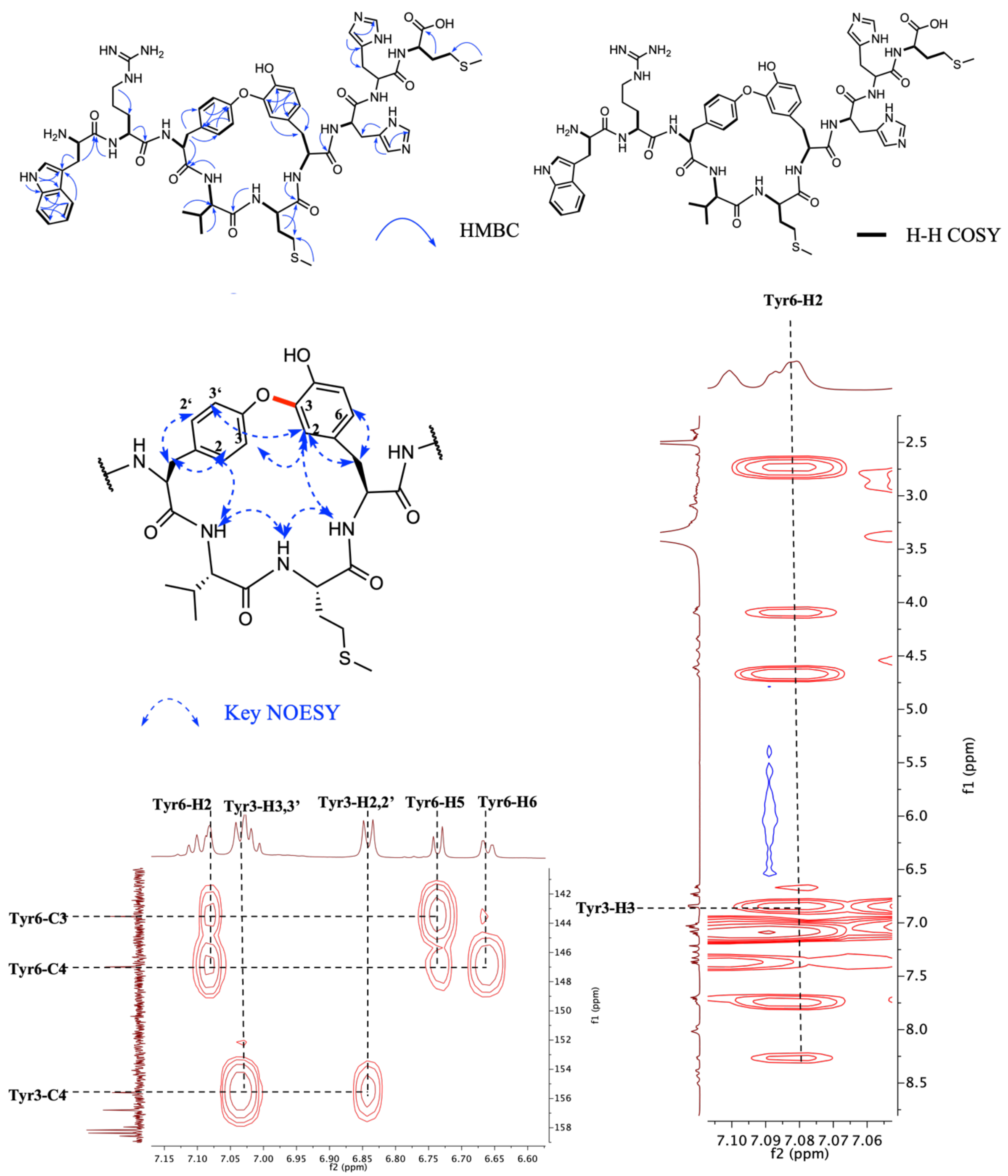

**Structure elucidation of typertide**

Tandem mass analysis revealed a 2 Da loss on the YVMY module. Marfey's analysis proved that the Met, His, Trp, Arg, and Val are all L-configured. We further subjected it to NMR analysis to elucidate the structure of Typertide. The NH protons ranging from 7.72 to 8.85 in $^{1}$H NMR are consistent with their peptide nature (WRYVMYHHM). The signal ($\delta_C$=143.5) in $^{13}$CNMR presented an unusual signal on the aromatic ring, we concluded for a single peak of Tyr6-C3, and we proposed that the carbon is connected with an electron-withdrawing group. Together, Tyr6-H2 showed an unusual singlet signal($\delta_H$=7.08, s). 2D NMR revealed its correlation with Tyr3-H3 in NOESY spectra, showing the close spatial orientation between Tyr3 and Tyr6. With all of the evidence together, we propose that a C-O bond was formed between Tyr3-O to Tyr6-C3.

**Figure S27.** GpeC enzymatic assay with WRYVMWHHM. A 5-hour reaction was performed before quenching by adding an equivalent volume of methanol. Tandem mass analysis revealed the compound (**8**) ions of $y_3$, $b_2$, $b_6$-2H, and $b_7$-2H. To isolate the cyclized product for structure elucidation, we performed a large-scale reaction of 25 mL and sent it for HPLC isolation. The compound (**8**) was marked as the target peak in green.

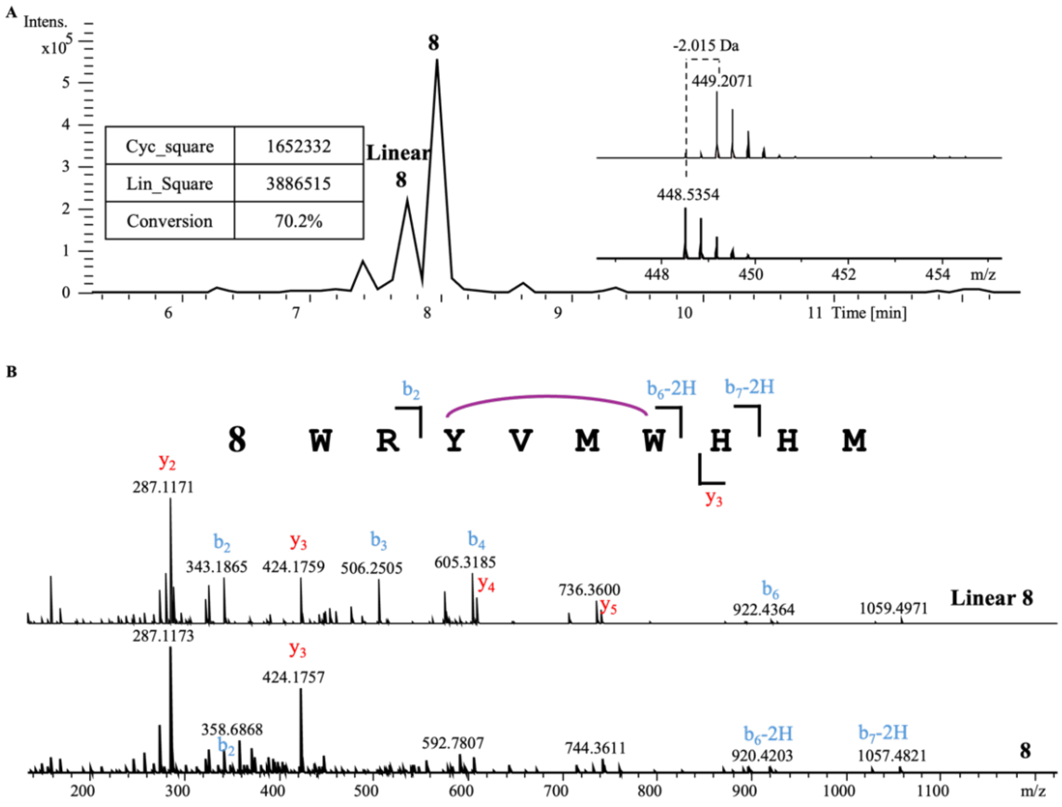


**Linear 8**

| Ions | Fragment | Observed mass | Calculated mass | z | Δppm |
|---|---|---|---|---|---|
| $y_2$ | HM | 287.1171 | 287.1172 | 1 | 0.35 |
| $y_3$ | HHM | 424.1759 | 424.1761 | 1 | 0.47 |
| $y_4$ | WHHM | 610.2544 | 610.2554 | 1 | 1.64 |
| $y_5$ | MWHHM | 741.2954 | 741.2959 | 1 | 0.67 |
| $b_2$ | WR | 343.1865 | 343.1877 | 1 | 3.5 |
| $b_3$ | WRY | 506.2505 | 506.251 | 1 | 0.99 |
| $b_4$ | WRYV | 605.3185 | 605.3194 | 1 | 1.49 |
| $b_6$ | WRYVMW | 922.4364 | 922.4392 | 1 | 3.04 |

**8**

| Ions | Fragment | Observed mass | Calculated mass | z | Δppm |
|---|---|---|---|---|---|
| $y_3$ | HHM | 424.1757 | 424.1761 | 1 | 0.94 |
| $b_2$ | WR | 343.1872 | 343.1877 | 1 | 1.46 |
| $b_6$-2H | WRYVMW | 920.4203 | 920.4236 | 1 | 3.59 |
| $b_7$-2H | WRYVMWH | 1057.4821 | 1057.483 | 1 | 0.85 |

C

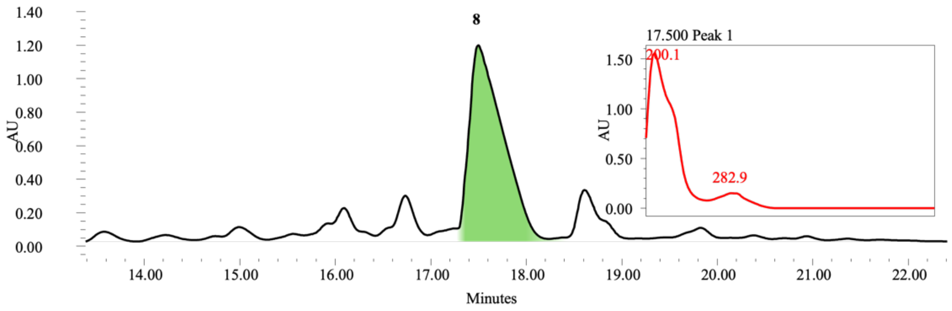


**Figure S28.** Structure elucidation of Gpentide (**8**)

| Residue | Derivative with L-FDLA | Derivative with D-FDLA | Configuration |
|---|---|---|---|
| Met | 15.1 | 16.6 | L |
| Val | 15.1 | 17.0 | L |
| His | 11.9 | 11.4 | L |
| Arg | 13.1 | 12.9 | L |
| Trp | 15.9 | 16.8 | L |

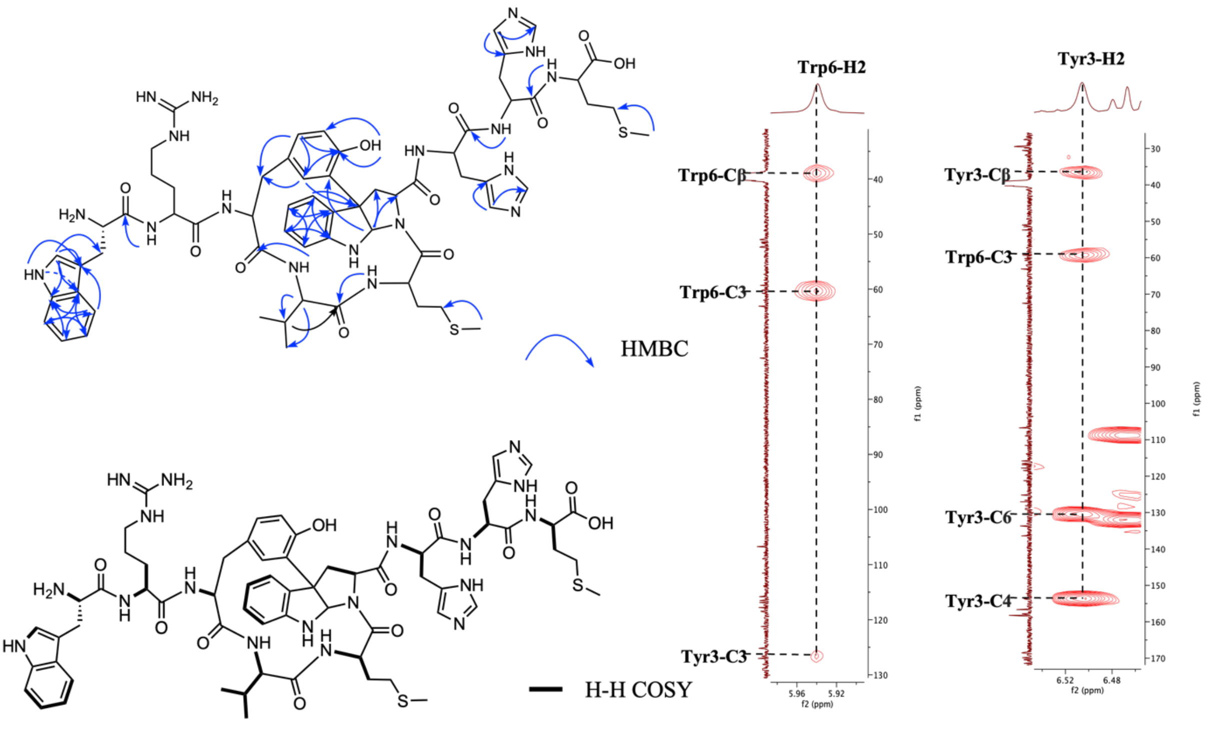

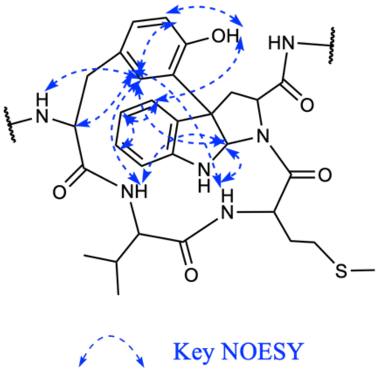


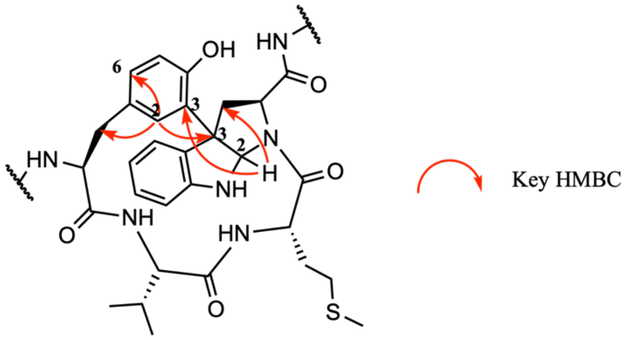


**Structure elucidation of Gpentide**

Tandem mass analysis revealed a 2 Da loss on the YVMW module. Marfey's analysis proved that the Met, His, Trp, Arg, and Val are all L-configured. We further subjected it to NMR analysis to elucidate the structure of Typertide. The NH protons ranging from 7.75 to 8.84 in $^{1}$H NMR are consistent with their peptide nature (WRYVMWHHM). The signal ($\delta_C$=81.5, $\delta_C$=59.3) in $^{13}$CNMR and ($\delta_H$=5.94,s) in $^{1}$HNMR, presented to be unusual signals. Comparative analysis of literature data identified these signatures as diagnostic of a C–N bond between the C-2 position of tryptophan and the corresponding backbone amide NH. This kind of structure has been found in several natural products containing tryptophan, eg, tryptorubin. So we proposed that the unusual signal in $^{13}$CNMR and $^{1}$HNMR was due to the C-N (Trp6-C2 to Trp6-N) and C-C (Tyr3-C3 to Trp6-C3) bonds forming. 2D NMR revealed its correlation with Tyr3-H2 to Trp6-C3 and Trp6-H2 to Tyr3-C3, which verified the C-C bond forming between Tyr3 and Trp6. Meanwhile, the HMBC correlation between Trp6-H2 to Trp6-C$\alpha$ and C$\beta$ proved the C-N bond between Trp6-C2 and Trp6-N. Noesy signal between Trp6-H4 to Tyr3-OH showcasing the close spatial distance between Tyr3 and Trp6, further verifying the complex biaryl motif within YVMW.

**Figure S29.** GpeC enzymatic assay of a series of bioinformatic-guided 9-mer substrates. A) Sequence logo of GpeA-like precursors. B) Based on the biosynthetic logic of the *Gpe* gene cluster and precursor sequence logo, we prioritize 6 precursors that may undergo a similar enzymatic process and chemically synthesized the 9-mer substrates based on the chosen precursor sequences. C) Information about the host of the gene clusters, full-length precursor sequences, and characters in bold represent the 9-mer substrate synthesized as the crosslinking residues marked in red. The conversion ratio is listed. The successful ratio is high, as 4/5 of them can be catalyzed by GpeC. We assumed GpeC was a promiscuous enzyme with a broad substrate scope.

**A**

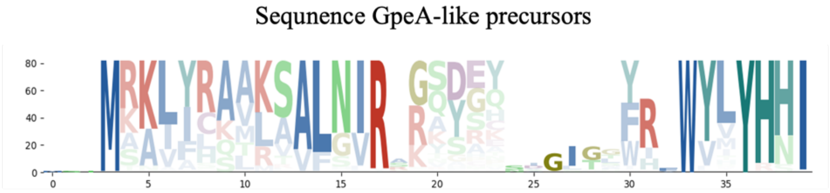


**B**

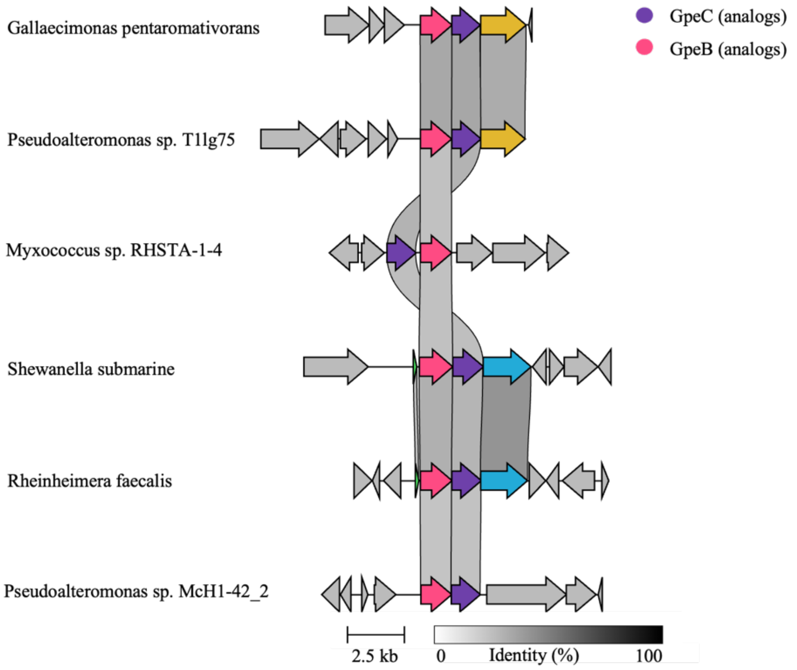


**C**

| HOST | Precursor | Modification | Conversion |
|---|---|---|---|
| >Pseudoalteromonas sp. T1lg75 | MKKIKQAMYRAVGIRSNAFGCNGNNEPW**WRWVHYHNM** | YES | 95% |
| >Myxococcus sp. RHSTA-1-4 | MAKIFRSVMAALSIRRKDPL**YRWYLYHHI** | YES | 33% |
| >Shewanella submarine | MKNLMVSVKNAIGMRASASDGVSSWGCGGRGDIM**WRWYLYHSM** | YES | 76% |
| >Rheinheimera faecalis | MKALQQAFVRALGLRVSVSDQVKDCSASKSSADCC**WRWVHYHDL** | YES | 42% |
| >Pseudoalteromonas sp. McH1-42_2 | MKTLKTKVLRALGVRATSPSQSKR**YLWIMYNNI** | NO | - |

**Figure S30.** GpeC enzymatic assay with WRWVHYHNM. A 5-hour reaction was performed before quenching by adding an equivalent volume of methanol. Conversion of WRWVHYHNM was calculated by the abovementioned method. Tandem mass analysis revealed the peptide ions of $y_2$, $y_3$, $b_2$, and $b_6$-2H, proving the modification happened within the WVHY motif.

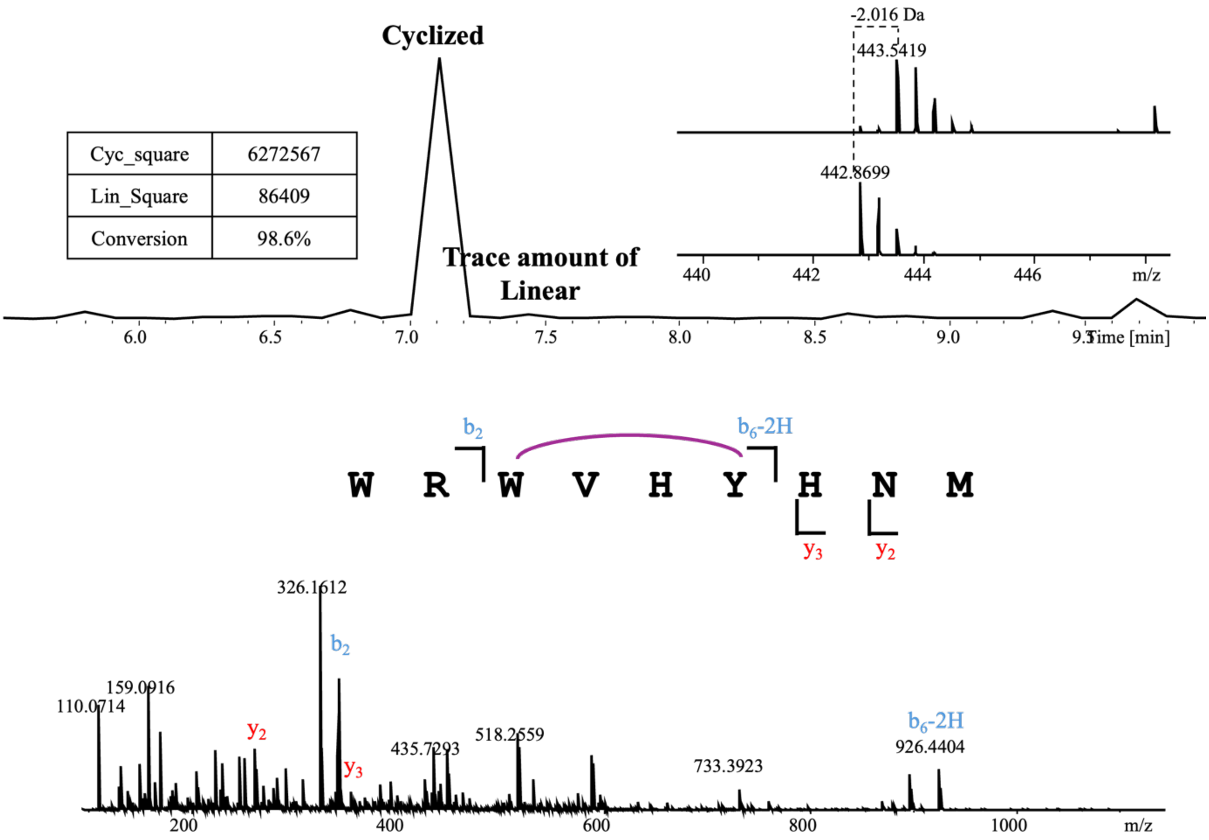


**Cyclized WRWVHYHNM**

| Ions | Fragment | Observed mass | Calculated mass | z | Δppm |
|---|---|---|---|---|---|
| $y_2$ | NM | 264.1015 | 264.1013 | 1 | 0.76 |
| $y_3$ | HNM | 401.1609 | 401.1602 | 1 | 1.74 |
| $b_2$ | WR | 343.1876 | 343.1877 | 1 | 0.29 |
| $b_6$-2H | WRWVHY | 926.4404 | 926.442 | 1 | 1.73 |

**Figure S31.** GpeC enzymatic assay with YRWYLYHHI. A 5-hour reaction was performed before quenching by adding an equivalent volume of methanol. Conversion of YRWYLYHHI was calculated by the abovementioned method. Tandem mass analysis revealed the peptide ions of $y_2$, $y_3$, $b_2$, and $b_6$-2H, proving the modification happened within the WYLY motif.

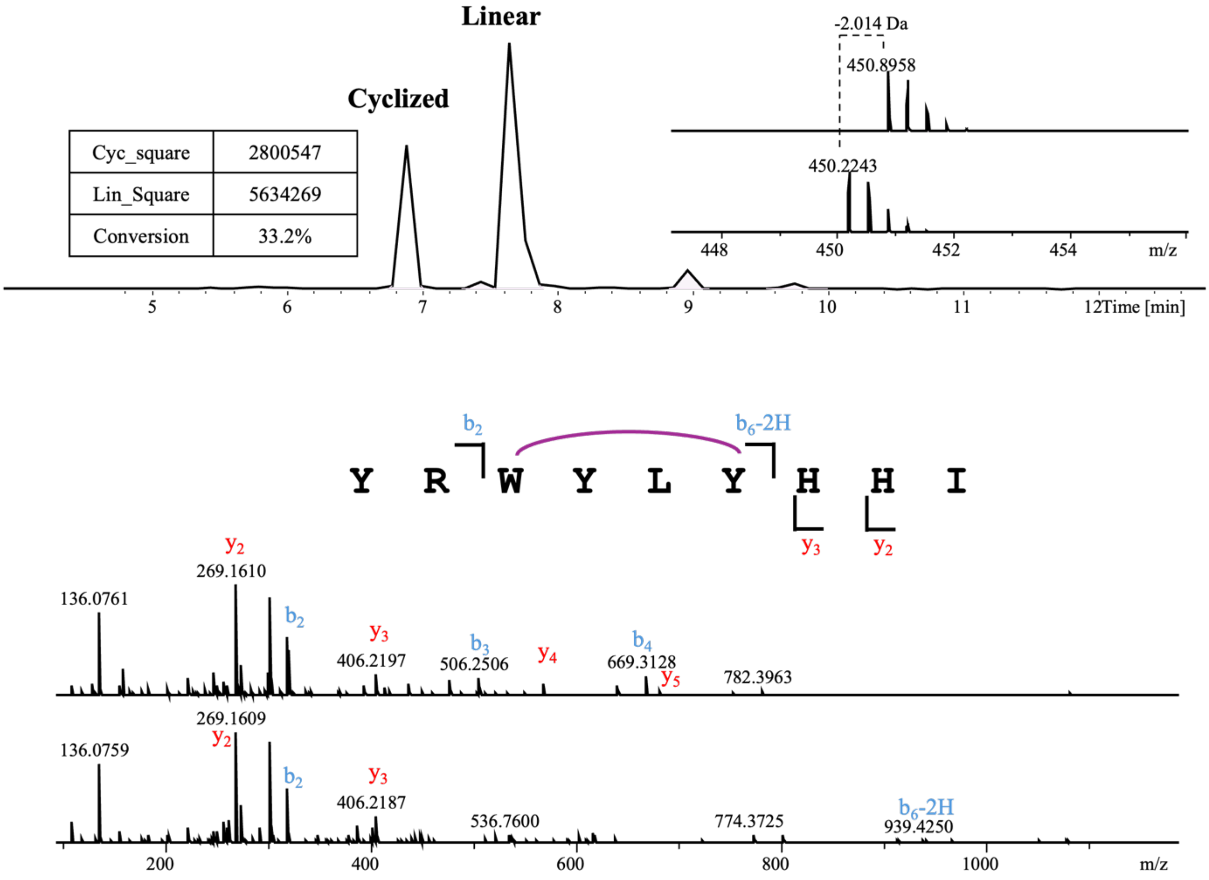


**Linear YRWYLYHHI**

| Ions | Fragment | Observed mass | Calculated mass | z | Δppm |
|---|---|---|---|---|---|
| $y_2$ | HI | 269.1610 | 269.1608 | 1 | 0.74 |
| $y_3$ | HHI | 406.2197 | 406.2197 | 1 | <0.1 |
| $y_4$ | YHHI | 569.2822 | 569.2831 | 1 | 1.58 |
| $y_5$ | LYHHI | 682.3649 | 682.3671 | 1 | 3.22 |
| $b_2$ | YR | 320.1715 | 320.1717 | 1 | 0.62 |
| $b_3$ | YRW | 506.2506 | 506.251 | 1 | 0.79 |
| $b_4$ | YRWY | 669.3128 | 669.3144 | 1 | 2.39 |

**Cyclized YRWYLYHHI**

| Ions | Fragment | Observed mass | Calculated mass | z | Δppm |
|---|---|---|---|---|---|
| $y_2$ | HI | 269.1609 | 269.1608 | 1 | 0.37 |
| $y_3$ | HHI | 406.2187 | 406.2197 | 1 | 2.46 |
| $b_2$ | YR | 320.1716 | 320.1717 | 1 | 0.31 |
| $b_6$-2H | YRWYLY | 943.4446 | 943.4461 | 1 | 1.59 |

**Figure S32.** GpeC enzymatic assay with WRWYLYHSM. A 5-hour reaction was performed before quenching by adding an equivalent volume of methanol. Conversion of WRWYLYHSM was calculated by the abovementioned method. Tandem mass analysis revealed the peptide ions of $y_2$, $y_3$, $b_2$, and $b_6$-2H, proving the modification happened within the WYLY motif.

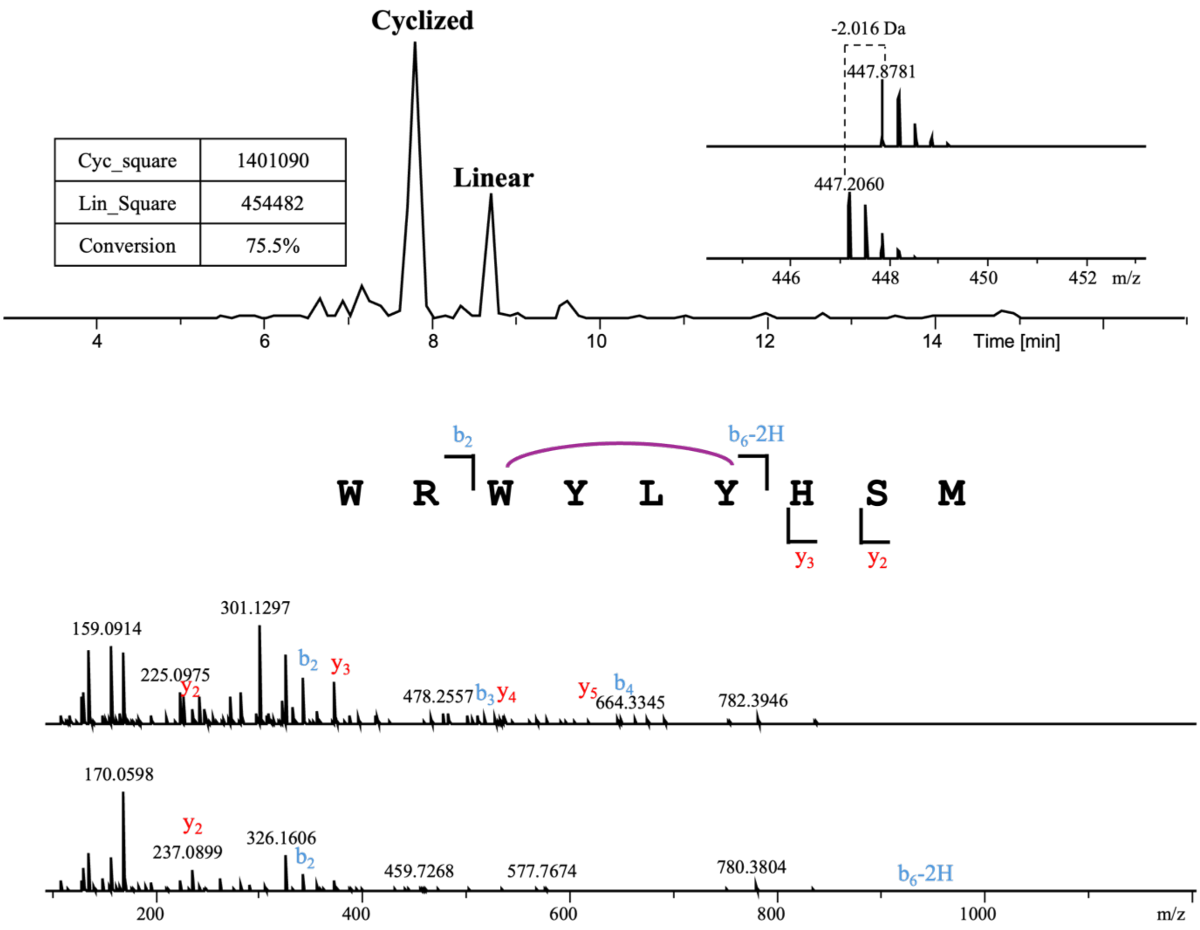


**Linear WRWYLYHSM**

| Ions | Fragment | Observed mass | Calculated mass | z | Δppm |
|---|---|---|---|---|---|
| $y_2$ | SM | 237.0914 | 237.0904 | 1 | 4.22 |
| $y_3$ | HSM | 374.1480 | 374.1493 | 1 | 3.47 |
| $y_4$ | YHSM | 537.2112 | 537.2126 | 1 | 2.61 |
| $y_5$ | LYHSM | 650.2938 | 650.2967 | 1 | 4.46 |
| $b_2$ | WR | 343.1867 | 343.1877 | 1 | 2.91 |
| $b_3$ | WRW | 529.2665 | 529.267 | 1 | 0.94 |
| $b_4$ | WRWY | 692.3276 | 692.3303 | 1 | 3.9 |

**Cyclized WRWYLYHSM**

| Ions | Fragment | Observed mass | Calculated mass | z | Δppm |
|---|---|---|---|---|---|
| $y_2$ | SM | 237.0899 | 237.0904 | 1 | 2.11 |
| $y_3$ | HSM | 374.1485 | 374.1493 | 1 | 2.14 |
| $b_2$ | WR | 343.1865 | 343.1877 | 1 | 3.5 |
| $b_6$-2H | WRWYLY | 966.4606 | 966.4621 | 1 | 1.55 |

**Figure S33.** GpeC enzymatic assay with WRWVHYHDL. A 5-hour reaction was performed before quenching by adding an equivalent volume of methanol. Conversion of WRWVHYHDL was calculated by the abovementioned method. Tandem mass analysis revealed the peptide ions of $y_2$, $y_3$, $b_2$, and $b_6$-2H, proving the modification happened within the WVHY motif.

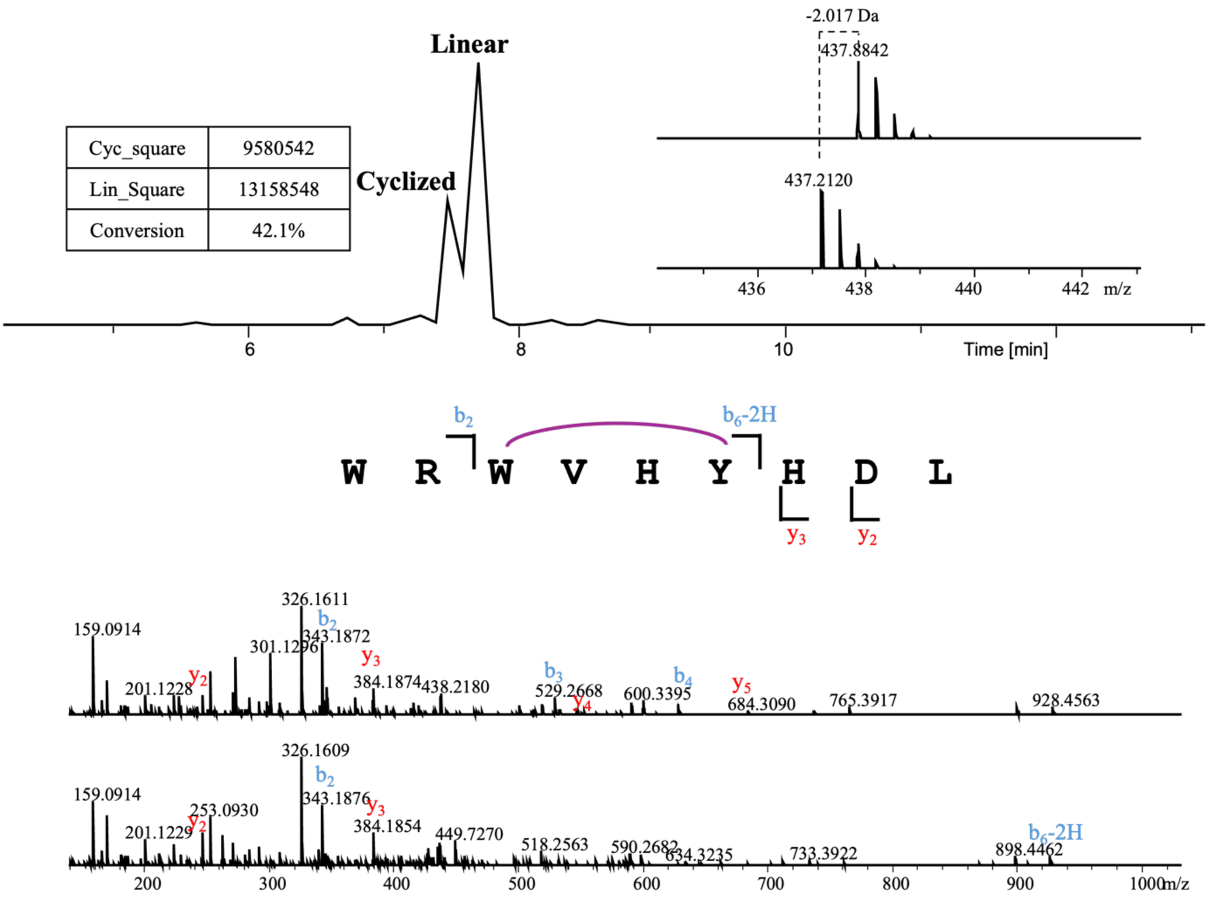


**Linear WRWVHYHDL**

| Ions | Fragment | Observed mass | Calculated mass | z | Δppm |
|---|---|---|---|---|---|
| $y_2$ | DL | 247.1282 | 247.1288 | 1 | 2.43 |
| $y_3$ | HDL | 384.1874 | 384.1878 | 1 | 1.04 |
| $y_4$ | YHDL | 547.2517 | 547.2511 | 1 | 1.1 |
| $y_5$ | HYHDL | 684.3090 | 684.31 | 1 | 1.46 |
| $b_2$ | WR | 343.1876 | 343.1877 | 1 | 0.29 |
| $b_3$ | WRW | 529.2668 | 529.267 | 1 | 0.38 |
| $b_4$ | WRWV | 628.3347 | 628.3354 | 1 | 1.11 |

**Cyclized WRWVHYHDL**

| Ions | Fragment | Observed mass | Calculated mass | z | Δppm |
|---|---|---|---|---|---|
| $y_2$ | DL | 247.1290 | 247.1288 | 1 | 0.81 |
| $y_3$ | HDL | 384.1854 | 384.1878 | 1 | 6.25 |
| $b_2$ | WR | 343.1872 | 343.1877 | 1 | 1.46 |
| $b_6$-2H | WRWVHY | 926.4406 | 926.442 | 1 | 1.51 |

**Figure S34.** GpeC enzymatic assay with YIWIMYNNI. A 5-hour reaction was performed before quenching by adding an equivalent volume of methanol. Conversion of YIWIMYNNI was not observed, indicating the unsuccessful catalytic reaction. Tandem mass analysis revealed the linear peptide ions of $y_2$, $y_3$, $y_4$, $y_5$, $b_2$, $b_3$, and $b_4$.

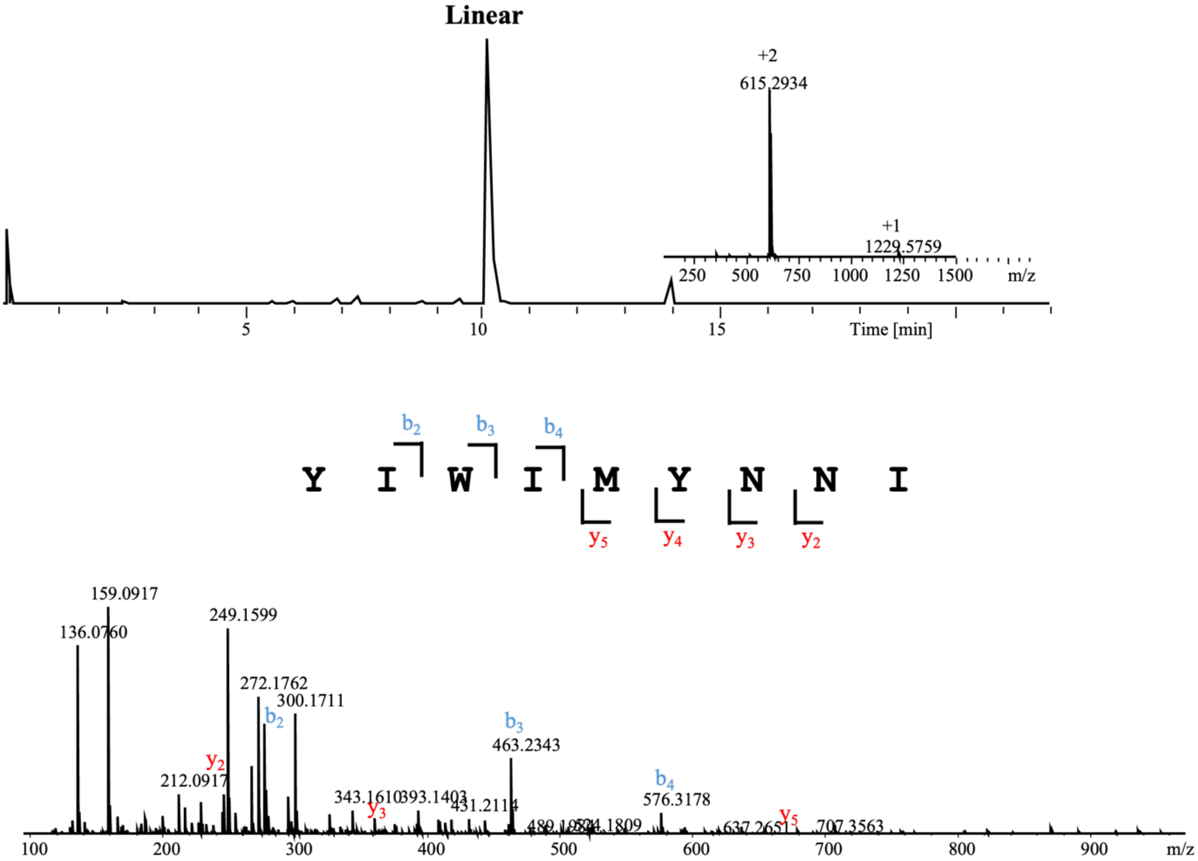


| Linear YIWIMYNNI | | | | | |
|---|---|---|---|---|---|
| **Ions** | **Fragment** | **Observed mass** | **Calculated mass** | **z** | **Δppm** |
| $y_2$ | NI | 246.1445 | 246.1448 | 1 | 2.43 |
| $y_3$ | NNI | 360.1876 | 360.1878 | 1 | 1.04 |
| $y_5$ | MYNNI | 654.2918 | 654.2916 | 1 | 1.46 |
| $b_2$ | YI | 277.1553 | 277.1547 | 1 | 0.29 |
| $b_3$ | YIW | 463.2313 | 463.234 | 1 | 0.38 |
| $b_4$ | YIWI | 576.3178 | 576.318 | 1 | 1.11 |

**Figure S35.** General synthesis method for the substrate scope characterization. We grouped the 20 canonical amino acids into 5 groups by their side chain properties. Peptide synthesis was performed as the abovementioned method.

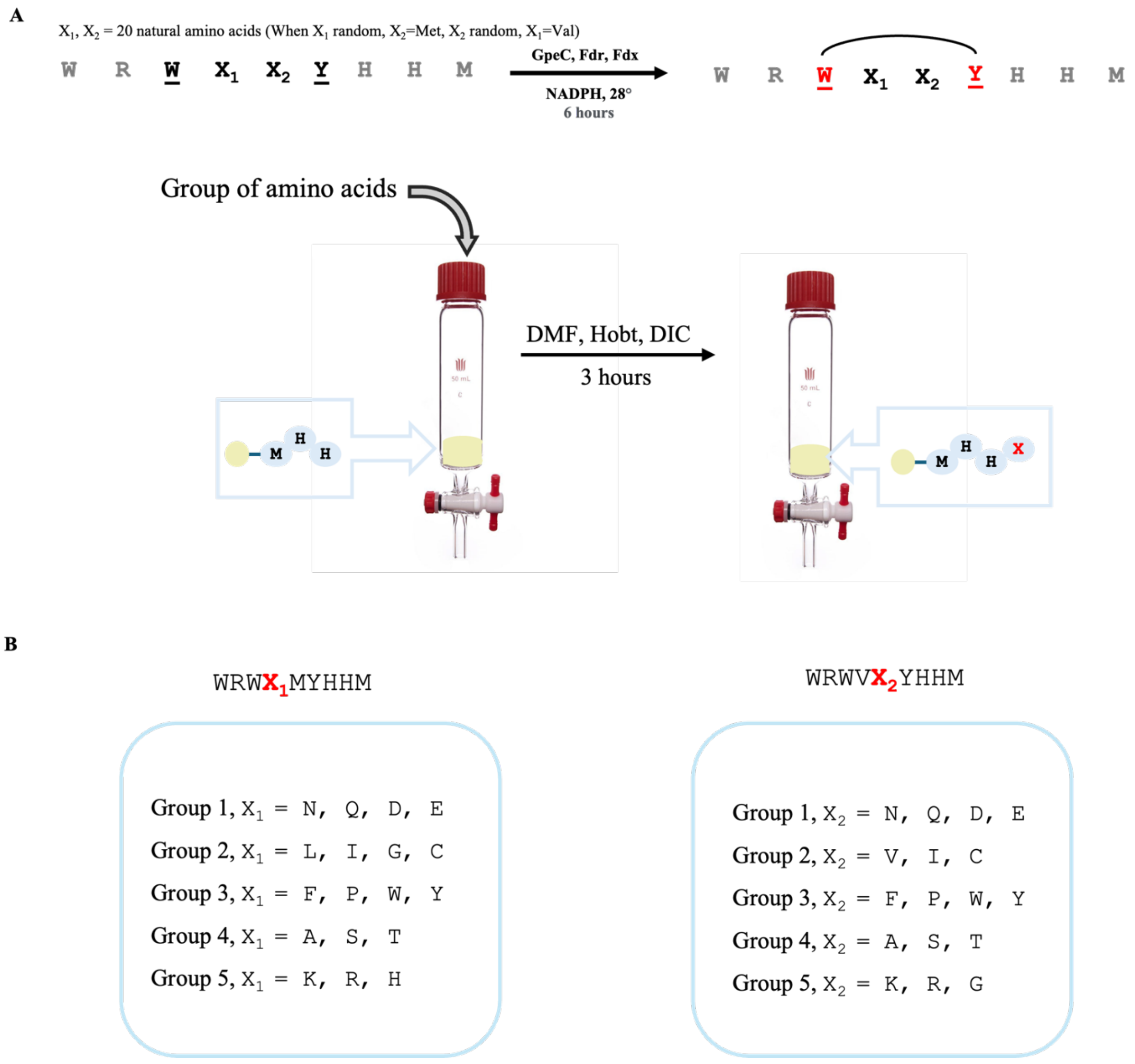

**Figure S36.** Conformation of GpeC with substrates WRWPMYHHM and WRWVPYHHM

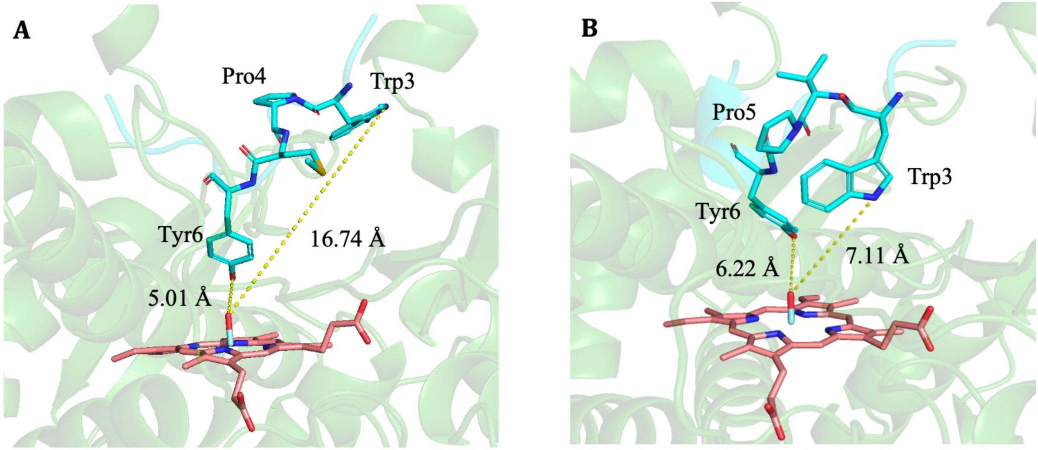


GpeC with WRWPMYHHM GpeC with WRWVPYHHM

A) Conformation of GpeC with substrates WRWPMYHHM, Trp3 and Tyr6 are placed far from the catalytic center, leading to the inefficient catalysis. B) Conformation of GpeC with substrates WRWVPYHHM, Trp3 and Tyr6 are placed far from the catalytic center, leading to the inefficient catalysis.

**Figure S37.** GpeC enzymatic assay with WRWYMYHHM (**25-X₁**). A 5-hour reaction was performed before quenching by adding an equivalent volume of methanol. Conversion of (**25-X₁**) was calculated by the abovementioned method. Tandem mass analysis revealed the peptide ions of $y_2$, $y_3$, $b_2$, and $b_6$-2H, proving the modification happened within the WYMY motif.

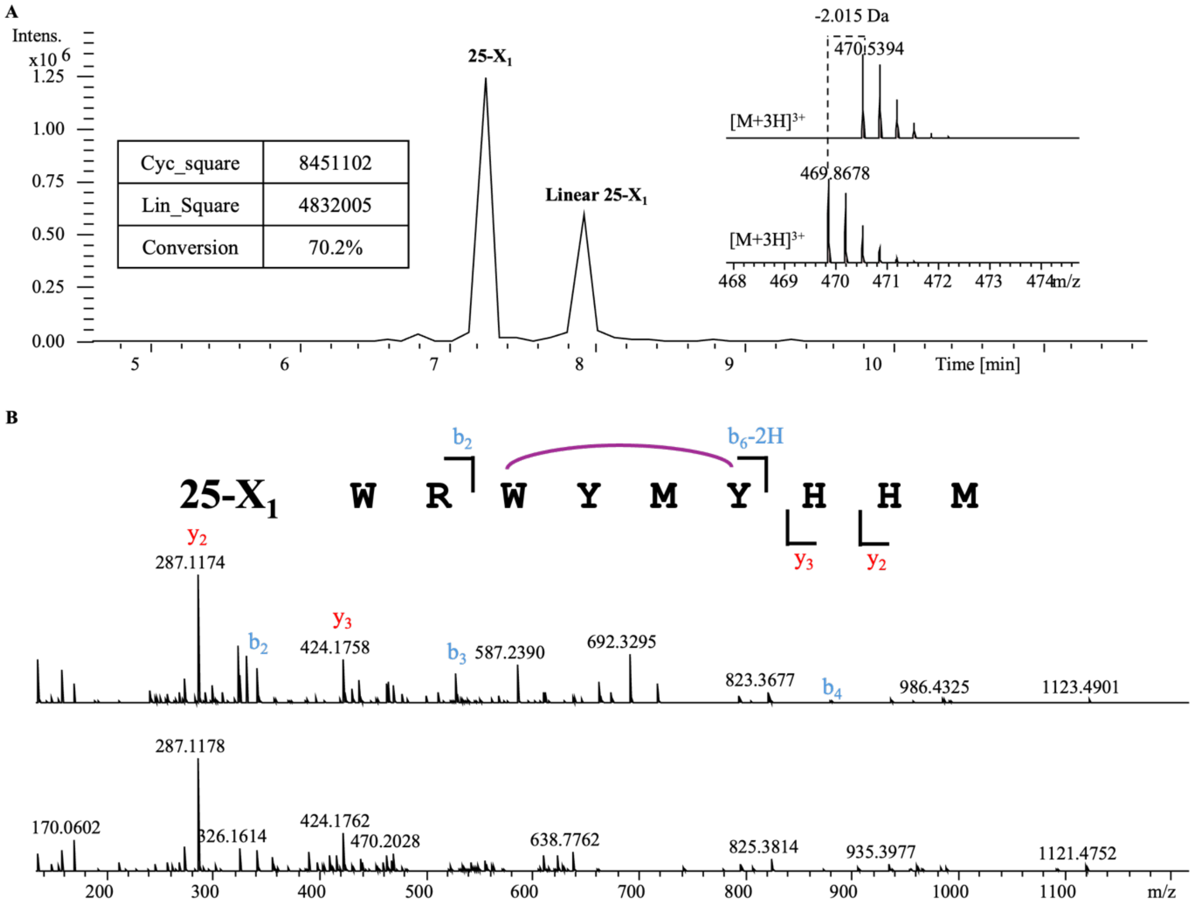


**Linear 25-X₁**

| Ions | Fragment | Observed mass | Calculated mass | z | Δppm |
|---|---|---|---|---|---|
| $y_2$ | HM | 287.1174 | 287.1172 | 1 | 1.39 |
| $y_3$ | HHM | 424.1758 | 424.1761 | 1 | 0.24 |
| $y_4$ | YHHM | 587.2398 | 587.2395 | 1 | 0.51 |
| $y_5$ | MYHHM | 644.2607 | 644.2609 | 1 | 0.31 |
| $b_2$ | WR | 343.1874 | 343.1877 | 1 | 0.87 |
| $b_3$ | WRW | 529.2670 | 529.267 | 1 | <0.1 |
| $b_6$ | WRWYMY | 848.4210 | 848.4202 | 1 | 0.94 |

**25-X₁**

| Ions | Fragment | Observed mass | Calculated mass | z | Δppm |
|---|---|---|---|---|---|
| $y_2$ | HM | 287.1178 | 287.1172 | 1 | 1.39 |
| $y_3$ | HHM | 424.1762 | 424.1761 | 1 | 1.89 |
| $b_2$ | WR | 343.1874 | 343.1877 | 1 | 0.87 |
| $b_6$-2H | WRWYMY | 846.4042 | 846.4046 | 1 | 0.47 |

**Figure S38.** GpeC enzymatic assay with WRWKMYHHM (**10-X**$_1$). A 5-hour reaction was performed before quenching by adding an equivalent volume of methanol. Conversion of (**10-X**$_1$) was calculated by the abovementioned method. Tandem mass analysis revealed the peptide ions of $y_2$, $y_3$, $b_2$, and $b_6$-2H, proving the modification happened within the WKMY motif.

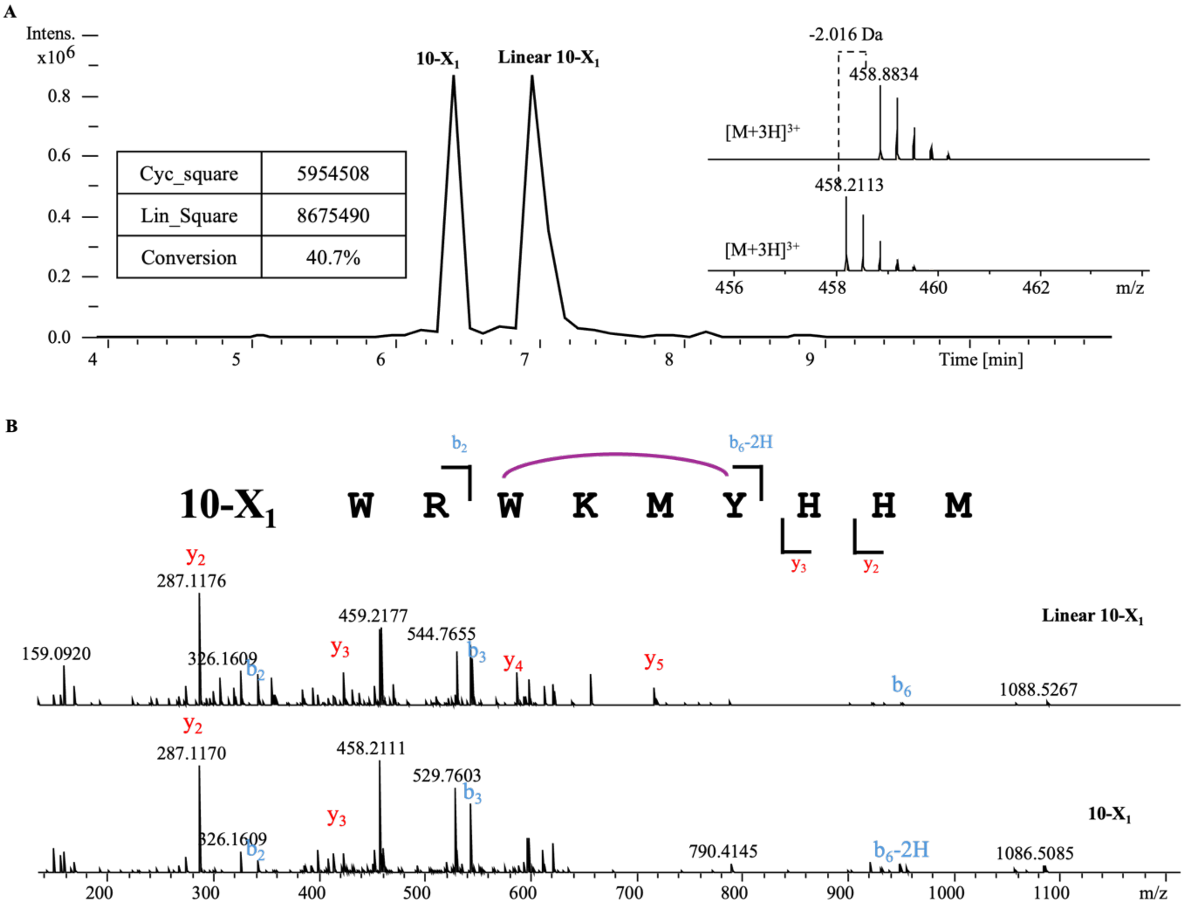


**Linear 10**

| Ions | Fragment | Observed mass | Calculated mass | z | Δppm |
|---|---|---|---|---|---|
| $y_2$ | HM | 287.1176 | 287.1172 | 1 | 1.39 |
| $y_3$ | HHM | 424.1759 | 424.1761 | 1 | 0.47 |
| $y_4$ | YHHM | 587.2392 | 587.2395 | 1 | 0.51 |
| $y_5$ | MYHHM | 718.2795 | 718.28 | 1 | 0.7 |
| $b_2$ | WR | 343.1872 | 343.1877 | 1 | 1.46 |
| $b_3$ | WRW | 529.2663 | 529.267 | 1 | 1.32 |
| $b_6$ | WRWKMY | 951.4665 | 951.4658 | 1 | 0.74 |

**10**

| Ions | Fragment | Observed mass | Calculated mass | z | Δppm |
|---|---|---|---|---|---|
| $y_2$ | HM | 287.1170 | 287.1172 | 1 | 0.7 |
| $y_3$ | HHM | 424.1755 | 424.1761 | 1 | 1.41 |
| $b_2$ | WR | 343.1878 | 343.1877 | 1 | 0.29 |
| $b_6$-2H | WRWKMY | 949.4536 | 949.4501 | 1 | 3.69 |

**Figure S39.** GpeC enzymatic assay with WRWVKYHHM(**10-X₂**). A 5-hour reaction was performed before quenching by adding an equivalent volume of methanol. Almost all of the linear substrate was catalyzed to the biaryl cyclized product (**10-X₂**). Tandem mass analysis revealed the peptide ions of $y_2$, $y_3$, $b_2$, and $b_6$-2H, proving the modification happened within the WVKY motif.

A

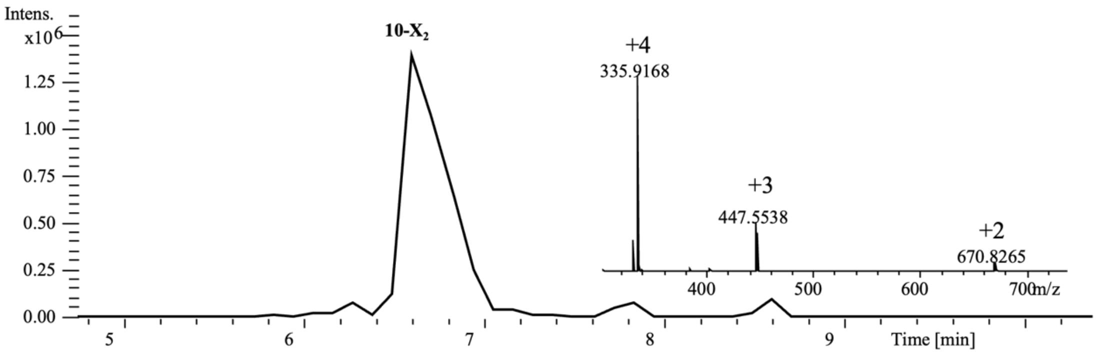


B

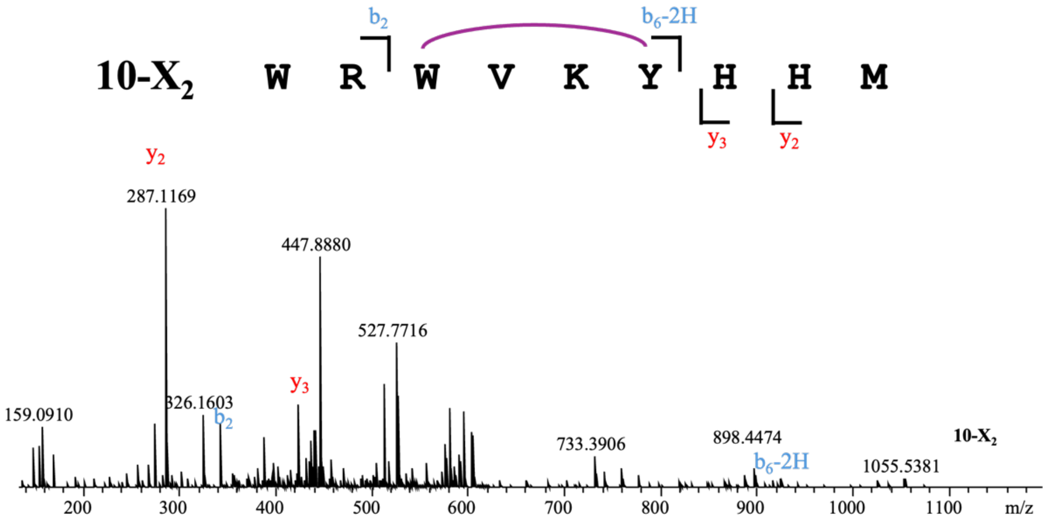


| 10-X₂ | | | | | |
|---|---|---|---|---|---|
| Ions | Fragment | Observed mass | Calculated mass | z | Δppm |
| $y_2$ | HM | 287.1169 | 287.1172 | 1 | 1.04 |
| $y_3$ | HHM | 424.1764 | 424.1761 | 1 | 0.71 |
| $b_2$ | WR | 343.1876 | 343.1877 | 1 | 0.29 |
| $b_6$-2H | WRWVKY | 917.4767 | 917.4781 | 1 | 1.53 |

**Figure S40.** GpeC enzymatic assay with WRWVEYHHM (**13-X₂**). A 5-hour reaction was performed before quenching by adding an equivalent volume of methanol. Tandem mass analysis revealed the peptide ions of $y_2$, $y_3$, $b_2$, and $b_6$-2H, proving the modification happened within the WVEY motif.

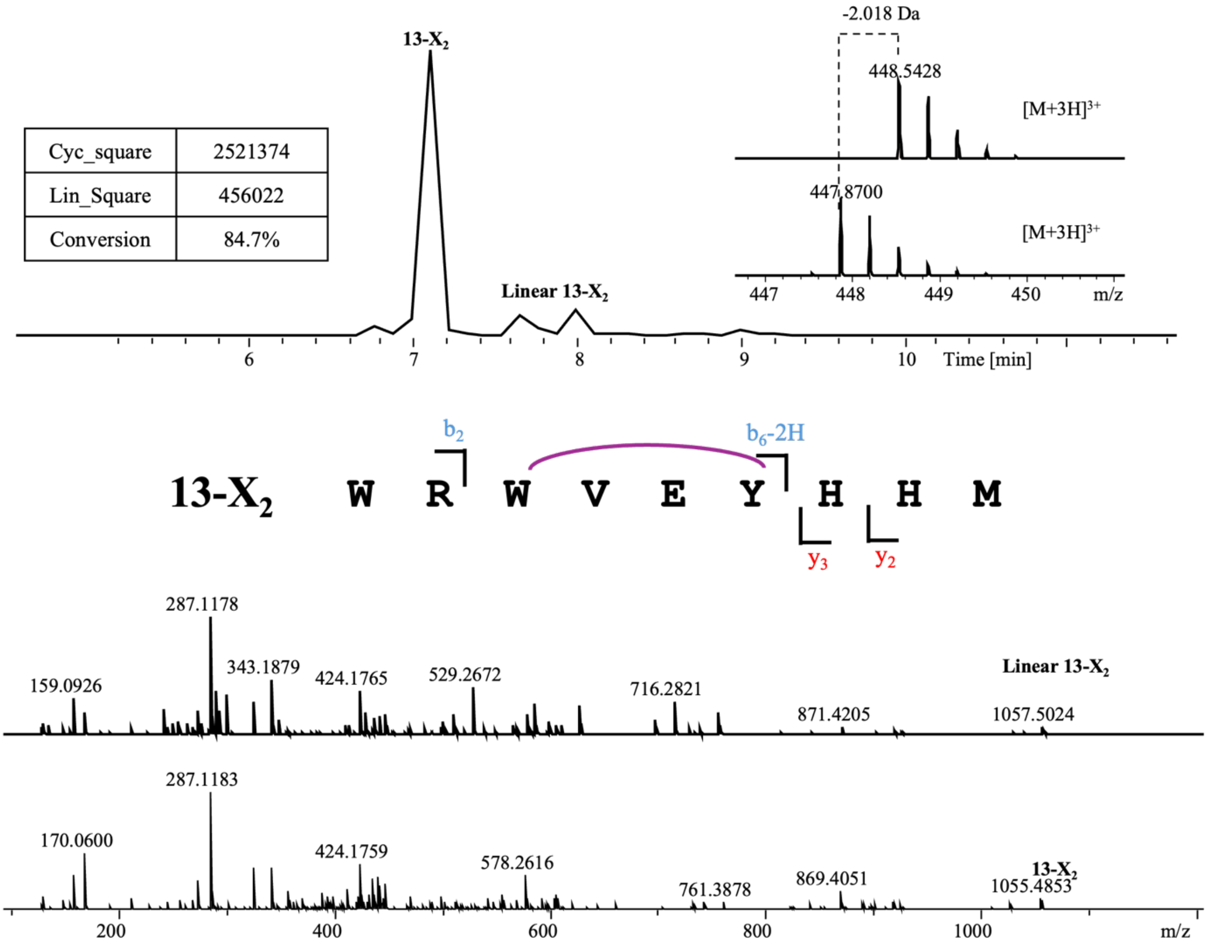


**Linear 13-X₂**

| Ions | Fragment | Observed mass | Calculated mass | z | Δppm |
|---|---|---|---|---|---|
| $y_2$ | HM | 287.1178 | 287.1172 | 1 | 2.09 |
| $y_3$ | HHM | 424.1765 | 424.1761 | 1 | 0.94 |
| $y_4$ | YHHM | 587.2400 | 587.2395 | 1 | 0.85 |
| $y_5$ | EYHHM | 716.2821 | 716.2821 | 1 | 0 |
| $b_2$ | WR | 343.1879 | 343.1877 | 1 | 0.58 |
| $b_3$ | WRW | 529.2672 | 529.267 | 1 | 0.38 |
| $b_6$ | WRWVEY | 920.4422 | 920.4413 | 1 | 0.98 |

**13-X₂**

| Ions | Fragment | Observed mass | Calculated mass | z | Δppm |
|---|---|---|---|---|---|
| $y_2$ | HM | 287.1183 | 287.1172 | 1 | 3.83 |
| $y_3$ | HHM | 424.1759 | 424.1761 | 1 | 0.47 |
| $b_2$ | WR | 343.1871 | 343.1877 | 1 | 1.75 |
| $b_6$-2H | WRWVEY | 918.4272 | 918.4257 | 1 | 1.63 |

**Figure S41.** GpeC enzymatic assay with WRWV-Dmet-YHHM (**27**). A 5-hour reaction was performed before quenching by adding an equivalent volume of methanol. Tandem mass analysis revealed the peptide ions of $y_3$, $b_2$, and $b_6$-2H, proving the modification happened within the WV-Dmet-Y motif.

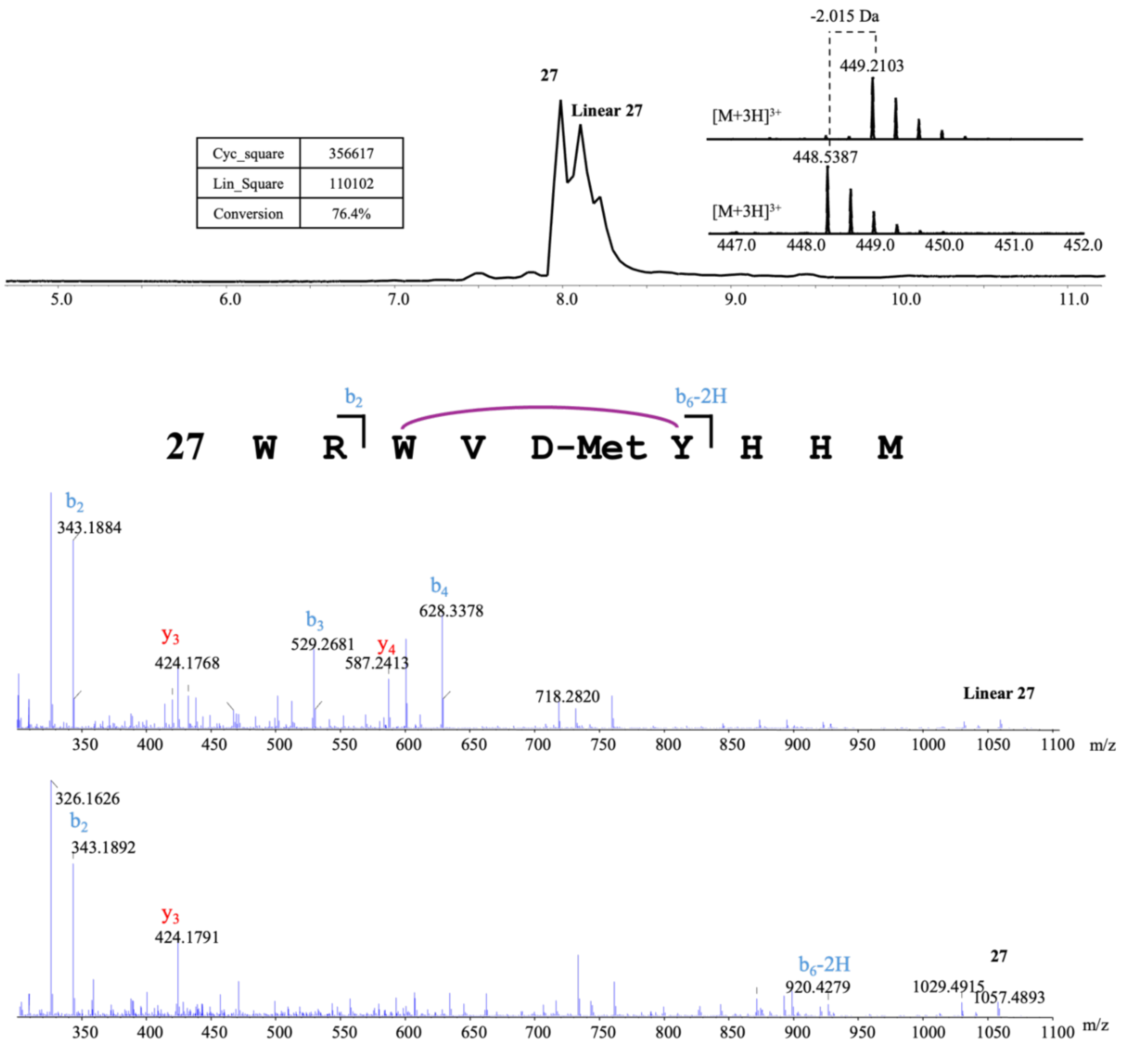


**Linear 27**

| Ions | Fragment | Observed mass | Calculated mass | z | Δppm |
|---|---|---|---|---|---|
| $y_3$ | HHM | 424.1768 | 424.1761 | 1 | 1.65 |
| $y_4$ | YHHM | 587.2413 | 587.2395 | 1 | 3.07 |
| $b_2$ | WR | 343.1884 | 343.1877 | 1 | 2.04 |
| $b_3$ | WRW | 529.2681 | 529.267 | 1 | 2.08 |
| $b_4$ | WRWV | 628.3378 | 628.3354 | 1 | 3.82 |

**27**

| Ions | Fragment | Observed mass | Calculated mass | z | Δppm |
|---|---|---|---|---|---|
| $y_3$ | HHM | 424.1791 | 424.1761 | 1 | 7.07 |
| $b_2$ | WR | 343.1892 | 343.1877 | 1 | 4.37 |
| $b_6$-2H | WRWV-DMet-Y | 920.4279 | 920.4236 | 1 | 4.67 |

**Figure S42.** GpeC enzymatic assay with WRWV-(3$NO_2$)Y-YHHM (**28**). A 5-hour reaction was performed before quenching by adding an equivalent volume of methanol. One isomer was formed, marked as (**36-1**). Tandem mass analysis revealed the peptide ions of $y_3$, $b_2$, and $b_6$-2H, proving the modification happened within the WV-(3$NO_2$)Y-Ymotif.

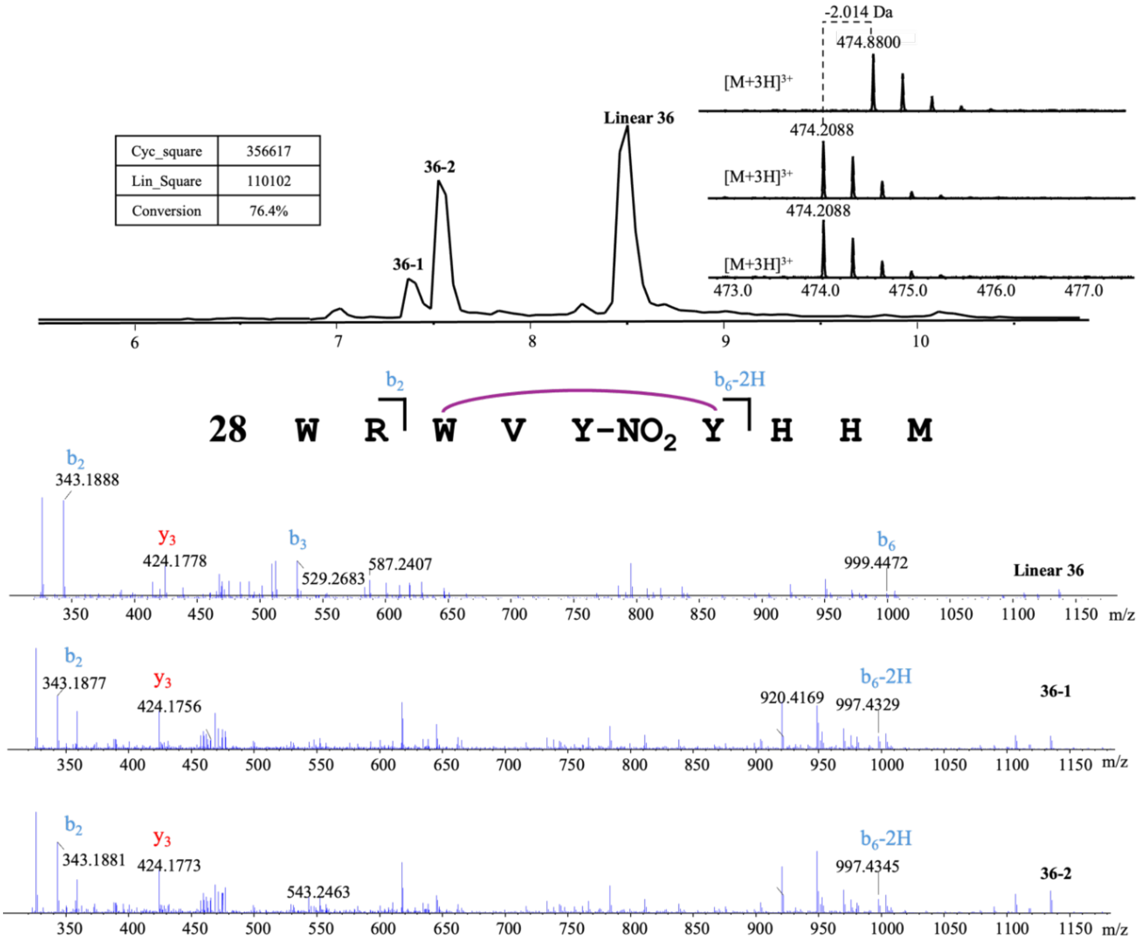


**Linear 36**

| Ions | Fragment | Observed mass | Calculated mass | z | Δppm |
|---|---|---|---|---|---|
| $y_3$ | HHM | 424.1778 | 424.1761 | 1 | 4.01 |
| $b_2$ | WR | 343.1888 | 343.1877 | 1 | 3.21 |
| $b_3$ | WRW | 529.2683 | 529.267 | 1 | 2.46 |
| $b_6$ | WRWV-(3$NO_2$)Y-Y | 999.4518 | 999.4472 | 1 | 4.6 |

**36-1**

| Ions | Fragment | Observed mass | Calculated mass | z | Δppm |
|---|---|---|---|---|---|
| $y_3$ | HHM | 424.1756 | 424.1761 | 1 | 1.18 |
| $b_2$ | WR | 343.1877 | 343.1877 | 1 | <0.1 |
| $b_6$-2H | WV-(3$NO_2$)Y-YHHM | 997.4329 | 997.4315 | 1 | 1.4 |

**36-2**

| Ions | Fragment | Observed mass | Calculated mass | z | Δppm |
|---|---|---|---|---|---|
| $y_3$ | HHM | 424.1773 | 424.1761 | 1 | 2.83 |
| $b_2$ | WR | 343.1881 | 343.1877 | 1 | 1.17 |
| $b_6$-2H | WV-(3$NO_2$)Y-YHHM | 997.4366 | 997.4315 | 1 | 5.11 |

**Figure S43.** GpeC enzymatic assay with WRWV-4OH-Dphg-Y-HHM (**29**). A 5-hour reaction was performed before quenching by adding an equivalent volume of methanol. Tandem mass analysis revealed the peptide ions of $y_3$, $b_2$, and $b_6$-2H, proving the modification happened within the WV-4OH-Dphg-Y motif.

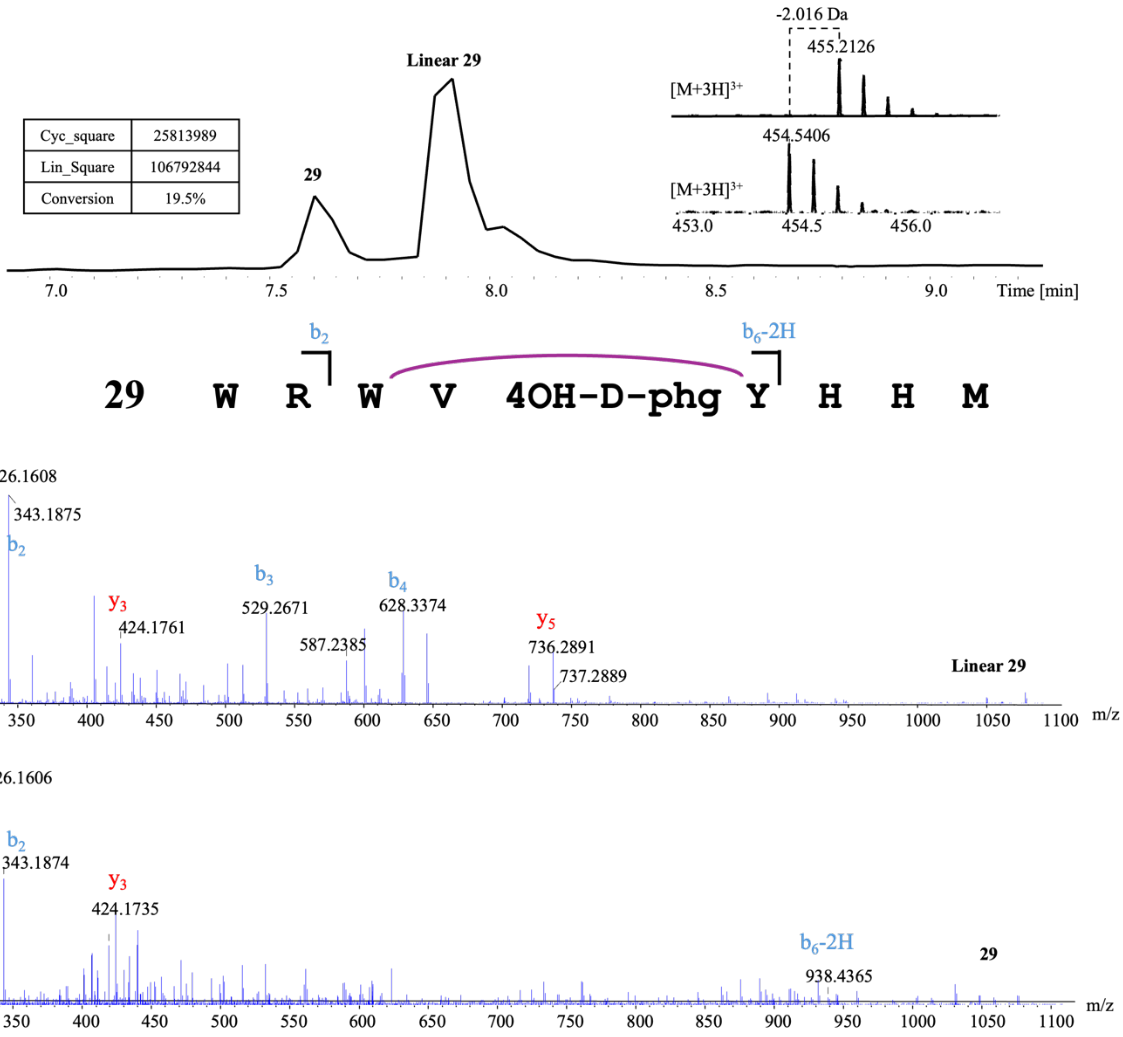


| Linear 29 | | | | | |
|---|---|---|---|---|---|
| Ions | Fragment | Observed mass | Calculated mass | z | Δppm |
| $y_3$ | HHM | 424.1768 | 424.1761 | 1 | 1.6 |
| $y_5$ | 4OH-Dphg-YHHM | 736.2891 | 736.2872 | 1 | 2.58 |
| $b_2$ | WR | 343.1875 | 343.1877 | 1 | 0.58 |
| $b_3$ | WRW | 529.2671 | 529.267 | 1 | 0.18 |
| $b_4$ | WRWV | 628.3374 | 628.3354 | 1 | 3.18 |

| 29 | | | | | |
|---|---|---|---|---|---|
| Ions | Fragment | Observed mass | Calculated mass | z | Δppm |
| $y_3$ | HHM | 424.1735 | 424.1761 | 1 | 6.13 |
| $b_2$ | WR | 343.1874 | 343.1877 | 1 | 0.87 |
| $b_6$-2H | WRWV-4OH-Dphg-Y | 938.4365 | 938.4308 | 1 | 6.07 |

**Figure S44.** GpeC enzymatic assay with WRWV-d-Propargylglycine-YHHM (**31**). A 5-hour reaction was performed before quenching by adding an equivalent volume of methanol. Conversion of (**31**) was calculated by the abovementioned method. Tandem mass analysis revealed the peptide ions of $y_2$, $y_3$, $b_2$, and $b_7$-2H, proving the modification happened within the WV- d-Propargylglycine-Y motif.

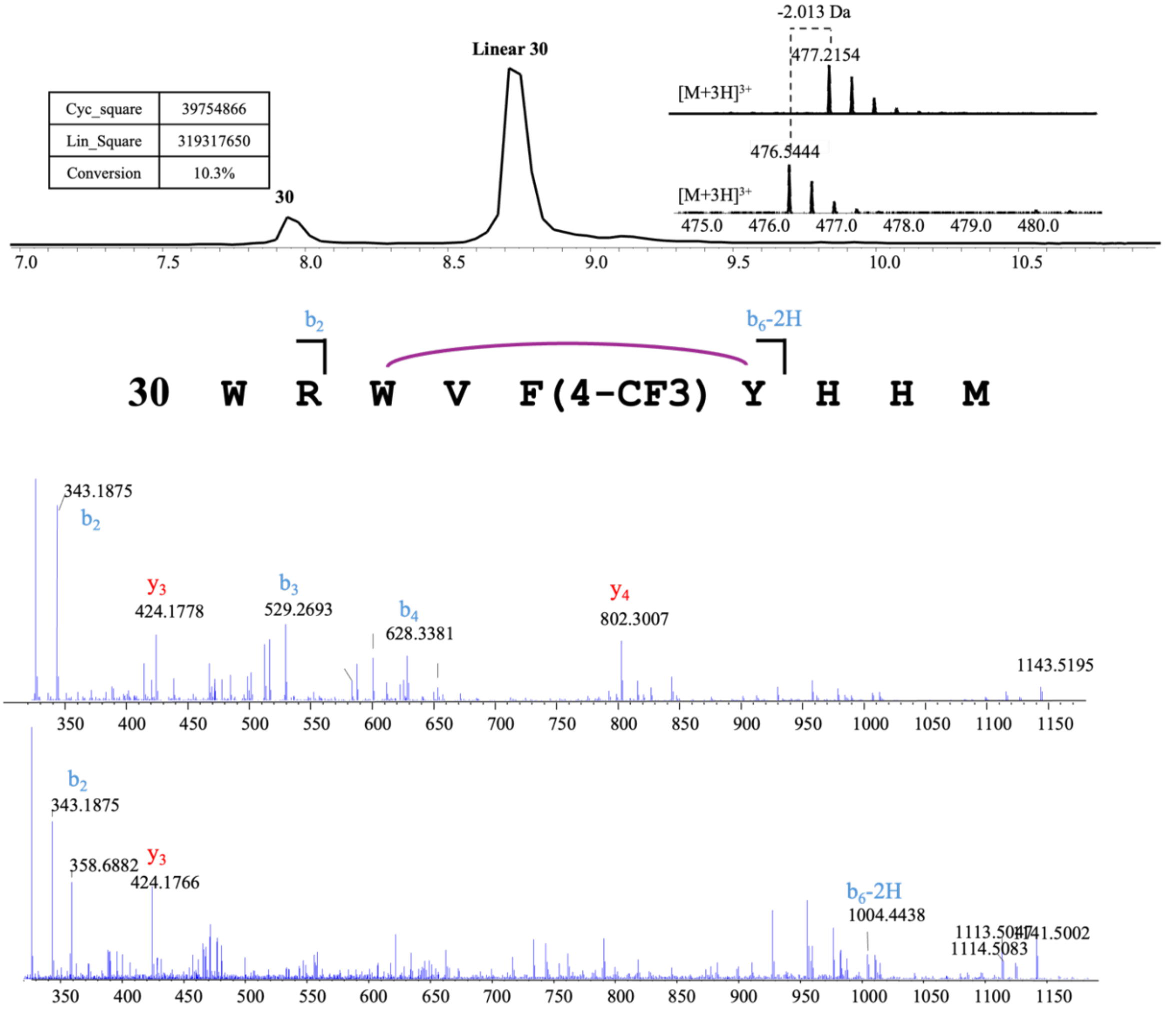


**Linear 30**

| Ions | Fragment | Observed mass | Calculated mass | z | Δppm |
|---|---|---|---|---|---|
| $y_3$ | HHM | 424.1778 | 424.1761 | 1 | 4.01 |
| $y_4$ | 4OH-Dphg-YHHM | 802.3007 | 802.2953 | 1 | 6.73 |
| $b_2$ | WR | 343.1888 | 343.1877 | 1 | 3.21 |
| $b_3$ | WRW | 529.2693 | 529.267 | 1 | 4.35 |
| $b_4$ | WRWV | 628.3381 | 628.3354 | 1 | 4.3 |

**30**

| Ions | Fragment | Observed mass | Calculated mass | z | Δppm |
|---|---|---|---|---|---|
| $y_3$ | HHM | 424.1766 | 424.1761 | 1 | 1.18 |
| $b_2$ | WR | 343.1875 | 343.1877 | 1 | 0.58 |
| $b_6$-2H | WRWV-4OH-Dphg-Y | 1004.4438 | 1004.4389 | 1 | 4.88 |

**Figure S45.** GpeC enzymatic assay with WRWV-d-Propargylglycine-YHHM (**31**). A 5-hour reaction was performed before quenching by adding an equivalent volume of methanol. Conversion of (**31**) was calculated by the abovementioned method. Tandem mass analysis revealed the peptide ions of $y_2$, $y_3$, $b_2$, and $b_7$-2H, proving the modification happened within the WV- d-Propargylglycine-Y motif.

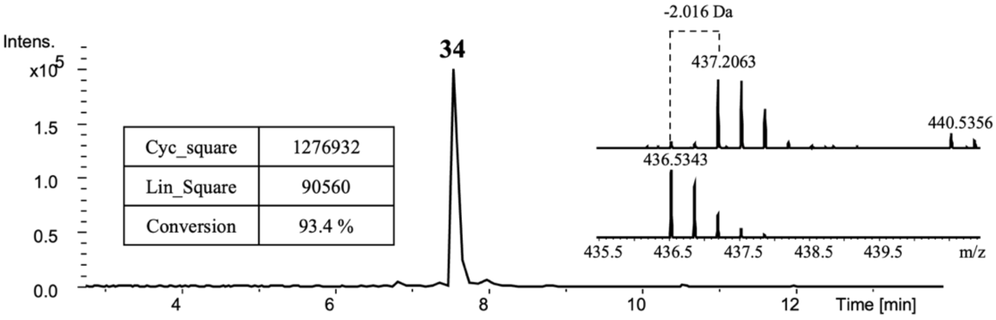


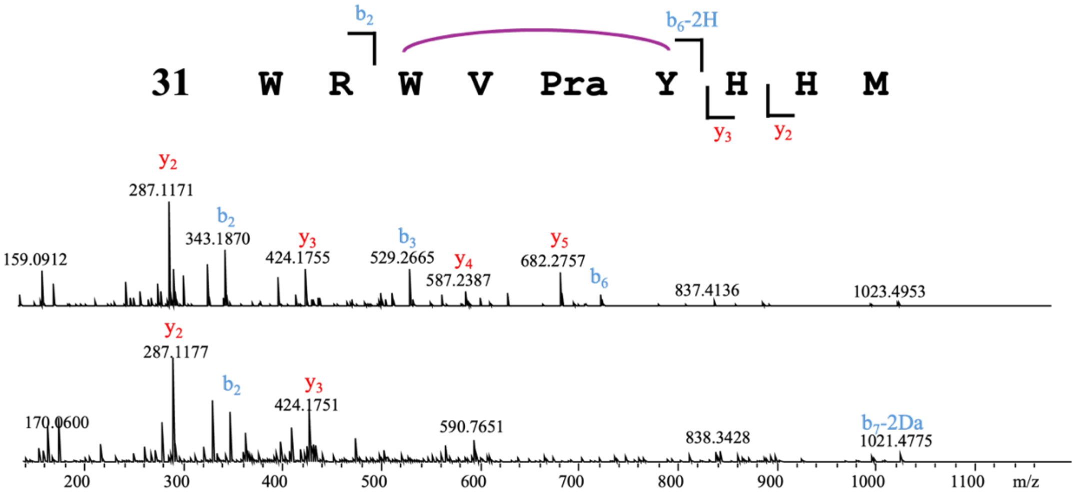


**Linear 34 (Ref.)**

| Ions | Fragment | Observed mass | Calculated mass | z | Δppm |
|---|---|---|---|---|---|
| $y_2$ | HM | 287.1171 | 287.1172 | 1 | 0.35 |
| $y_3$ | HHM | 424.1755 | 424.1761 | 1 | 1.41 |
| $y_4$ | YHHM | 587.2387 | 587.2395 | 1 | 1.36 |
| $y_5$ | d-Pra-YHHM | 682.2757 | 682.2715 | 1 | 6.16 |
| $b_2$ | WR | 343.1870 | 343.1877 | 1 | 2.04 |
| $b_3$ | WRW | 529.2665 | 529.267 | 1 | 0.94 |
| $b_6$ | WRWV-d-Pra-Y | 723.3723 | 723.3731 | 1 | 1.11 |

**Cyclized 34**

| Ions | Fragment | Observed mass | Calculated mass | z | Δppm |
|---|---|---|---|---|---|
| $y_2$ | HM | 287.1177 | 287.1172 | 1 | 1.74 |
| $y_3$ | HHM | 424.1751 | 424.1761 | 1 | 2.36 |
| $b_2$ | WR | 343.1870 | 343.1877 | 1 | 2.04 |
| $b_6$-2H | WRWHMY | 1021.4788 | 1021.4790 | 1 | 0.2 |

**Figure S46.** GpeC enzymatic assay with WRWV- L- Homoallylglycine-YHHM (**32**). A 5-hour reaction was performed before quenching by adding an equivalent volume of methanol. Conversion of (**32**) was calculated by the the abovementioned method. Tandem mass analysis revealed the peptide ions of $y_2$, $y_3$, $b_2$, and $b_6$-2H, proving the modification happened within the WV- L-Homoallylglycine-Y motif.

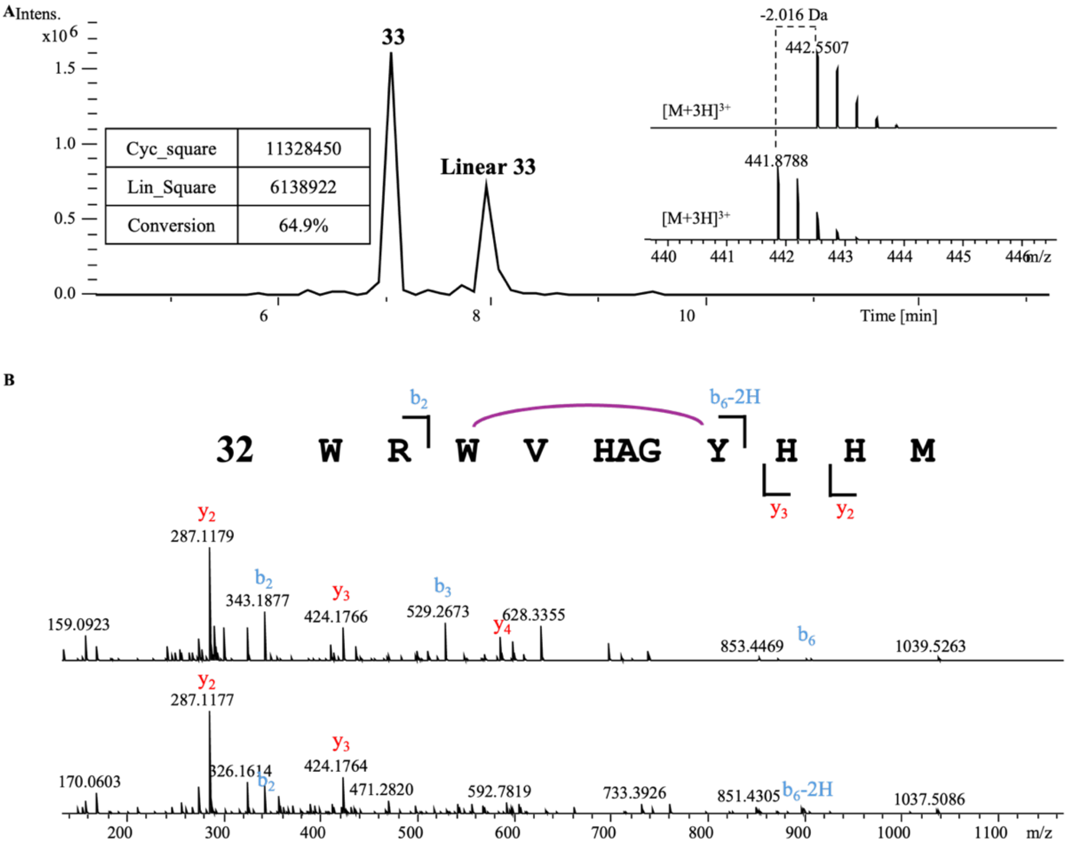


**Linear 33**

| Ions | Fragment | Observed mass | Calculated mass | z | Δppm |
|---|---|---|---|---|---|
| $y_2$ | HM | 287.1179 | 287.1172 | 1 | 2.44 |
| $y_3$ | HHM | 424.1766 | 424.1761 | 1 | 1.18 |
| $y_4$ | YHHM | 587.2390 | 587.2395 | 1 | 0.85 |
| $b_2$ | WR | 343.1877 | 343.1877 | 1 | <0.1 |
| $b_3$ | WRW | 529.2673 | 529.267 | 1 | 0.57 |
| $b_6$ | WRWV-L-HAG-Y | 902.4672 | 902.4672 | 1 | <0.1 |

**33**

| Ions | Fragment | Observed mass | Calculated mass | z | Δppm |
|---|---|---|---|---|---|
| $y_2$ | HM | 287.1177 | 287.1172 | 1 | 1.74 |
| $y_3$ | HHM | 424.1764 | 424.1761 | 1 | 0.71 |
| $b_2$ | WR | 343.1876 | 343.1877 | 1 | 0.29 |
| $b_6$-2H | WRWV-L-HAG-Y | 900.4513 | 900.4513 | 1 | <0.1 |

**Figure S47.** GpeC enzymatic assay with WRWV- L-4-Chlorophenylglycine-YHHM (**33**). A 5-hour reaction was performed before quenching by adding an equivalent volume of methanol. Conversion of (**35**) was calculated by the abovementioned method. Tandem mass analysis revealed the peptide ions of $y_2$, $y_3$, $b_2$, and $b_6$-2H, proving the modification happened within the WV-L-4-Chlorophenylglycine-Y -Y motif.

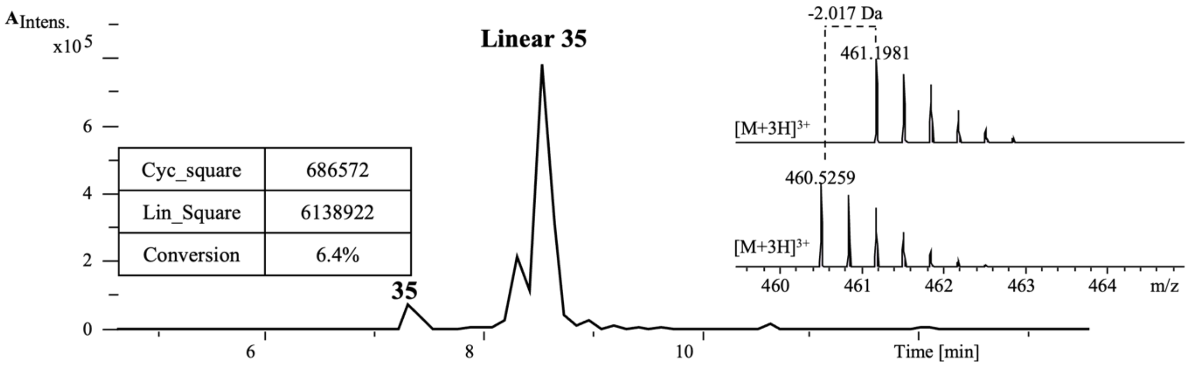


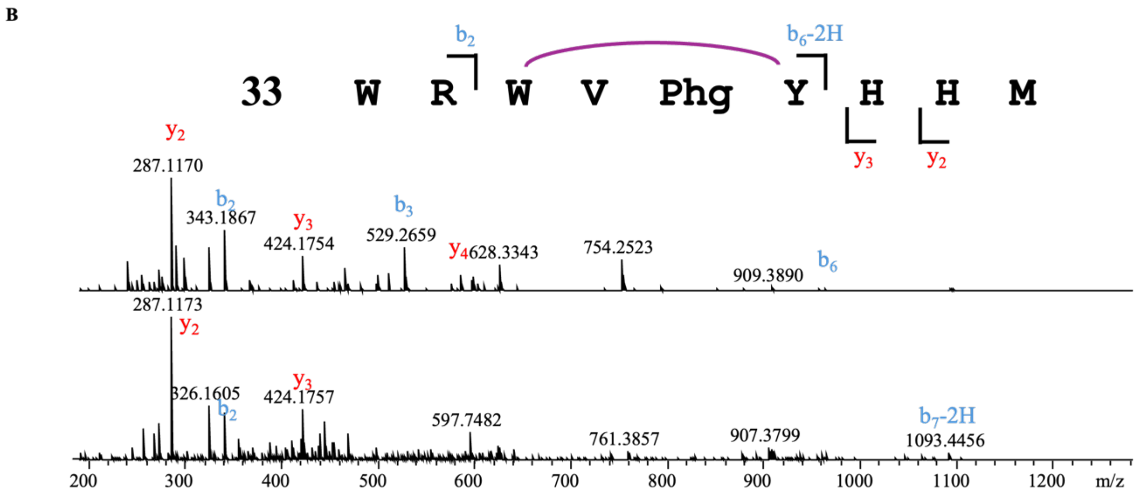


| Linear 35 | | | | | |
|---|---|---|---|---|---|
| Ions | Fragment | Observed mass | Calculated mass | z | Δppm |
| $y_2$ | HM | 287.1170 | 287.1172 | 1 | 0.7 |
| $y_3$ | HHM | 424.1754 | 424.1761 | 1 | 1.65 |
| $y_4$ | YHHM | 587.2383 | 587.2395 | 1 | 2.04 |
| $b_2$ | WR | 343.1867 | 343.1877 | 1 | 2.91 |
| $b_3$ | WRW | 529.2659 | 529.267 | 1 | 2.08 |
| $b_6$ | WRWV-L-Phg-Y | 958.4112 | 958.4125 | 1 | 1.36 |

| 35 | | | | | |
|---|---|---|---|---|---|
| Ions | Fragment | Observed mass | Calculated mass | z | Δppm |
| $y_2$ | HM | 287.1173 | 287.1172 | 1 | 0.35 |
| $y_3$ | HHM | 424.1757 | 424.1761 | 1 | 0.94 |
| $b_2$ | WR | 343.1858 | 343.1877 | 1 | 5.54 |
| $b_7$-2H | WRWV-L-Phg-YH | 1093.4456 | 1093.4558 | 1 | 9.33 |

**Figure S48.** GpeC enzymatic assay with WRWV-Chg-Y-HHM (**34**). A 5-hour reaction was performed before quenching by adding an equivalent volume of methanol. Tandem mass analysis revealed the peptide ions of $y_3$, $b_2$, and $b_6$-2H, proving the modification happened within the WV-Chg-Y motif.

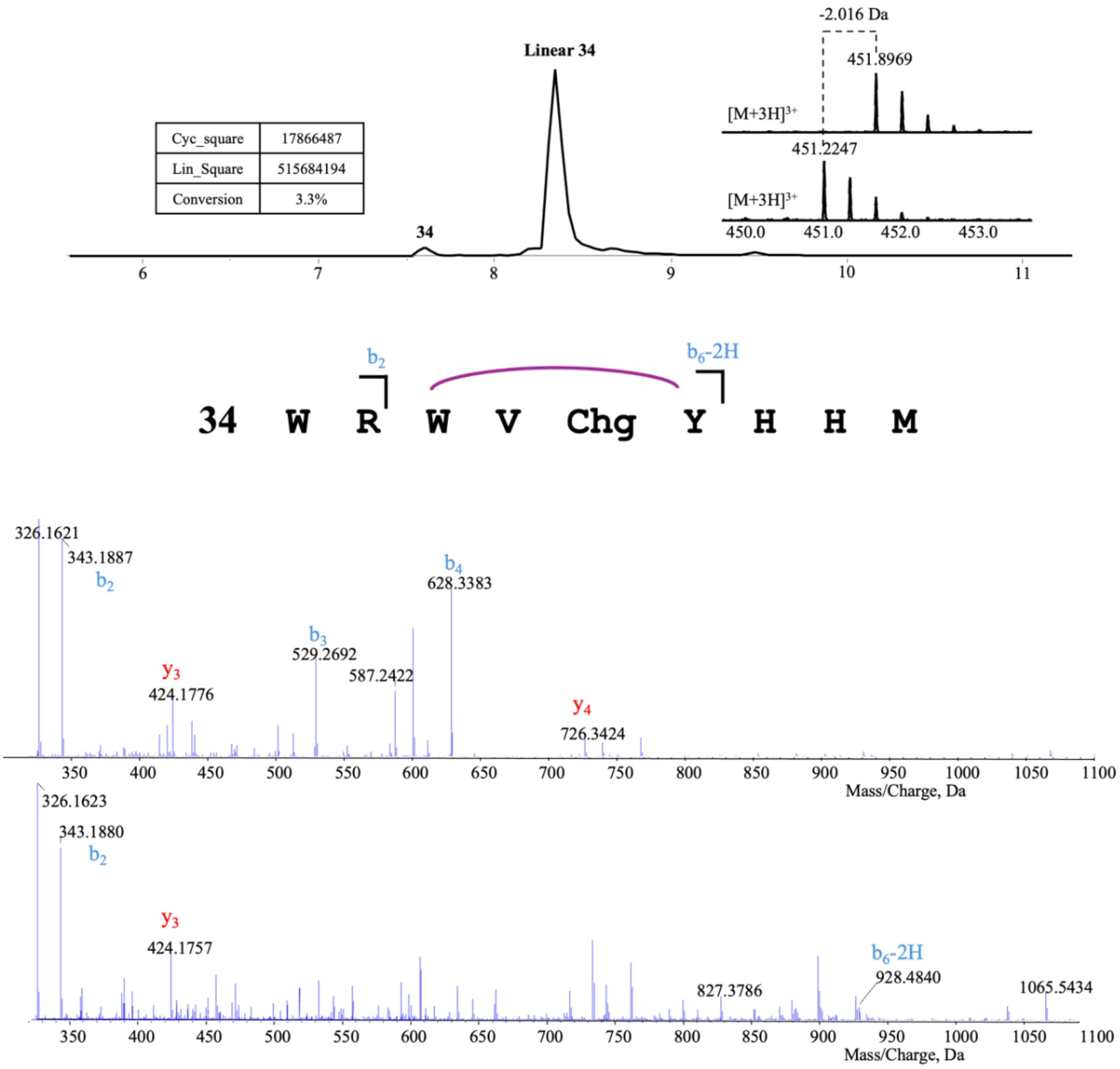


**Linear 34**

| Ions | Fragment | Observed mass | Calculated mass | z | Δppm |
|---|---|---|---|---|---|
| $y_3$ | HHM | 424.1776 | 424.1761 | 1 | 3.54 |
| $y_4$ | Chg-YHHM | 726.3424 | 726.3392 | 1 | 4.41 |
| $b_2$ | WR | 343.1887 | 343.1877 | 1 | 2.91 |
| $b_3$ | WRW | 529.2692 | 529.267 | 1 | 4.16 |
| $b_6$ | WRWV | 628.3383 | 628.3354 | 1 | 4.62 |

**34**

| Ions | Fragment | Observed mass | Calculated mass | z | Δppm |
|---|---|---|---|---|---|
| $y_3$ | HHM | 424.1766 | 424.1761 | 1 | 1.18 |
| $b_2$ | WR | 343.1875 | 343.1877 | 1 | 0.58 |
| $b_6$-2H | WRWV-Chg-Y | 928.4840 | 928.4828 | 1 | 1.29 |

**Figure S49.** GpeC enzymatic assay with WRWV-3-Bta-Y-HHM (**35**). A 5-hour reaction was performed before quenching by adding an equivalent volume of methanol. Tandem mass analysis revealed the peptide ions of $y_3$, $b_2$, and $b_6$-2H, proving the modification happened within the WV-3Bta-Y motif.

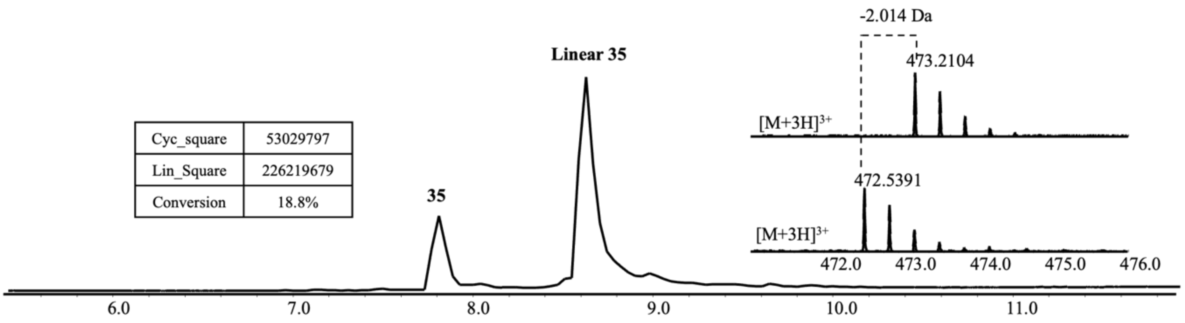


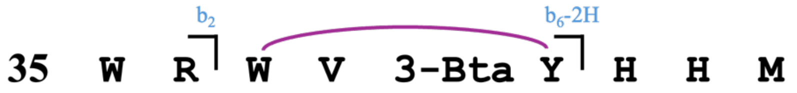


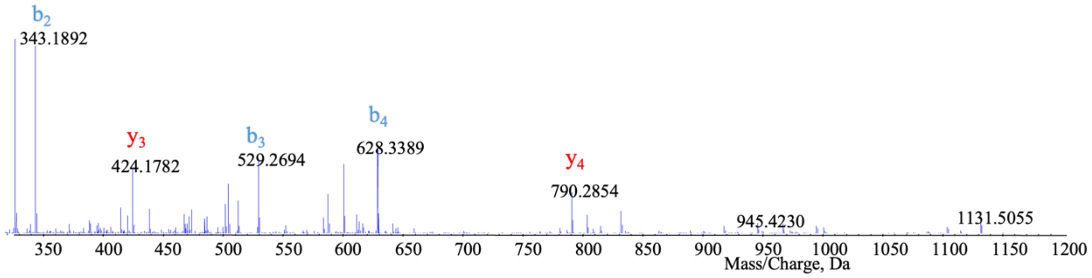


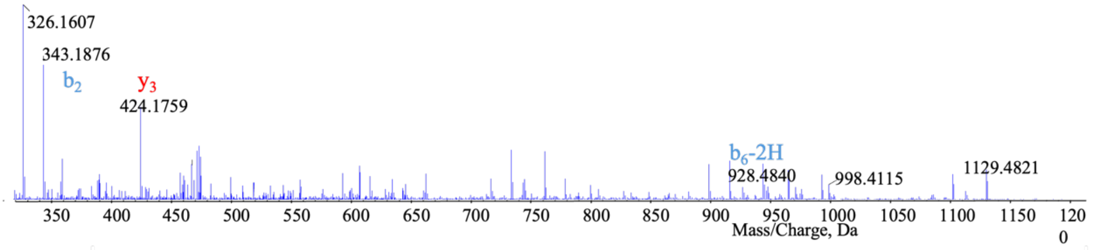


**Linear 35**

| Ions | Fragment | Observed mass | Calculated mass | z | Δppm |
|---|---|---|---|---|---|
| $y_3$ | HHM | 424.1782 | 424.1761 | 1 | 4.95 |
| $y_4$ | 3-Bta-YHHM | 790.2854 | 790.2800 | 1 | 6.83 |
| $b_2$ | WR | 343.1892 | 343.1877 | 1 | 4.37 |
| $b_3$ | WRW | 529.2694 | 529.267 | 1 | 4.53 |
| $b_4$ | WRWV | 628.3389 | 628.3354 | 1 | 5.57 |

**35**

| Ions | Fragment | Observed mass | Calculated mass | z | Δppm |
|---|---|---|---|---|---|
| $y_3$ | HHM | 424.1759 | 424.1761 | 1 | 0.47 |
| $b_2$ | WR | 343.1876 | 343.1877 | 1 | 0.29 |
| $b_6$-2H | WRWV-3Bta-Y | 928.4840 | 928.4828 | 1 | 1.29 |

**Figure S50.** GpeC enzymatic assay with WR-5F-Trp-VMYHHM (**36**). A 5-hour reaction was performed before quenching by adding an equivalent volume of methanol. One isomer was formed, marked as **36-1**. Tandem mass analysis revealed the peptide ions of $y_3$, $b_2$, and $b_6$-2H, proving the modification happened within the 5F-Trp-VMY motif.

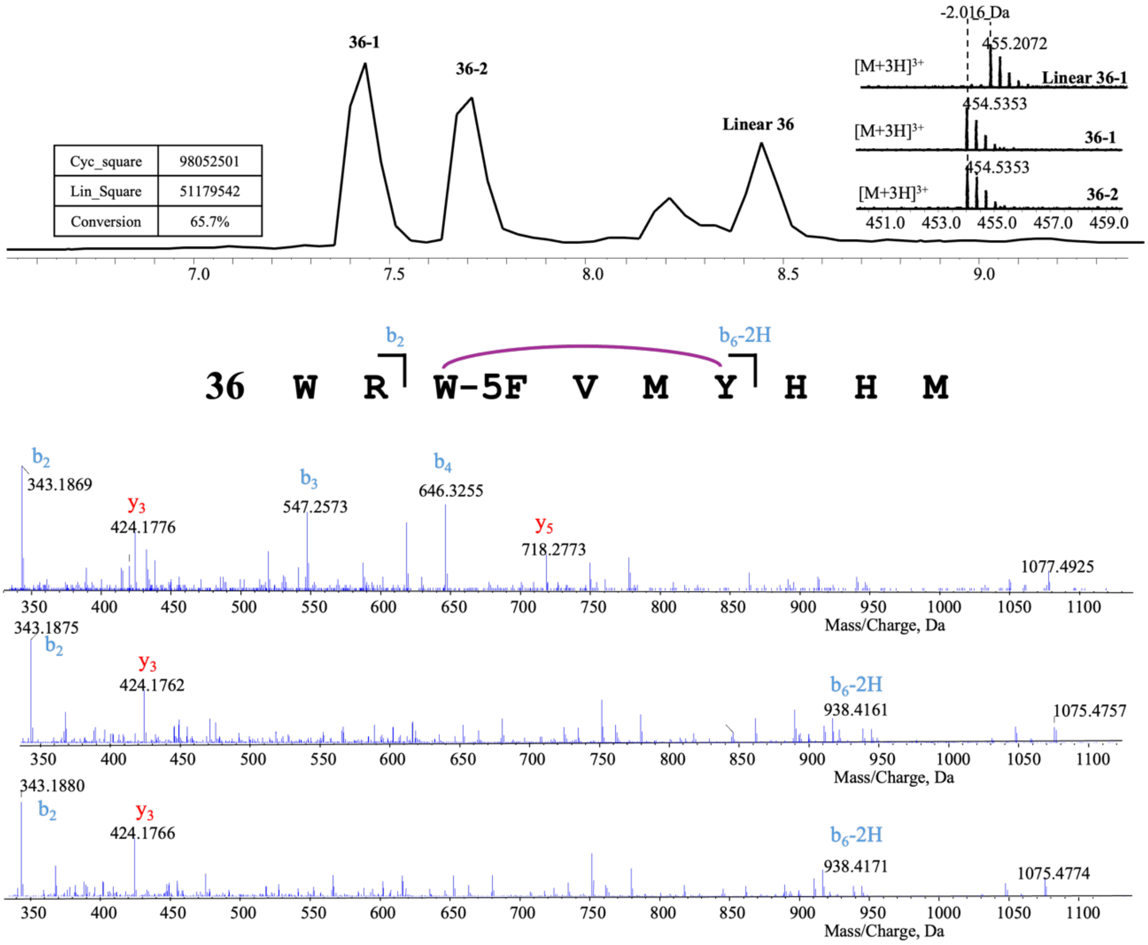


| Linear 36 | | | | | |
|---|---|---|---|---|---|
| Ions | Fragment | Observed mass | Calculated mass | z | Δppm |
| $y_3$ | HHM | 424.1776 | 424.1761 | 1 | 3.54 |
| $y_5$ | MYHHM | 718.2773 | 718.2800 | 1 | 3.76 |
| $b_2$ | WR | 343.1869 | 343.1877 | 1 | 2.33 |
| $b_3$ | WR-5F-Trp | 547.2573 | 547.2576 | 1 | 0.55 |
| $b_4$ | WR-5F-Trp-V | 646.3255 | 646.3260 | 1 | 0.77 |

| 36-1 | | | | | |
|---|---|---|---|---|---|
| Ions | Fragment | Observed mass | Calculated mass | z | Δppm |
| $y_3$ | HHM | 424.1762 | 424.1761 | 1 | 0.24 |
| $b_2$ | WR | 343.1875 | 343.1877 | 1 | 0.58 |
| $b_6$-2H | WR-5F-Trp-VMY | 938.4161 | 938.4142 | 1 | 2.02 |

| 36-2 | | | | | |
|---|---|---|---|---|---|
| Ions | Fragment | Observed mass | Calculated mass | z | Δppm |
| $y_3$ | HHM | 424.1766 | 424.1761 | 1 | 1.18 |
| $b_2$ | WR | 343.1880 | 343.1877 | 1 | 0.87 |
| $b_6$-2H | WR-5F-Trp-VMY | 938.4171 | 928.4828 | 1 | 3.09 |

**Figure S51.** GpeC enzymatic assay with WR-5Br-Trp-VMYHHM (**37**). A 5-hour reaction was performed before quenching by adding an equivalent volume of methanol. Trace amount of cyclic product was found, marked as **37**. Tandem mass analysis revealed the peptide ions of $y_3$, $b_2$, and $b_6$-2H, proving the modification happened within the 5Br-Trp-VMY motif.

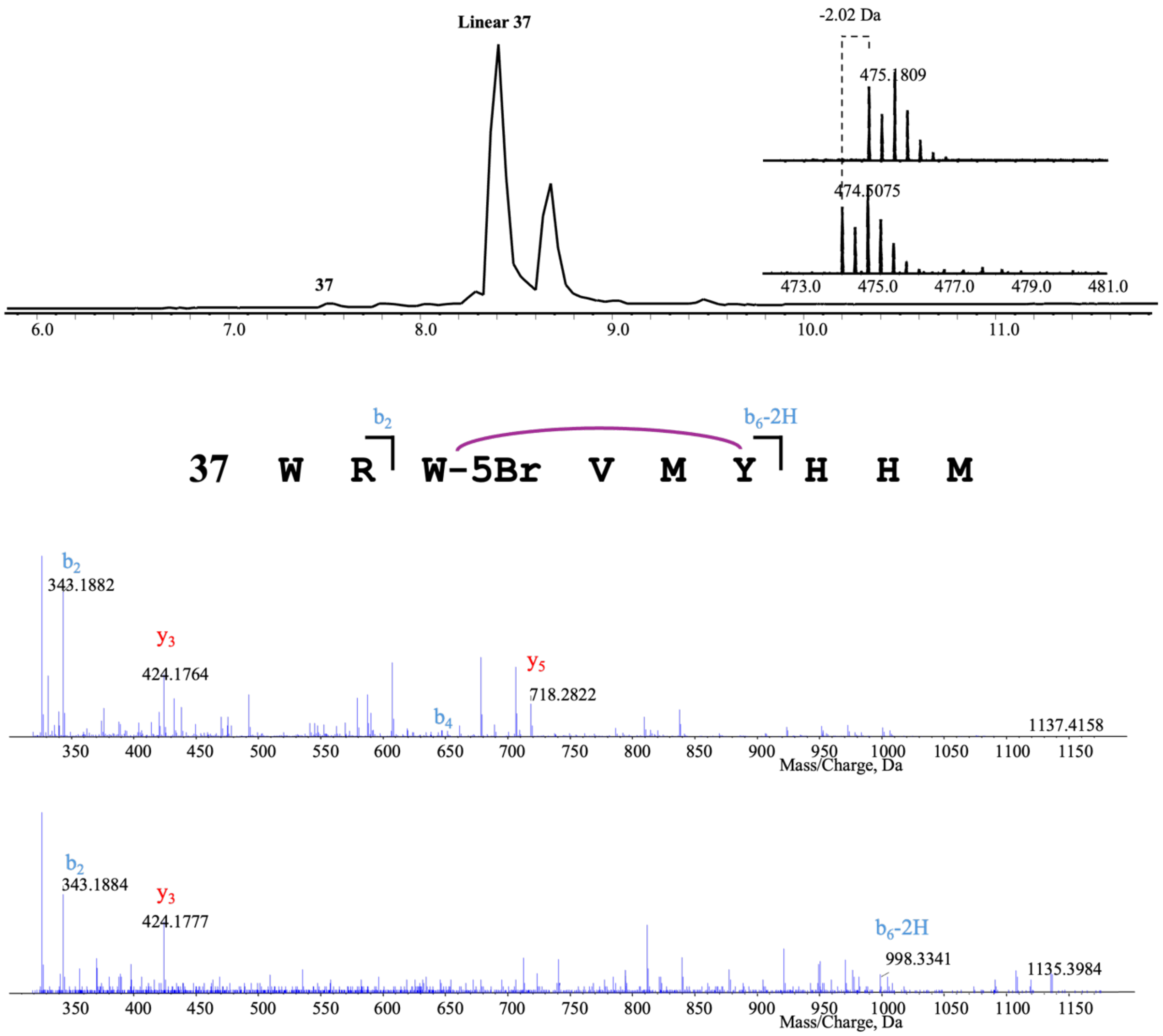


| **Linear 37** | | | | | |
|---|---|---|---|---|---|
| Ions | Fragment | Observed mass | Calculated mass | z | Δppm |
| $y_3$ | HHM | 424.1764 | 424.1761 | 1 | 0.71 |
| $y_5$ | MYHHM | 718.2822 | 718.2800 | 1 | 3.06 |
| $b_2$ | WR | 343.1882 | 343.1877 | 1 | 1.46 |
| $b_4$ | WR-5Br-Trp-V | 646.3255 | 646.3260 | 1 | 0.77 |
| **37-1** | | | | | |
| Ions | Fragment | Observed mass | Calculated mass | z | Δppm |
| $y_3$ | HHM | 424.1777 | 424.1761 | 1 | 3.77 |
| $b_2$ | WR | 343.1884 | 343.1877 | 1 | 2.04 |
| $b_6$-2H | WR-5Br-TrpVMY | 998.3420 | 998.3341 | 1 | 7.91 |

**Figure S52.** GpeC enzymatic assay with WR-6Cl-Trp-VMYHHM (**38**). A 5-hour reaction was performed before quenching by adding an equivalent volume of methanol. Tandem mass analysis revealed the peptide ions of $y_9$-2H and $b_7$-2H, proving the modification happened within the 6Cl-Trp-VMY motif.

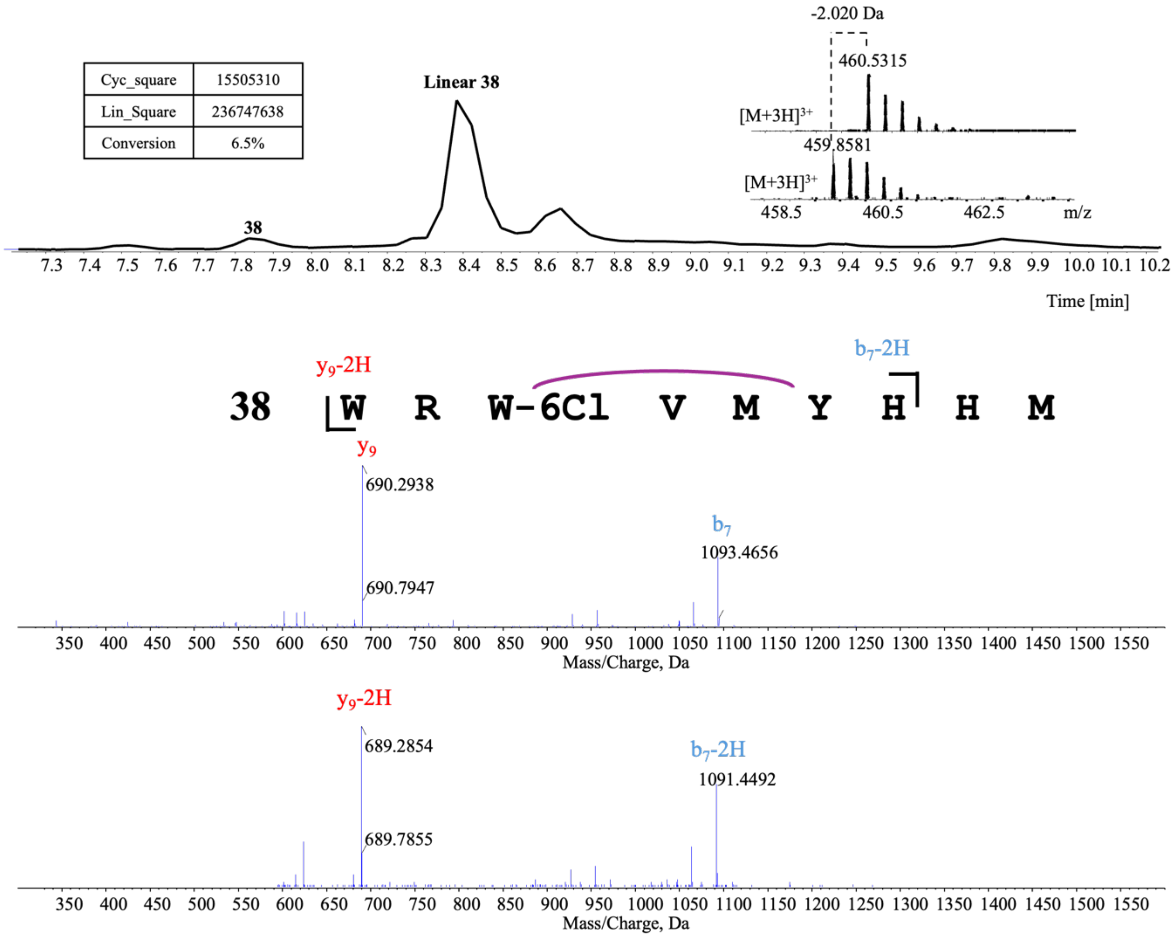


**Linear 38**

| Ions | Fragment | Observed mass | Calculated mass | z | Δppm |
|---|---|---|---|---|---|
| $y_9$ | WR-6Cl-Trp-VMYHHM | 690.2938 | 690.2882 | 2 | 0.71 |
| $b_7$ | WR-6Cl-Trp-VMYH | 1093.4656 | 1093.4592 | 1 | 0.77 |

**38**

| Ions | Fragment | Observed mass | Calculated mass | z | Δppm |
|---|---|---|---|---|---|
| $y_9$-2H | WR-6Cl-Trp-VMYHHM | 689.2854 | 689.2804 | 2 | 0.71 |
| $b_7$-2H | WR-6Cl-Trp-VMYH | 1091.4492 | 1091.4435 | 1 | 7.91 |

**Figure S53.** GpeC enzymatic assay with WRWVM-(3NO$_2$)Y-HHM (**39**). A 5-hour reaction was performed before quenching by adding an equivalent volume of methanol. Tandem mass analysis revealed the peptide ions of $y_3$, $b_2$, and $b_6$-2H, proving the modification happened within the WVM-(3NO$_2$)Ymotif.

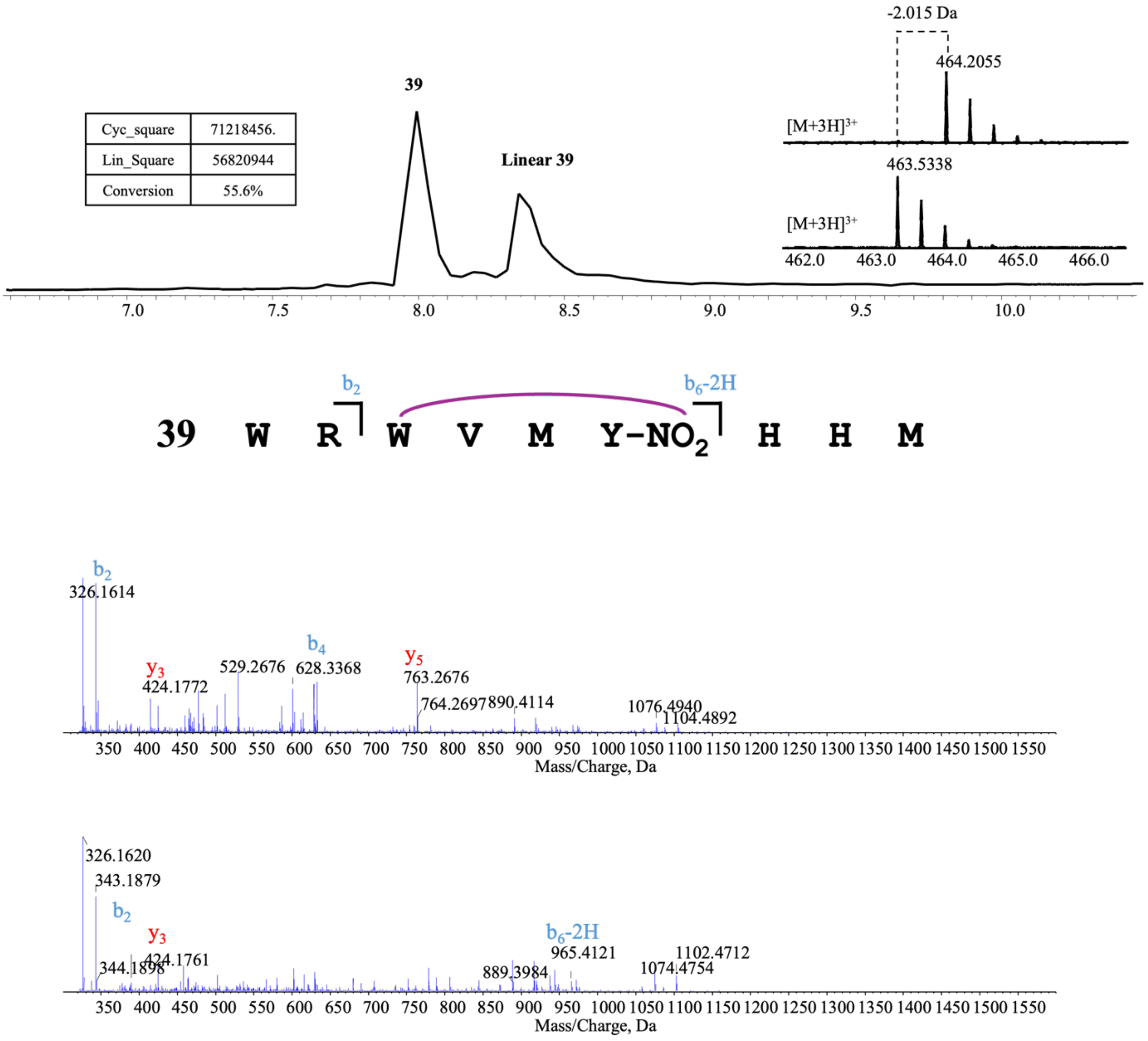


**Linear 39**

| Ions | Fragment | Observed mass | Calculated mass | z | Δppm |
|---|---|---|---|---|---|
| $y_3$ | HHM | 424.1772 | 424.1761 | 1 | 2.59 |
| $y_4$ | M-(3-NO$_2$)Y-HHM | 763.2676 | 763.2650 | 1 | 3.41 |
| $b_2$ | WR | 343.1874 | 343.1877 | 1 | 0.87 |
| $b_4$ | WRWV | 628.3368 | 628.3354 | 1 | 2.23 |

**39**

| Ions | Fragment | Observed mass | Calculated mass | z | Δppm |
|---|---|---|---|---|---|
| $y_3$ | HHM | 424.1761 | 424.1761 | 1 | <0.1 |
| $b_2$ | WR | 343.1879 | 343.1877 | 1 | 0.58 |
| $b_6$-2H | WRWVM-(3-NO$_2$)Y | 965.4121 | 965.4087 | 1 | 3.52 |

**Figure S54.** GpeC enzymatic assay with WRWVM-(3,5-DiI)Y-HHM (**40**). A 5-hour reaction was performed before quenching by adding an equivalent volume of methanol. Tandem mass analysis revealed the peptide ions of $y_3$, $y_5$, $b_2$, and $b_4$, while cyclic product displayed ions fragment of $y_3$, $b_2$, and $b_6$-2H-1I, proving the modification happened within the WVM-(3,5-DiI)Y motif wih a -2H and a indine atom loss.

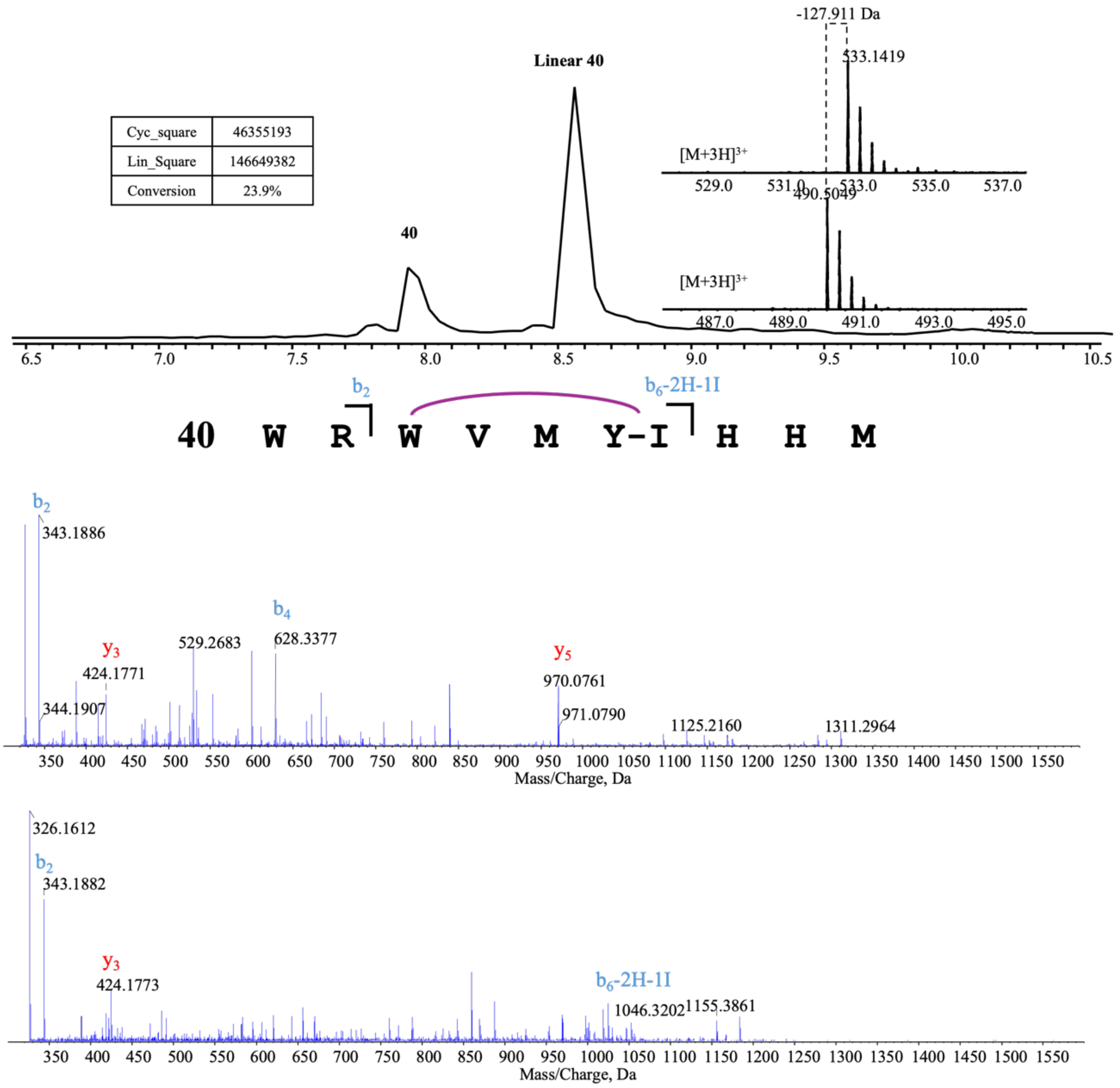


**Linear 40**

| Ions | Fragment | Observed mass | Calculated mass | z | Δppm |
|---|---|---|---|---|---|
| $y_3$ | HHM | 424.1771 | 424.1761 | 1 | 2.36 |
| $y_5$ | M-(3,5-DiI)Y-HHM | 970.0761 | 970.0732 | 1 | 2.99 |
| $b_2$ | WR | 343.1886 | 343.1877 | 1 | 2.62 |
| $b_4$ | WRWV | 628.3377 | 628.3354 | 1 | 3.66 |

**40**

| Ions | Fragment | Observed mass | Calculated mass | z | Δppm |
|---|---|---|---|---|---|
| $y_3$ | HHM | 424.1773 | 424.1761 | 1 | 2.83 |
| $b_2$ | WR | 343.1882 | 343.1877 | 1 | 1.46 |
| $b_6$-2H-1I | WRWVM(Y-I) | 1046.3241 | 1046.3202 | 1 | 3.73 |

**Figure S55.** GpeC enzymatic assay with WRWVM-(m-OH)Y-HHM (**41**). A 5-hour reaction was performed before quenching by adding an equivalent volume of methanol. Tandem mass analysis revealed the peptide ions $y_9$-2H and $b_7$-2H two cyclic products with different retention times marked as **41-1** and **41-2.**

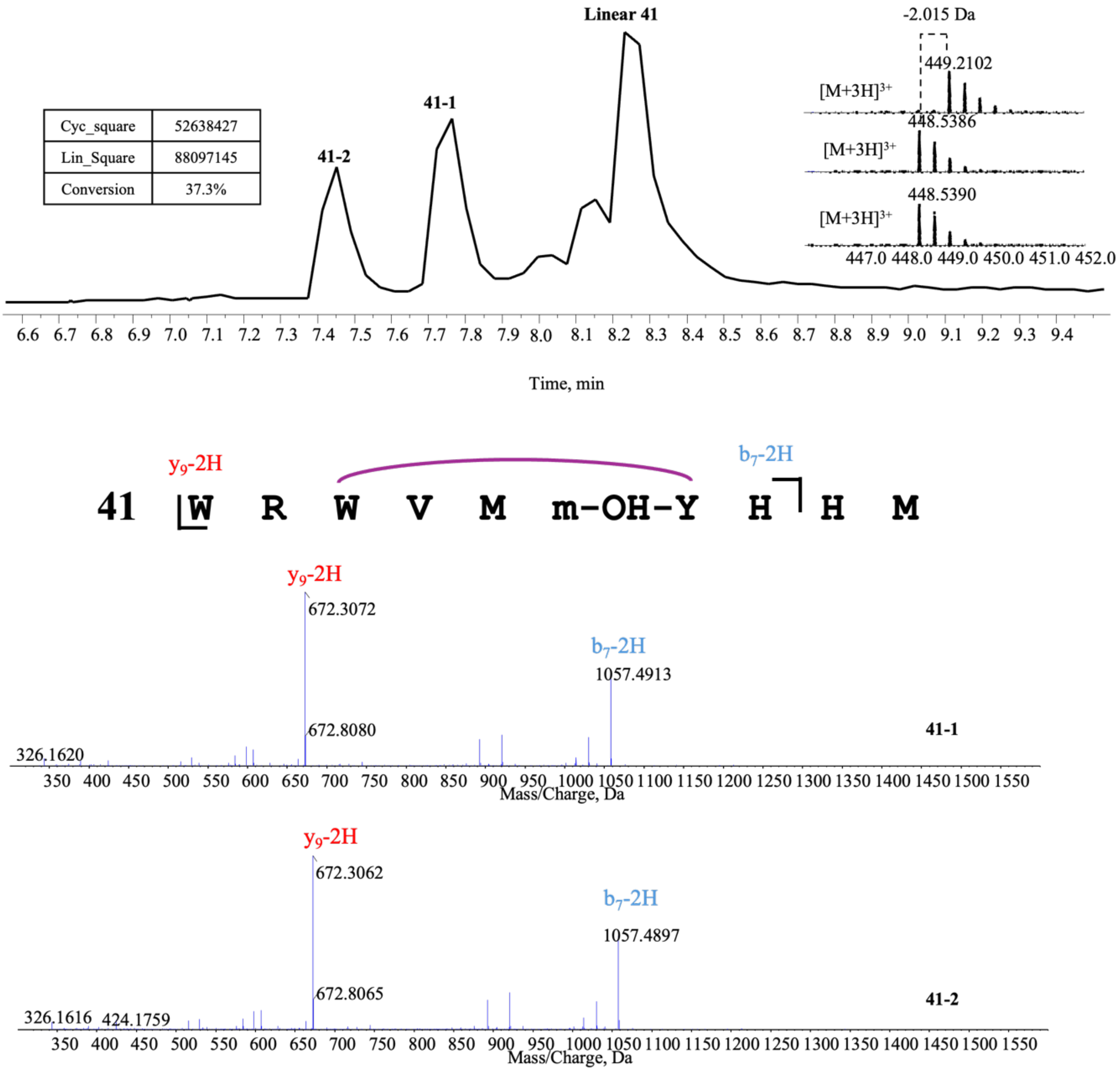


**41-1**

| Ions | Fragment | Observed mass | Calculated mass | z | Δppm |
|---|---|---|---|---|---|
| $y_9$-2H | WRWVM-(m-OH)Y-HHM | 672.3072 | 672.2998 | 2 | 11 |
| $b_7$-2H | WRWVM-(m-OH)Y-H | 1057.4913 | 1057.4825 | 1 | 8.32 |

**41-2**

| Ions | Fragment | Observed mass | Calculated mass | z | Δppm |
|---|---|---|---|---|---|
| $y_9$-2H | WRWVM-(m-OH)Y-HHM | 672.3072 | 672.2998 | 2 | 11 |
| $b_7$-2H | WRWVM-(m-OH)Y-H | 1057.4897 | 1057.4825 | 1 | 6.81 |

**Figure S56.** The method was established for building the tetrapeptide module (LTT). A) Chemical synthesis of an ether-substituted substrate and UPLC-HRMS verification of the synthesized substrate (**43**), the key ether bond was confirmed with $y_3$ ion on tandem mass (Obs. 425.1610, Exa. 425.16). C) UPLC-HRMS verification of the catalyzed product under the Fdx, Fdr, and NADPH system. (**44**) represents the cyclized product. We also observed a Trp-Tyr cyclized product WRWVMY (**44'**) and a linear WRWVMY (**43'**), and the lytic HHM. D) HRMS verification of the product after being treated with trypsin and NaOH, resulting in the final tetracyclic core part (**45**).

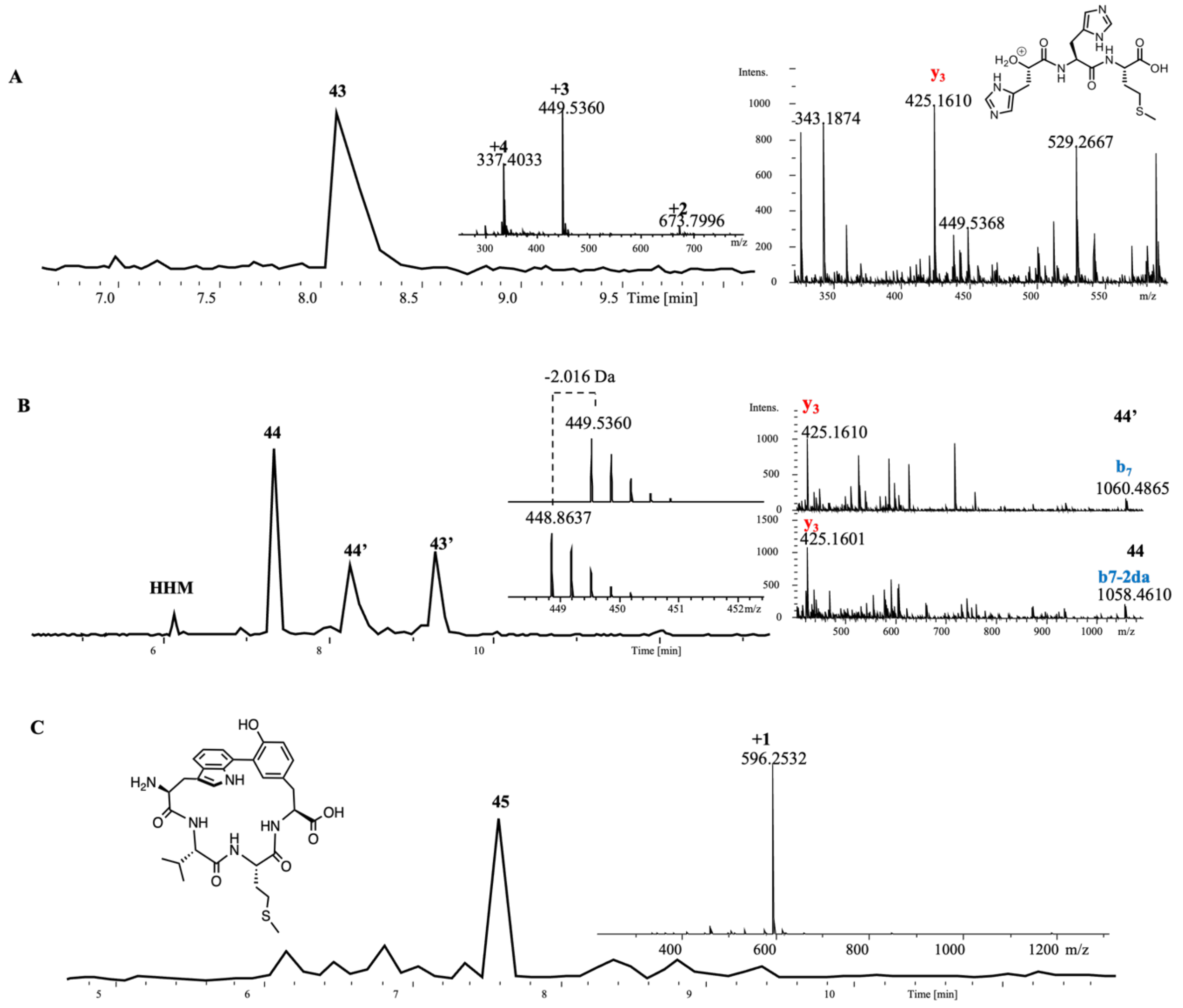

#### UPLC- HRMS data

##### Truncated leader peptide assay

GpeC enzymatic assay with Shortened-1 (RNQAVCYSKNPSWFWMLYHHM). A 5-hour reaction was performed before quenching by adding an equivalent volume of methanol. Tandem mass analysis revealed the peptide ions of $y_2$, $y_3$, $y_7$-2H, and $y_8$-2H, proving the modification happened within the WMLY motif.

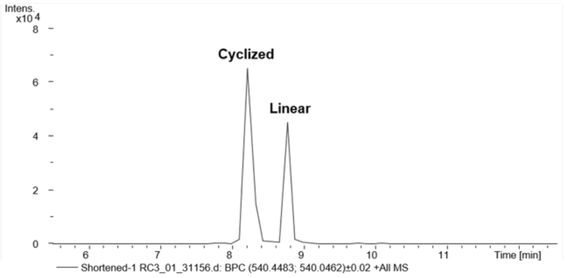


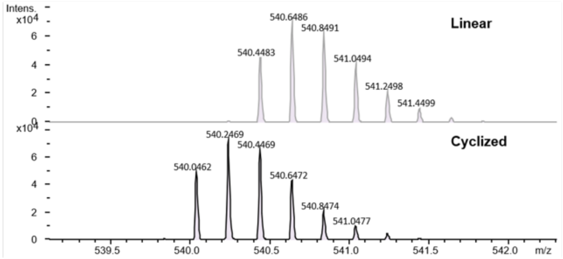


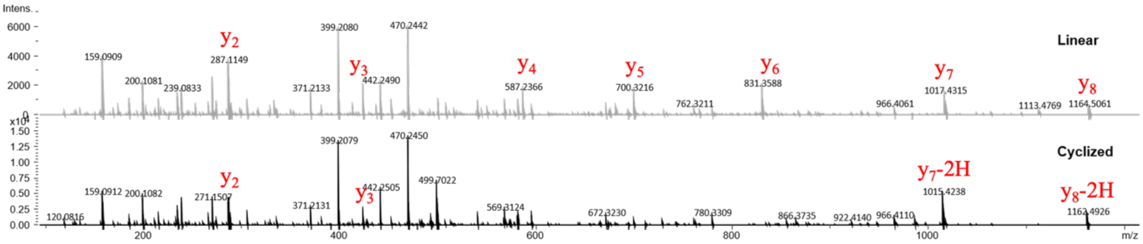


**Linear shortened-1**

| ions | Fragment | Observed mass | Calculated mass | Z | Δppm |
|---|---|---|---|---|---|
| $y_2$ | HM | 287.1149 | 287.1172 | 1 | 8.01 |
| $y_3$ | HHM | 424.1758 | 424.1761 | 1 | 0.7 |
| $y_4$ | YHHM | 587.2366 | 587.2395 | 1 | 4.93 |
| $y_5$ | LYHHM | 700.3216 | 700.3235 | 1 | 2.71 |
| $y_6$ | MLYHHM | 831.3588 | 831.364 | 1 | 6.25 |
| $y_7$ | WMLYHHM | 1017.4315 | 1017.443 | 1 | 11.3 |
| $y_8$ | FWMLYHHM | 1164.5061 | 1164.512 | 1 | 5.06 |

**Cyclized shrtened-1**

| ions | Fragment | Observed mass | Calculated mass | Z | Δppm |
|---|---|---|---|---|---|
| $y_2$ | HM | 287.1163 | 287.1172 | 1 | 3.13 |
| $y_3$ | HHM | 424.1768 | 424.1761 | 1 | 1.65 |
| $y_7$-2H | WMLYHHM | 1015.4238 | 1015.428 | 1 | 4.13 |
| $y_8$-2H | FWMLYHHM | 1162.4926 | 1162.496 | 1 | 2.92 |

GpeC enzymatic assay with Shortened-2 (AVCYSKNPSWFWMLYHHM). A 5-hour reaction was performed before quenching by adding an equivalent volume of methanol. Tandem mass analysis revealed the peptide ions of $y_2$, $y_3$, $y_7$-2H, and $y_8$-2H, proving the modification happened within the WMLY motif.

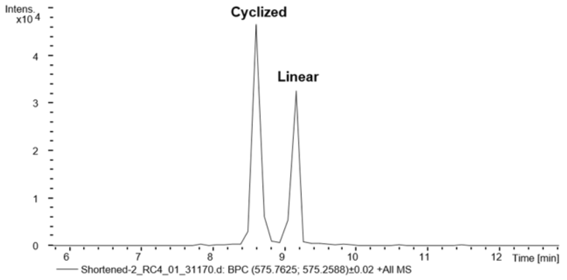


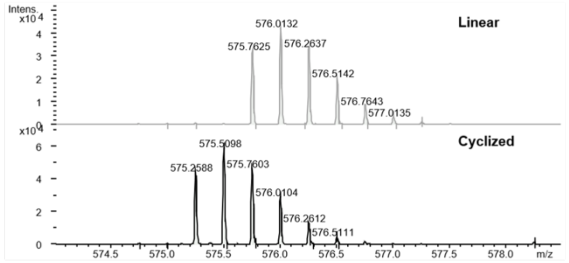


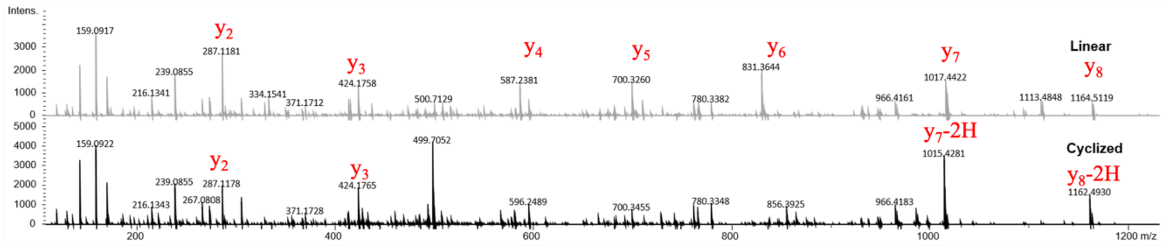


**Linear shortened-2**

| ions | Fragment | Observed mass | Calculated mass | Z | Δppm |
|---|---|---|---|---|---|
| $y_2$ | HM | 287.1181 | 287.1172 | 1 | 3.13 |
| $y_3$ | HHM | 424.1758 | 424.1761 | 1 | 0.7 |
| $y_4$ | YHHM | 587.2381 | 587.2395 | 1 | 2.38 |
| $y_5$ | LYHHM | 700.3260 | 700.3235 | 1 | 3.56 |
| $y_6$ | MLYHHM | 831.3644 | 831.364 | 1 | 0.48 |
| $y_7$ | WMLYHHM | 1017.4422 | 1017.443 | 1 | 0.78 |
| $y_8$ | FWMLYHHM | 1164.5119 | 1164.512 | 1 | 0.08 |

**Cyclized shortened-2**

| ions | Fragment | Observed mass | Calculated mass | Z | Δppm |
|---|---|---|---|---|---|
| $y_2$ | HM | 287.1178 | 287.1172 | 1 | 2.08 |
| $y_3$ | HHM | 424.1765 | 424.1761 | 1 | 0.94 |
| $y_7$-2H | WMLYHHM | 1015.4281 | 1015.428 | 1 | 0.09 |
| $y_8$-2H | FWMLYHHM | 1162.4930 | 1162.496 | 1 | 2.58 |

GpeC enzymatic assay with Shortened-3 (YSKNPSWFWMLYHHM). A 5-hour reaction was performed before quenching by adding an equivalent volume of methanol. Tandem mass analysis revealed the peptide ions of $y_2$, $y_3$, $y_7$-2H, and $y_8$-2H, proving the modification happened within the WMLY motif.

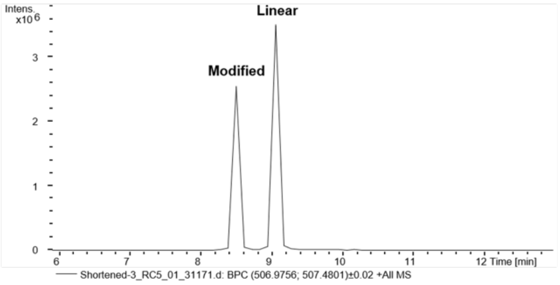


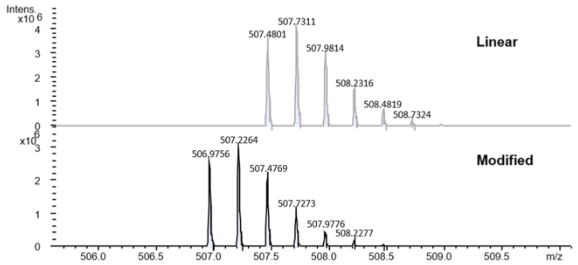


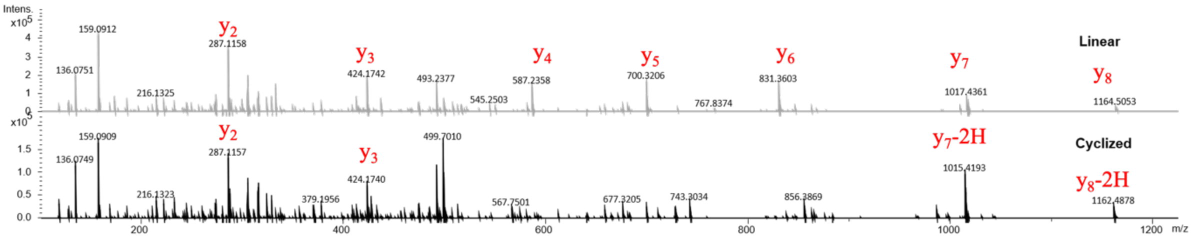


**Linear shortened-3**

| ions | Fragment | Observed mass | Calculated mass | Z | Δppm |
|---|---|---|---|---|---|
| $y_2$ | HM | 287.1158 | 287.1172 | 1 | 4.87 |
| $y_3$ | HHM | 424.1742 | 424.1761 | 1 | 4.47 |
| $y_4$ | YHHM | 587.2358 | 587.2395 | 1 | 6.30 |
| $y_5$ | LYHHM | 700.3206 | 700.3235 | 1 | 4.14 |
| $y_6$ | MLYHHM | 831.3603 | 831.364 | 1 | 4.45 |
| $y_7$ | WMLYHHM | 1017.4361 | 1017.443 | 1 | 6.78 |
| $y_8$ | FWMLYHHM | 1164.5053 | 1164.512 | 1 | 5.75 |

**Cyclized shortened-3**

| ions | Fragment | Observed mass | Calculated mass | Z | Δppm |
|---|---|---|---|---|---|
| $y_2$ | HM | 287.1157 | 287.1172 | 1 | 5.22 |
| $y_3$ | HHM | 424.1740 | 424.1761 | 1 | 4.95 |
| $y_7$-2H | WMLYHHM | 1015.4193 | 1015.428 | 1 | 8.56 |
| $y_8$-2H | FWMLYHHM | 1162.4878 | 1162.496 | 1 | 7.05 |

GpeC enzymatic assay with Shortened-4 (NPSWFWMLYHHM). A 5-hour reaction was performed before quenching by adding an equivalent volume of methanol. Tandem mass analysis revealed the peptide ions of $y_2$, $y_3$, $y_7$-2H, and $y_8$-2H, proving the modification happened within the WMLY motif.

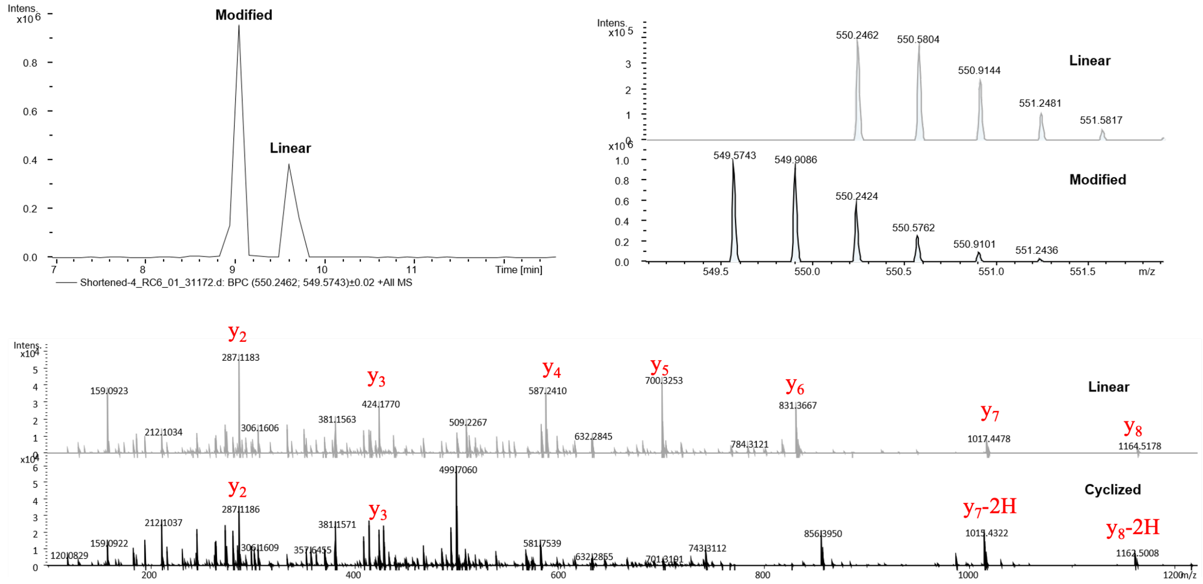


**Linear shortened-4**

| ions | Fragment | Observed mass | Calculated mass | Z | Δppm |
|---|---|---|---|---|---|
| $y_2$ | HM | 287.1183 | 287.1172 | 1 | 3.83 |
| $y_3$ | HHM | 424.1770 | 424.1761 | 1 | 2.12 |
| $y_4$ | YHHM | 587.2410 | 587.2395 | 1 | 2.55 |
| $y_5$ | LYHHM | 700.3253 | 700.3235 | 1 | 2.57 |
| $y_6$ | MLYHHM | 831.3667 | 831.364 | 1 | 3.24 |
| $y_7$ | WMLYHHM | 1017.4478 | 1017.443 | 1 | 4.71 |
| $y_8$ | FWMLYHHM | 1164.5178 | 1164.512 | 1 | 4.98 |

**Cyclized shortened-4**

| ions | Fragment | Observed mass | Calculated mass | Z | Δppm |
|---|---|---|---|---|---|
| $y_2$ | HM | 287.1186 | 287.1172 | 1 | 4.87 |
| $y_3$ | HHM | 424.1772 | 424.1761 | 1 | 2.59 |
| $y_7$-2H | WMLYHHM | 1015.4322 | 1015.428 | 1 | 4.13 |
| $y_8$-2H | FWMLYHHM | 1162.5008 | 1162.496 | 1 | 4.12 |

**Mutations on the ring-forming residues**

GpeC enzymatic assay with WRHVMYHHM. A 5-hour reaction was performed before quenching by adding an equivalent volume of methanol. No cyclized product was observed. Tandem mass analysis revealed the linear peptide ions of $y_2$, $y_3$, and $b_3$.

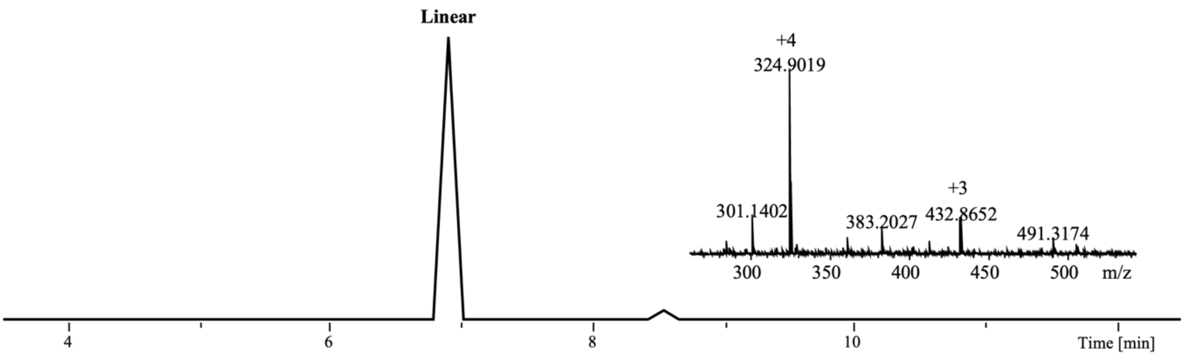


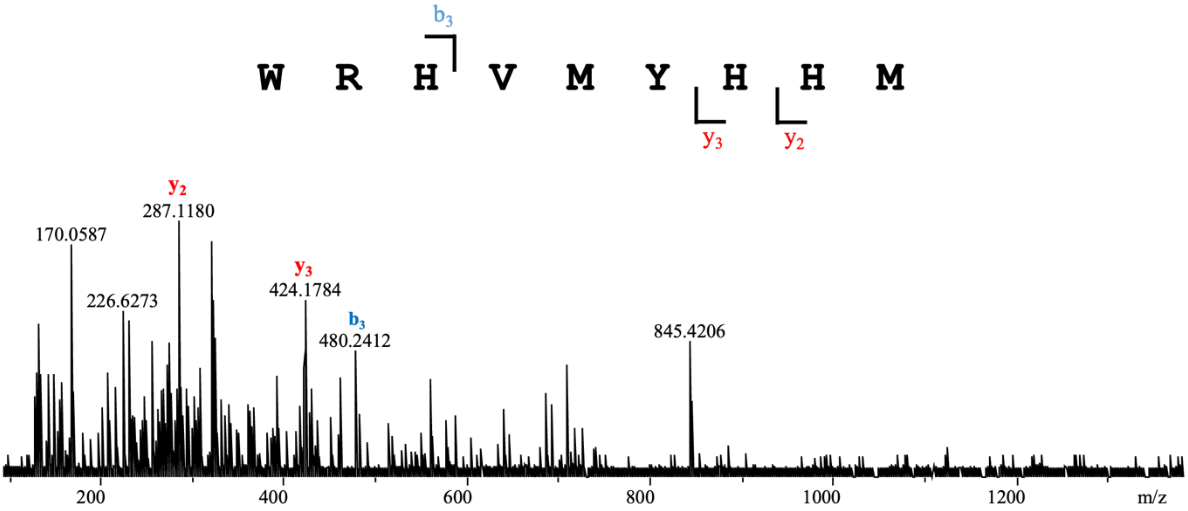


**Linear**

| Ions | Fragment | Observed mass | Calculated mass | z | Δppm |
|---|---|---|---|---|---|
| $y_2$ | HM | 287.1180 | 287.1172 | 1 | 2.79 |
| $y_3$ | HHM | 424.1784 | 424.1761 | 1 | 5.42 |
| $b_3$ | WRH | 480.2412 | 480.2466 | 1 | 11.2 |

GpeC enzymatic assay with WRHVMWHHM. A 5-hour reaction was performed before quenching by adding an equivalent volume of methanol. No cyclized product was observed. Tandem mass analysis revealed the linear peptide ions of $y_2$, $y_3$, $y_4$, $y_5$, $b_2$, and $b_6$.

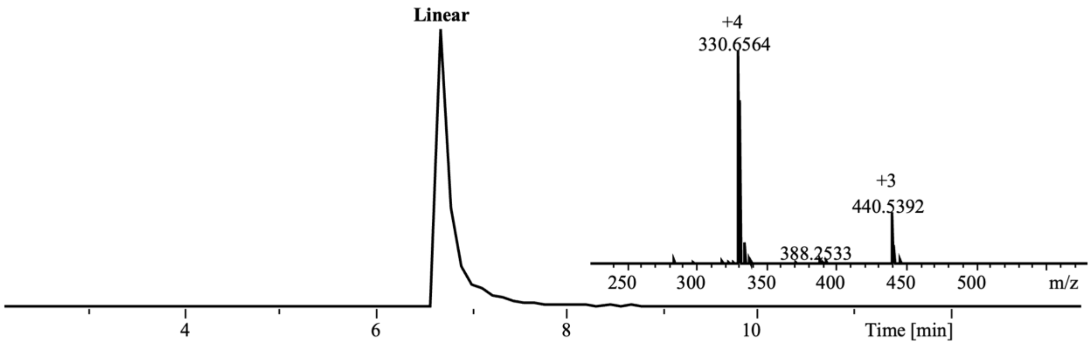


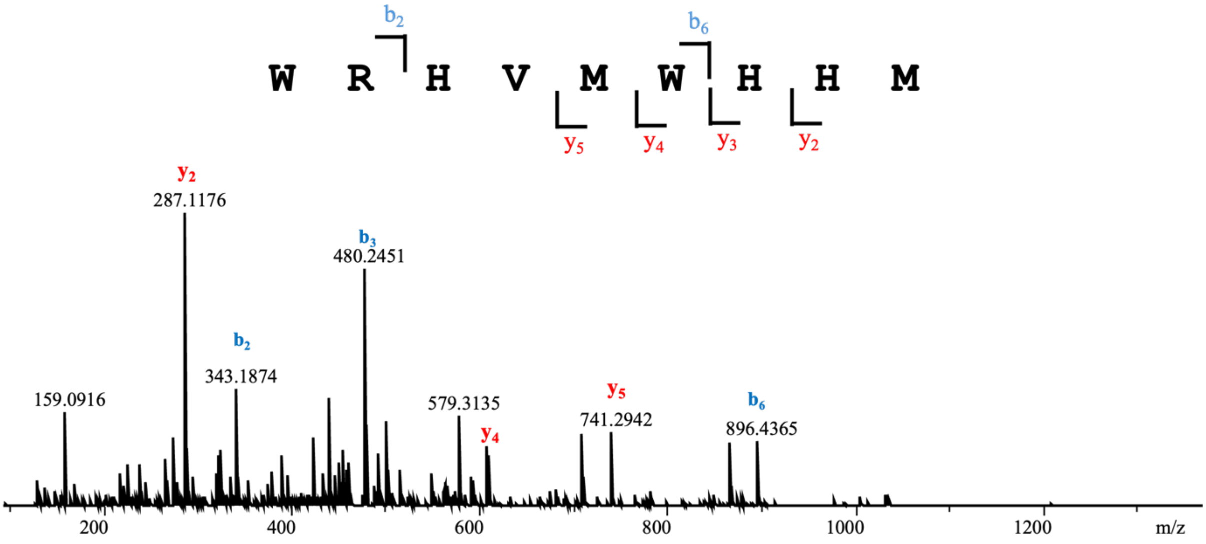


| Linear | | | | | |
|---|---|---|---|---|---|
| Ions | Fragment | Observed mass | Calculated mass | z | Δppm |
| $y_2$ | HM | 287.1176 | 287.1172 | 1 | 1.39 |
| $y_3$ | HHM | 424.1753 | 424.1761 | 1 | 1.89 |
| $y_4$ | WHHM | 610.2517 | 610.2555 | 1 | 6.23 |
| $y_5$ | MWHHM | 741.2942 | 741.2959 | 1 | 2.29 |
| $b_2$ | WR | 343.1874 | 343.1877 | 1 | 0.87 |
| $b_3$ | WRH | 480.2451 | 480.2466 | 1 | 3.12 |
| $b_6$ | WRHVMW | 896.4365 | 896.4348 | 1 | 1.9 |

GpeC enzymatic assay with WRFVMWHHM. A 5-hour reaction was performed before quenching by adding an equivalent volume of methanol. No cyclized product was observed. Tandem mass analysis revealed the linear peptide ions of $y_2$, $y_3$, $y_4$, $y_5$, $b_2$, and $b_6$.

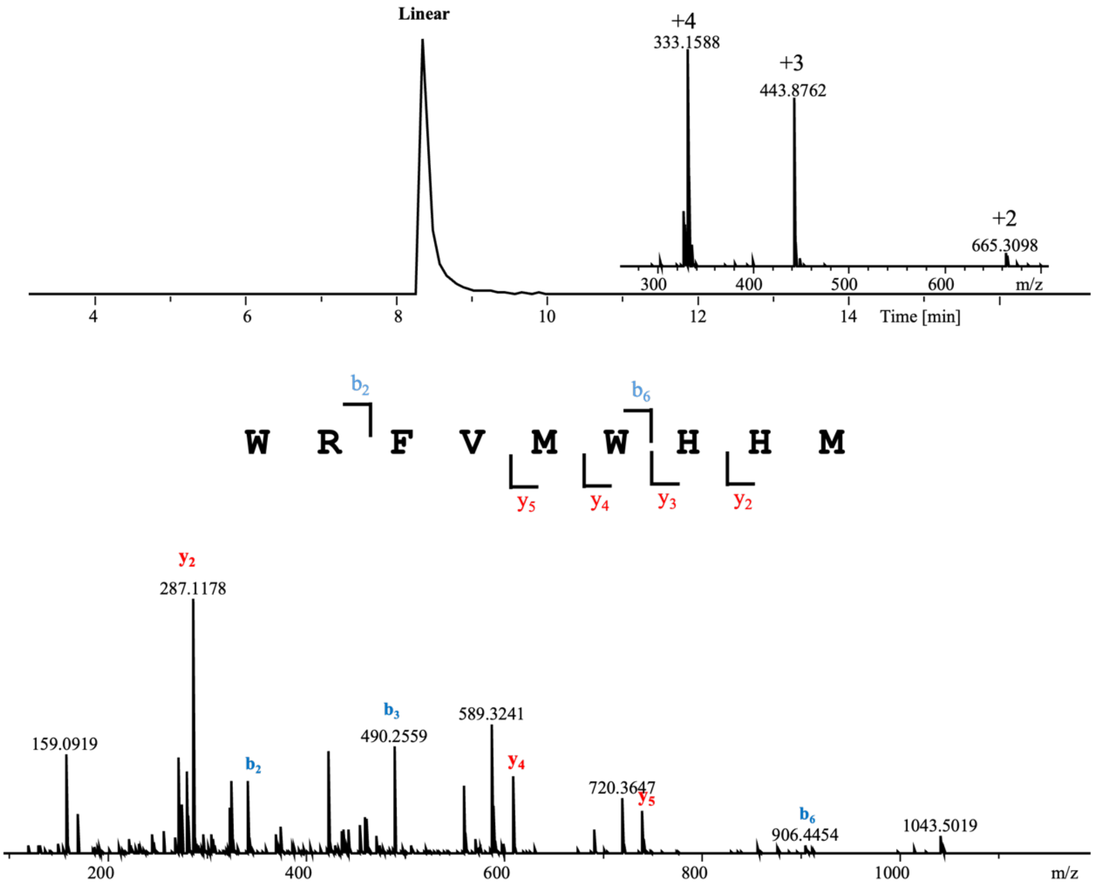


| Linear | | | | | |
|---|---|---|---|---|---|
| Ions | Fragment | Observed mass | Calculated mass | z | Δppm |
| $y_2$ | HM | 287.1178 | 287.1172 | 1 | 2.09 |
| $y_3$ | HHM | 424.1768 | 424.1761 | 1 | 1.65 |
| $y_4$ | WHHM | 610.2552 | 610.2555 | 1 | 0.49 |
| $y_5$ | MWHHM | 741.2954 | 741.2959 | 1 | 0.67 |
| $b_2$ | WR | 343.1877 | 343.1877 | 1 | <0.1 |
| $b_3$ | WRF | 490.2559 | 490.2561 | 1 | 0.41 |
| $b_6$ | WRFVMW | 906.4454 | 906.4443 | 1 | 1.21 |

GpeC enzymatic assay with WRYVMFHHM. A 5-hour reaction was performed before quenching by adding an equivalent volume of methanol. No cyclized product was observed. Tandem mass analysis revealed the linear peptide ions of $y_2$, $y_3$, $y_4$, $y_5$, $y_7$, $b_2$, $b_3$, and $b_6$.

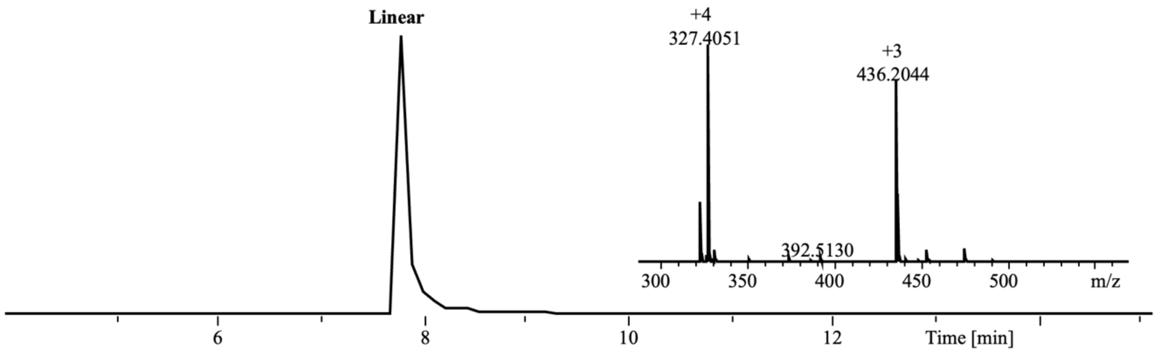


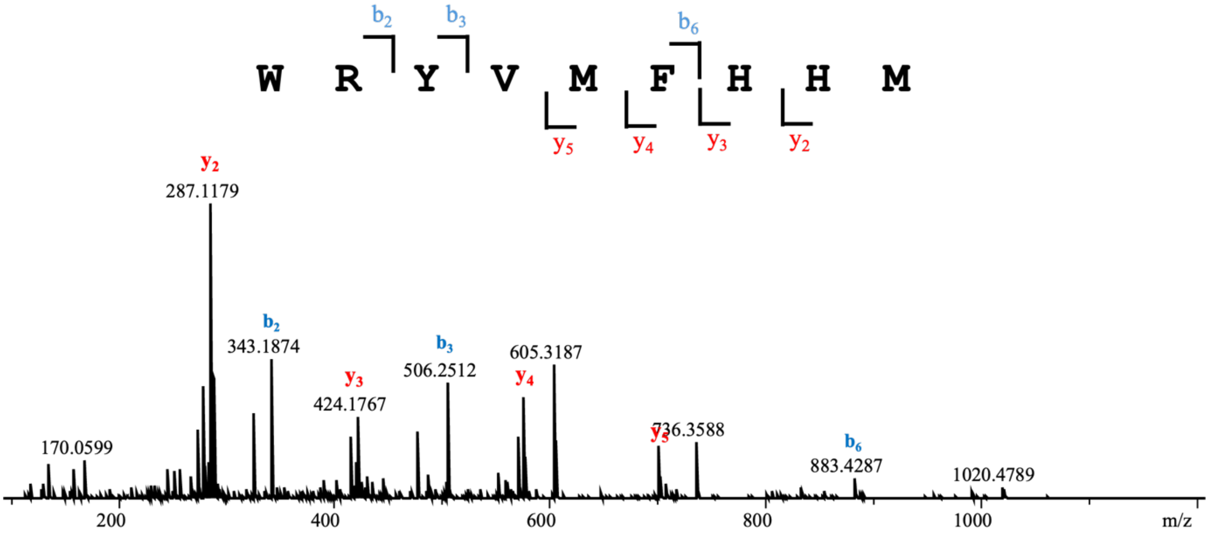


| Linear | | | | | |
|---|---|---|---|---|---|
| Ions | Fragment | Observed mass | Calculated mass | z | Δppm |
| $y_2$ | HM | 287.1179 | 287.1172 | 1 | 2.44 |
| $y_3$ | HHM | 424.1767 | 424.1761 | 1 | 1.41 |
| $y_4$ | FHHM | 571.2423 | 571.2446 | 1 | 4.03 |
| $y_5$ | MFHHM | 702.2836 | 702.285 | 1 | 1.99 |
| $b_2$ | WR | 343.1874 | 343.1877 | 1 | 0.87 |
| $b_3$ | WRY | 506.2512 | 506.251 | 1 | 0.4 |
| $b_6$ | WRYVMF | 883.4287 | 883.4283 | 1 | 0.45 |

GpeC enzymatic assay with WRHVMFHHM. A 5-hour reaction was performed before quenching by adding an equivalent volume of methanol. No cyclized product was observed. Tandem mass analysis revealed the linear peptide ions of $y_2$, $y_3$, $y_4$, $y_5$, $y_7$, $b_2$, $b_3$, and $b_6$.

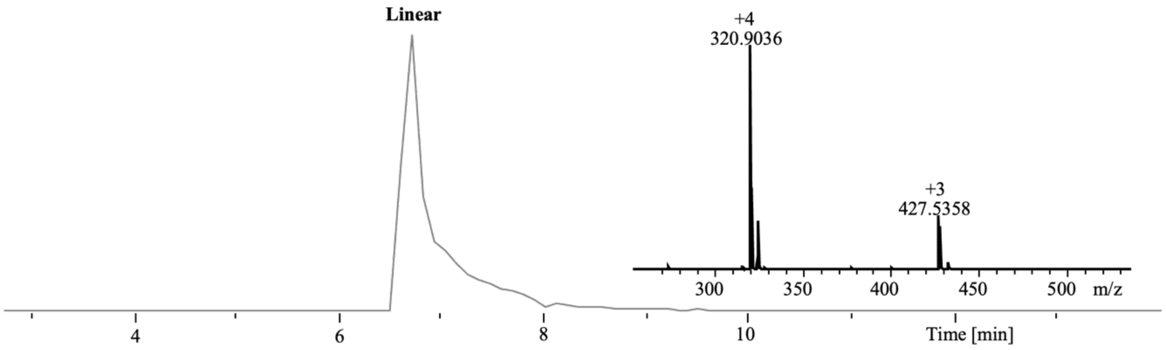


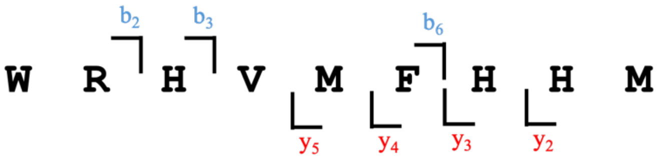


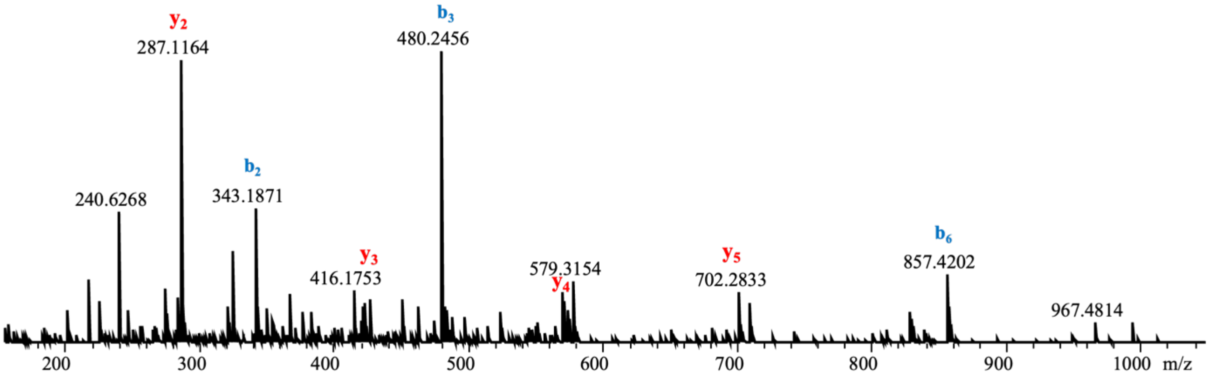


| Linear | | | | | |
|---|---|---|---|---|---|
| Ions | Fragment | Observed mass | Calculated mass | z | Δppm |
| $y_2$ | HM | 287.1164 | 287.1172 | 1 | 2.79 |
| $y_3$ | HHM | 424.1754 | 424.1761 | 1 | 1.65 |
| $y_4$ | FHHM | 571.2446 | 571.2446 | 1 | <0.1 |
| $y_5$ | MFHHM | 702.2833 | 702.285 | 1 | 2.42 |
| $b_2$ | WR | 343.1871 | 343.1877 | 1 | 1.75 |
| $b_3$ | WRH | 480.2456 | 480.2466 | 1 | 2.08 |
| $b_6$ | WRHVMF | 857.4202 | 857.4239 | 1 | 4.32 |

GpeC enzymatic assay with FRWVMYHHM. A 5-hour reaction was performed before quenching by adding an equivalent volume of methanol. No cyclized product was observed. Tandem mass analysis revealed the linear peptide ions of $y_2$, $y_3$, $y_4$, $y_5$, $y_7$, $b_2$, and $b_3$.

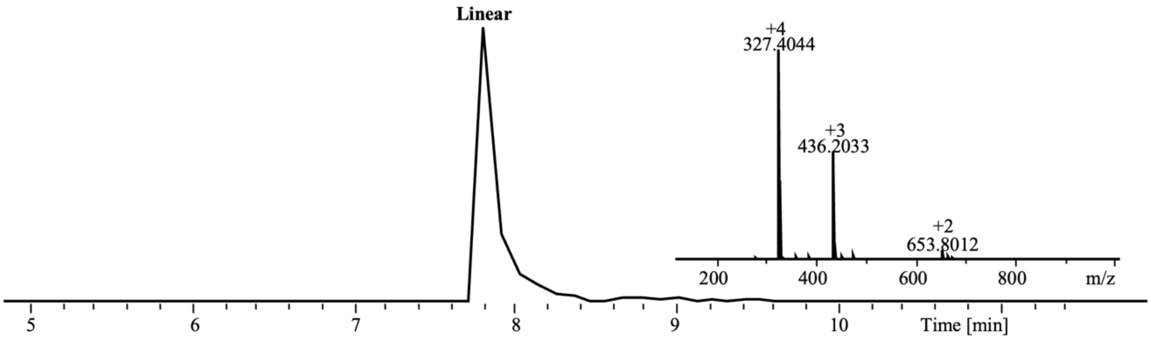


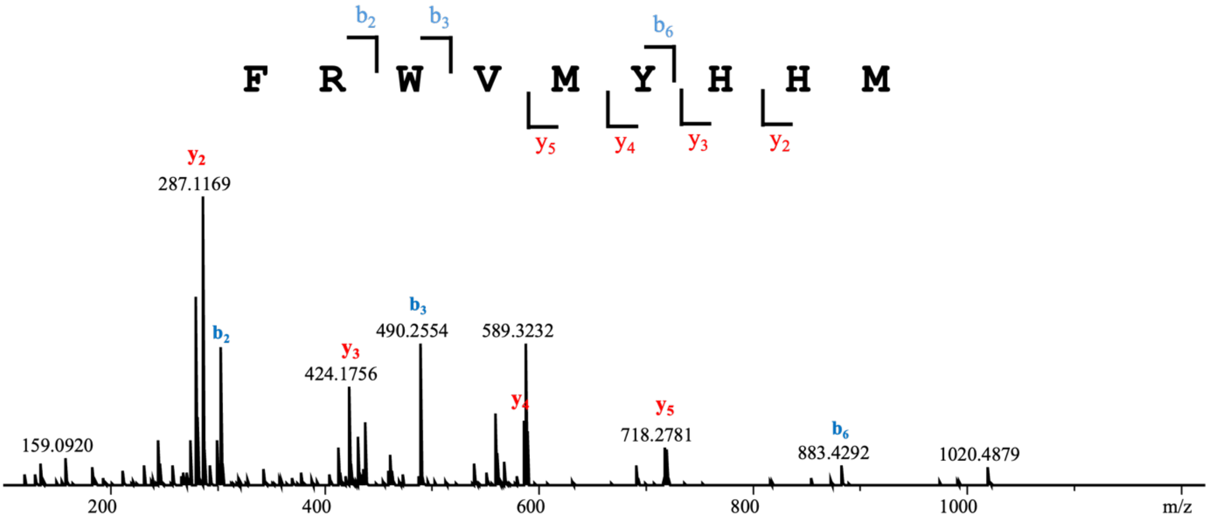


**Linear**

| Ions | Fragment | Observed mass | Calculated mass | z | Δppm |
|---|---|---|---|---|---|
| $y_2$ | HM | 287.1169 | 287.1172 | 1 | 1.04 |
| $y_3$ | HHM | 424.1756 | 424.1761 | 1 | 1.18 |
| $y_4$ | YHHM | 587.2373 | 587.2395 | 1 | 3.75 |
| $y_5$ | MYHHM | 718.2781 | 718.280 | 1 | 2.65 |
| $b_2$ | FR | 304.1763 | 304.1768 | 1 | 1.64 |
| $b_3$ | FRW | 490.2554 | 490.2561 | 1 | 1.43 |
| $b_6$ | FRWVMY | 883.4292 | 883.4283 | 1 | 1.02 |

GpeC enzymatic assay with WRWVMYFHM. A 5-hour reaction was performed before quenching by adding an equivalent volume of methanol. No cyclized product was observed. Tandem mass analysis revealed the linear peptide ions of $y_2$, $y_3$, $y_4$, $y_5$, $y_7$, $b_2$, and $b_3$.

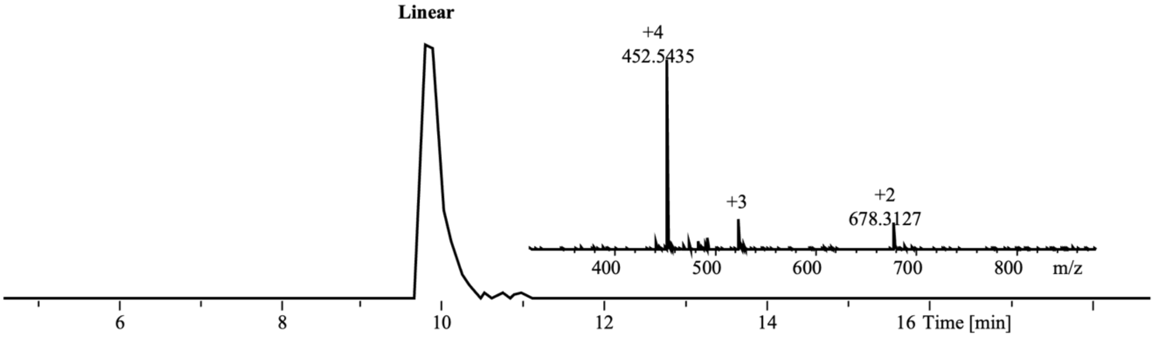


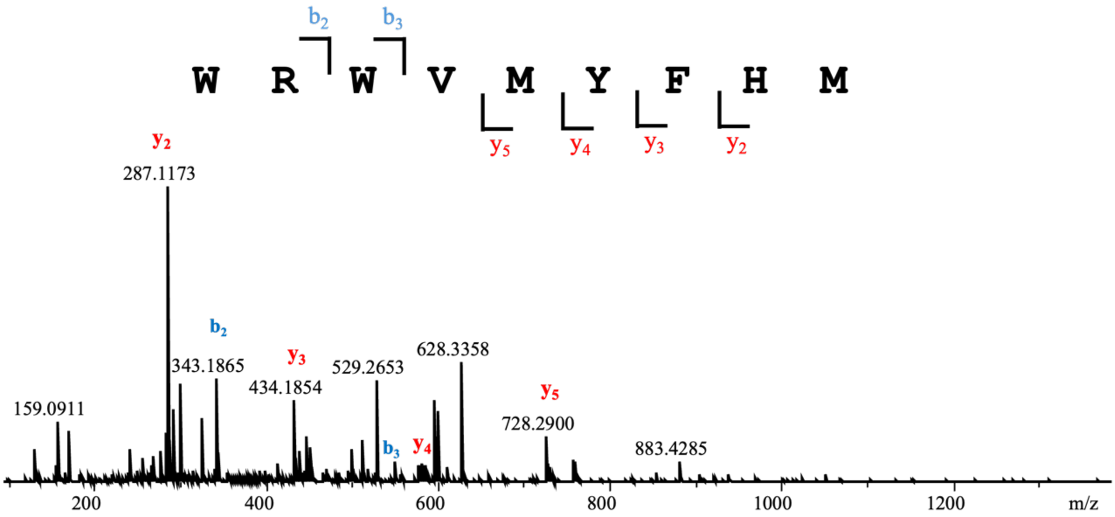


| Linear | | | | | |
|---|---|---|---|---|---|
| Ions | Fragment | Observed mass | Calculated mass | z | Δppm |
| $y_2$ | HM | 287.1173 | 287.1172 | 1 | 0.35 |
| $y_3$ | FHM | 434.1854 | 434.1857 | 1 | 0.69 |
| $y_4$ | YFHM | 597.2461 | 597.249 | 1 | 4.86 |
| $y_5$ | MYFHM | 728.2900 | 728.2895 | 1 | 0.69 |
| $b_2$ | WR | 343.1865 | 343.1877 | 1 | 3.5 |
| $b_3$ | WRW | 529.2653 | 529.267 | 1 | 3.21 |

**Random amino acids incorporated in the intracyclic amino acids within the substrate**

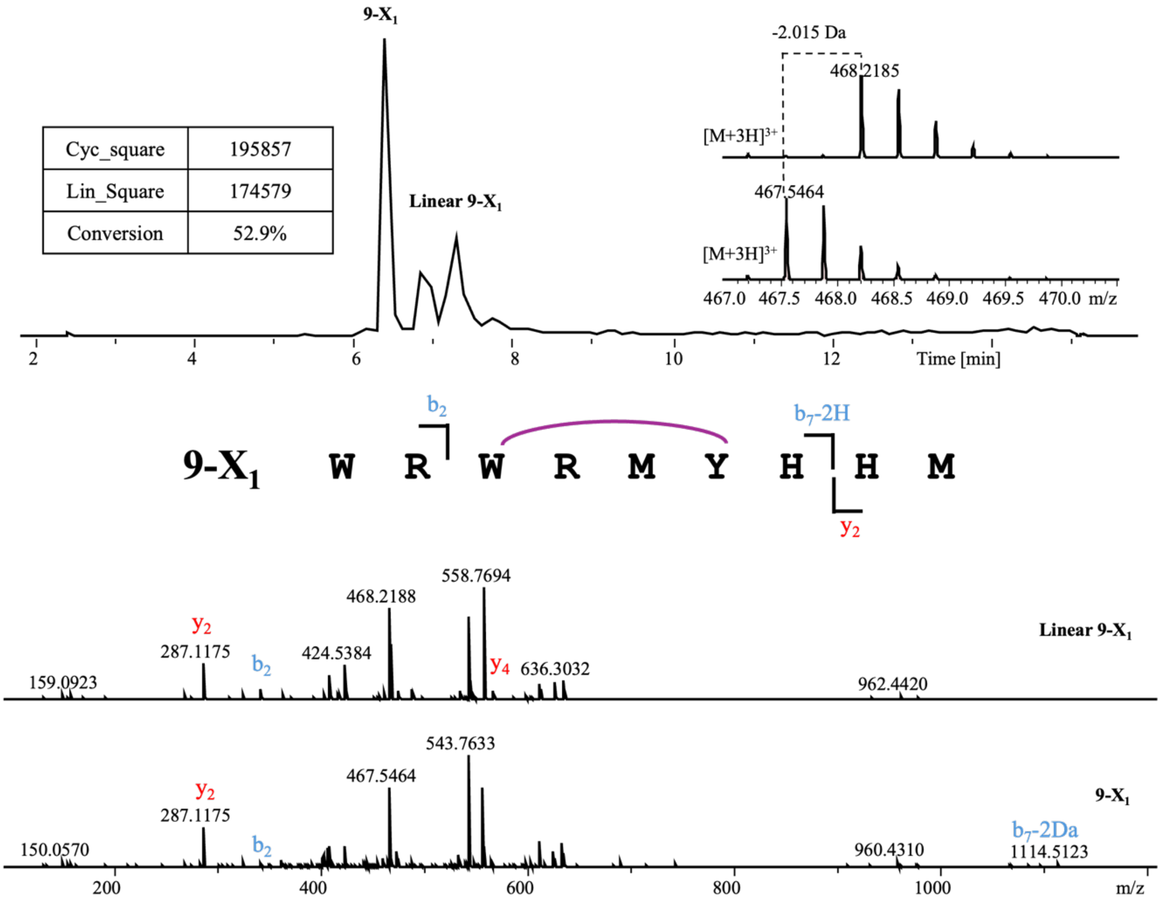


**Linear 9-$X_1$**

| Ions | Fragment | Observed mass | Calculated mass | z | Δppm |
|---|---|---|---|---|---|
| $y_2$ | HM | 287.1175 | 287.1172 | 1 | 1.04 |
| $y_4$ | YHHM | 587.2400 | 587.2395 | 1 | 0.85 |
| $y_5$ | MYHHM | 718.2767 | 718.28 | 1 | 4.59 |
| $b_2$ | WR | 343.1878 | 343.1877 | 1 | 0.29 |
| $b_6$ | WRWRMY | 979.4709 | 979.4719 | 1 | 1.02 |

**Cyclized 9-$X_1$**

| Ions | Fragment | Observed mass | Calculated mass | z | Δppm |
|---|---|---|---|---|---|
| $y_2$ | HM | 287.1175 | 287.1172 | 1 | 1.04 |
| $b_2$ | WR | 343.1891 | 343.1877 | 1 | 4.08 |
| $b_7$-2H | WRWRMYH | 1114.5123 | 1114.515 | 1 | 2.42 |

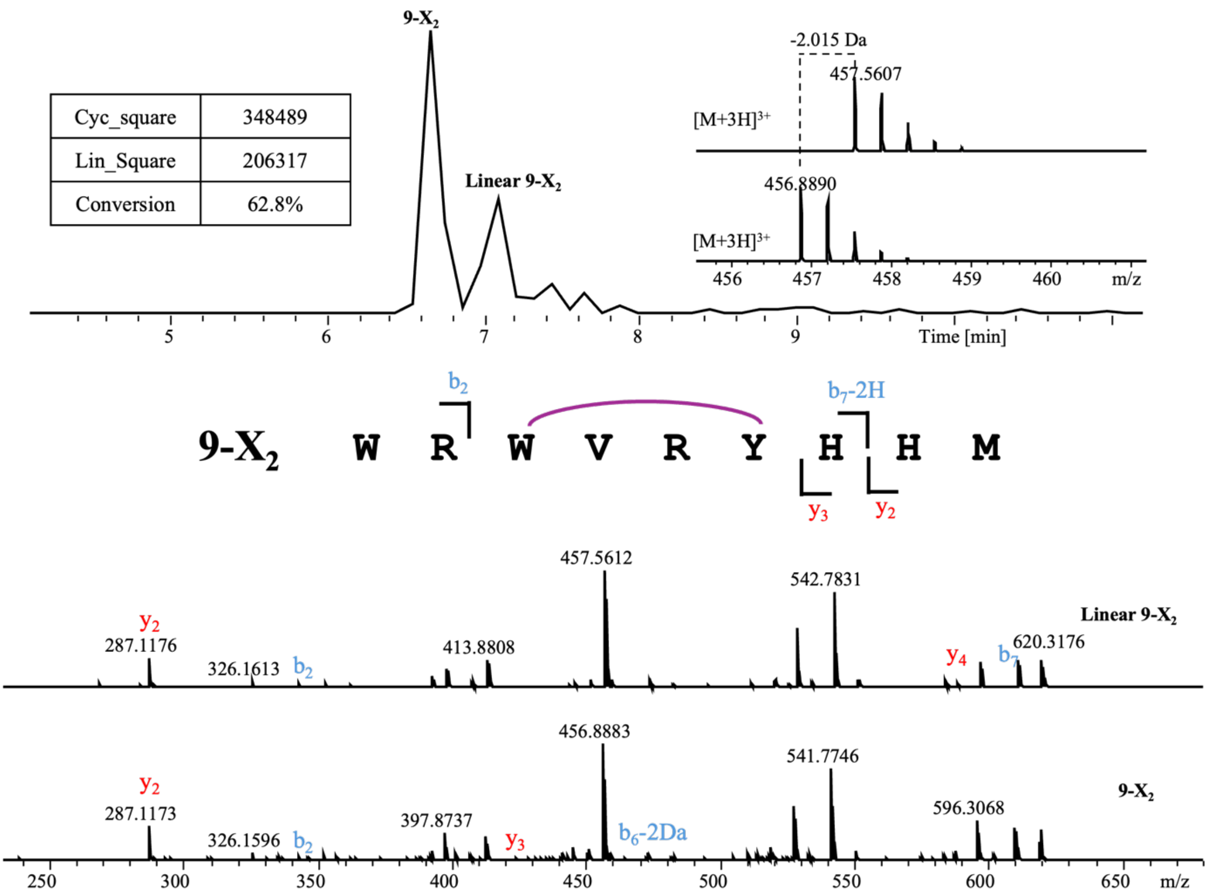


**Linear 9-$X_2$**

| Ions | Fragment | Observed mass | Calculated mass | z | Δppm |
|---|---|---|---|---|---|
| $y_2$ | HM | 287.1176 | 287.1172 | 1 | 1.39 |
| $y_5$ | YHHM | 587.2387 | 587.2395 | 1 | 1.36 |
| $b_2$ | WR | 343.1867 | 343.1877 | 1 | 2.91 |
| $b_7$ | WRWVRYH | 611.3120 | 611.3125 | 2 | 0.82 |

**Cyclized 9-$X_2$**

| Ions | Fragment | Observed mass | Calculated mass | z | Δppm |
|---|---|---|---|---|---|
| $y_2$ | HM | 287.1173 | 287.1172 | 1 | 0.35 |
| $y_3$ | HHM | 424.1766 | 424.1761 | 1 | 1.18 |
| $b_2$ | WR | 343.1876 | 343.1877 | 1 | 0.29 |
| $b_6$-2H | WRWVRY | 473.2471 | 473.2457 | 2 | 2.96 |

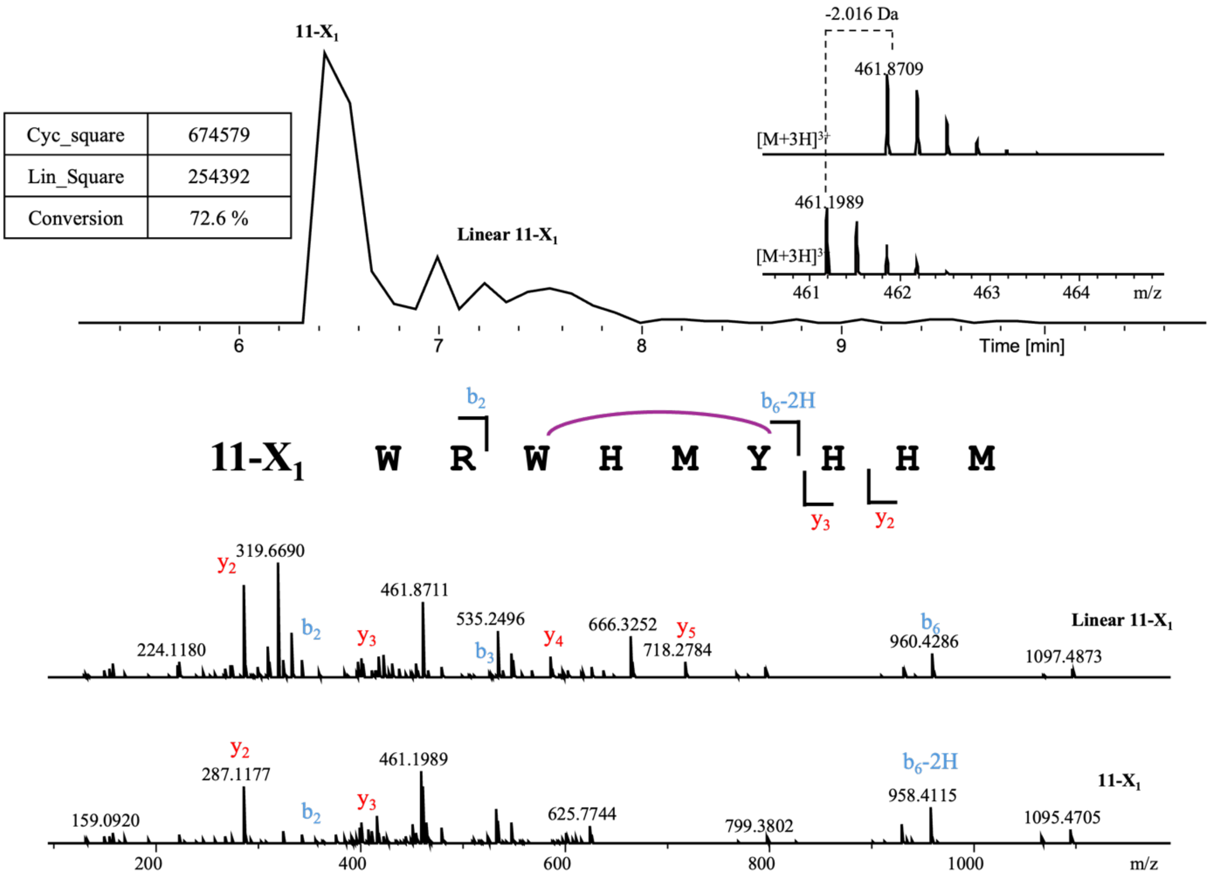


| linear 11-X$_1$ | | | | | |
|---|---|---|---|---|---|
| Ions | Fragment | Observed mass | Calculated mass | z | Δppm |
| $y_2$ | HM | 287.1172 | 287.1172 | 1 | 0 |
| $y_3$ | HHM | 424.1758 | 424.1761 | 1 | 0.71 |
| $y_4$ | YHHM | 587.2390 | 587.2395 | 1 | 0.85 |
| $y_5$ | MYHHM | 718.2784 | 718.28 | 1 | 2.23 |
| $b_2$ | WR | 343.1869 | 343.1877 | 1 | 2.33 |
| $b_3$ | WRW | 529.2671 | 529.267 | 1 | 0.19 |
| $b_6$ | WRWHMY | 960.4286 | 960.4297 | 1 | 1.15 |

| 11-X$_1$ | | | | | |
|---|---|---|---|---|---|
| Ions | Fragment | Observed mass | Calculated mass | z | Δppm |
| $y_2$ | HM | 287.1177 | 287.1172 | 1 | 1.74 |
| $y_3$ | HHM | 424.1741 | 424.1761 | 1 | 4.72 |
| $b_2$ | WR | 343.1860 | 343.1877 | 1 | 4.95 |
| $b_6$-2H | WRWHMY | 958.4115 | 958.4141 | 1 | 2.71 |

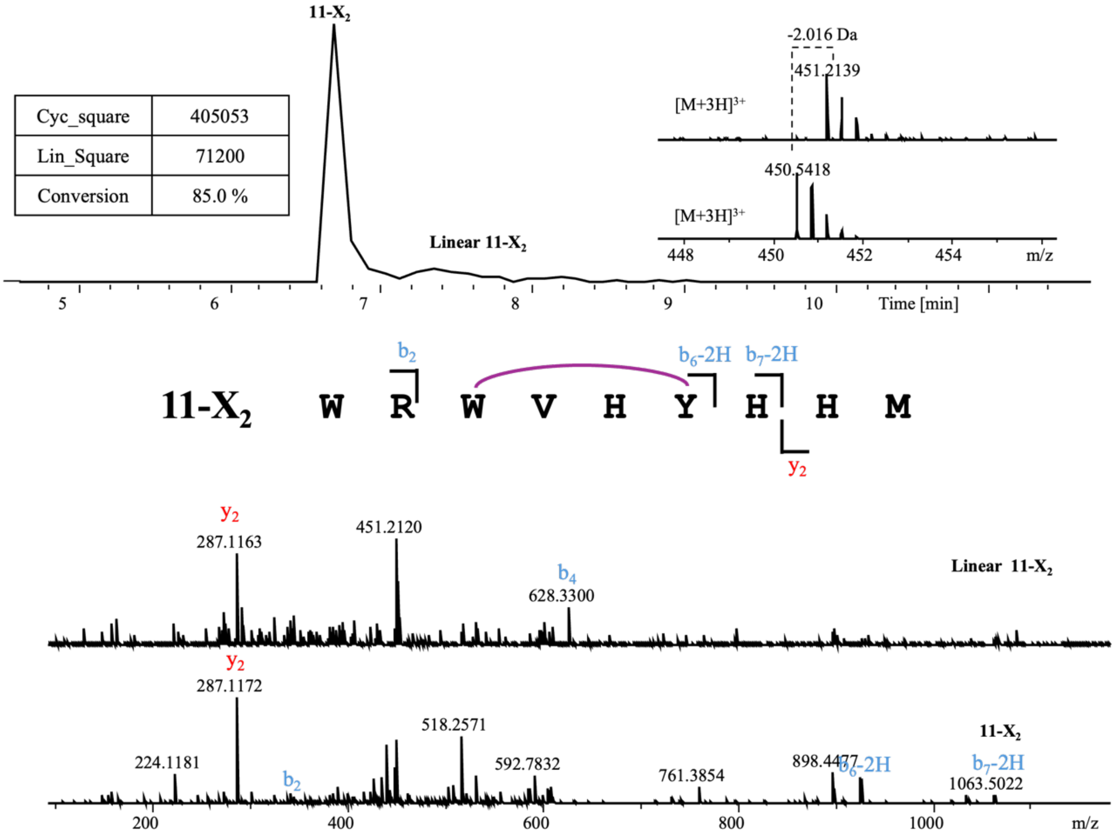


**Linear 11-$X_2$**

| Ions | Fragment | Observed mass | Calculated mass | z | Δppm |
|---|---|---|---|---|---|
| $y_2$ | HM | 287.1163 | 287.1172 | 1 | 3.13 |
| $b_4$ | WRWV | 628.3300 | 628.3354 | 1 | 8.59 |

**11-$X_2$**

| Ions | Fragment | Observed mass | Calculated mass | z | Δppm |
|---|---|---|---|---|---|
| $y_2$ | HHM | 424.1758 | 424.1761 | 1 | 0.71 |
| $b_2$ | WR | 343.1902 | 343.1877 | 1 | 7.28 |
| $b_6$-2H | WRWVHY | 926.4427 | 926.442 | 1 | 0.76 |
| $b_7$-2H | WRWVHYH | 1063.5022 | 1063.501 | 1 | 1.13 |

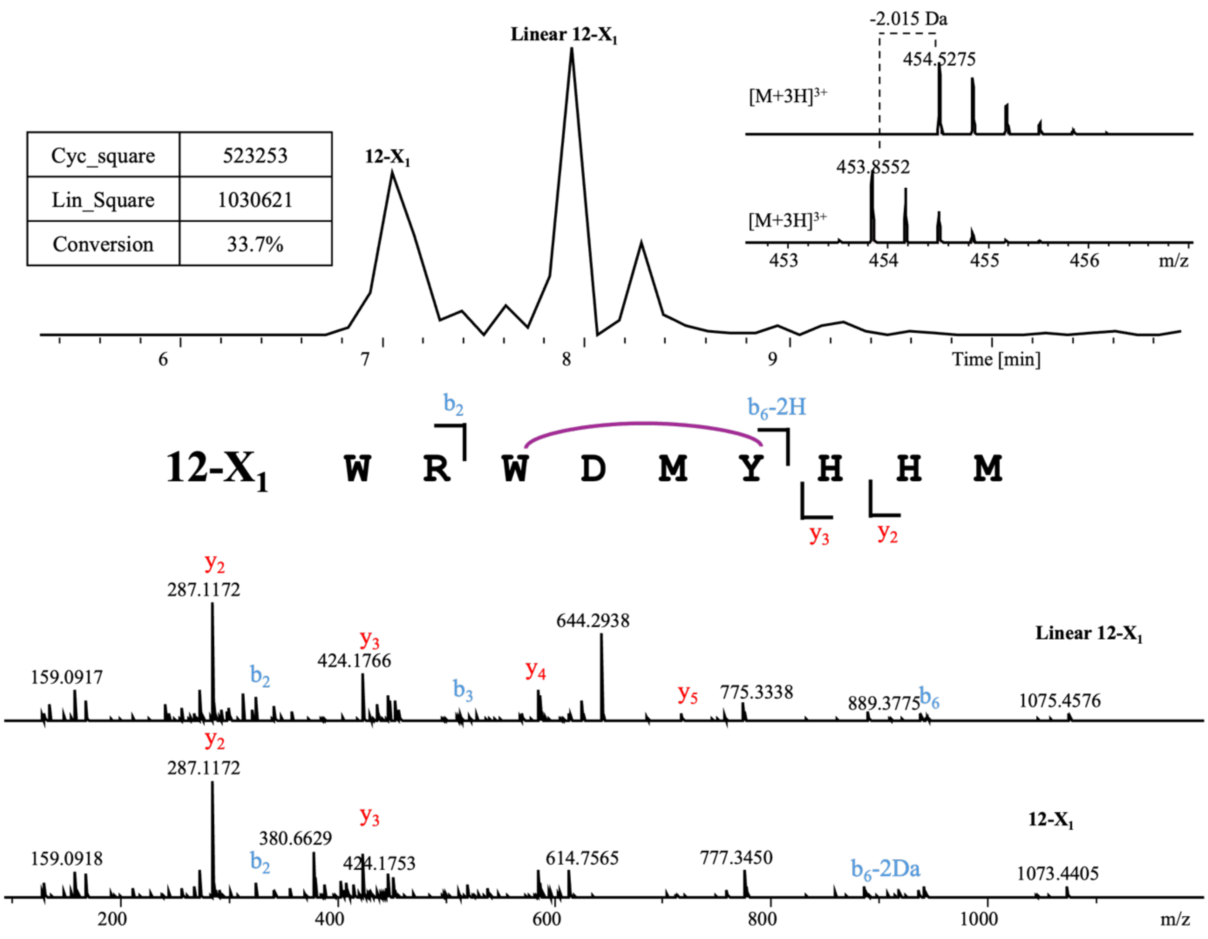


**Linear 12-X$_1$**

| Ions | Fragment | Observed mass | Calculated mass | z | Δppm |
|---|---|---|---|---|---|
| $y_2$ | HM | 287.1174 | 287.1172 | 1 | 0.7 |
| $y_3$ | HHM | 424.1766 | 424.1761 | 1 | 1.18 |
| $y_4$ | YHHM | 587.2394 | 587.2395 | 1 | 0.17 |
| $y_5$ | MYHHM | 718.2799 | 718.28 | 1 | 0.14 |
| $b_2$ | WR | 343.1873 | 343.1877 | 1 | 1.17 |
| $b_3$ | WRW | 529.2664 | 529.267 | 1 | 1.13 |
| $b_6$ | WRWDMY | 938.3988 | 938.3948 | 1 | 4.26 |

**12-X$_1$**

| Ions | Fragment | Observed mass | Calculated mass | z | Δppm |
|---|---|---|---|---|---|
| $y_2$ | HM | 287.1172 | 287.1172 | 1 | <0.1 |
| $y_3$ | HHM | 424.1753 | 424.1761 | 1 | 1.89 |
| $b_2$ | WR | 343.1888 | 343.1877 | 1 | 3.21 |
| $b_6$-2H | WRWDMY | 936.3780 | 936.3821 | 1 | 4.38 |

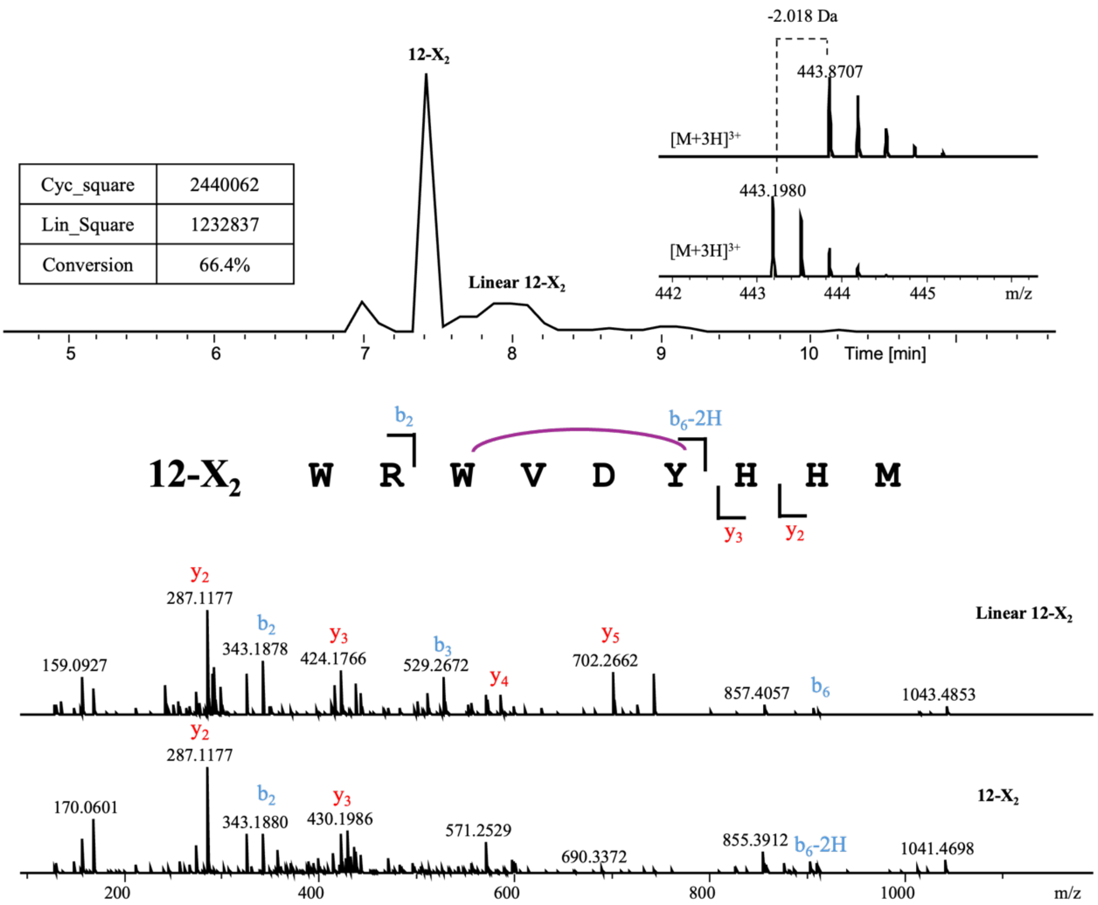


| Linear 12-$X_2$ | | | | | |
|---|---|---|---|---|---|
| Ions | Fragment | Observed mass | Calculated mass | z | Δppm |
| $y_2$ | HM | 287.1177 | 287.1172 | 1 | 1.74 |
| $y_3$ | HHM | 424.1766 | 424.1761 | 1 | 1.18 |
| $y_4$ | YHHM | 587.2400 | 587.2395 | 1 | 0.85 |
| $y_5$ | DYHHM | 702.2662 | 702.2667 | 1 | 0.71 |
| $b_2$ | WR | 343.1878 | 343.1877 | 1 | 0.29 |
| $b_3$ | WRW | 529.2672 | 529.267 | 1 | 0.38 |
| $b_6$ | WRWVDY | 906.4264 | 906.4257 | 1 | 0.77 |

| 12-$X_2$ | | | | | |
|---|---|---|---|---|---|
| Ions | Fragment | Observed mass | Calculated mass | z | Δppm |
| $y_2$ | HM | 287.1177 | 287.1172 | 1 | 1.74 |
| $y_3$ | HHM | 424.1763 | 424.1761 | 1 | 0.47 |
| $b_2$ | WR | 343.1880 | 343.1877 | 1 | 0.87 |
| $b_6$-2H | WRWVDY | 904.4109 | 904.41 | 1 | 1 |

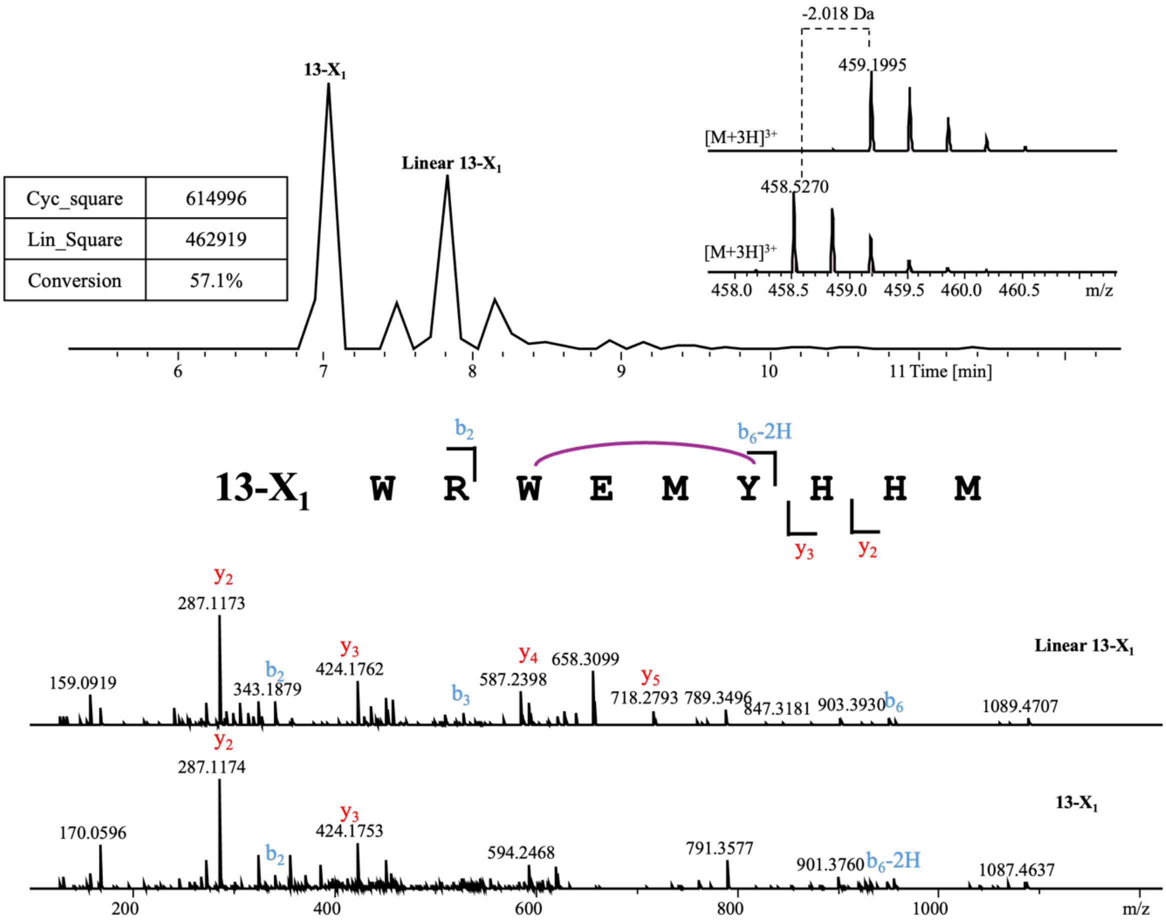


**Linear 13-$X_1$**

| Ions | Fragment | Observed mass | Calculated mass | z | Δppm |
|---|---|---|---|---|---|
| $y_2$ | HM | 287.1173 | 287.1172 | 1 | 0.35 |
| $y_3$ | HHM | 424.1762 | 424.1761 | 1 | 0.24 |
| $y_4$ | YHHM | 587.2398 | 587.2395 | 1 | 0.51 |
| $y_5$ | MYHHM | 718.2793 | 718.28 | 1 | 0.97 |
| $b_2$ | WR | 343.1879 | 343.1877 | 1 | 0.58 |
| $b_3$ | WRW | 529.2603 | 529.267 | 1 | 12.7 |
| $b_6$ | WRWEMY | 952.4132 | 952.4134 | 1 | 0.21 |

**13-$X_1$**

| Ions | Fragment | Observed mass | Calculated mass | z | Δppm |
|---|---|---|---|---|---|
| $y_2$ | HM | 287.1174 | 287.1172 | 1 | 0.7 |
| $y_3$ | HHM | 424.1753 | 424.1761 | 1 | 1.89 |
| $b_2$ | WR | 343.1865 | 343.1877 | 1 | 3.5 |
| $b_6$-2H | WRWEMY | 950.4009 | 950.3978 | 1 | 3.26 |

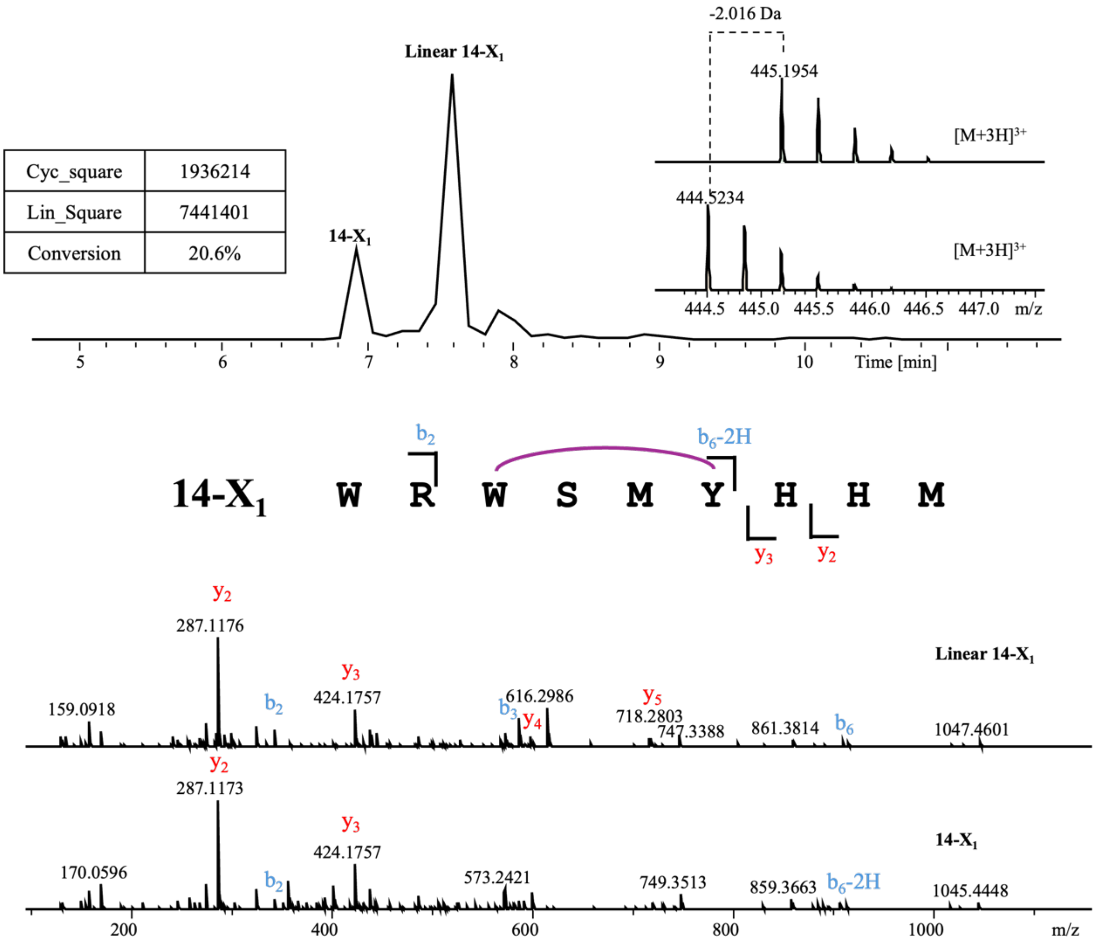


**Linear 14-$X_1$**

| Ions | Fragment | Observed mass | Calculated mass | z | Δppm |
|---|---|---|---|---|---|
| $y_2$ | HM | 287.1176 | 287.1172 | 1 | 1.39 |
| $y_3$ | HHM | 424.1757 | 424.1761 | 1 | 0.94 |
| $y_4$ | YHHM | 587.2395 | 587.2395 | 1 | 0 |
| $y_5$ | MYHHM | 718.2803 | 718.28 | 1 | 0.42 |
| $b_2$ | WR | 343.1869 | 343.1877 | 1 | 2.33 |
| $b_3$ | WRW | 529.2660 | 529.267 | 1 | 1.89 |
| $b_6$ | WRWSMY | 910.4003 | 910.4029 | 1 | 2.86 |

**14-$X_1$**

| Ions | Fragment | Observed mass | Calculated mass | z | Δppm |
|---|---|---|---|---|---|
| $y_2$ | HM | 287.1173 | 287.1172 | 1 | 0.35 |
| $y_3$ | HHM | 424.1757 | 424.1761 | 1 | 0.94 |
| $b_2$ | WR | 343.1876 | 343.1877 | 1 | 0.29 |
| $b_6$-2H | WRWSMY | 908.3865 | 908.3872 | 1 | 0.77 |

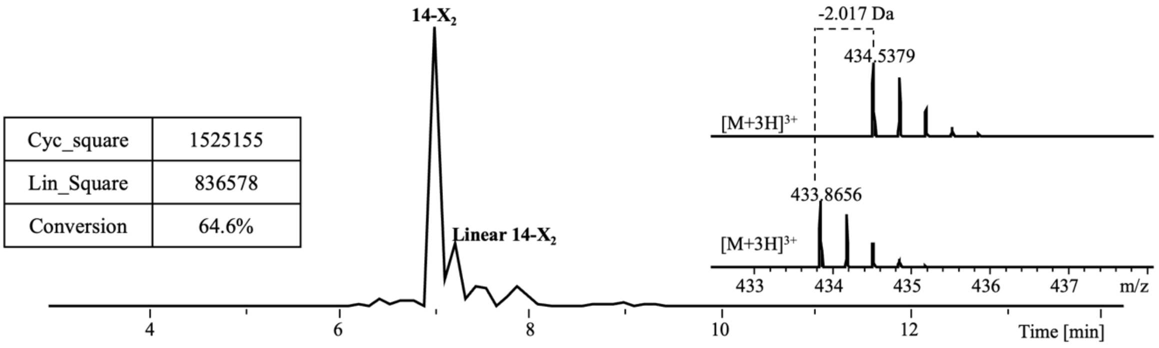


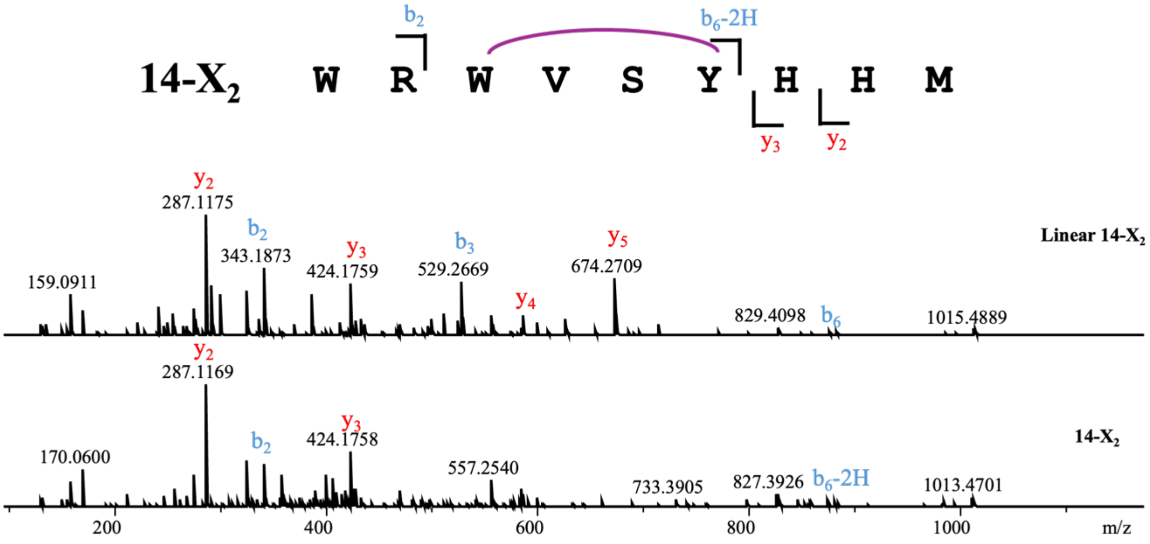


**Linear 14-X$_2$**

| Ions | Fragment | Observed mass | Calculated mass | z | Δppm |
|---|---|---|---|---|---|
| $y_2$ | HM | 287.1175 | 287.1172 | 1 | 1.04 |
| $y_3$ | HHM | 424.1759 | 424.1761 | 1 | 0.47 |
| $y_4$ | YHHM | 587.2391 | 587.2395 | 1 | 0.68 |
| $y_5$ | SYHHM | 674.2709 | 674.2715 | 1 | 0.89 |
| $b_2$ | WR | 343.1873 | 343.1877 | 1 | 1.17 |
| $b_3$ | WRW | 529.2669 | 529.267 | 1 | 0.19 |
| $b_6$ | WRWVSY | 878.4293 | 878.4308 | 1 | 1.71 |

**14-X$_2$**

| Ions | Fragment | Observed mass | Calculated mass | z | Δppm |
|---|---|---|---|---|---|
| $y_2$ | HM | 287.1169 | 287.1172 | 1 | 1.04 |
| $y_3$ | HHM | 424.1758 | 424.1761 | 1 | 0.71 |
| $b_2$ | WR | 343.1870 | 343.1877 | 1 | 2.04 |
| $b_6$-2H | WRWVSY | 876.4121 | 876.4151 | 1 | 3.42 |

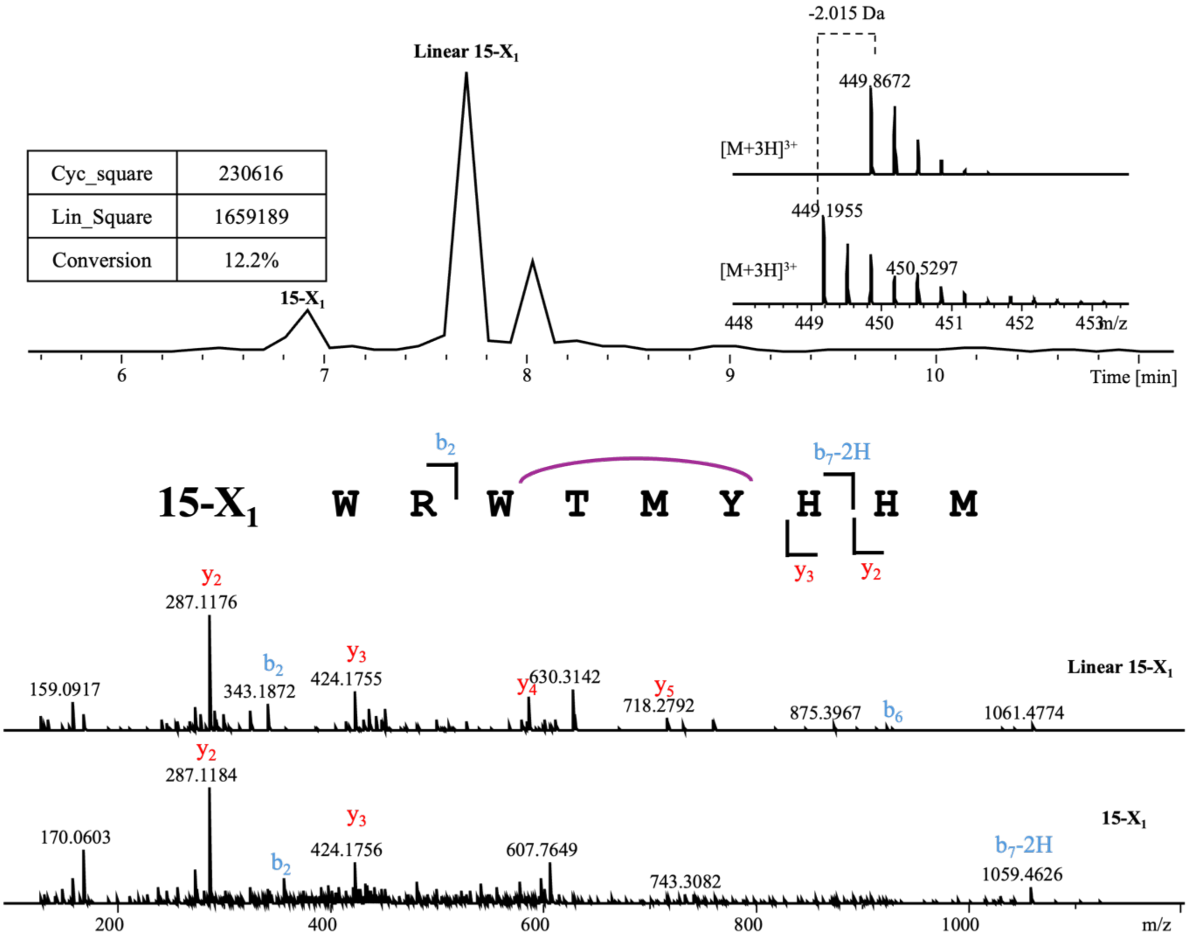


| **Linear 15-$X_1$** | | | | | |
|---|---|---|---|---|---|
| Ions | Fragment | Observed mass | Calculated mass | z | Δppm |
| $y_2$ | HM | 287.1176 | 287.1172 | 1 | 1.39 |
| $y_3$ | HHM | 424.1755 | 424.1761 | 1 | 1.41 |
| $y_4$ | YHHM | 587.2388 | 587.2395 | 1 | 1.19 |
| $y_5$ | MYHHM | 718.2792 | 718.28 | 1 | 1.11 |
| $b_2$ | WR | 343.1872 | 343.1877 | 1 | 1.46 |
| $b_3$ | WRW | 529.2665 | 529.267 | 1 | 0.94 |
| $b_6$ | WRWTMY | 924.4187 | 924.4185 | 1 | 0.22 |

| **15-$X_1$** | | | | | |
|---|---|---|---|---|---|
| Ions | Fragment | Observed mass | Calculated mass | z | Δppm |
| $y_2$ | HM | 287.1184 | 287.1172 | 1 | 4.18 |
| $y_3$ | HHM | 424.1756 | 424.1761 | 1 | 1.18 |
| $b_2$ | WR | 343.1861 | 343.1877 | 1 | 4.66 |
| $b_6$-2H | WRWTMYH | 1059.4626 | 1059.462 | 1 | 0.57 |

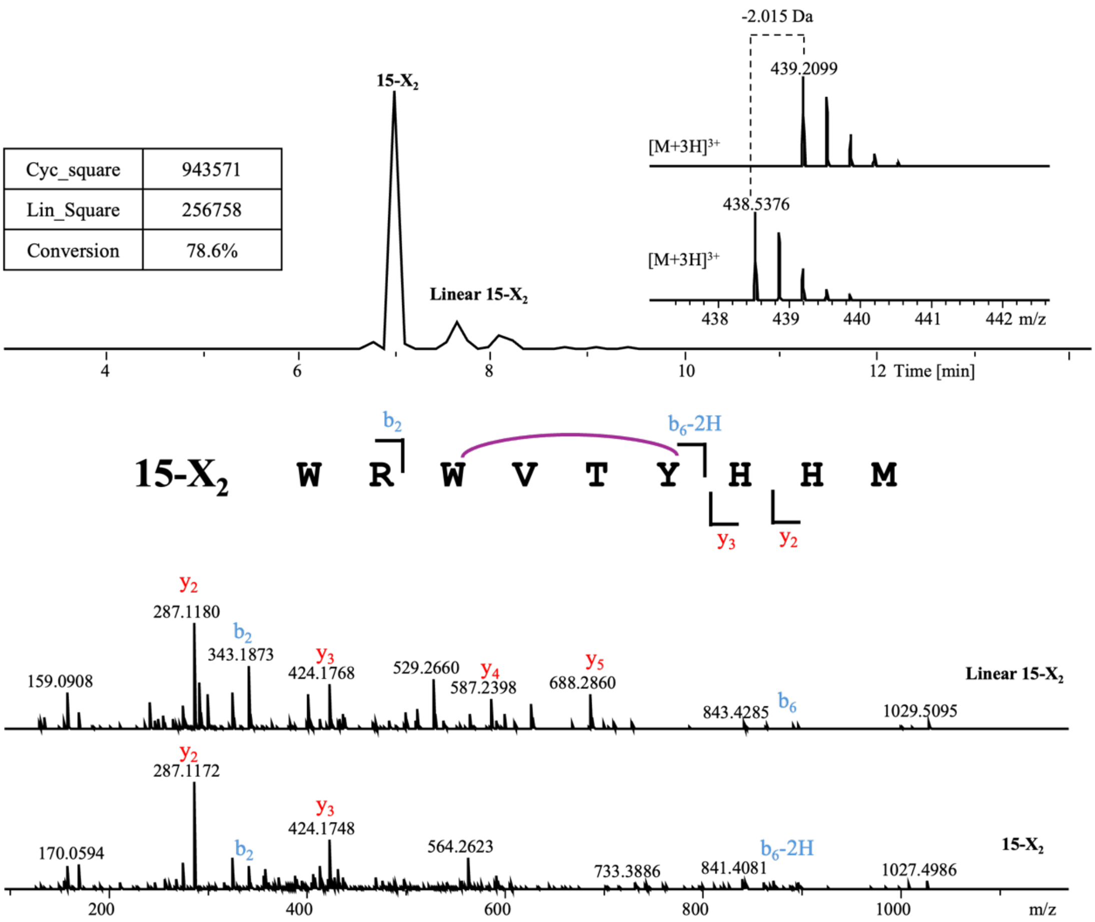


| | |
|---|---|
| Cyc_square | 943571 |
| Lin_Square | 256758 |
| Conversion | 78.6% |

**Linear 15-$X_2$**

| Ions | Fragment | Observed mass | Calculated mass | z | Δppm |
|---|---|---|---|---|---|
| $y_2$ | HM | 287.1180 | 287.1172 | 1 | 2.79 |
| $y_3$ | HHM | 424.1768 | 424.1761 | 1 | 1.65 |
| $y_4$ | YHHM | 587.2398 | 587.2395 | 1 | 0.51 |
| $y_5$ | TYHHM | 688.2860 | 688.2872 | 1 | 1.74 |
| $b_2$ | WR | 343.1873 | 343.1877 | 1 | 1.17 |
| $b_3$ | WRW | 529.2660 | 529.267 | 1 | 1.89 |
| $b_6$ | WRWVTY | 892.4451 | 892.4464 | 1 | 1.46 |

**15-$X_2$**

| Ions | Fragment | Observed mass | Calculated mass | z | Δppm |
|---|---|---|---|---|---|
| $y_2$ | HM | 287.1172 | 287.1172 | 1 | <0.1 |
| $y_3$ | HHM | 424.1748 | 424.1761 | 1 | 3.06 |
| $b_2$ | WR | 343.1873 | 343.1877 | 1 | 1.17 |
| $b_6$-2H | WRWVTY | 890.4222 | 890.4308 | 1 | 9.66 |

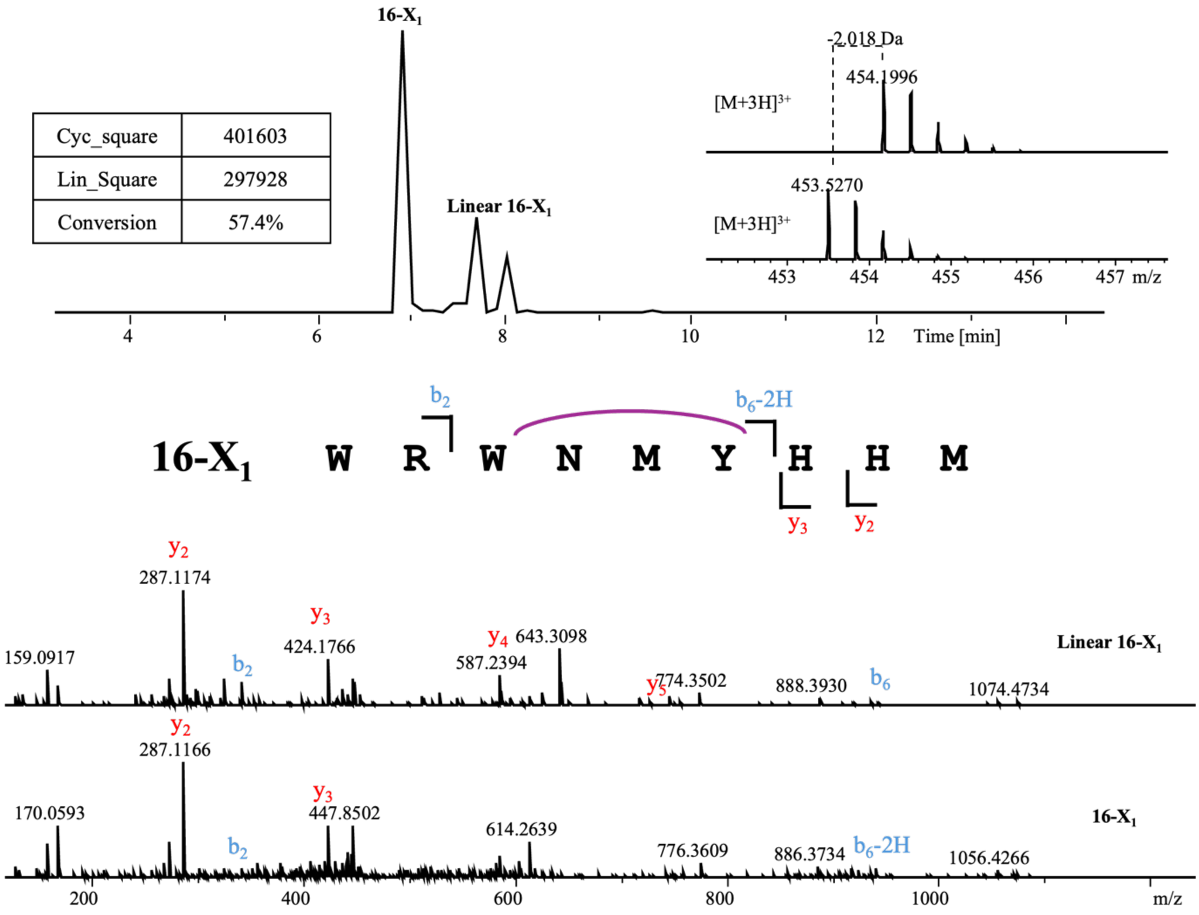


**Linear 16-X$_1$**

| Ions | Fragment | Observed mass | Calculated mass | z | Δppm |
|---|---|---|---|---|---|
| $y_2$ | HM | 287.1174 | 287.1172 | 1 | 0.7 |
| $y_3$ | HHM | 424.1766 | 424.1761 | 1 | 1.18 |
| $y_4$ | YHHM | 587.2394 | 587.2395 | 1 | 0.17 |
| $y_5$ | MYHHM | 718.2799 | 718.28 | 1 | 0.14 |
| $b_2$ | WR | 343.1879 | 343.1877 | 1 | 0.58 |
| $b_3$ | WRW | 529.2664 | 529.267 | 1 | 1.13 |
| $b_6$ | WRWNMY | 937.4142 | 937.4138 | 1 | 0.43 |

**16-X$_1$**

| Ions | Fragment | Observed mass | Calculated mass | z | Δppm |
|---|---|---|---|---|---|
| $y_2$ | HM | 287.1166 | 287.1172 | 1 | 2.09 |
| $y_3$ | HHM | 424.1752 | 424.1761 | 1 | 2.12 |
| $b_2$ | WR | 343.1870 | 343.1877 | 1 | 2.04 |
| $b_6$-2H | WRWNMY | 935.3993 | 935.3981 | 1 | 1.28 |

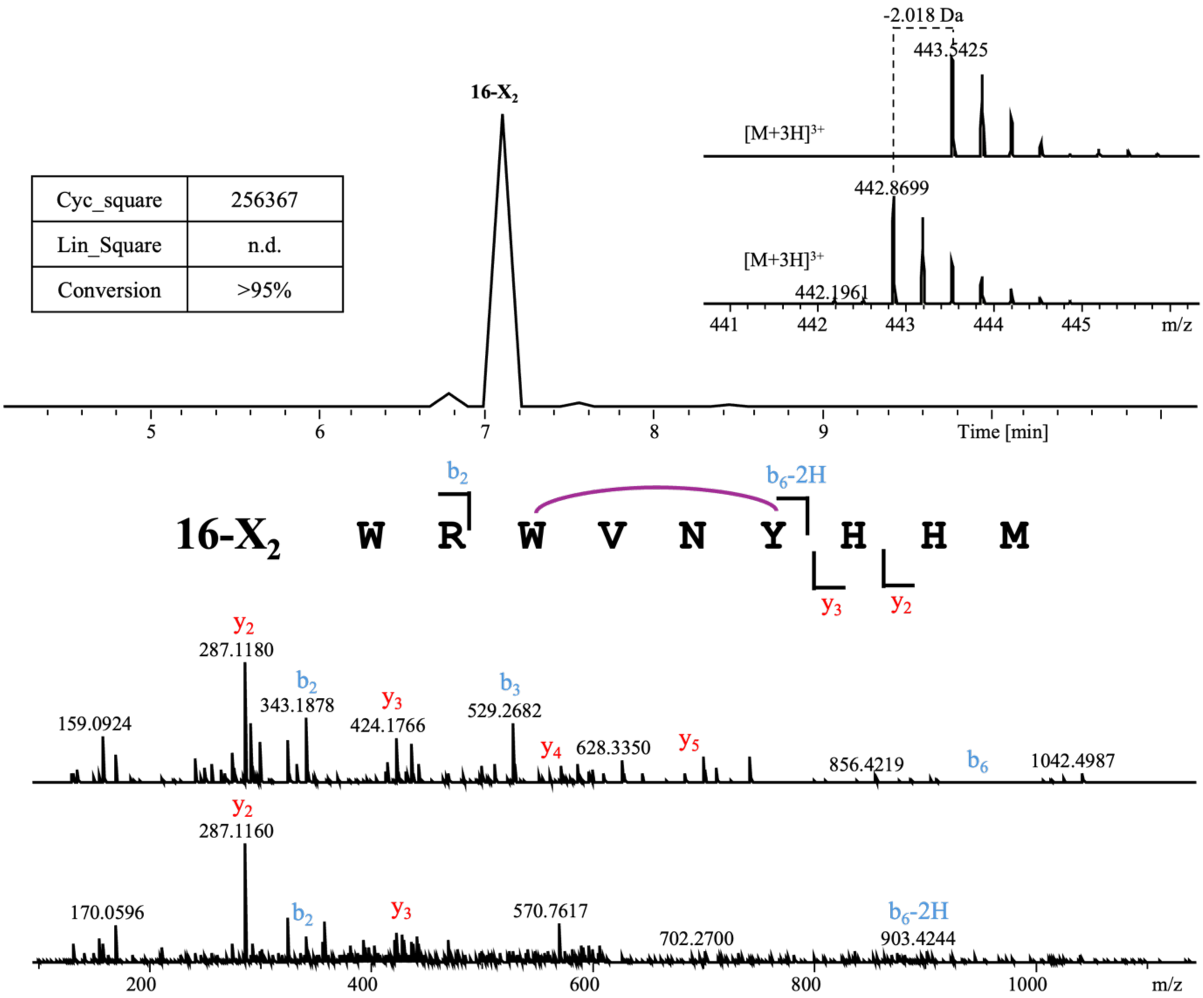


**Linear 16-$X_2$**

| Ions | Fragment | Observed mass | Calculated mass | z | Δppm |
|---|---|---|---|---|---|
| $y_2$ | HM | 287.1180 | 287.1172 | 1 | 2.79 |
| $y_3$ | HHM | 424.1766 | 424.1761 | 1 | 1.18 |
| $y_4$ | YHHM | 587.2408 | 587.2395 | 1 | 2.21 |
| $y_5$ | NYHHM | 701.2824 | 701.2824 | 1 | <0.1 |
| $b_2$ | WR | 343.1878 | 343.1877 | 1 | 0.29 |
| $b_3$ | WRW | 529.2682 | 529.267 | 1 | 2.27 |

**16-$X_2$**

| Ions | Fragment | Observed mass | Calculated mass | z | Δppm |
|---|---|---|---|---|---|
| $y_2$ | HM | 287.1160 | 287.1172 | 1 | 4.18 |
| $y_3$ | HHM | 424.1784 | 424.1761 | 1 | 5.42 |
| $b_2$ | WR | 343.1889 | 343.1877 | 1 | 3.5 |
| $b_6$-2H | WRWVNY | 903.4244 | 903.426 | 1 | 1.77 |

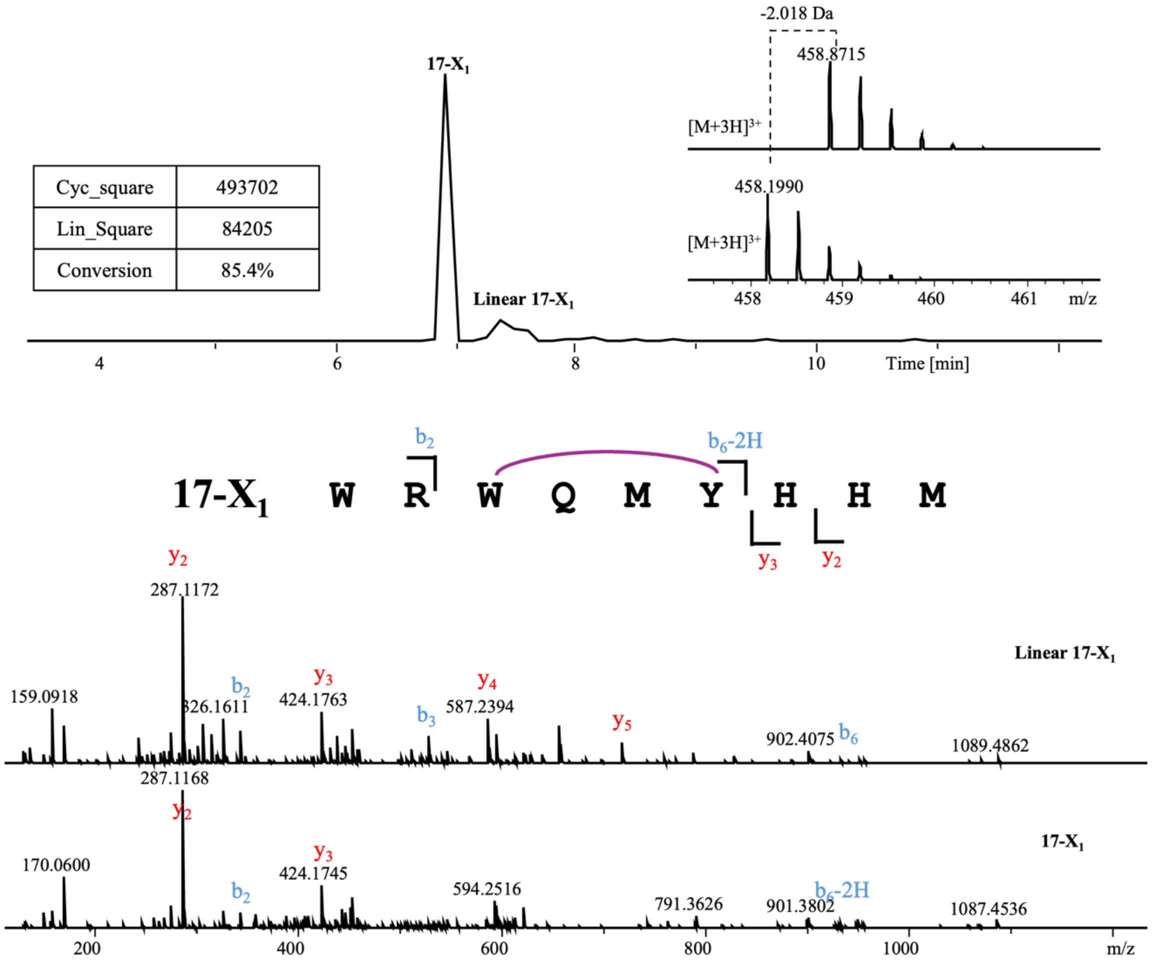


| | | Linear 17-$X_1$ | | | |
|---|---|---|---|---|---|
| Ions | Fragment | Observed mass | Calculated mass | z | Δppm |
| $y_2$ | HM | 287.1172 | 287.1172 | 1 | <0.1 |
| $y_3$ | HHM | 424.1763 | 424.1761 | 1 | 0.47 |
| $y_4$ | YHHM | 587.2394 | 587.2395 | 1 | 0.17 |
| $y_5$ | MYHHM | 718.2782 | 718.28 | 1 | 2.51 |
| $b_2$ | WR | 343.1871 | 343.1877 | 1 | 1.75 |
| $b_3$ | WRW | 529.2665 | 529.267 | 1 | 0.94 |
| $b_6$ | WRWQMY | 951.4304 | 951.4294 | 1 | 1.05 |

| | | 17-$X_1$ | | | |
|---|---|---|---|---|---|
| Ions | Fragment | Observed mass | Calculated mass | z | Δppm |
| $y_2$ | HM | 287.1168 | 287.1172 | 1 | 1.39 |
| $y_3$ | HHM | 424.1745 | 424.1761 | 1 | 3.77 |
| $b_2$ | WR | 343.1863 | 343.1877 | 1 | 4.08 |
| $b_6$-2H | WRWQMY | 949.4138 | 949.4138 | 1 | <0.1 |

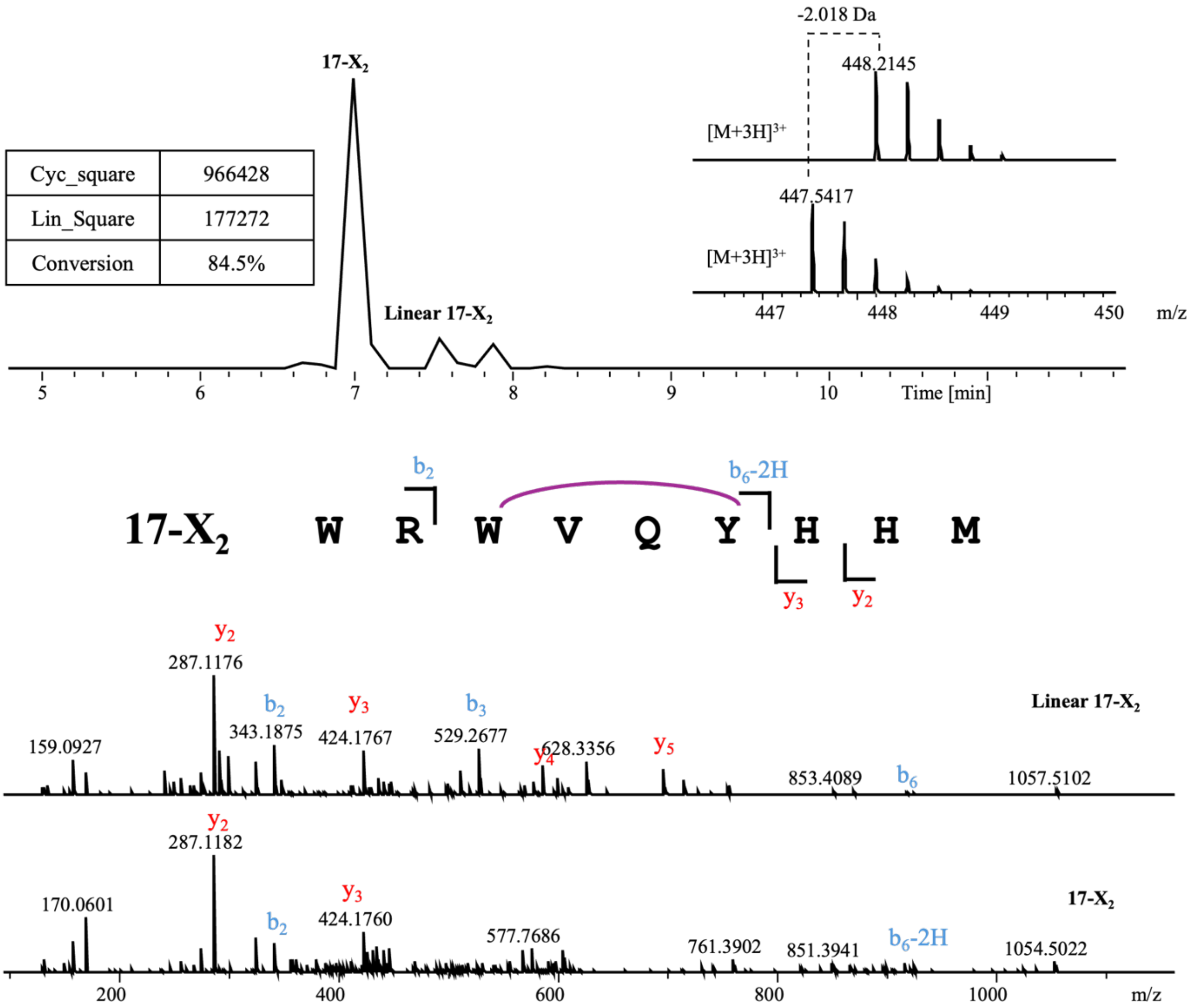


| **Linear 17-X$_2$** | | | | | |
|---|---|---|---|---|---|
| Ions | Fragment | Observed mass | Calculated mass | z | Δppm |
| $y_2$ | HM | 287.1176 | 287.1172 | 1 | 1.39 |
| $y_3$ | HHM | 424.1767 | 424.1761 | 1 | 1.41 |
| $y_4$ | YHHM | 587.2393 | 587.2395 | 1 | 0.34 |
| $y_5$ | QYHHM | 715.2987 | 715.2981 | 1 | 0.84 |
| $b_2$ | WR | 343.1875 | 343.1877 | 1 | 0.58 |
| $b_3$ | WRW | 529.2677 | 529.267 | 1 | 1.32 |
| $b_6$ | WRWVQY | 919.4576 | 919.4573 | 1 | 0.33 |

| **17-X$_2$** | | | | | |
|---|---|---|---|---|---|
| Ions | Fragment | Observed mass | Calculated mass | z | Δppm |
| $y_2$ | HM | 287.1182 | 287.1172 | 1 | 3.48 |
| $y_3$ | HHM | 424.1760 | 424.1761 | 1 | 0.24 |
| $b_2$ | WR | 343.1876 | 343.1877 | 1 | 0.29 |
| $b_6$-2H | WRWVQY | 917.4412 | 917.4417 | 1 | 0.54 |

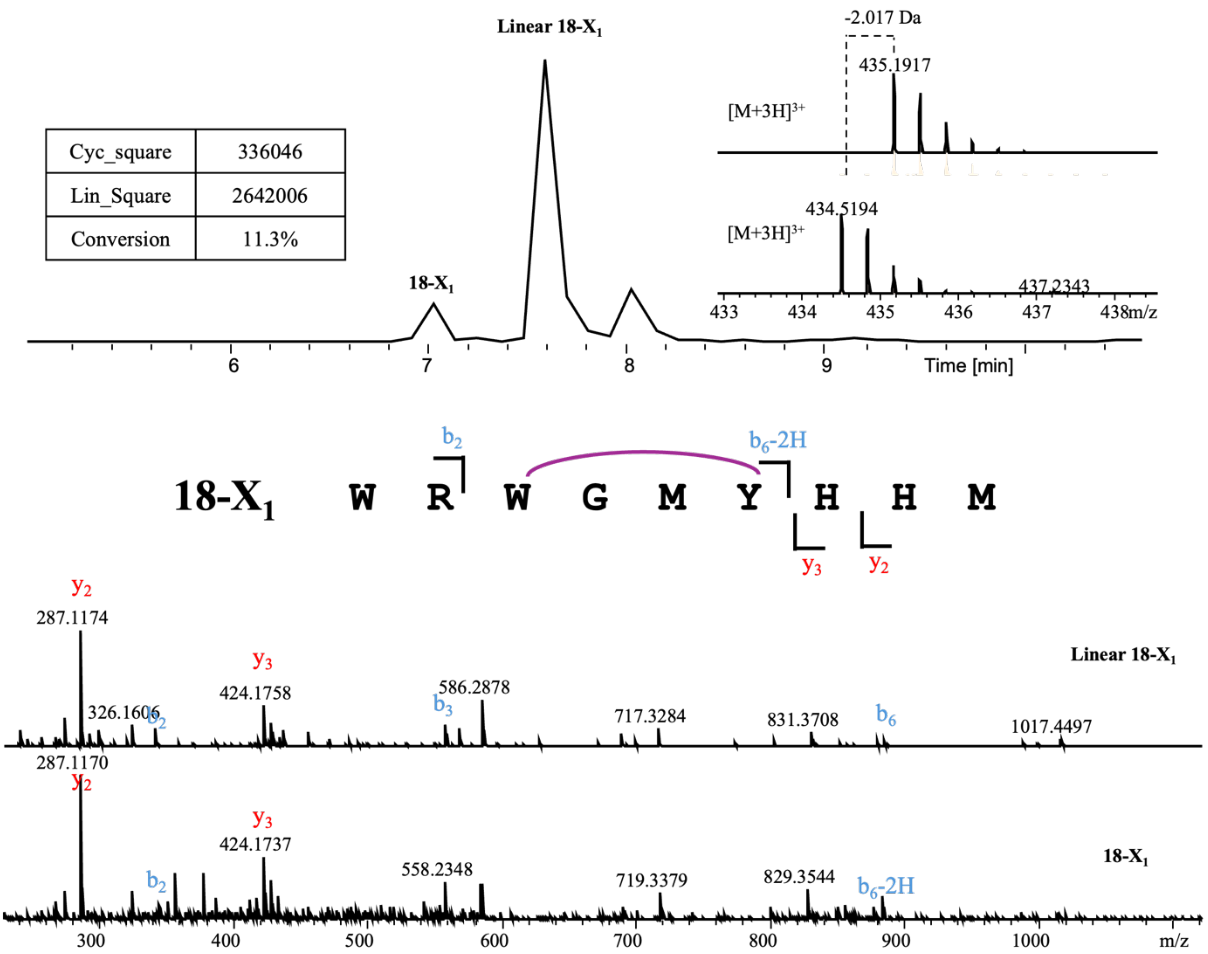


**Linear 18-X$_1$**

| Ions | Fragment | Observed mass | Calculated mass | z | Δppm |
|---|---|---|---|---|---|
| $y_2$ | HM | 287.1174 | 287.1172 | 1 | 0.7 |
| $y_3$ | HHM | 424.1758 | 424.1761 | 1 | 0.71 |
| $y_4$ | YHHM | 587.2403 | 587.2395 | 1 | 1.36 |
| $b_2$ | WR | 343.1871 | 343.1877 | 1 | 1.75 |
| $b_3$ | WRW | 529.2667 | 529.267 | 1 | 0.57 |
| $b_6$ | WRWGMY | 880.3919 | 880.3923 | 1 | 0.45 |

**18-X$_1$**

| Ions | Fragment | Observed mass | Calculated mass | z | Δppm |
|---|---|---|---|---|---|
| $y_2$ | HM | 287.1170 | 287.1172 | 1 | 0.7 |
| $y_3$ | HHM | 424.1737 | 424.1761 | 1 | 5.66 |
| $b_2$ | WR | 343.1863 | 343.1877 | 1 | 4.08 |
| $b_6$-2H | WRWGMY | 878.3727 | 878.3766 | 1 | 4.44 |

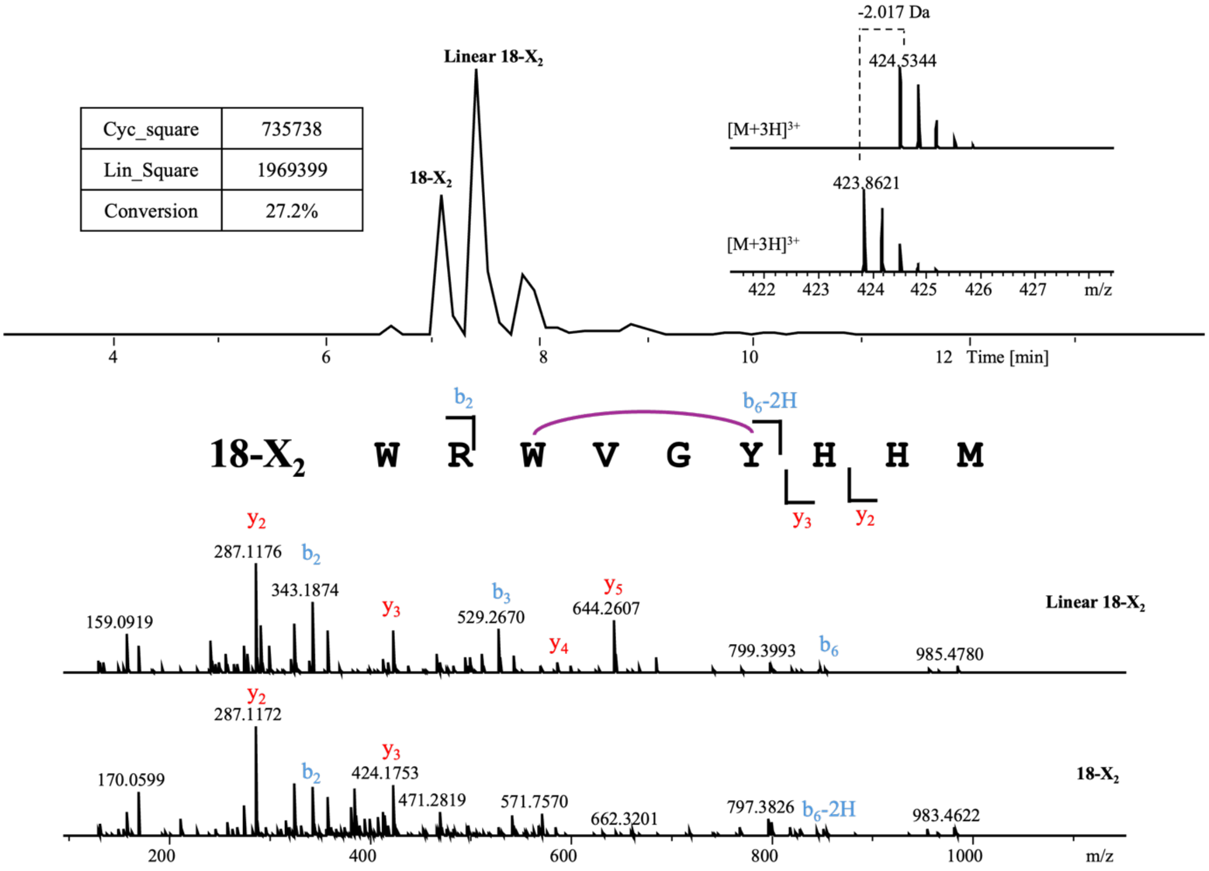


**Linear 18-$X_2$**

| Ions | Fragment | Observed mass | Calculated mass | z | Δppm |
|---|---|---|---|---|---|
| $y_2$ | HM | 287.1176 | 287.1172 | 1 | 1.39 |
| $y_3$ | HHM | 424.1762 | 424.1761 | 1 | 0.24 |
| $y_4$ | YHHM | 587.2398 | 587.2395 | 1 | 0.51 |
| $y_5$ | GYHHM | 644.2607 | 644.2609 | 1 | 0.31 |
| $b_2$ | WR | 343.1874 | 343.1877 | 1 | 0.87 |
| $b_3$ | WRW | 529.2670 | 529.267 | 1 | <0.1 |
| $b_6$ | WRWVGY | 848.4210 | 848.4202 | 1 | 0.94 |

**18-$X_2$**

| Ions | Fragment | Observed mass | Calculated mass | z | Δppm |
|---|---|---|---|---|---|
| $y_2$ | HM | 287.1176 | 287.1172 | 1 | 1.39 |
| $y_3$ | HHM | 424.1753 | 424.1761 | 1 | 1.89 |
| $b_2$ | WR | 343.1874 | 343.1877 | 1 | 0.87 |
| $b_6$-2H | WRWVGY | 846.4042 | 846.4046 | 1 | 0.47 |

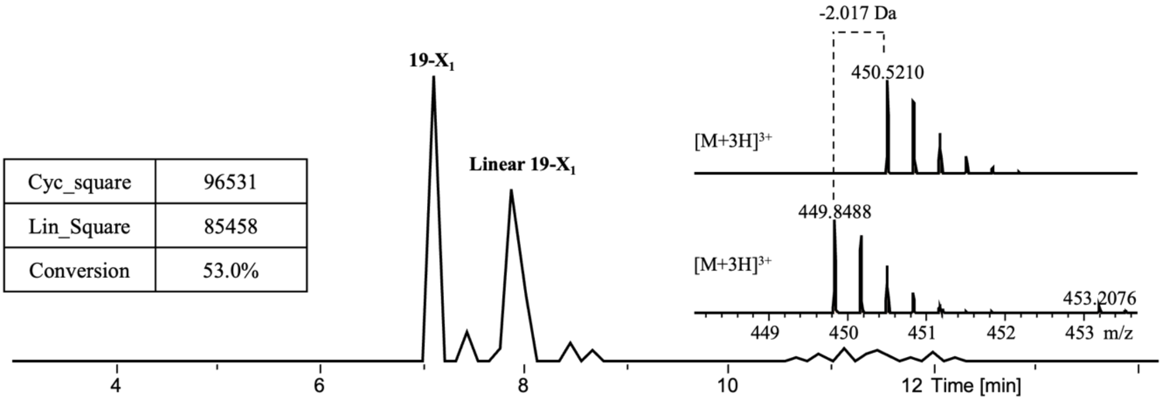


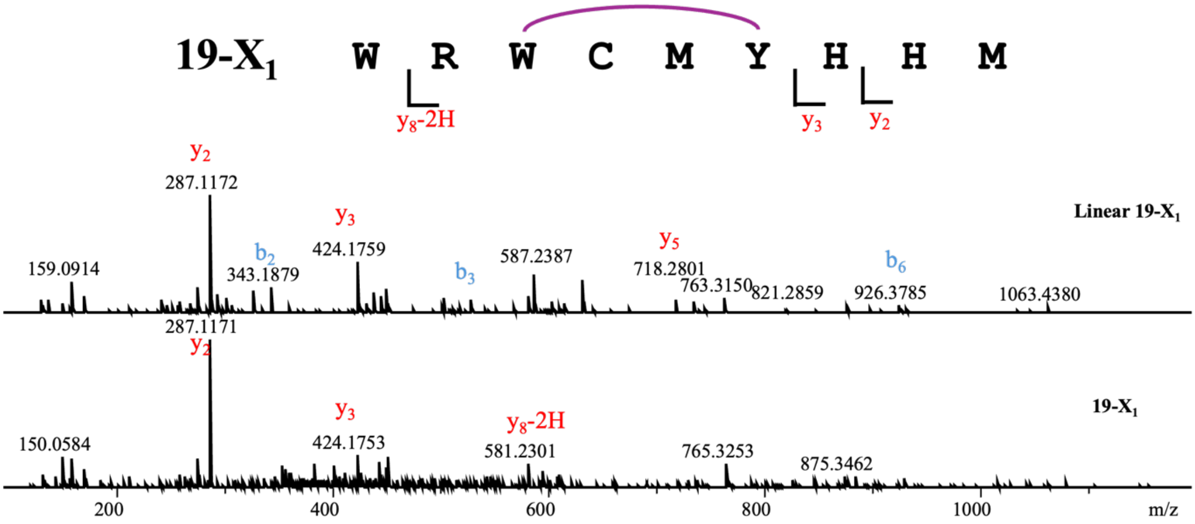


**Linear 19-X$_1$**

| Ions | Fragment | Observed mass | Calculated mass | z | Δppm |
|---|---|---|---|---|---|
| $y_2$ | HM | 287.1172 | 287.1172 | 1 | <0.1 |
| $y_3$ | HHM | 424.1759 | 424.1761 | 1 | 0.47 |
| $y_4$ | YHHM | 587.2387 | 587.2395 | 1 | 1.36 |
| $y_5$ | MYHHM | 718.2801 | 718.28 | 1 | 0.14 |
| $b_2$ | WR | 343.1879 | 343.1877 | 1 | 0.58 |
| $b_3$ | WRW | 529.2683 | 529.267 | 1 | 2.46 |
| $b_6$ | WRWCMY | 926.3785 | 926.38 | 1 | 1.62 |

**19-X$_1$**

| Ions | Fragment | Observed mass | Calculated mass | z | Δppm |
|---|---|---|---|---|---|
| $y_2$ | HM | 287.1171 | 287.1172 | 1 | 0.35 |
| $y_3$ | HHM | 424.1753 | 424.1761 | 1 | 1.89 |
| $y_8$-2H | RWCMYHHM | 581.2301 | 581.2306 | 2 | 0.86 |

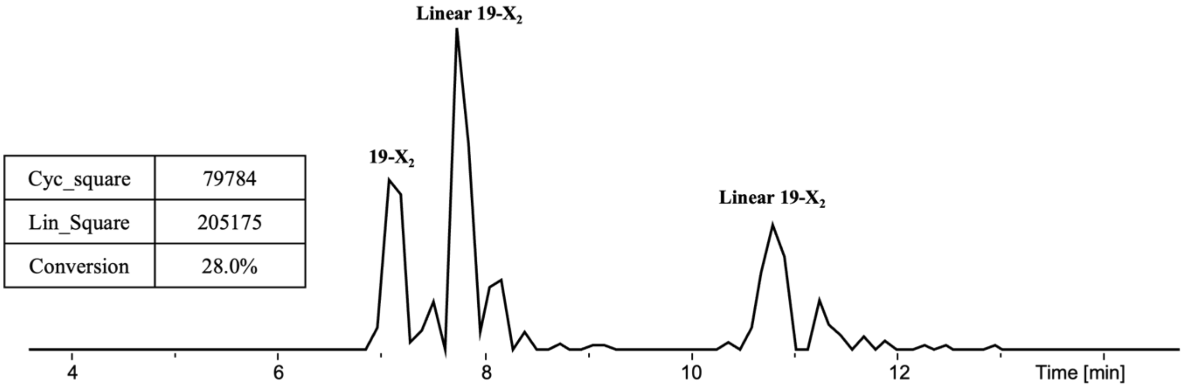


| Cyc_square | 79784 |
|---|---|
| Lin_Square | 205175 |
| Conversion | 28.0% |

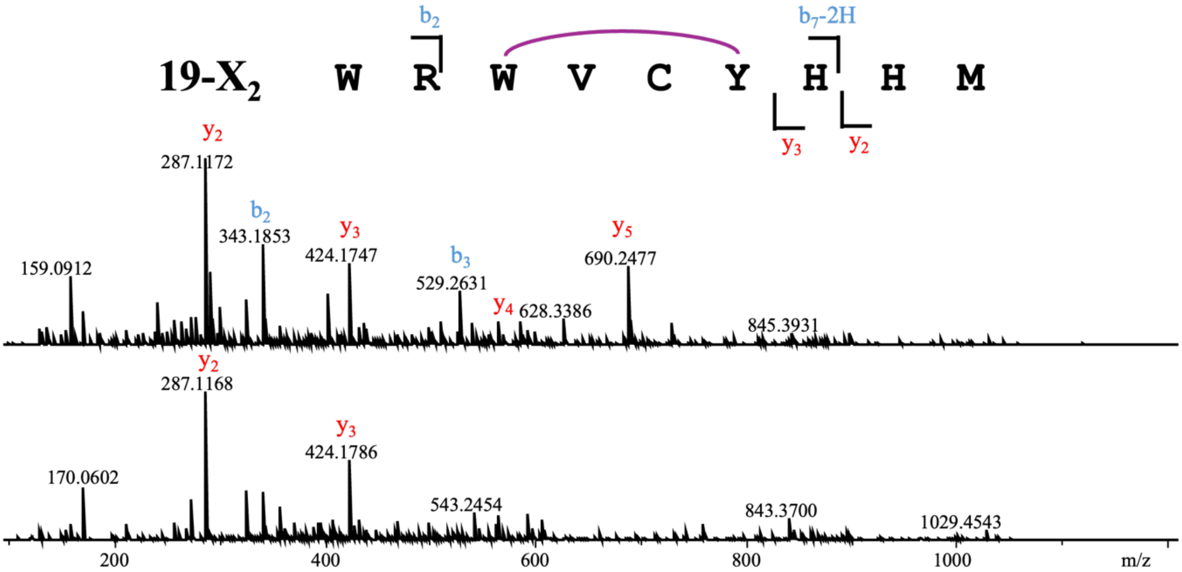


**Linear 19-$X_2$**

| Ions | Fragment | Observed mass | Calculated mass | z | Δppm |
|---|---|---|---|---|---|
| $y_2$ | HM | 287.1172 | 287.1172 | 1 | <0.1 |
| $y_3$ | HHM | 424.1747 | 424.1761 | 1 | 3.3 |
| $y_4$ | YHHM | 587.2412 | 587.2395 | 1 | 2.89 |
| $y_5$ | CYHHM | 690.2477 | 690.2487 | 1 | 1.45 |
| $b_2$ | WR | 343.1853 | 343.1877 | 1 | 6.99 |
| $b_3$ | WRW | 529.2631 | 529.267 | 1 | 7.37 |

**19-$X_2$**

| Ions | Fragment | Observed mass | Calculated mass | z | Δppm |
|---|---|---|---|---|---|
| $y_2$ | HM | 287.1168 | 287.1172 | 1 | 1.39 |
| $y_3$ | HHM | 424.1786 | 424.1761 | 1 | 5.89 |
| $b_7$-2H | WRWVCYH | 1029.4543 | 1029.451 | 1 | 3.21 |
| $b_2$ | WR | 343.1866 | 343.1877 | 1 | 3.21 |

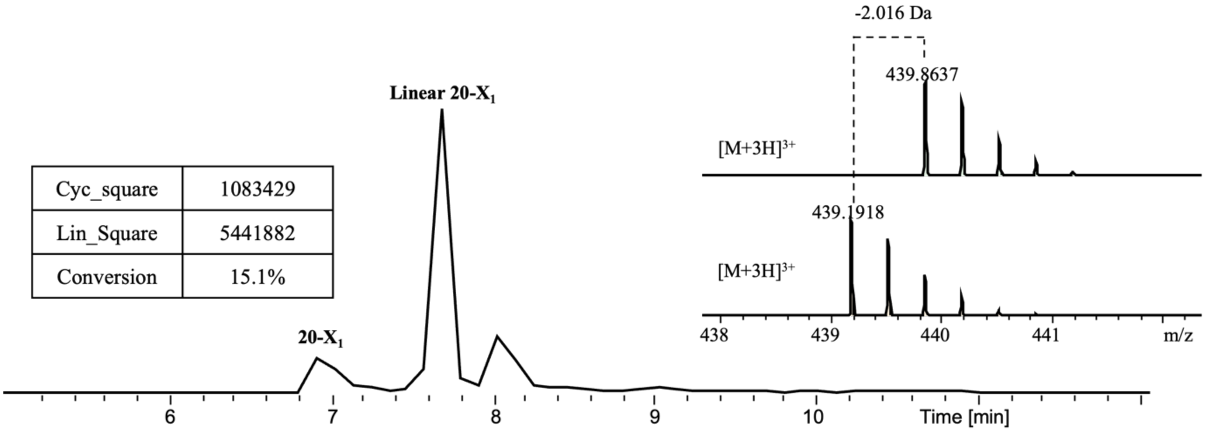


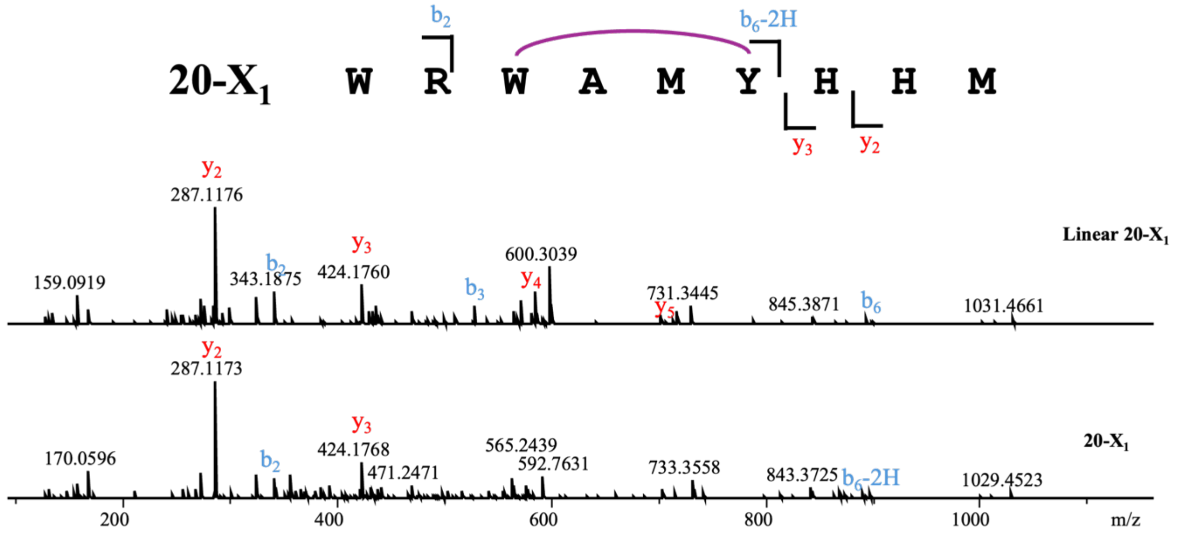


**Linear 20-$X_1$**

| Ions | Fragment | Observed mass | Calculated mass | z | Δppm |
|---|---|---|---|---|---|
| $y_2$ | HM | 287.1176 | 287.1172 | 1 | 1.39 |
| $y_3$ | HHM | 424.1760 | 424.1761 | 1 | 0.24 |
| $y_4$ | YHHM | 587.2395 | 587.2395 | 1 | <0.1 |
| $y_5$ | MYHHM | 718.2798 | 718.28 | 1 | 0.28 |
| $b_2$ | WR | 343.1875 | 343.1877 | 1 | 0.58 |
| $b_3$ | WRW | 529.2669 | 529.267 | 1 | 0.19 |
| $b_6$ | WRWWMY | 894.4076 | 894.4079 | 1 | 0.34 |

**20-$X_1$**

| Ions | Fragment | Observed mass | Calculated mass | z | Δppm |
|---|---|---|---|---|---|
| $y_2$ | HM | 287.1173 | 287.1172 | 1 | 0.35 |
| $y_3$ | HHM | 424.1768 | 424.1761 | 1 | 1.65 |
| $b_2$ | WR | 343.1874 | 343.1877 | 1 | 0.87 |
| $b_6$-2H | WRWAMY | 892.3939 | 892.3923 | 1 | 1.79 |

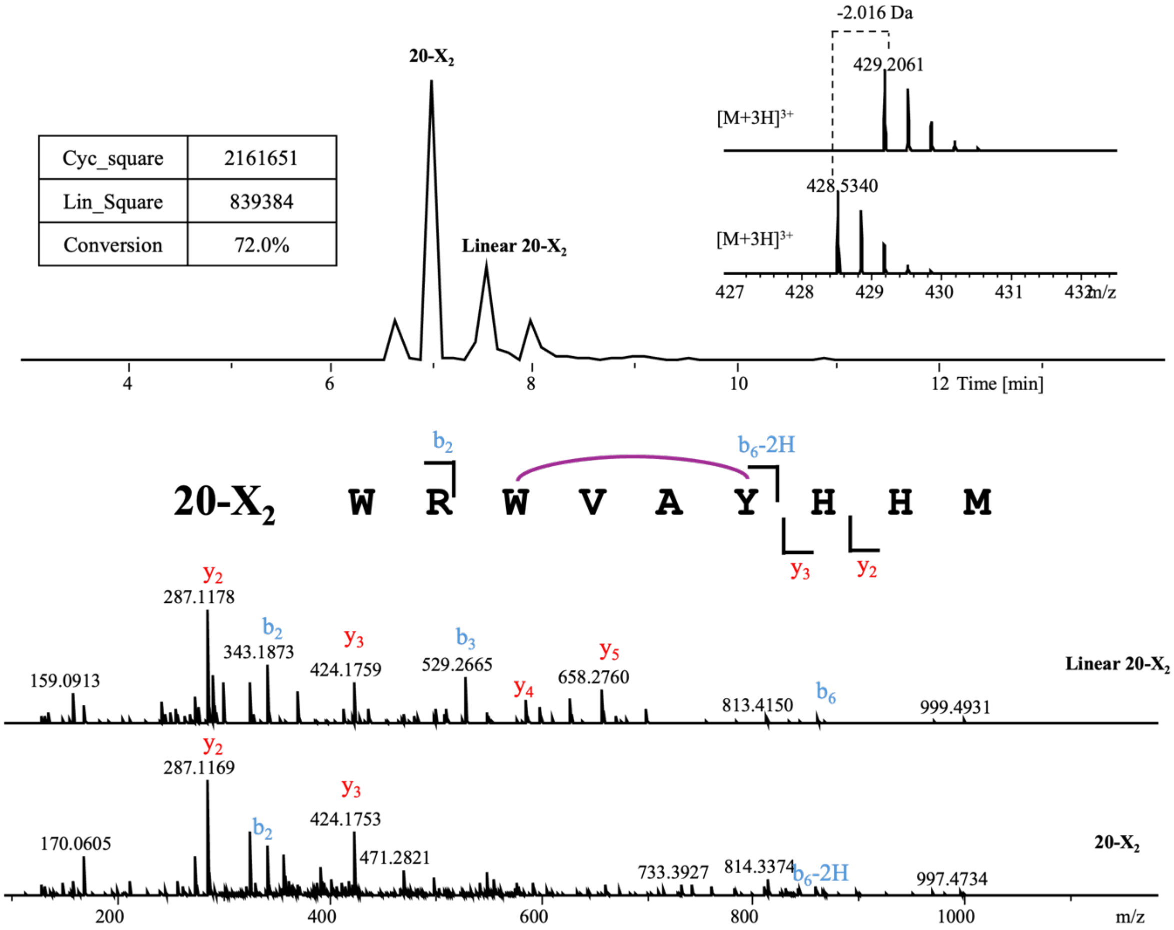


**Linear 20-X$_2$**

| Ions | Fragment | Observed mass | Calculated mass | z | Δppm |
|---|---|---|---|---|---|
| $y_2$ | HM | 287.1178 | 287.1172 | 1 | 2.09 |
| $y_3$ | HHM | 424.1759 | 424.1761 | 1 | 0.47 |
| $y_4$ | YHHM | 587.2391 | 587.2395 | 1 | 0.68 |
| $y_5$ | AYHHM | 658.2760 | 658.2766 | 1 | 0.91 |
| $b_2$ | WR | 343.1873 | 343.1877 | 1 | 1.17 |
| $b_3$ | WRW | 529.2665 | 529.267 | 1 | 0.94 |
| $b_6$ | WRWVAY | 862.4355 | 862.4359 | 1 | 0.46 |

**20-X$_2$**

| Ions | Fragment | Observed mass | Calculated mass | z | Δppm |
|---|---|---|---|---|---|
| $y_2$ | HM | 287.1169 | 287.1172 | 1 | 1.04 |
| $y_3$ | HHM | 424.1753 | 424.1761 | 1 | 1.89 |
| $b_2$ | WR | 343.1869 | 343.1877 | 1 | 2.33 |
| $b_6$-2H | WRWVAY | 860.4159 | 860.4202 | 1 | 5 |

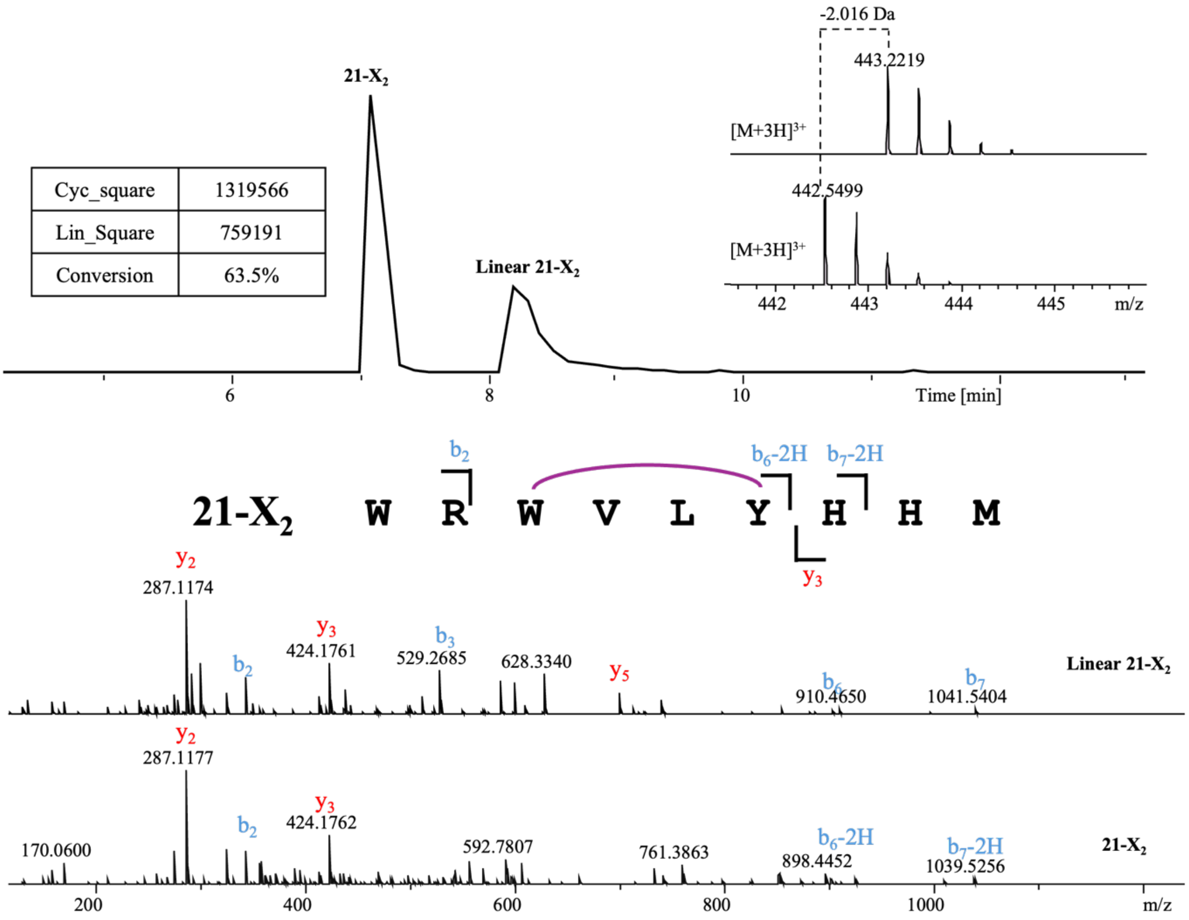


**Linear 21-$X_2$**

| Ions | Fragment | Observed mass | Calculated mass | z | Δppm |
|---|---|---|---|---|---|
| $y_2$ | HM | 287.1174 | 287.1172 | 1 | 0.7 |
| $y_3$ | HHM | 424.1761 | 424.1761 | 1 | <0.1 |
| $y_4$ | YHHM | 587.2387 | 587.2395 | 1 | 1.36 |
| $y_5$ | LYHHM | 700.3269 | 700.3235 | 1 | 4.85 |
| $b_2$ | WR | 343.1865 | 343.1877 | 1 | 3.5 |
| $b_3$ | WRW | 529.2685 | 529.267 | 1 | 2.83 |
| $b_6$ | WRWVLY | 904.4756 | 904.4828 | 1 | 7.96 |
| $b_7$ | WRWVLYH | 1041.5404 | 1041.542 | 1 | 1.54 |

**21-$X_2$**

| Ions | Fragment | Observed mass | Calculated mass | z | Δppm |
|---|---|---|---|---|---|
| $y_3$ | HHM | 424.1762 | 424.1761 | 1 | 0.24 |
| $b_2$ | WR | 343.1870 | 343.1877 | 1 | 2.04 |
| $b_6$-2H | WRWVLY | 902.4668 | 902.4672 | 1 | 0.44 |
| $b_7$-2H | WRWVLYH | 1039.5256 | 1039.526 | 1 | 0.38 |

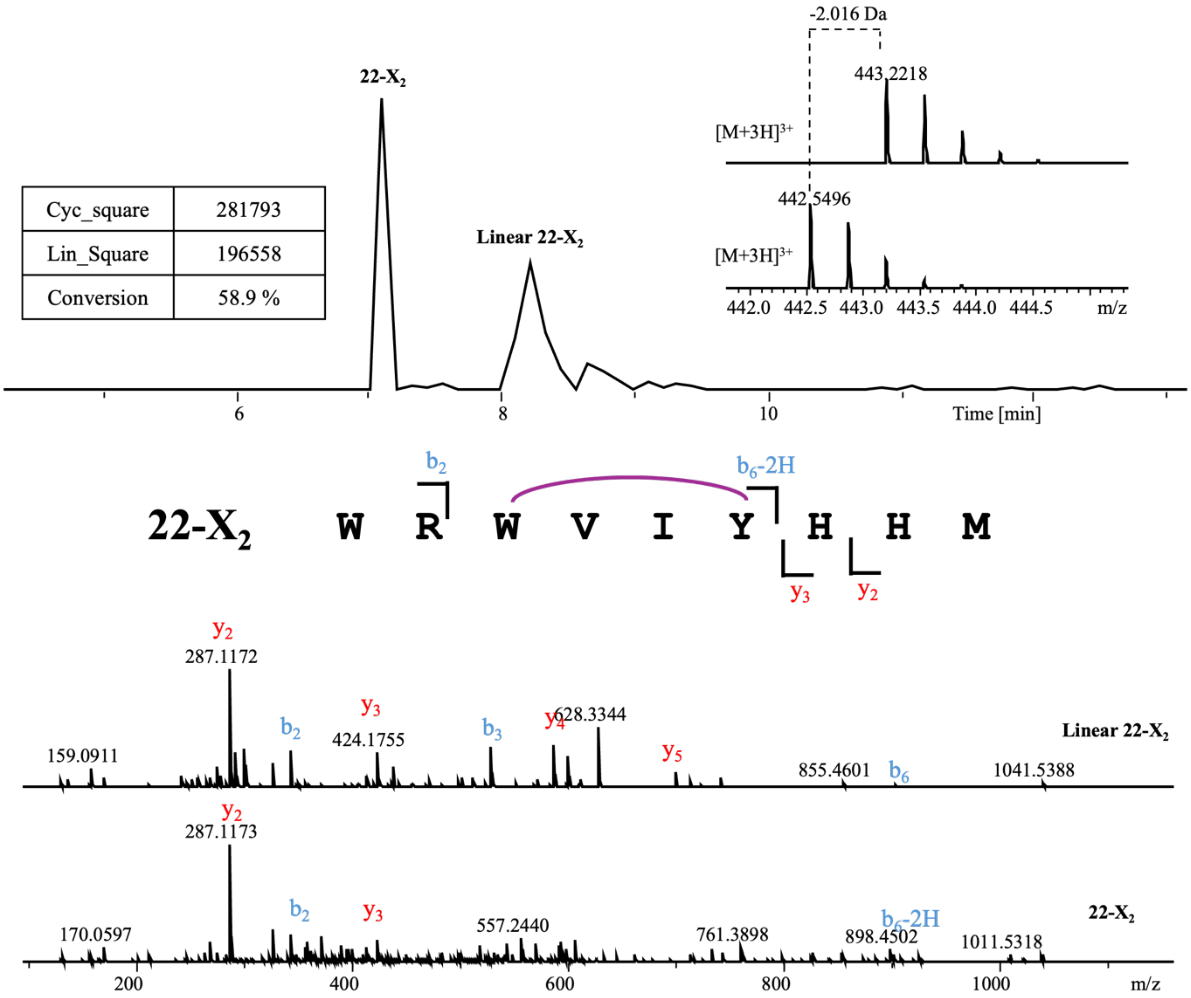


**Linear 22-$X_2$**

| Ions | Fragment | Observed mass | Calculated mass | z | Δppm |
|---|---|---|---|---|---|
| $y_2$ | HM | 287.1172 | 287.1172 | 1 | 0 |
| $y_3$ | HHM | 424.1755 | 424.1761 | 1 | 1.41 |
| $y_4$ | YHHM | 587.2389 | 587.2395 | 1 | 1.02 |
| $y_5$ | IYHHM | 700.3221 | 700.3235 | 1 | 2 |
| $b_2$ | WR | 343.1869 | 343.1877 | 1 | 2.33 |
| $b_3$ | WRW | 529.2662 | 529.267 | 1 | 1.51 |
| $b_6$ | WRWVIY | 904.4813 | 904.4828 | 1 | 1.66 |

**22-$X_2$**

| Ions | Fragment | Observed mass | Calculated mass | z | Δppm |
|---|---|---|---|---|---|
| $y_2$ | HM | 287.1173 | 287.1172 | 1 | 0.35 |
| $y_3$ | HHM | 424.1764 | 424.1761 | 1 | 0.71 |
| $b_2$ | WR | 343.1850 | 343.1877 | 1 | 7.87 |
| $b_6$-2H | WRWVIY | 902.4597 | 902.4672 | 1 | 8.31 |

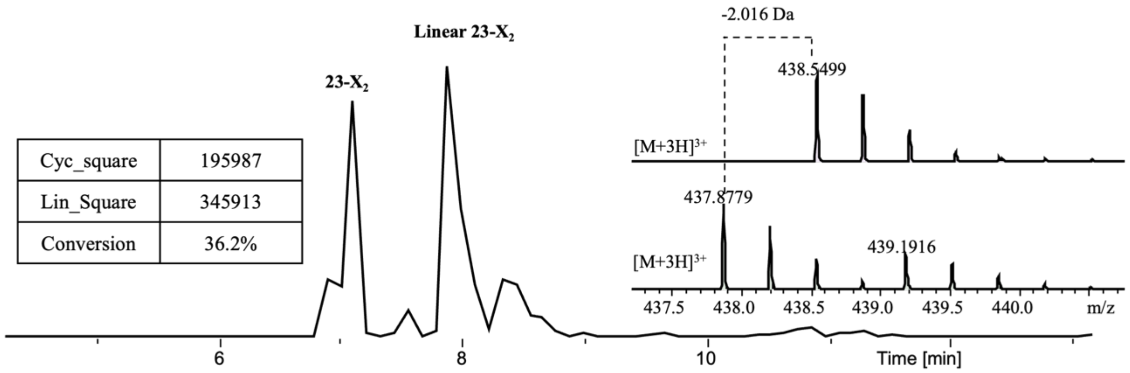


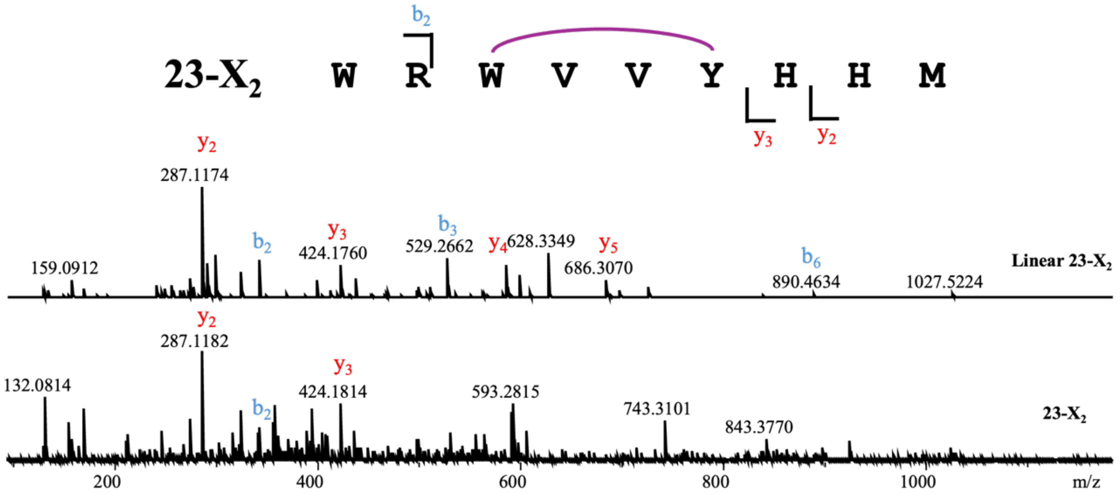


**Linear 23-X$_2$**

| Ions | Fragment | Observed mass | Calculated mass | z | Δppm |
|---|---|---|---|---|---|
| $y_2$ | HM | 287.1174 | 287.1172 | 1 | 0.7 |
| $y_3$ | HHM | 424.1760 | 424.1761 | 1 | 0.24 |
| $y_4$ | YHHM | 587.2387 | 587.2395 | 1 | 1.36 |
| $y_5$ | VYHHM | 686.3070 | 686.3079 | 1 | 1.31 |
| $b_2$ | WR | 343.1870 | 343.1877 | 1 | 2.04 |
| $b_3$ | WRW | 529.2662 | 529.267 | 1 | 1.51 |
| $b_6$ | WRWVVY | 890.4634 | 890.4672 | 1 | 4.27 |

**23-X$_2$**

| Ions | Fragment | Observed mass | Calculated mass | z | Δppm |
|---|---|---|---|---|---|
| $y_2$ | HM | 287.1183 | 287.1172 | 1 | 3.83 |
| $y_3$ | HHM | 424.1759 | 424.1761 | 1 | 0.47 |
| $b_2$ | WR | 343.1871 | 343.1877 | 1 | 1.75 |
| $b_6$-2H | WRWVVY | - | 888.4515 | 1 | - |

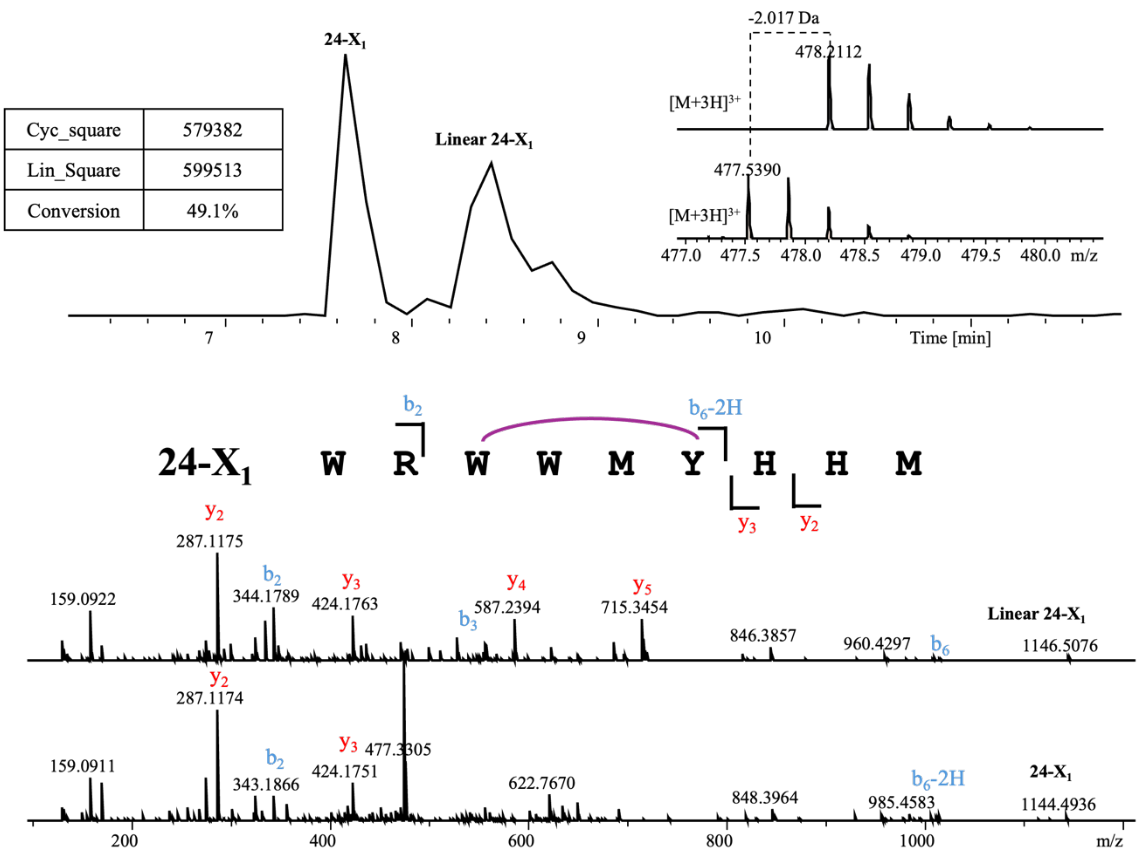


**Linear 24-X$_1$**

| Ions | Fragment | Observed mass | Calculated mass | z | Δppm |
|---|---|---|---|---|---|
| $y_2$ | HM | 287.1175 | 287.1172 | 1 | 1.04 |
| $y_3$ | HHM | 424.1763 | 424.1761 | 1 | 0.47 |
| $y_4$ | YHHM | 587.2394 | 587.2395 | 1 | 0.17 |
| $y_5$ | MYHHM | 718.2779 | 718.28 | 1 | 2.92 |
| $b_2$ | WR | 343.1867 | 343.1877 | 1 | 2.91 |
| $b_3$ | WRW | 529.2668 | 529.267 | 1 | 0.38 |
| $b_6$ | WRWWMY | 1009.4572 | 1009.45 | 1 | 7.13 |

**24-X$_1$**

| Ions | Fragment | Observed mass | Calculated mass | z | Δppm |
|---|---|---|---|---|---|
| $y_2$ | HM | 287.1174 | 287.1172 | 1 | 0.7 |
| $y_3$ | HHM | 424.1751 | 424.1761 | 1 | 2.36 |
| $b_2$ | WR | 343.1866 | 343.1877 | 1 | 3.21 |
| $b_6$-2H | WRWWMY | 1007.4321 | 1007.4345 | 1 | 2.38 |

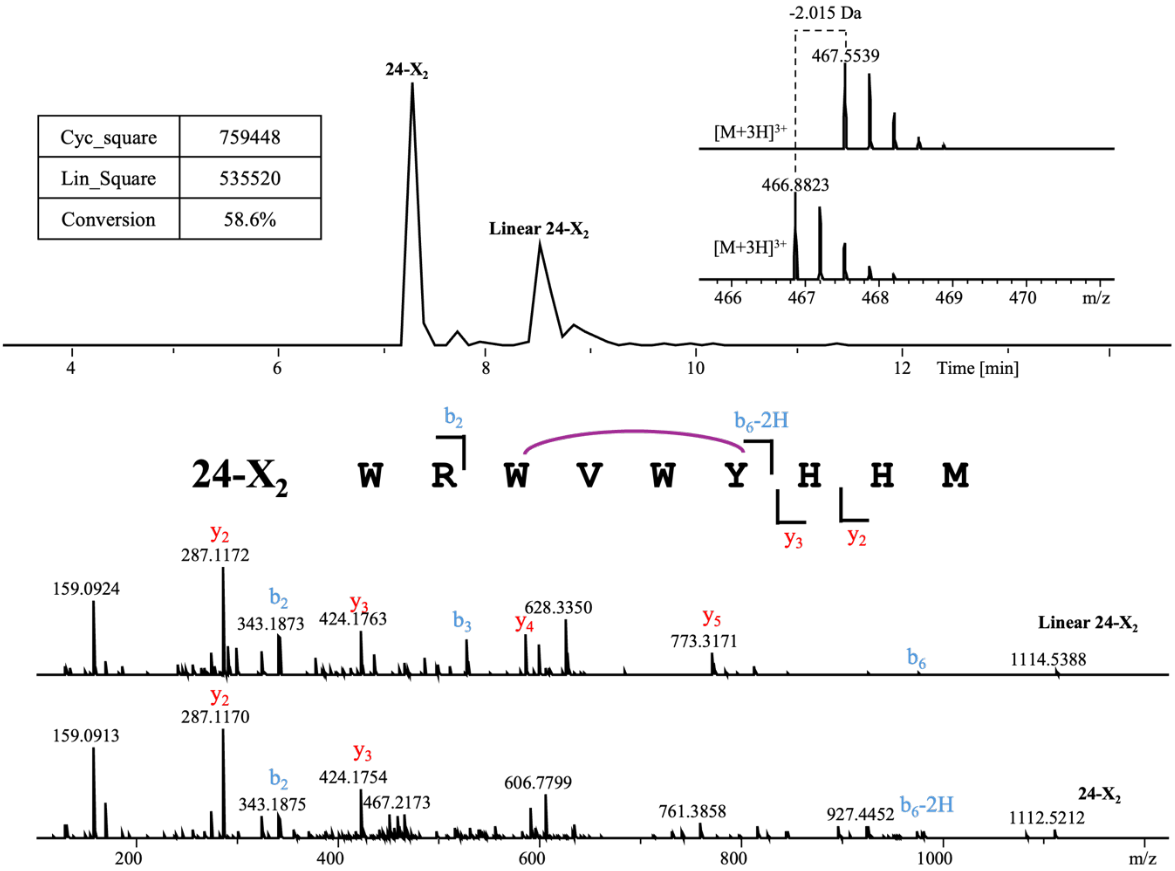


**Linear 24-$X_2$**

| Ions | Fragment | Observed mass | Calculated mass | z | Δppm |
|---|---|---|---|---|---|
| $y_2$ | HM | 287.1172 | 287.1172 | 1 | <0.1 |
| $y_3$ | HHM | 424.1763 | 424.1761 | 1 | 0.47 |
| $y_4$ | YHHM | 587.2393 | 587.2395 | 1 | 0.34 |
| $y_5$ | WYHHM | 773.3171 | 773.3188 | 1 | 2.2 |
| $b_2$ | WR | 343.1873 | 343.1877 | 1 | 1.17 |
| $b_3$ | WRW | 529.267 | 529.267 | 1 | <0.1 |
| $b_6$ | WRWVWY | 977.4777 | 977.4781 | 1 | 0.41 |

**24-$X_2$**

| Ions | Fragment | Observed mass | Calculated mass | z | Δppm |
|---|---|---|---|---|---|
| $y_2$ | HM | 287.1170 | 287.1172 | 1 | 0.7 |
| $y_3$ | HHM | 424.1754 | 424.1761 | 1 | 1.65 |
| $b_2$ | WR | 343.1875 | 343.1877 | 1 | 0.58 |
| $b_6$-2H | WRWVWY | 975.4634 | 975.4624 | 1 | 1.03 |

| | |
|---|---|
| Cyc_square | 1499384 |
| Lin_Square | 922402 |
| Conversion | 61.9% |

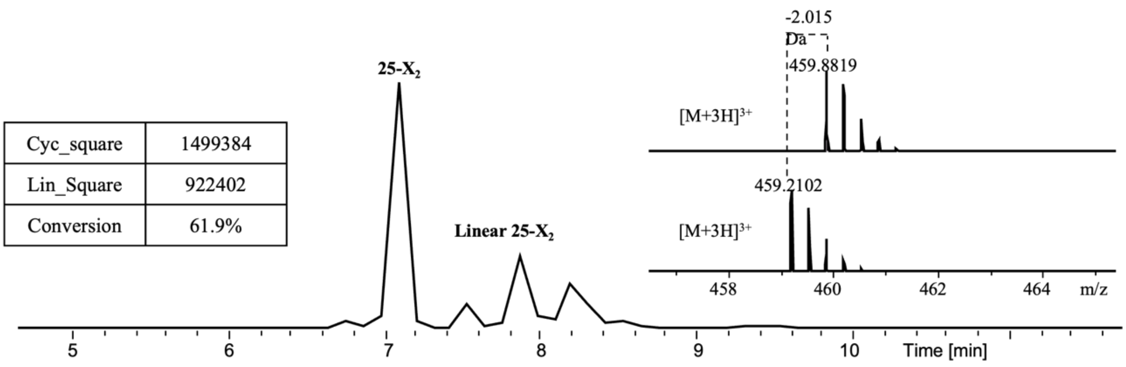


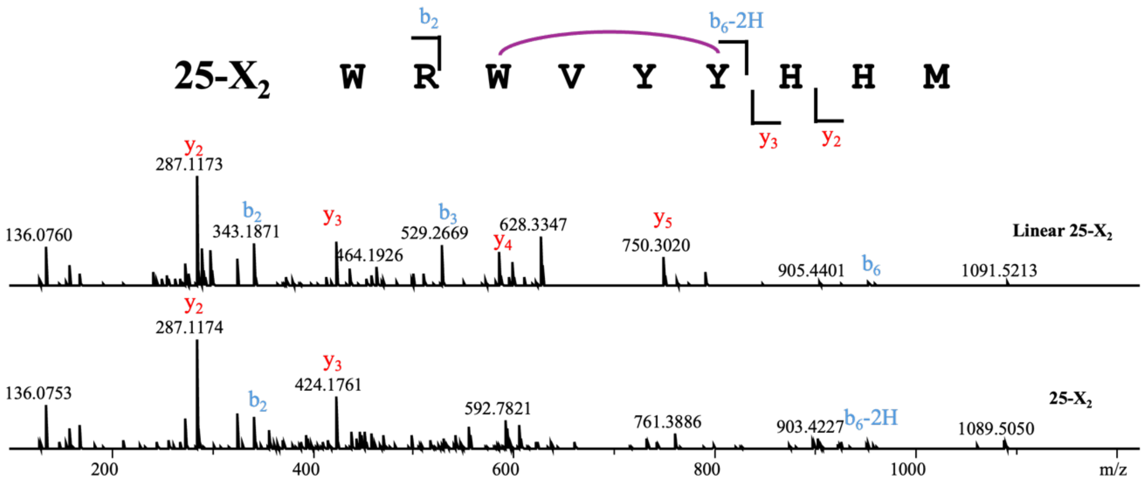


**Linear 25-$X_2$**

| Ions | Fragment | Observed mass | Calculated mass | z | Δppm |
|---|---|---|---|---|---|
| $y_2$ | HM | 287.1173 | 287.1172 | 1 | 0.35 |
| $y_3$ | HHM | 424.1760 | 424.1761 | 1 | 0.24 |
| $y_4$ | YHHM | 587.2391 | 587.2395 | 1 | 0.68 |
| $y_5$ | YYHHM | 750.3020 | 750.3028 | 1 | 1.07 |
| $b_2$ | WR | 343.1871 | 343.1877 | 1 | 1.75 |
| $b_3$ | WRW | 529.2669 | 529.267 | 1 | 0.19 |
| $b_6$ | WRWVYY | 954.4605 | 954.4621 | 1 | 1.68 |

**25-$X_2$**

| Ions | Fragment | Observed mass | Calculated mass | z | Δppm |
|---|---|---|---|---|---|
| $y_2$ | HM | 287.1174 | 287.1172 | 1 | 0.7 |
| $y_3$ | HHM | 424.1761 | 424.1761 | 1 | <0.1 |
| $b_2$ | WR | 343.1872 | 343.1877 | 1 | 1.46 |
| $b_6$-2H | WRWVYY | 952.4419 | 952.4464 | 1 | 4.72 |

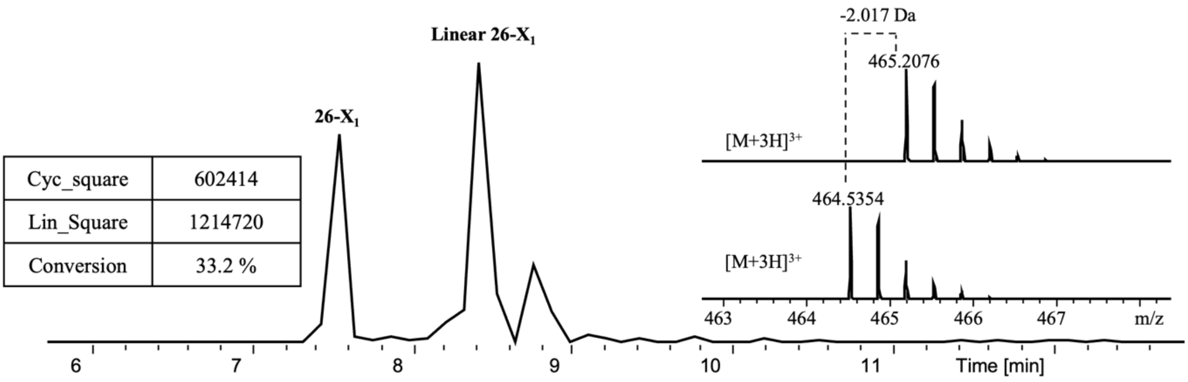


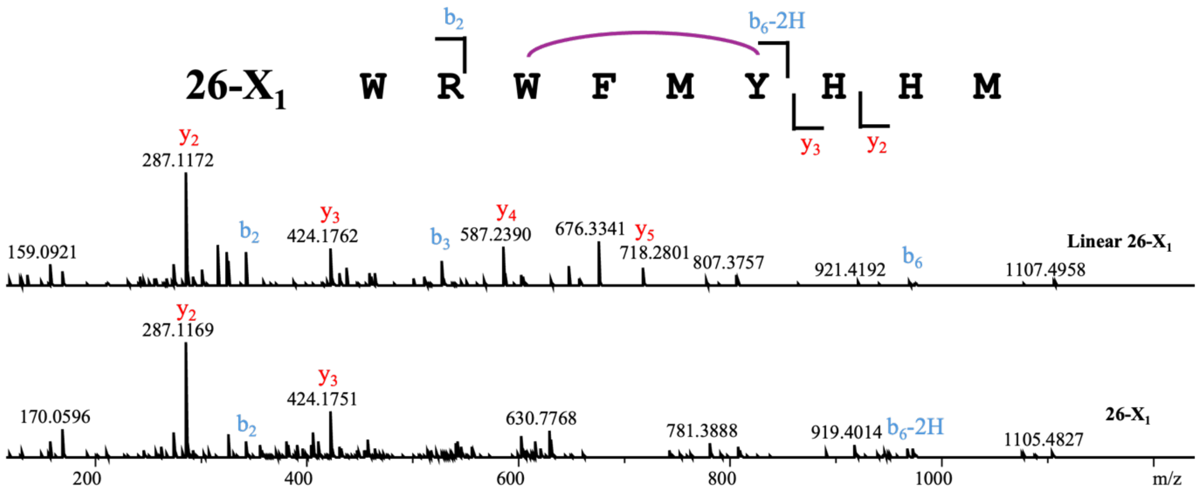


**Linear 26-$X_1$**

| Ions | Fragment | Observed mass | Calculated mass | z | Δppm |
|---|---|---|---|---|---|
| $y_2$ | HM | 287.1172 | 287.1172 | 1 | <0.1 |
| $y_3$ | HHM | 424.1762 | 424.1761 | 1 | 0.24 |
| $y_4$ | YHHM | 587.2390 | 587.2395 | 1 | 0.85 |
| $y_5$ | MYHHM | 718.2801 | 718.28 | 1 | 0.14 |
| $b_2$ | WR | 343.1875 | 343.1877 | 1 | 0.58 |
| $b_3$ | WRW | 529.2667 | 529.267 | 1 | 0.57 |
| $b_6$ | WRWFMY | 970.4369 | 970.4392 | 1 | 2.37 |

**26-$X_1$**

| Ions | Fragment | Observed mass | Calculated mass | z | Δppm |
|---|---|---|---|---|---|
| $y_2$ | HM | 287.1169 | 287.1172 | 1 | 1.04 |
| $y_3$ | HHM | 424.1751 | 424.1761 | 1 | 2.36 |
| $b_2$ | WR | 343.1890 | 343.1877 | 1 | 3.79 |
| $b_6$-2H | WRWFMY | 968.4268 | 968.4236 | 1 | 3.3 |

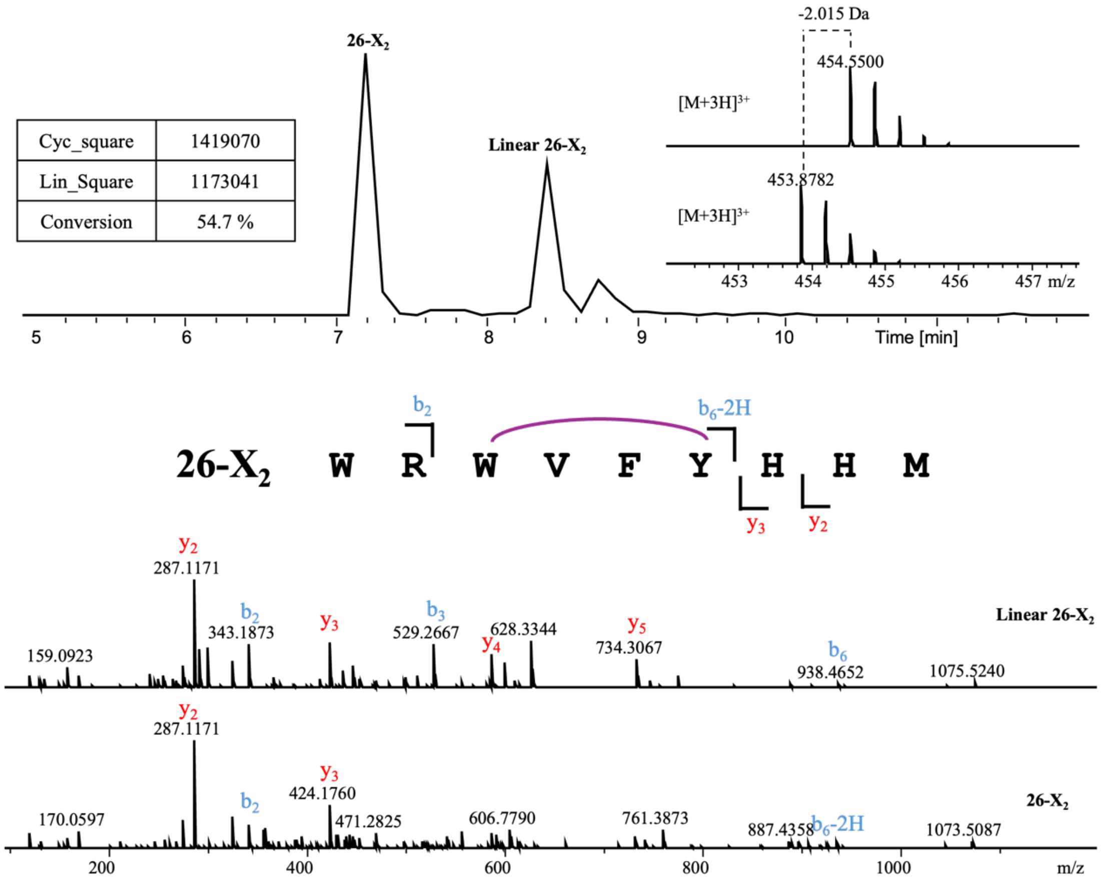


**Linear 26-$X_2$**

| Ions | Fragment | Observed mass | Calculated mass | z | Δppm |
|---|---|---|---|---|---|
| $y_2$ | HM | 287.1171 | 287.1172 | 1 | 0.35 |
| $y_3$ | HHM | 424.1757 | 424.1761 | 1 | 0.94 |
| $y_4$ | YHHM | 587.2388 | 587.2395 | 1 | 1.19 |
| $y_5$ | FYHHM | 734.3067 | 734.3079 | 1 | 1.63 |
| $b_2$ | WR | 343.1873 | 343.1877 | 1 | 1.17 |
| $b_3$ | WRW | 529.2667 | 529.267 | 1 | 0.57 |
| $b_6$ | WRWVFY | 938.4652 | 938.4672 | 1 | 2.13 |

**26-$X_2$**

| Ions | Fragment | Observed mass | Calculated mass | z | Δppm |
|---|---|---|---|---|---|
| $y_2$ | HM | 287.1171 | 287.1172 | 1 | 0.35 |
| $y_3$ | HHM | 424.1760 | 424.1761 | 1 | 0.24 |
| $b_2$ | WR | 343.1876 | 343.1877 | 1 | 0.29 |
| $b_6$-2H | WRWVFY | 936.4508 | 936.4515 | 1 | 0.75 |

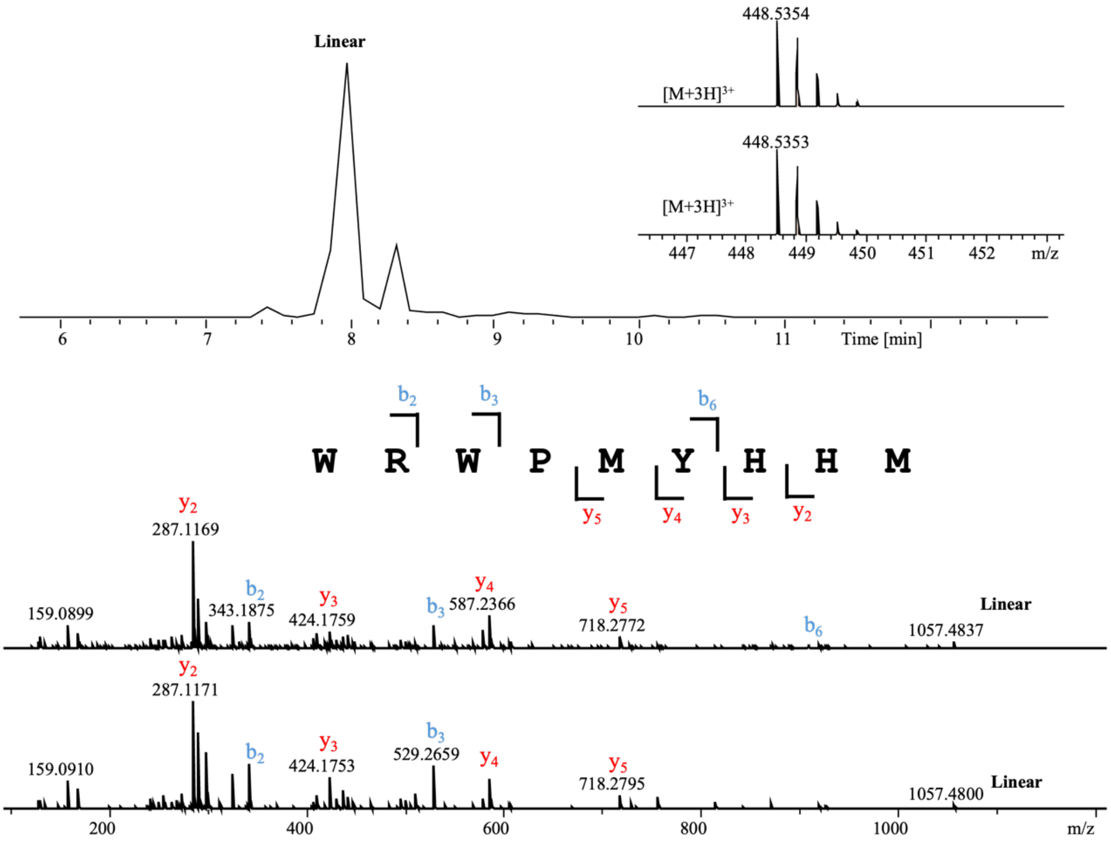


| Linear | | | | | |
|---|---|---|---|---|---|
| Ions | Fragment | Observed mass | Calculated mass | z | Δppm |
| $y_2$ | HM | 287.1169/287.1171 | 287.1172 | 1 | 1.04/0.34 |
| $y_3$ | HHM | 424.1759/424.1753 | 424.1761 | 1 | 0.47/1.89 |
| $y_4$ | YHHM | 587.2366/587.2390 | 587.2395 | 1 | 4.94/0.85 |
| $y_5$ | MYHHM | 718.2772/718.2795 | 718.28 | 1 | 3.89/0.70 |
| $b_2$ | WR | 343.1875/343.1872 | 343.1877 | 1 | 0.58/1.46 |
| $b_3$ | WRW | 529.2704/529.2659 | 529.267 | 1 | 6.42/2.08 |
| $b_6$ | WRWPMY | -/920.4235 | 920.4236 | 1 | -/0.1 |

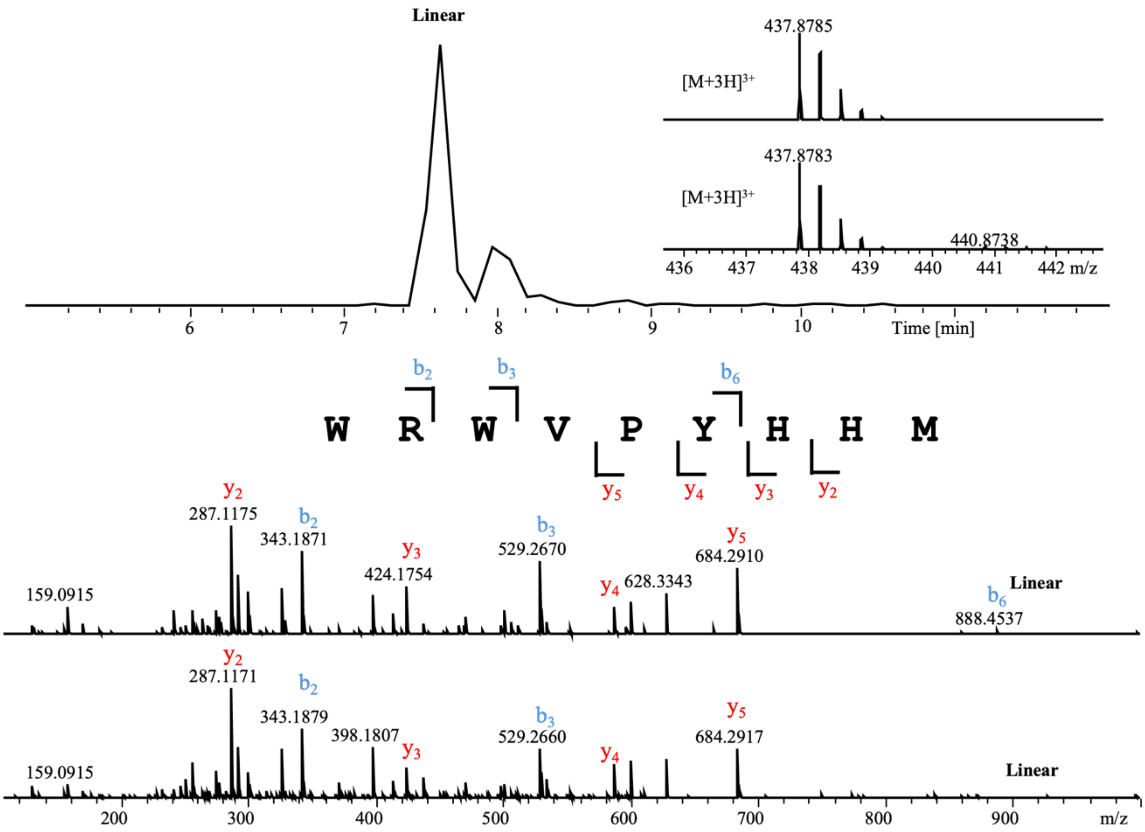


**Linear**

| Ions | Fragment | Observed mass | Calculated mass | z | Δppm |
|---|---|---|---|---|---|
| $y_2$ | HM | 287.1175/287.1171 | 287.1172 | 1 | 1.04/0.35 |
| $y_3$ | HHM | 424.1754/424.1785 | 424.1761 | 1 | 1.65/5.65 |
| $y_4$ | YHHM | 587.2396/587.2370 | 587.2395 | 1 | 0.17/4.25 |
| $y_5$ | PYHHM | 684.2910/684.2917 | 684.2922 | 1 | 1.75/0.73 |
| $b_2$ | WR | 343.1871/343.1879 | 343.1877 | 1 | 1.74/0.58 |
| $b_3$ | WRW | 529.2670/529.2660 | 529.267 | 1 | <0.1/1.88 |
| $b_6$ | WRWVPY | 888.4531/- | 888.4515 | 1 | 1.80/- |

#### NMR spectra

$^{1}$H NMR of **3** (600 MHz, DMSO-*d*$^{6}$):

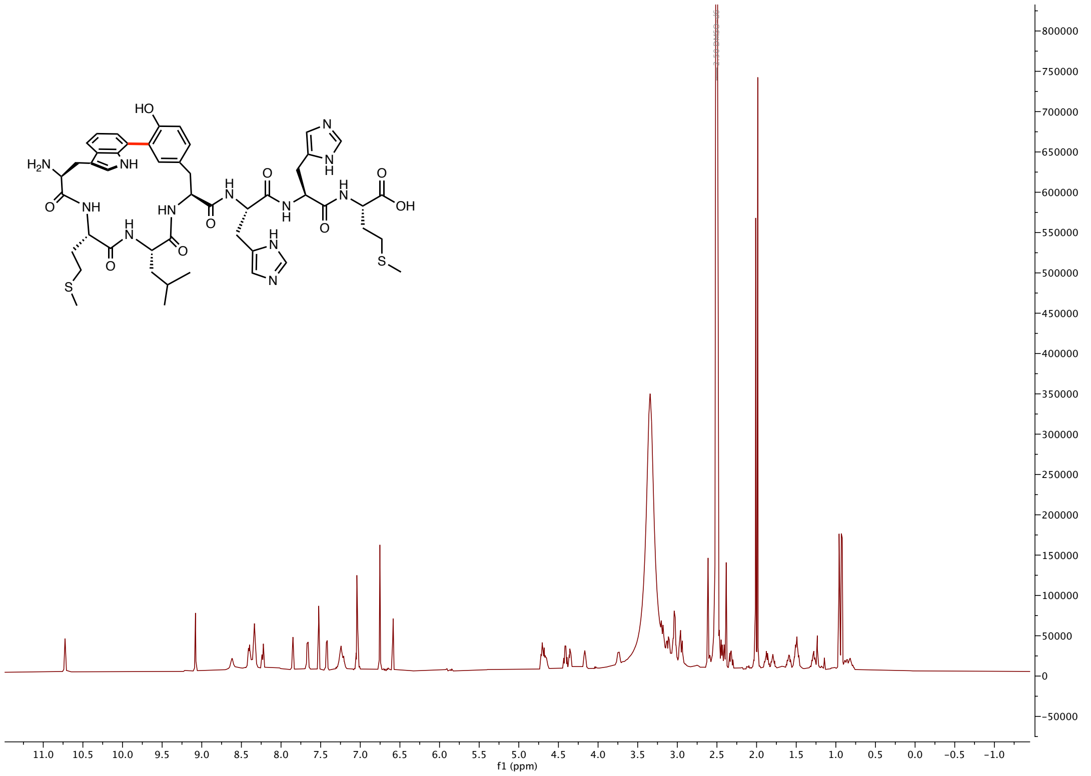


$^{13}$C NMR of **3** (150 MHz, DMSO-*d*$^{6}$):

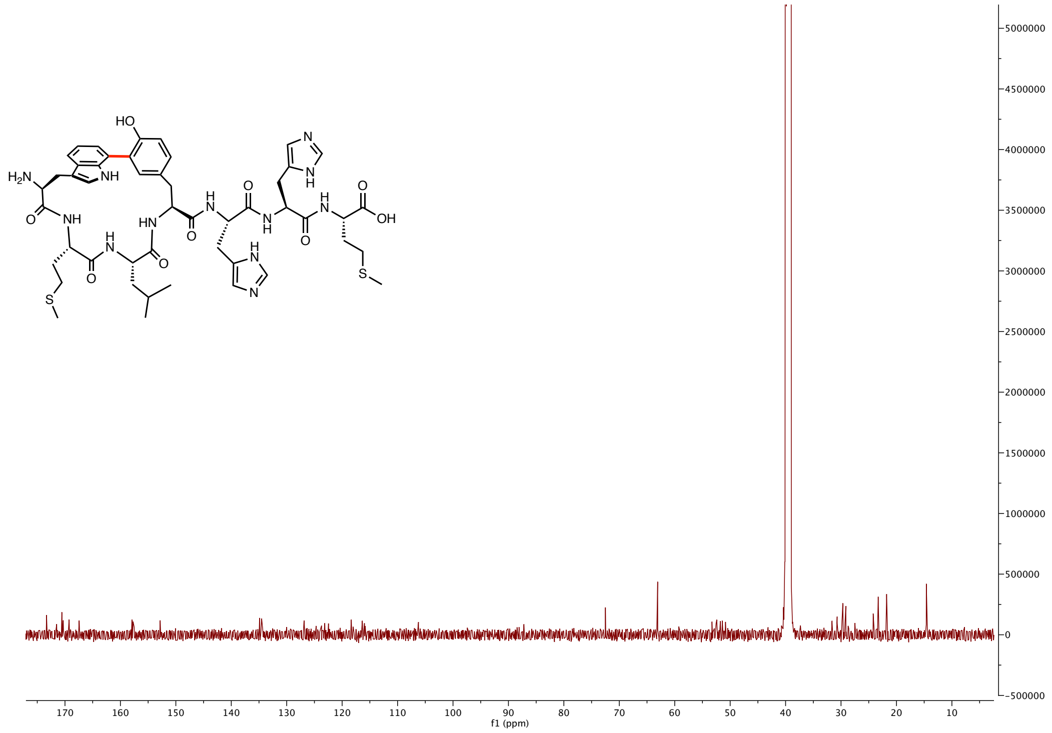

H-H cosy NMR of **3** (600 MHz, DMSO-$d^6$):

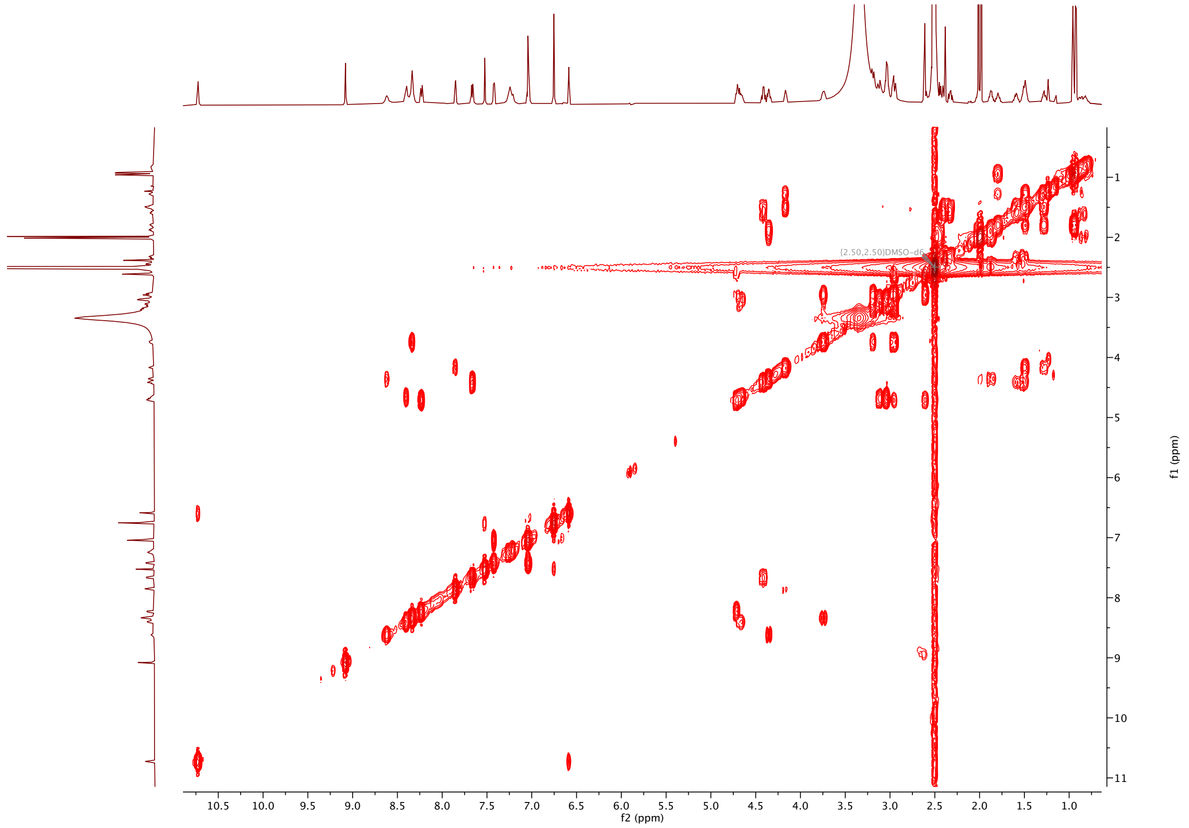


NOESY of **3** (600 MHz, DMSO-$d^6$):

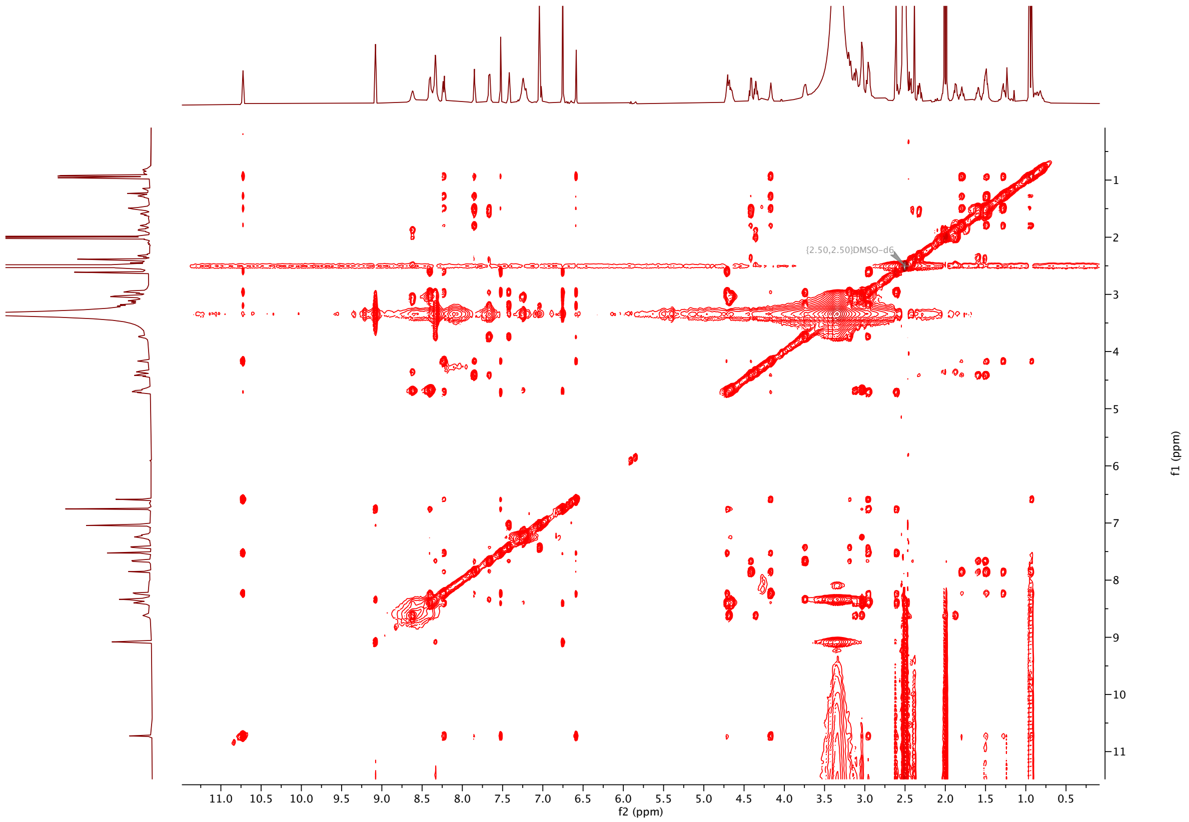

HSQC of **3** (600 MHz, DMSO-$d^6$):

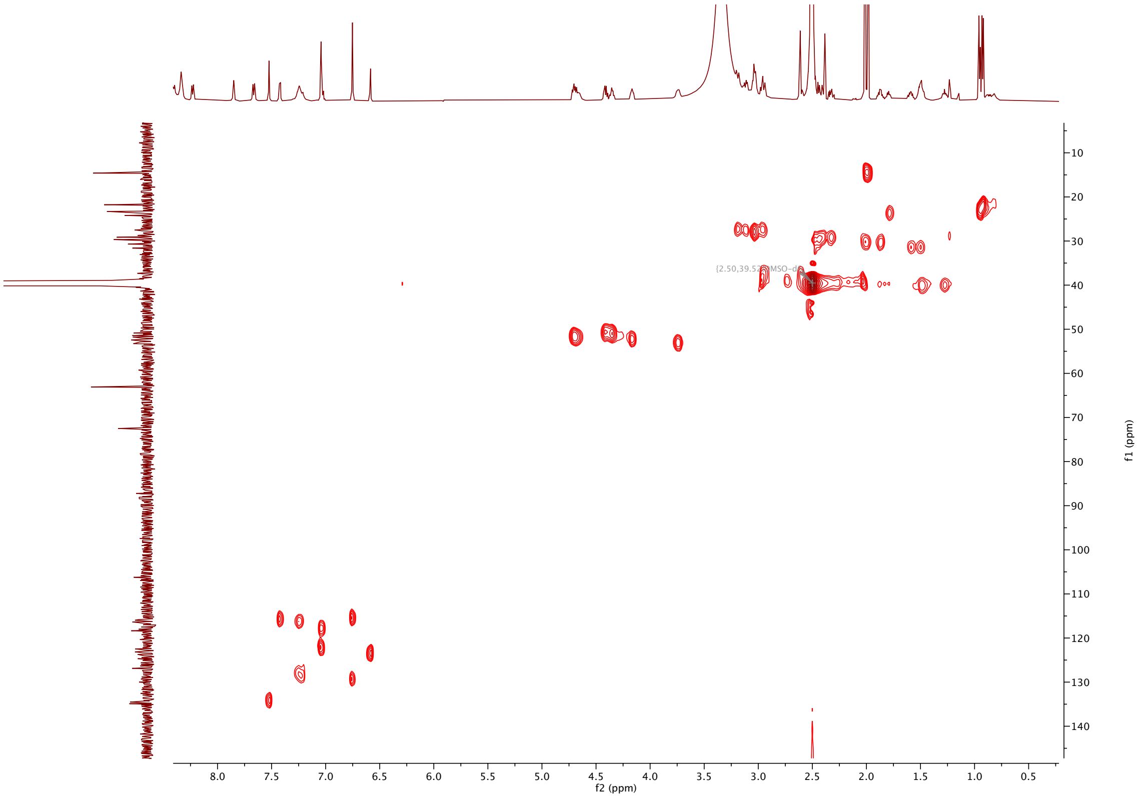


HMBC of **3** (600 MHz, DMSO-$d^6$):

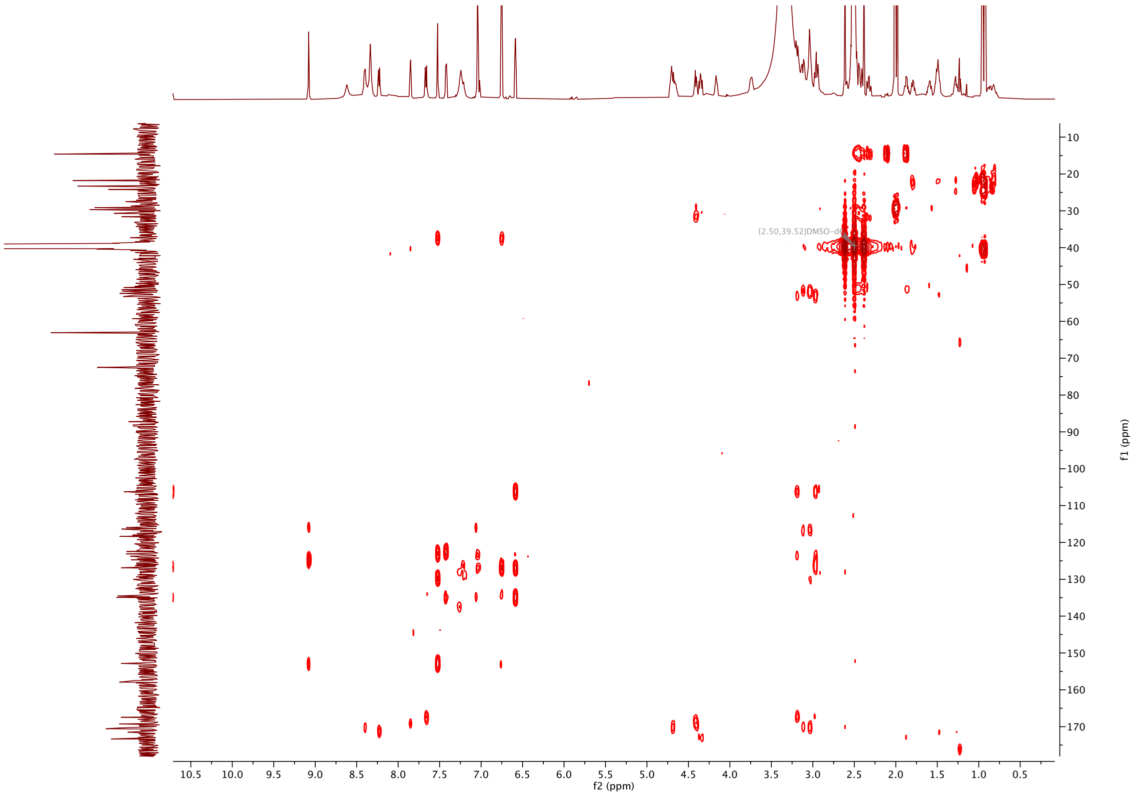

$^{1}$H NMR of **7** (600 MHz, DMSO-$d^6$):

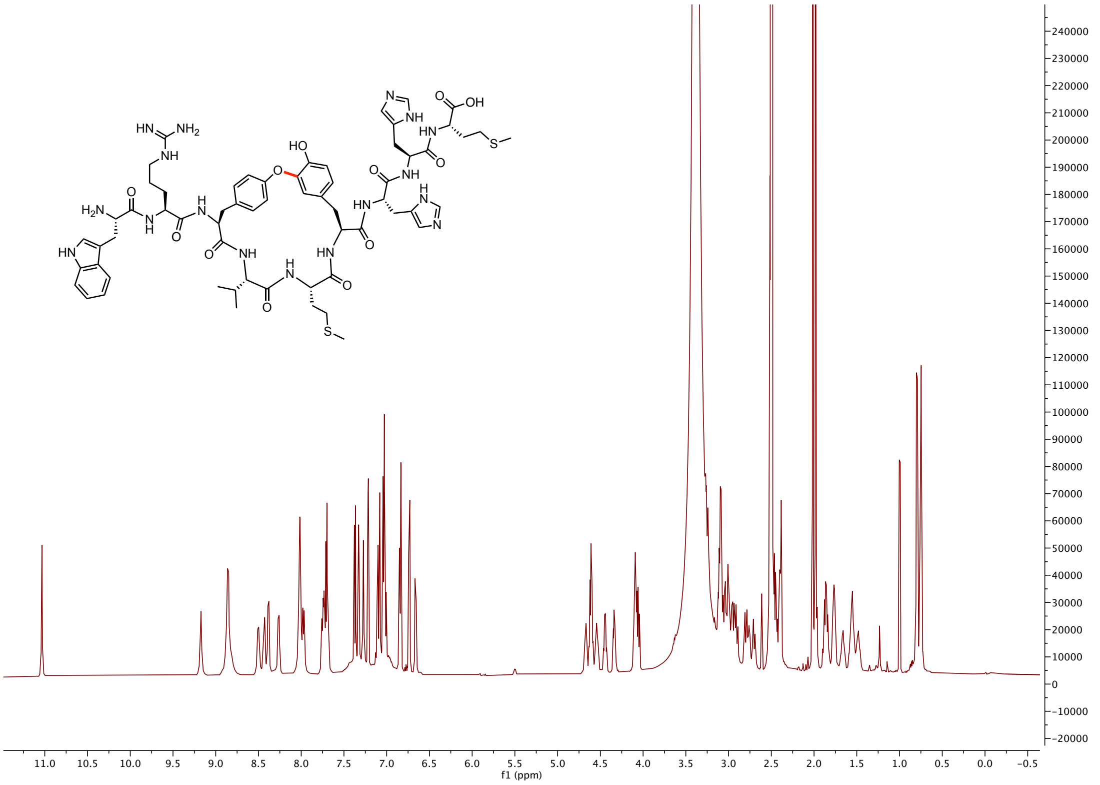

$^{13}$C NMR of **7** (150 MHz, DMSO-$d^6$):

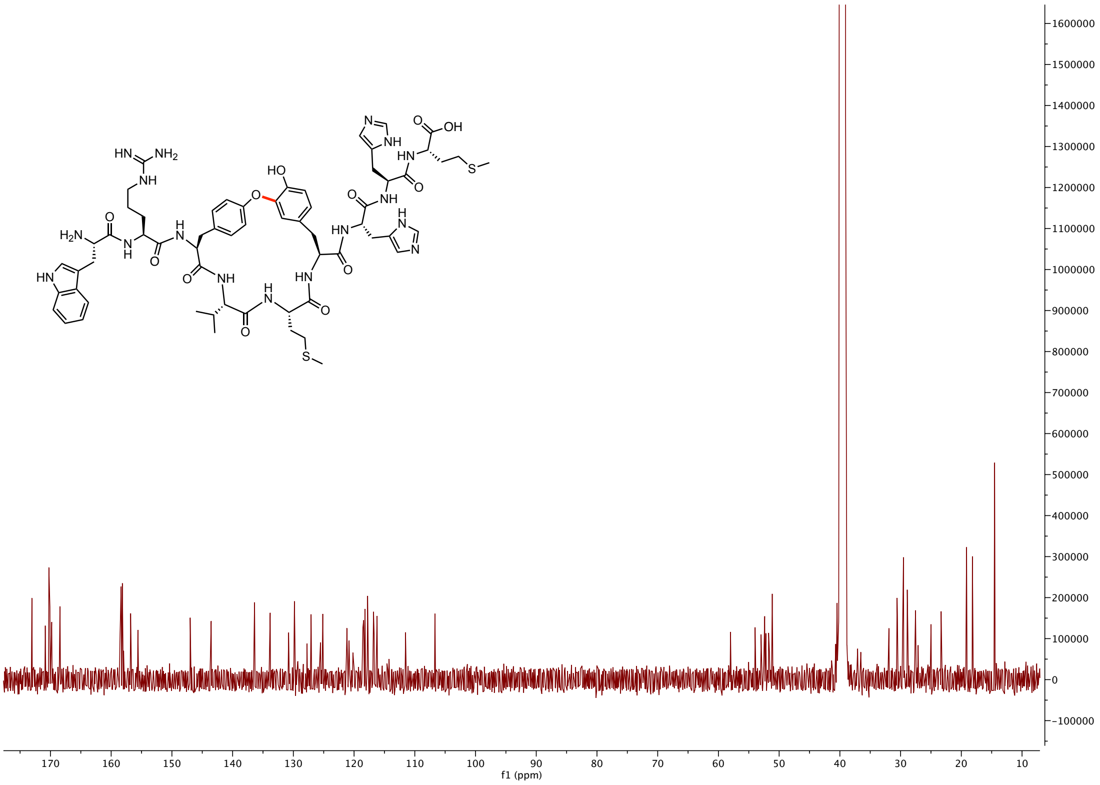

H-H cosy of **7** (600 MHz, DMSO-$d^6$):

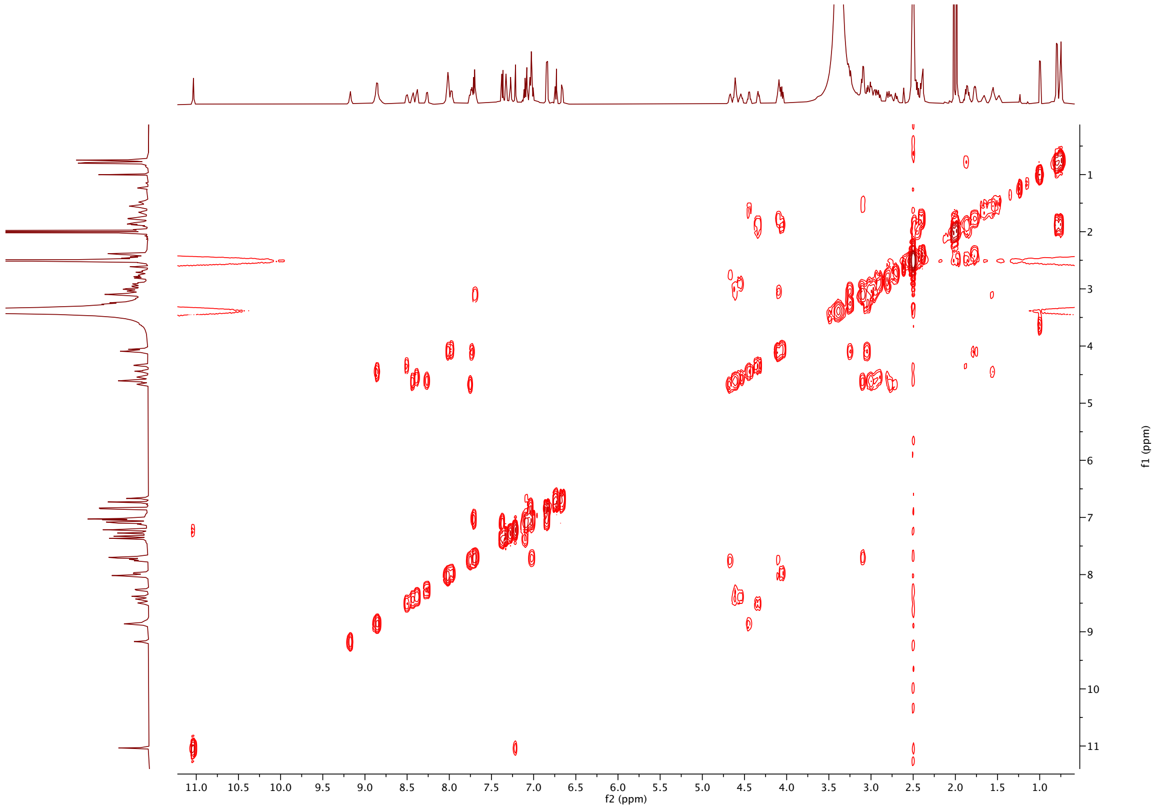


NOESY of **7** (600 MHz, DMSO-$d^6$):

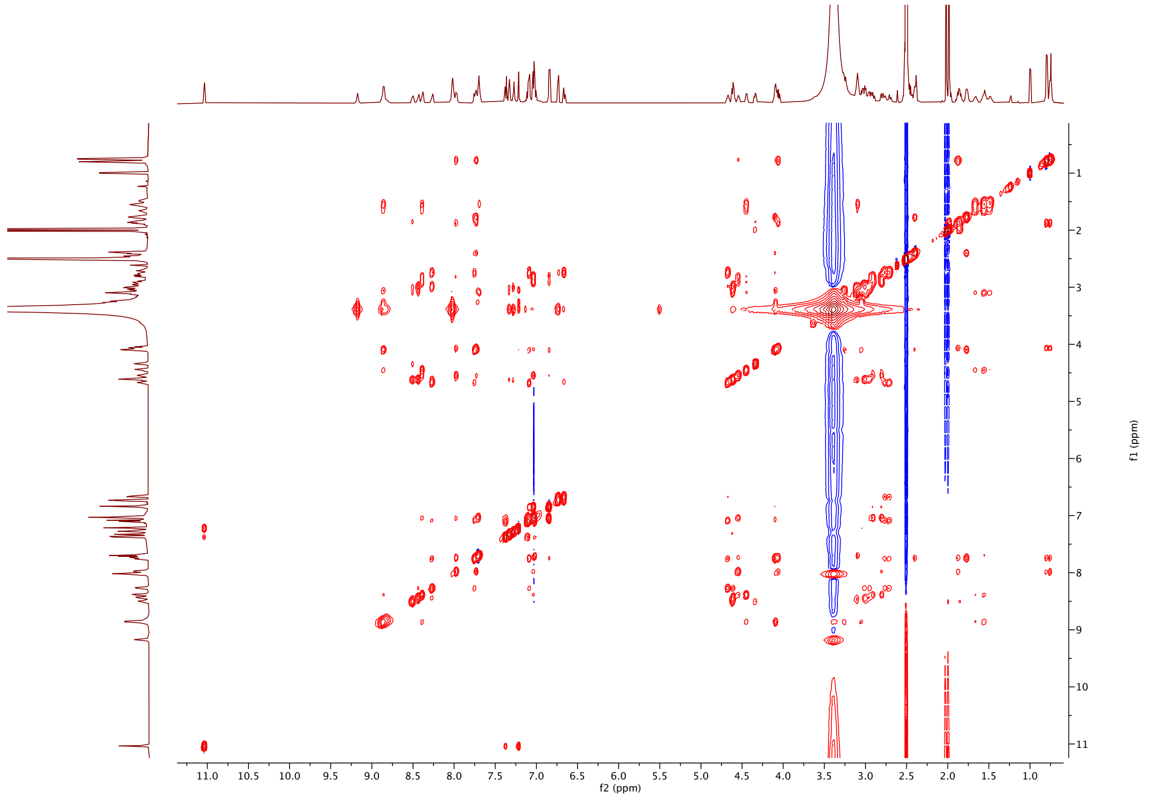

HSQC of **7** (600 MHz, DMSO-$d^6$):

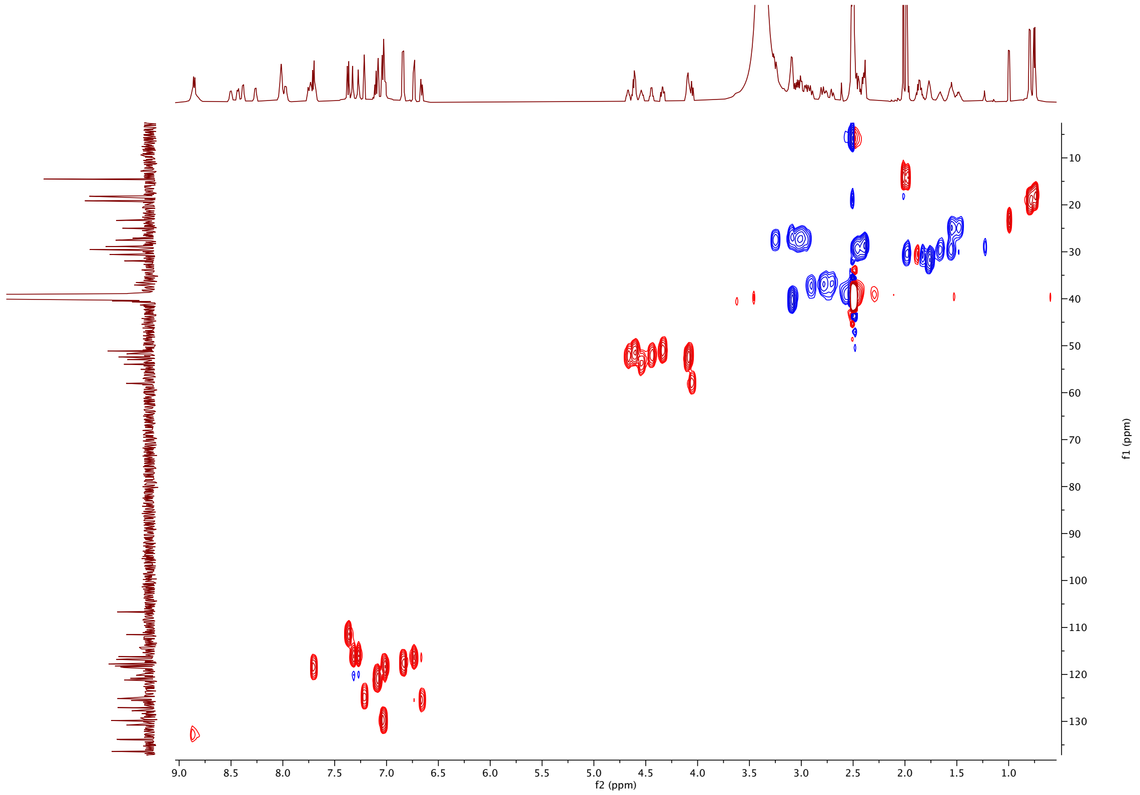


HMBC of **7** (600 MHz, DMSO-$d^6$):

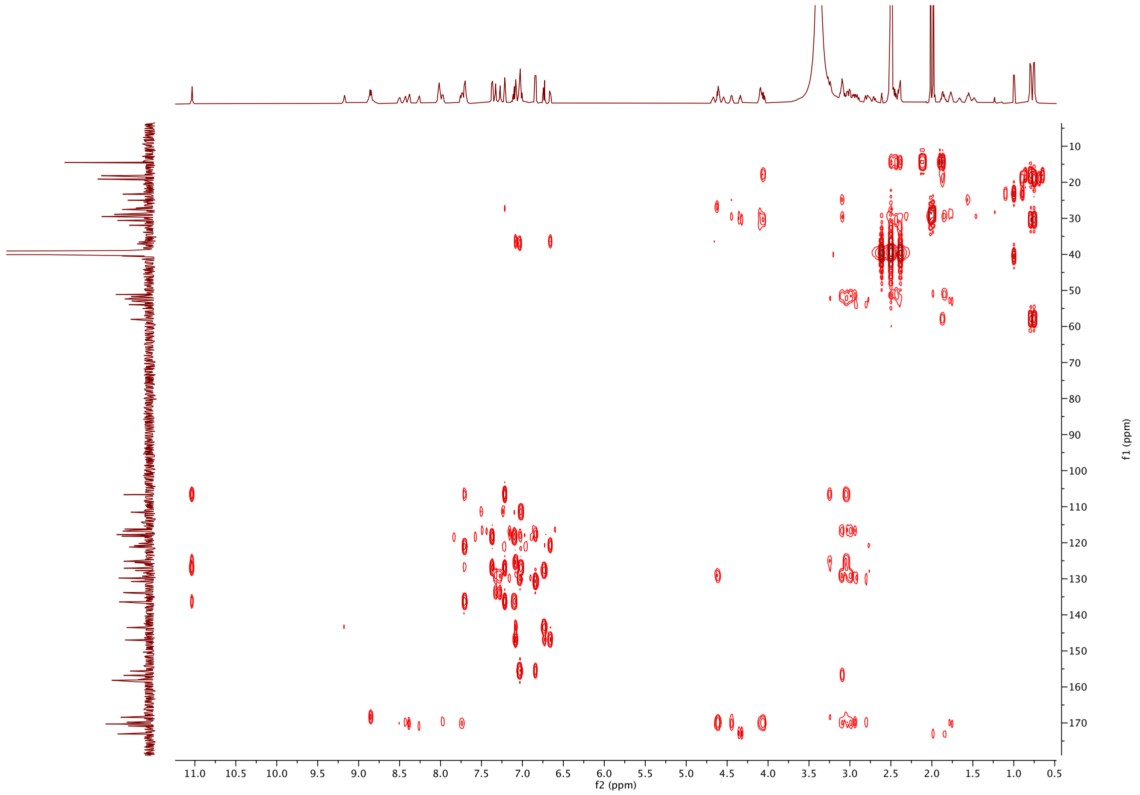

$^{1}H$ NMR of **8** (600 MHz, DMSO-$d^{6}$):

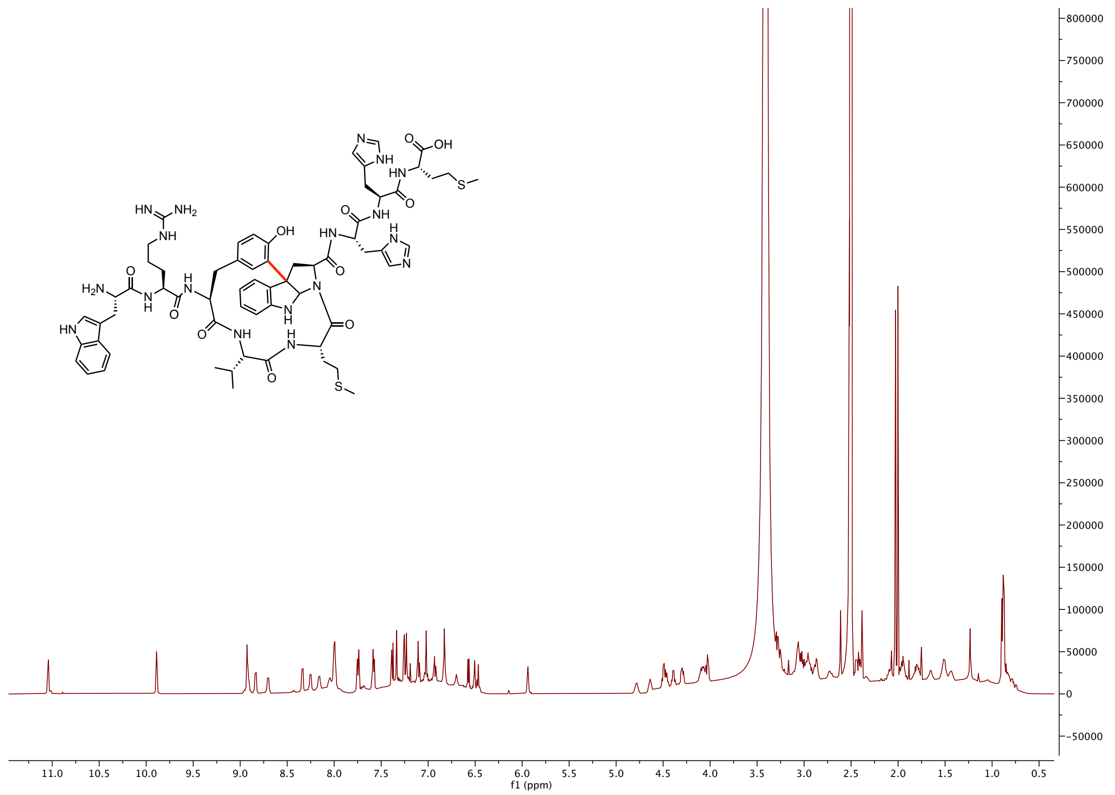


$^{13}C$ NMR of **8** (150 MHz, DMSO-$d^{6}$):

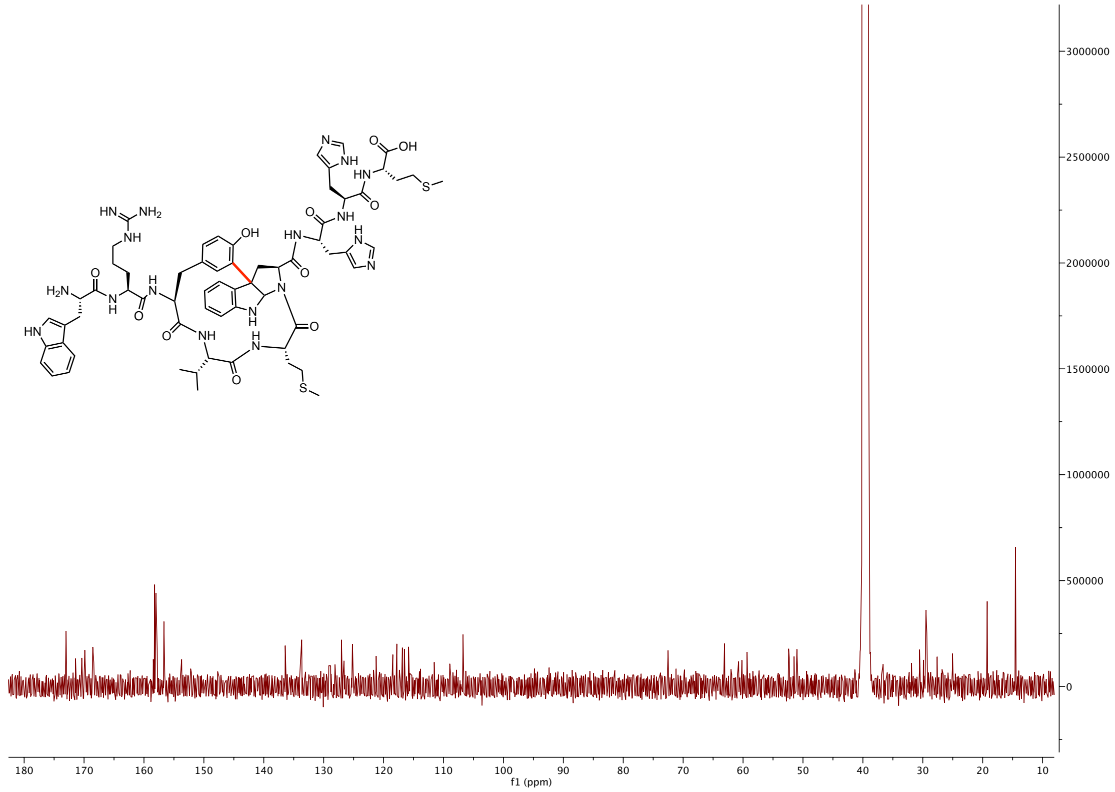

H-H COSY of **8** (600 MHz, DMSO-$d^6$):

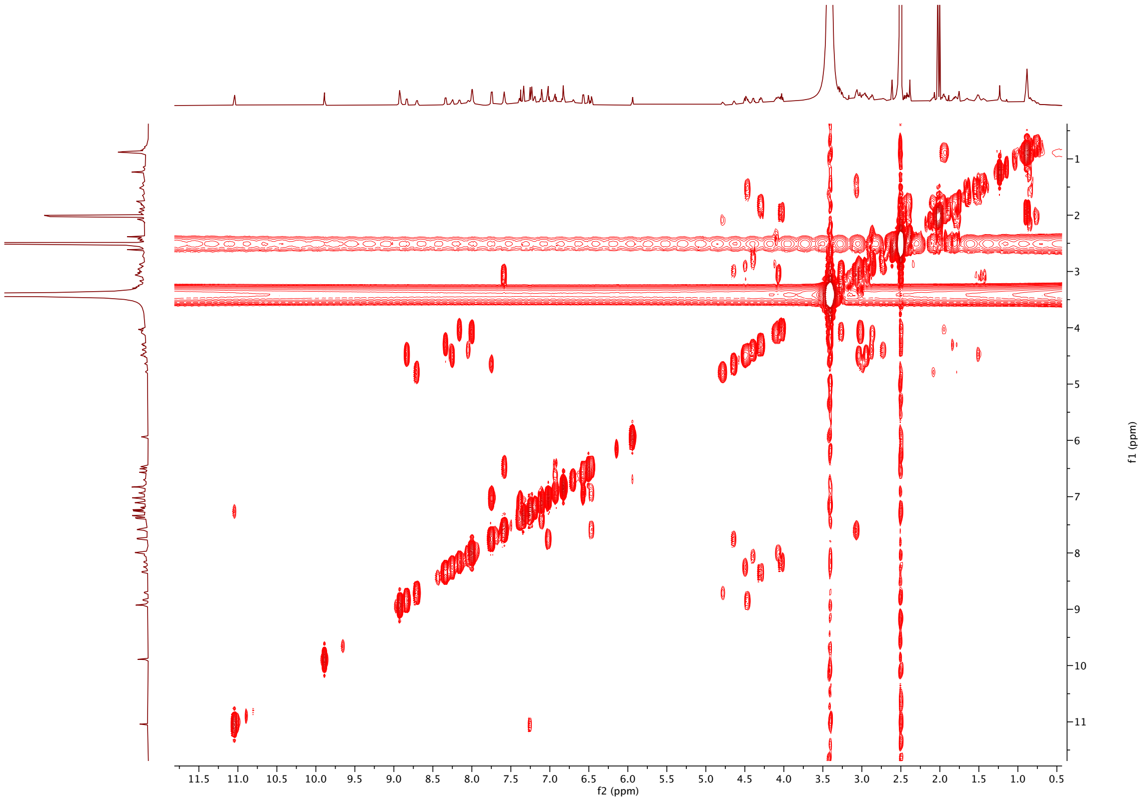


NOESY of **8** (600 MHz, DMSO-$d^6$):

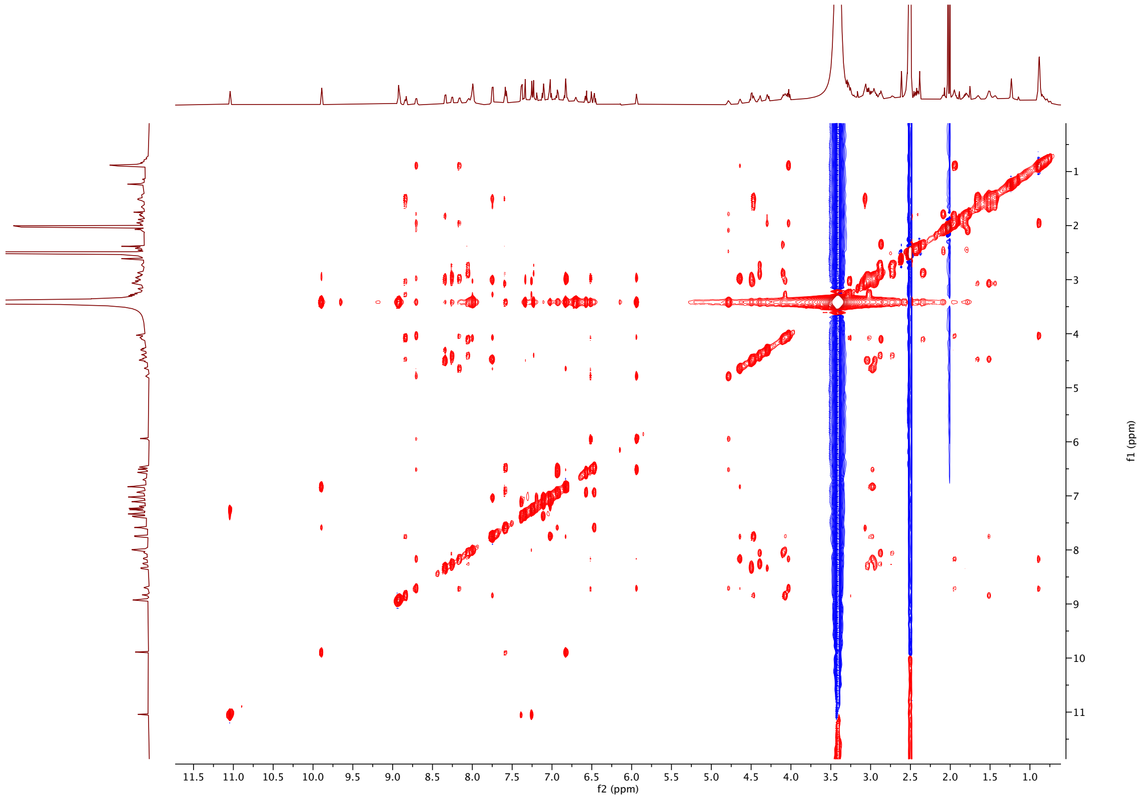

HSQC-DEPT of **8** (600 MHz, DMSO-$d^6$):

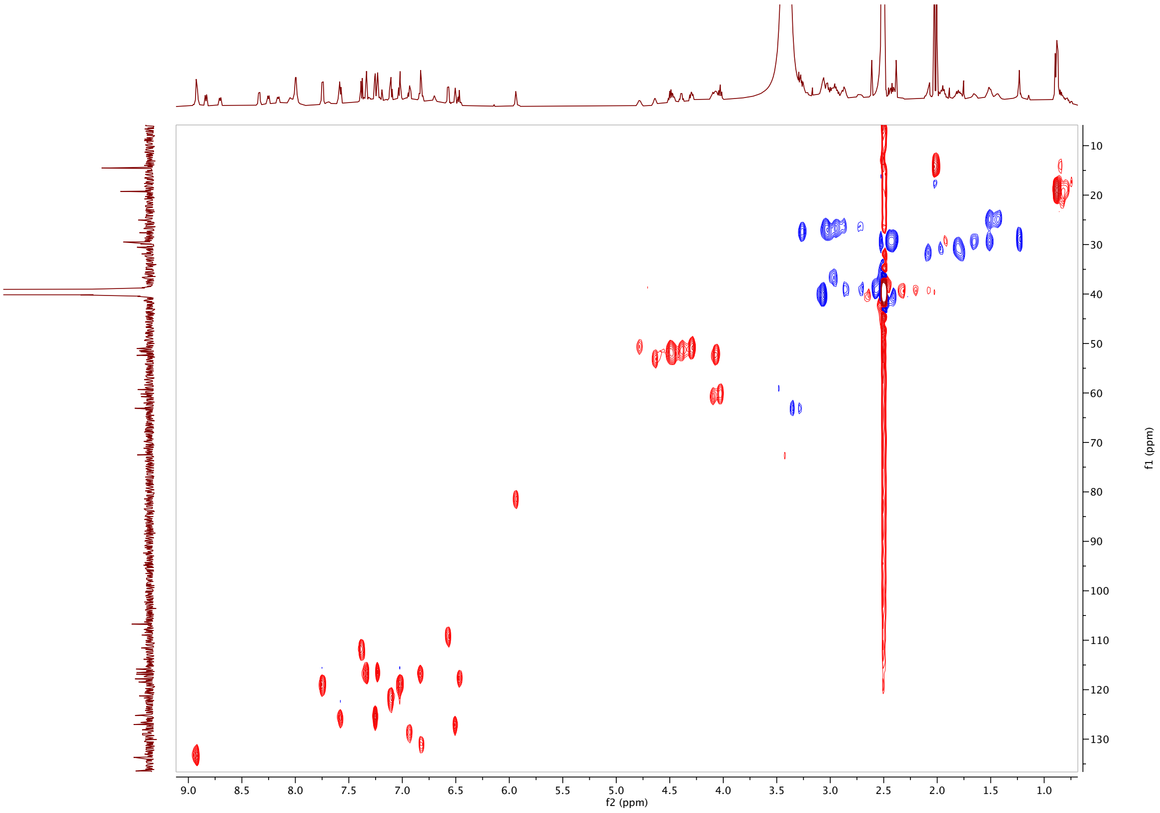


HMBC of **8** (600 MHz, DMSO-$d^6$):

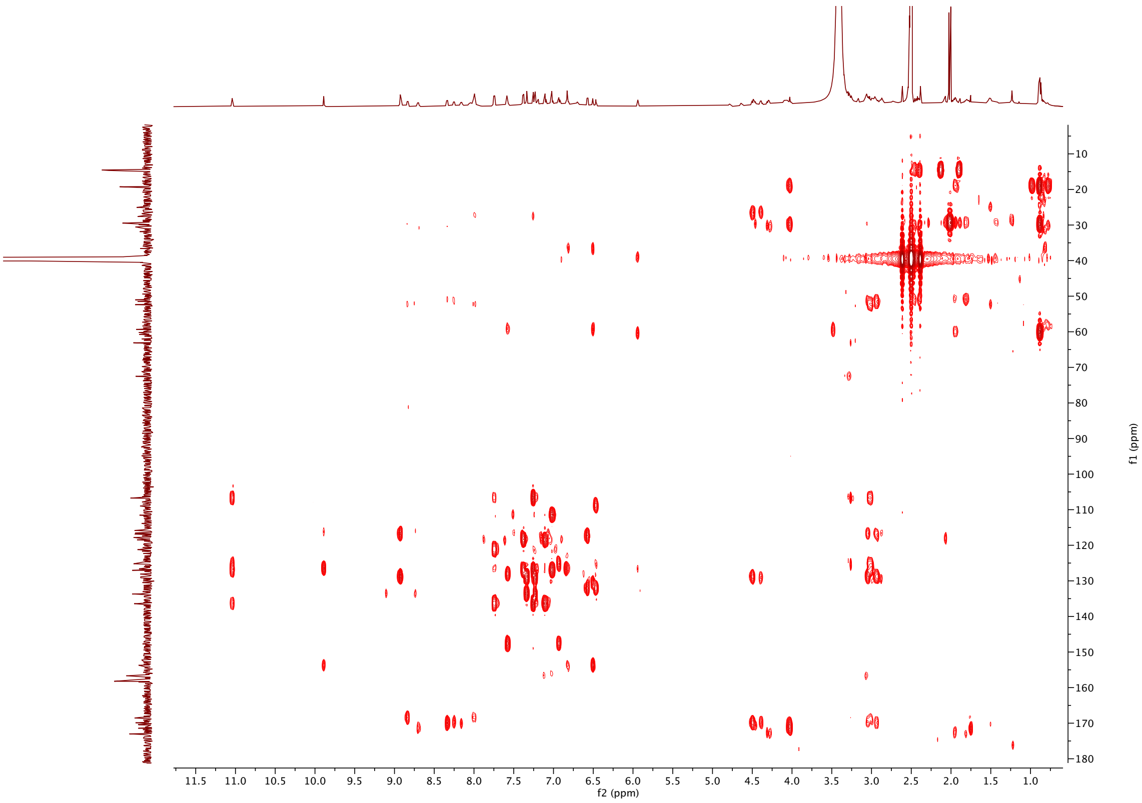

[1]H NMR of **42** (600 MHz, DMSO-*d*[6]):

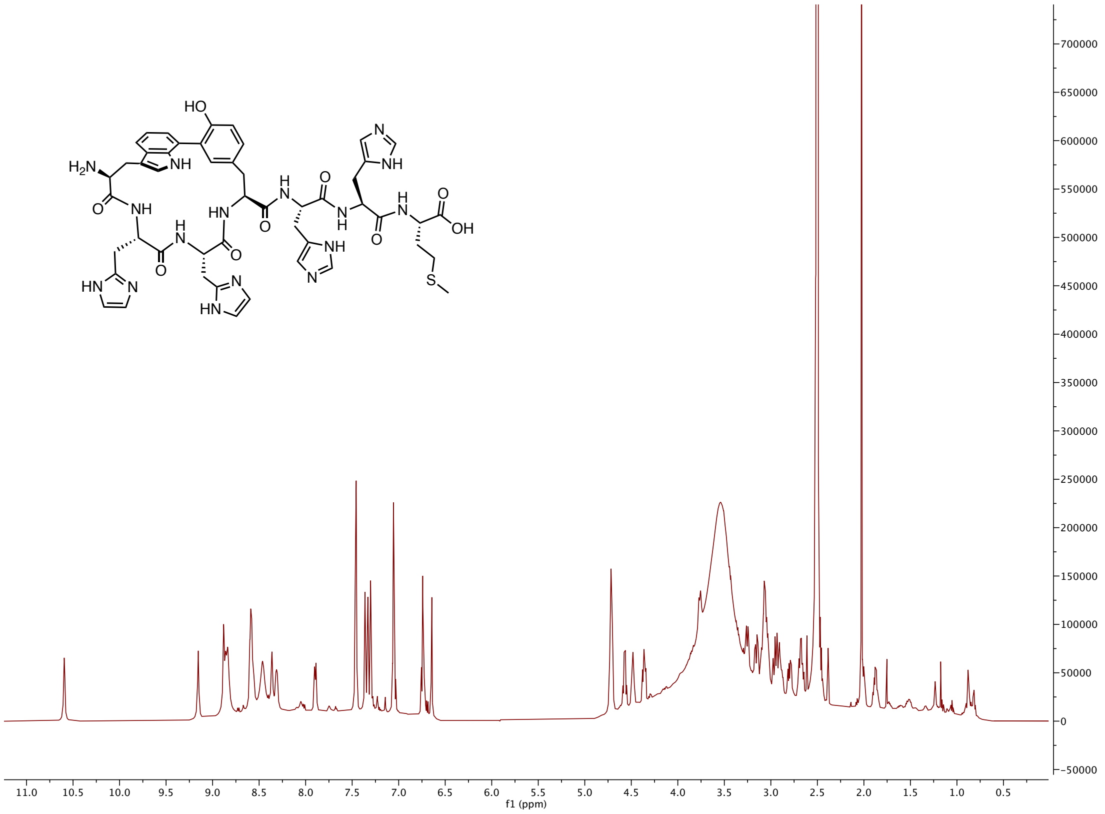


[13]C NMR of **42** (150 MHz, DMSO-*d*[6]):

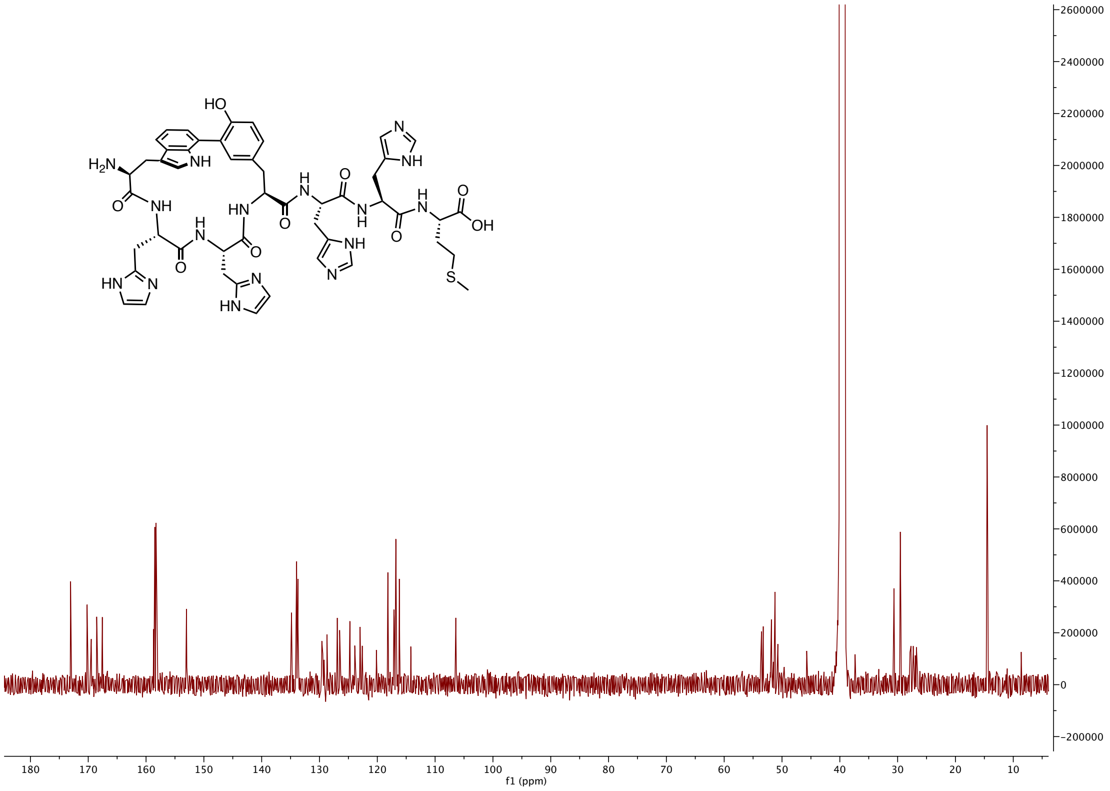

H-H COSY of **42** (600 MHz, DMSO-$d^6$):

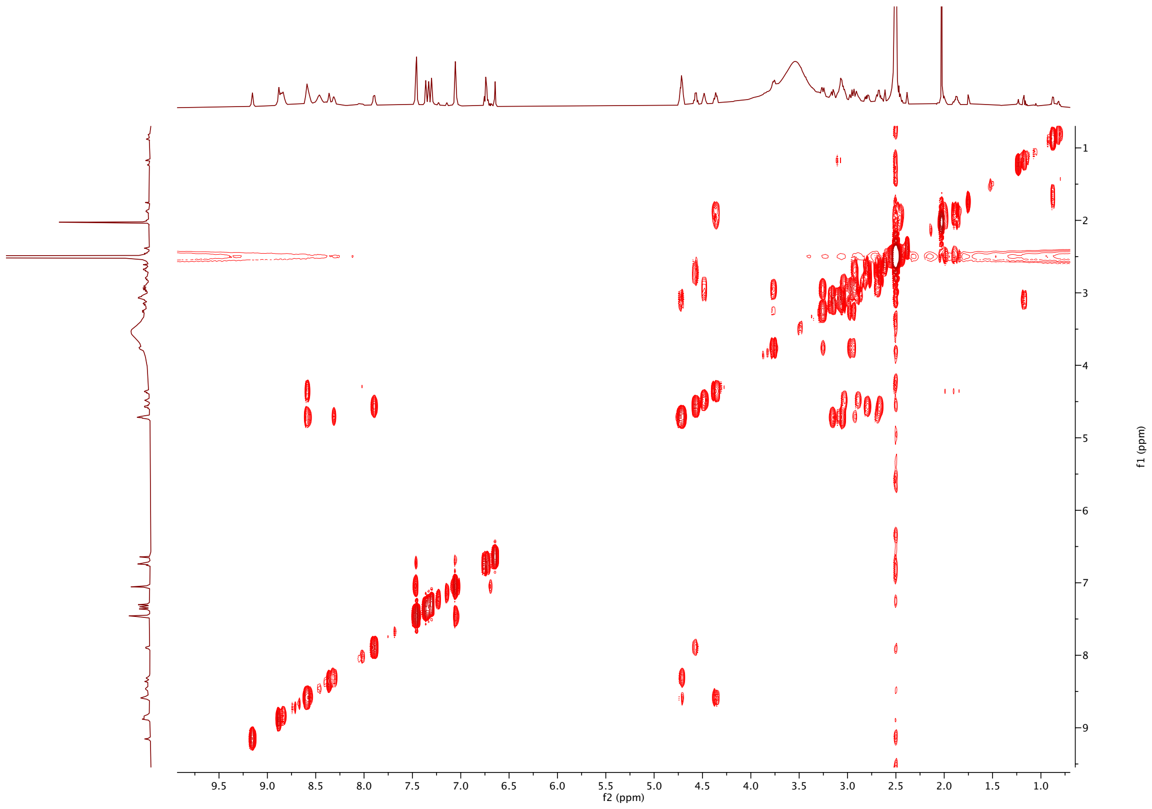


NOESY NMR of **42** (600 MHz, DMSO-$d^6$):

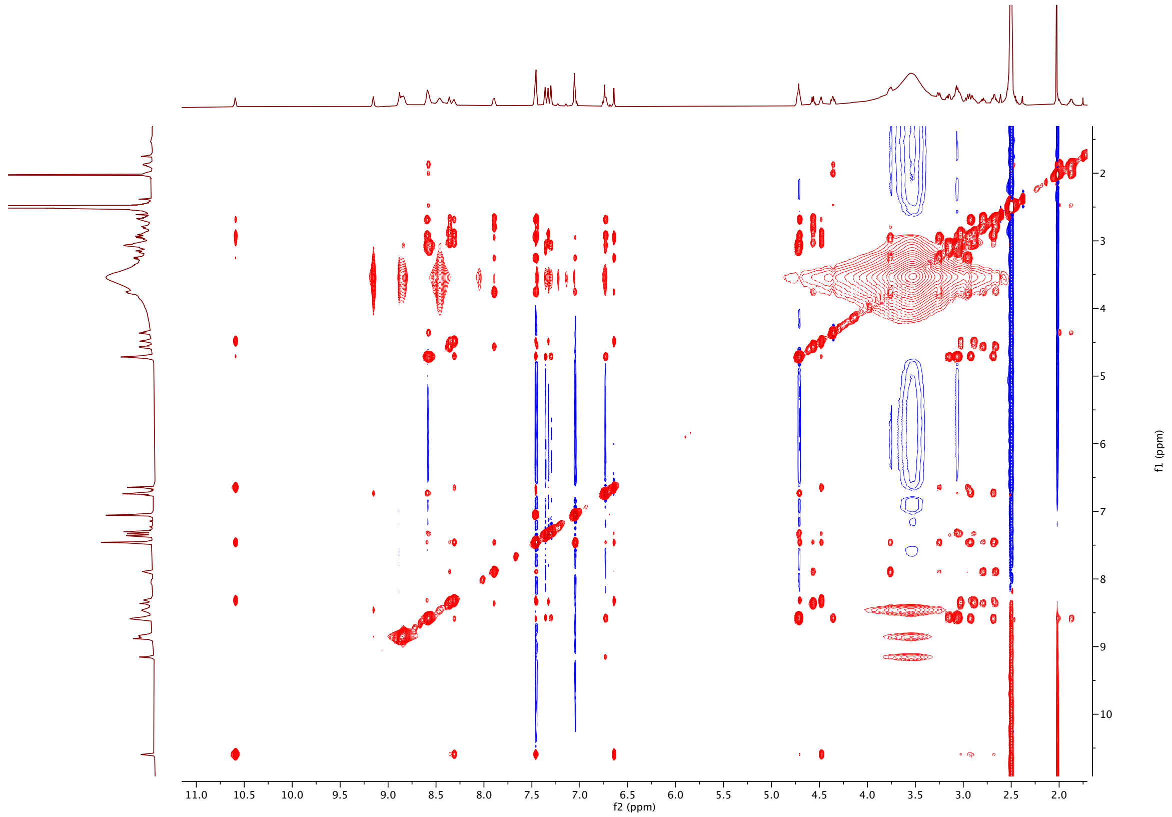

HSQC-DEPT of **42** (600 MHz, DMSO-$d^6$):

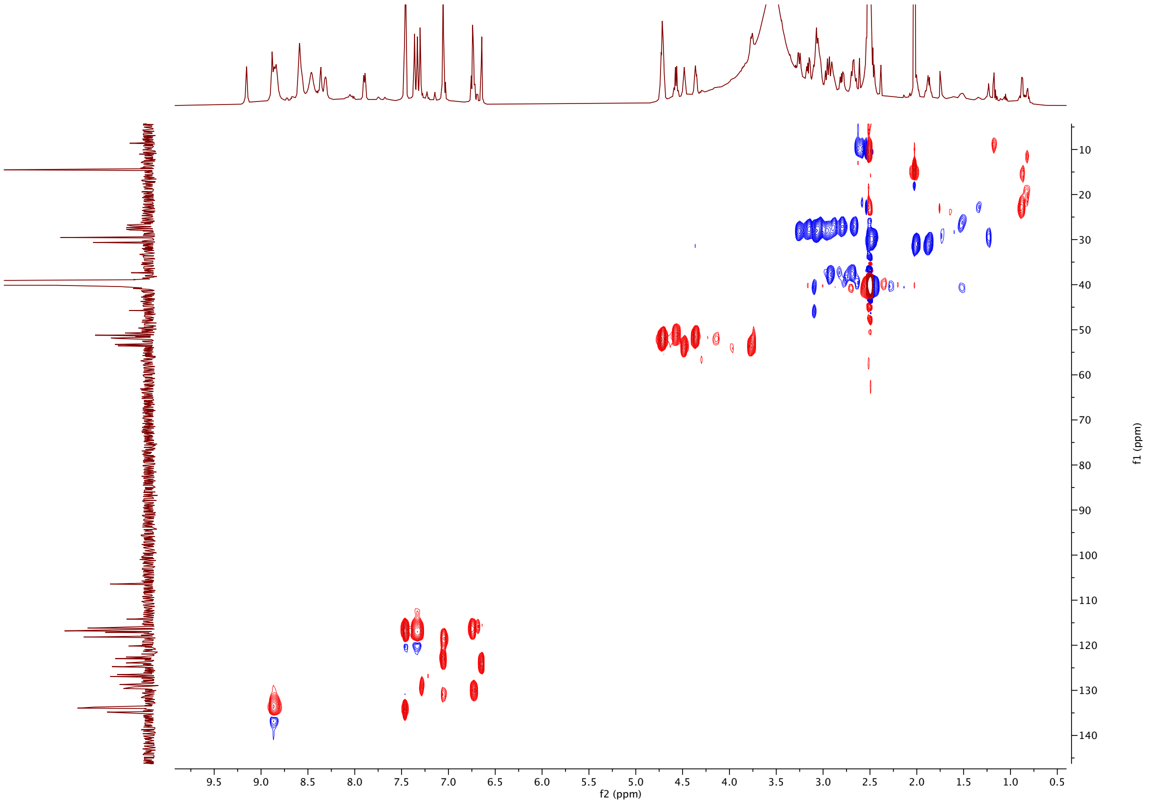


HMBC of **42** (600 MHz, DMSO-$d^6$):

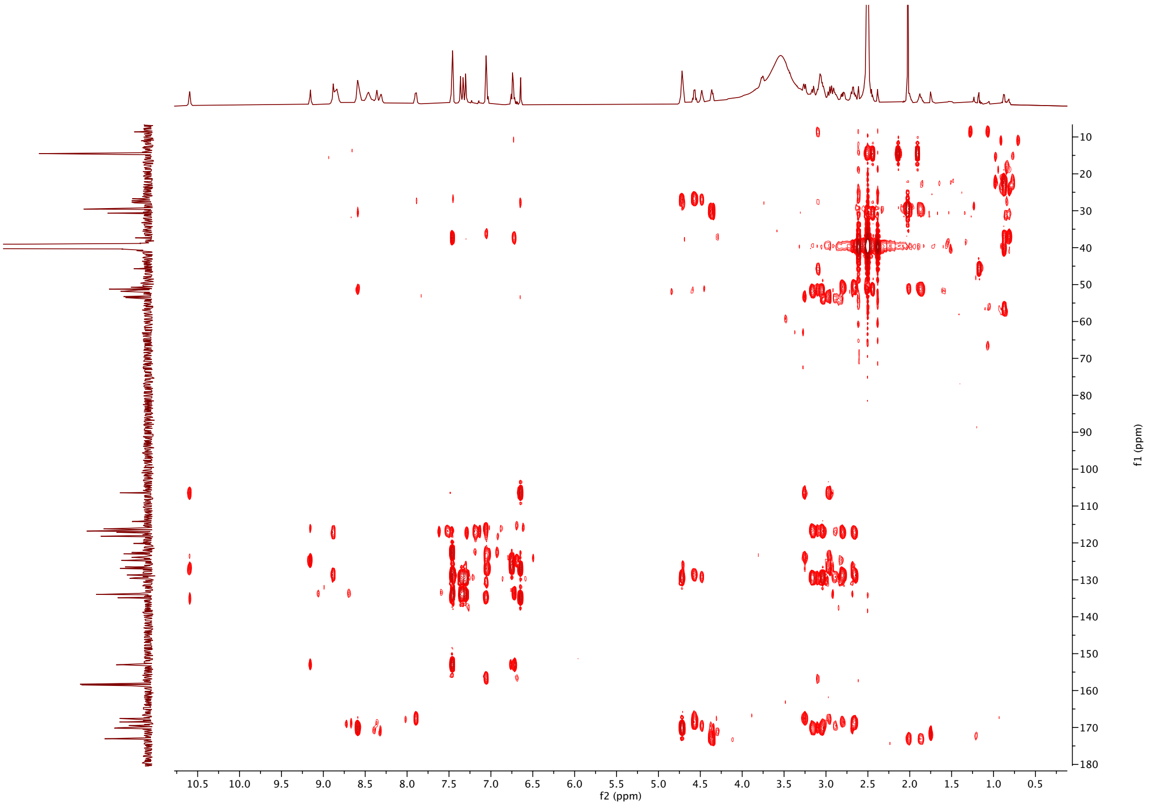

$^{1}H$ NMR of **His(OH)-DNP** (400 MHz, DMSO-$d^{6}$):

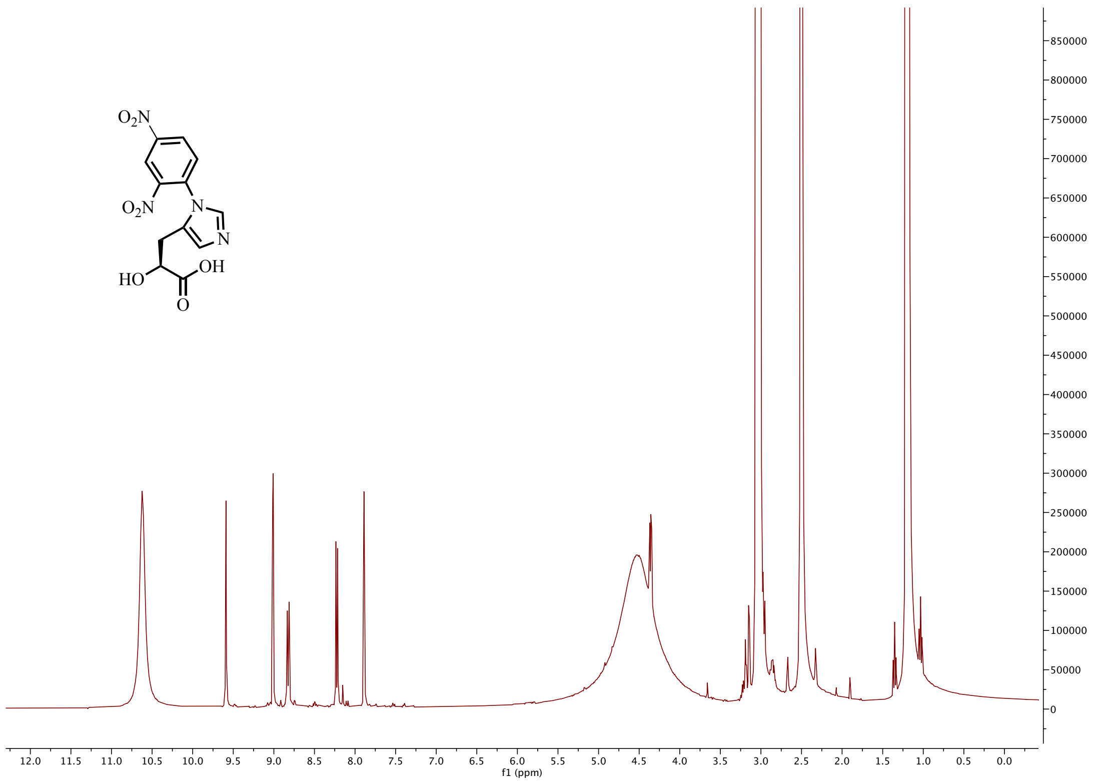


$^{13}C$ NMR of **His(OH)-DNP** (101 MHz, DMSO-$d^{6}$):

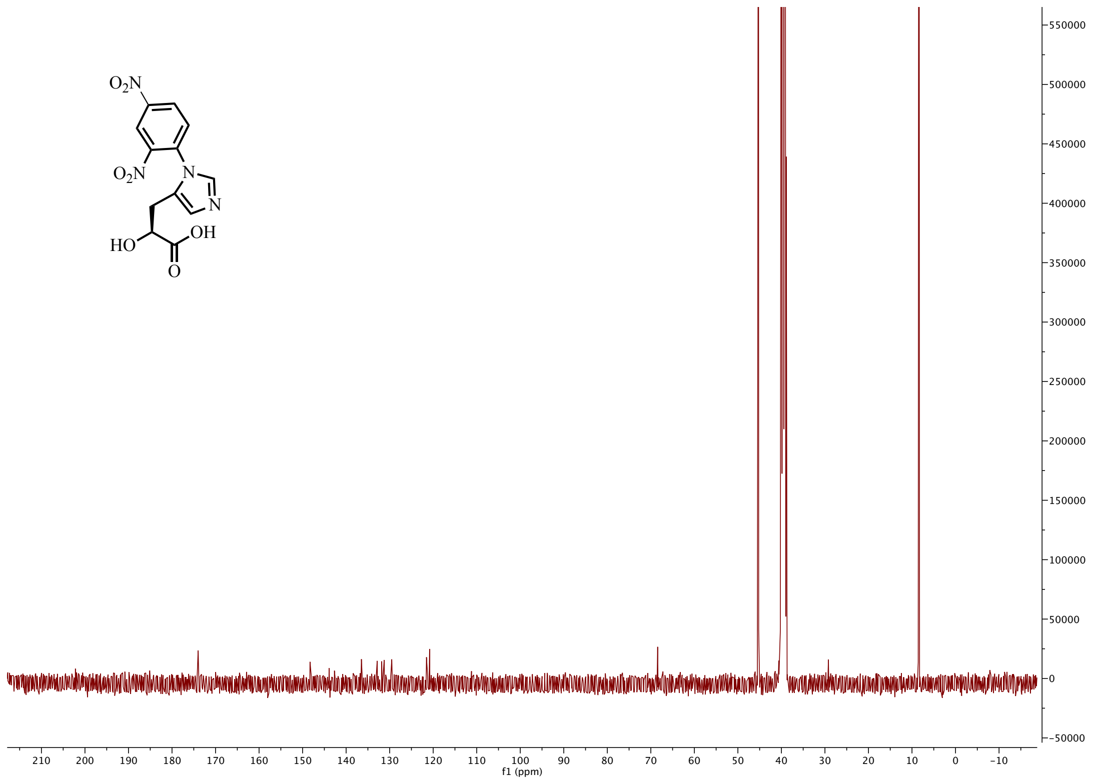
